# Essentialome-Wide Multigenerational Imaging Reveals Mechanistic Origins of Cell Growth Laws

**DOI:** 10.1101/2025.06.10.658525

**Authors:** Daniel S. Eaton, Carlos Sánchez, Luis Gutiérrez-López, Jacob Q. Shenker, Youlian Goulev, Brianna R. Watson, Véronique Henriot, Ethan C. Garner, Quincey A. Justman, Jeffrey R. Moffitt, Johan Paulsson

**Affiliations:** Department of Systems Biology, Harvard Medical School, Boston, MA 02115, USA; Program in Cellular and Molecular Medicine, Boston Children’s Hospital, Boston, MA 02115, USA; Institut Curie, Orsay, Essonne 91400, France; Department of Molecular and Cellular Biology, Harvard University, Boston, MA 02138, USA; Department of Microbiology, Blavatnik Institute, Harvard Medical School, Boston, MA 02115, USA

## Abstract

*Escherichia coli* is arguably the most thoroughly characterized organism, and yet ∼20% of its essential genes serve completely unknown functions,^1,2^ and many core quantitative physiological principles remain unexplained, including the famous nutrient ‘growth law’ where cell volume seems to depend exponentially on growth rate. Here we develop a platform for massive, multi-generational Optical Pooled Screening (OPS)^3–6^, and apply it to link image-based phenotypes to genotypes for 133,000 CRISPRi knockdowns of essential genes, tracking tens of millions of lineages and analyzing 1.6 billion cells. Our multi-dimensional dynamic phenotypes correlate exceptionally well with known gene functions, allowing us to identify many unknown roles of essential genes. Quantifying the relation between growth and cell size in turn identifies three distinct variants of the bacterial growth laws, which we explain mechanistically by discovering a new role for (p)ppGpp and SpoT as a sensor of translation elongation. Finally, we propose and systematically test an exceedingly simple, passive mechanism for the nutrient growth law, based on triggering cell division through the accumulation of a protein that, unlike ribosomes, is *not* controlled by (p)ppGpp. The resulting hyperbolic growth law fits the data from our three variants even better than the previously proposed exponential relationship, and with fewer parameters.

## Main

Time-lapse microscopy is a powerful tool for cell biology, but is often throughput-limited – finely tracking a few labeled components in a handful of genetic backgrounds, but only managing genome-level screens with great effort^7,8^. Optical Pooled Screening (OPS) promises to change that by leveraging randomly pooled rather than structurally arrayed cell libraries, and then simultaneously phenotyping and genotyping every cell under the microscope. The randomization reduces batch artifacts when comparing cells, and, by eliminating the need to keep custody of cells in parallel compartments, approaches the physical limit of experimental miniaturization where each individual cell serves as a biological replicate. However, current OPS also suffers several limitations. Pooling cells together makes it difficult to culture and track them over multiple generations of growth. OPS has therefore focused on single snapshots or short-term dynamics in constant environments. OPS is also challenging^9^ when studying bacteria or yeast, because of their hard-to-penetrate membranes and low per cell nucleic acid abundance. If it were not for these challenges, microbes would be exceptionally well suited for multigenerational OPS thanks to their small physical footprint, short generation times and the relative ease of building large, pooled libraries.

Here we present an optical screening platform that combines and extends the key benefits of both arrayed and pooled screens, allowing long-term tracking and subsequent genotypic identification of microbes. Specifically, we achieve large-scale OPS for microbes in a microfluidic cell culture format based on so-called “mother machine”^4,10–20^ designs. Cells are loaded as random pools (Fig. 1a) that self-organize into isogenic lineages by single-file growth in trenches, diffusively fed from continuously replenished media that also serves to evacuate surplus offspring (Fig. 1b). This enables tracking of the growth of >130,000 strains per experiment over hundreds of consecutive generations^15,16^ in local environments that can be finely controlled in time and space^17,19^. It also minimizes measurement artifacts^17^, allows for retrieval of cells of interest^18^, and has many other important features^11,20,4^. Early attempts to expand OPS to this format successfully used Fluorescent In situ Hybridization (FISH) to optically read out RNA barcodes at the end of the experiment in a low-throughput format that was limited to 84 strains per experiment.^4,20,21^ We have built on this foundation by systematically optimizing several aspects of the format.

**Fig. 1:**
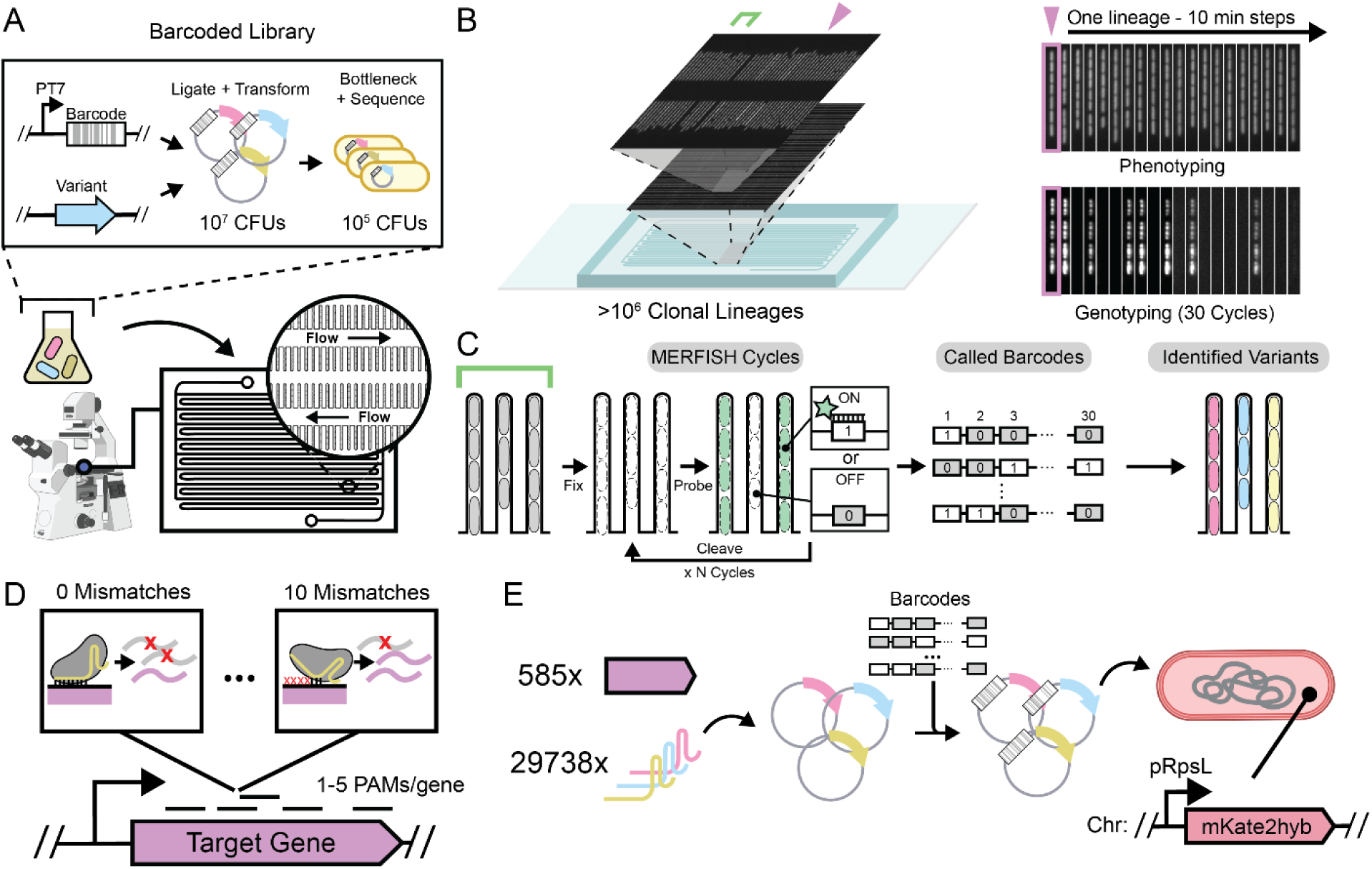
MARLIN - Multiplexed Assignment of RNA-barcoded LINeages. **a,** MARLIN libraries are constructed through combinatorial ligation of MERFISH readout sequences to genetic variants and subsequently bottlenecked to ∼10^5^ CFUs. During an experiment, libraries are loaded as a pool and cultivated on our high-throughput mother machine. **b,** In a single imaging run, our device enables continuous imaging of >1 million bacterial lineages every 10 minutes, enabling high precision measurement of growth dynamics at scale. **c,** As an endpoint, cells are fixed, and barcodes read using FISH. Dye signal is subsequently eliminated by a disulfide reduction allowing multiple cycles of sequential probing. Decoded barcode sequences are mapped back to their associated library members’ time-lapse data using a sequencing-established lookup table. **d,** Our library is designed according to a mismatch-CRISPRi strategy where gene expression is titrated between 0 and 10 mismatches between sgRNA and target. We included mismatch series for up to 5 PAM sites per gene, depending on the number available in the gene. **e,** Our essentialome-wide CRISPRi screen was designed targeting 585 essential genes in *E. coli* with 29738 sgRNAs, derived a previous set of matched sgRNAs^2^, which was barcoded and introduced into MG1655 *E. coli* containing a genomically integrated pRpsL-mKate2Hyb fluorescent reporter.

First, we developed a microfluidic architecture organizing cells to be tracked into over a million parallel lineages in a dense geometry that enables image acquisition for about ∼100,000 lineages per minute (Fig. 1b). The resulting multi-terabyte data volumes are segmented and analyzed using custom software^22^ for parallel computation on high-performance computing systems (Supplementary Methods). By doing this, we can track hundreds of millions of *Escherichia coli* cell divisions per day and determine the division phenotype.

Second, we link the observed phenotypes to genetically encoded barcodes by optically reading out the latter under the microscope. Traditional DNA barcodes of sequence length *n* can in principle separate between 4*^n^* variants, but in practice less since the number of nucleotide differences between barcodes (the Hamming distance) must be large compared to the average number of errors from sequencing. Here we instead use an expression-based strategy with *n* FISH rounds, which extends a previous FISH-based barcoding method for single-time-point OPS.^3^ Specifically, we use labeled probes against unique RNA targets, asking if each target is expressed (“1”) or not expressed (“0”) in a given cell lineage (Fig. 1c). By using three spectrally distinct probes simultaneously, we can in principle separate between 2^3*n*^=8*^n^* variants in *n* rounds of FISH. In practice the number is lower since we bottleneck our libraries to statistically ensure that the number of different FISH targets between barcodes (here termed the Herring distance) is typically much larger than the number of FISH errors.^3^ We have heavily optimized this process, placing our barcodes under strong, inducible expression post-phenotyping, extensively screening FISH probes for minimal adsorption into the device polymer, and adapting MERFISH chemistry^3,23,24^ to speed up each probe cycle – typically running 10 rounds of three-color imaging in 14 hours (Methods). To test accuracy, we assayed a pooled library of 120,000 different strains, each expressing a unique 30-bit barcode together with an intact or inactivated copy of a *gfp* gene, with fluorescence providing ground truth for the *gfp* genotype (Extended Data Fig. 2b,c). This revealed that our FISH barcode read-outs were >99.8% accurate, in a process that was straightforward to automate and costs around $180 per run (Supplementary Table 6).

Finally, we link the barcodes to the genetic regions of interest by sequencing both elements on the same read. This requires a challenging combination of long-read sequencing (>1kb) and a low error rate to capture even single point mutations. However, our barcodes help here as well: with typical Herring distances of 3-4 from the nearest barcode, and single-bit Herring distances corresponding to Hamming distances of ∼10bp, the whole-barcode Hamming distances are generally >30bp, providing a clear barcode consensus for every read. Grouping the reads by the barcodes then allows us to pinpoint other mutations with great accuracy, e.g., detecting the 2bp substitution distinguishing our *gfp* variants with an overall total accuracy of >99.8%, with as few as 20 grouped reads from an Oxford Nanopore run. Higher accuracy is likely to be possible with recent advances in sequencing chemistries (Extended Data Fig. 2a and Supplementary Methods).

This ‘big FISH’ platform – which we call MARLIN for <u>M</u>ultiplexed <u>A</u>ssignment of <u>R</u>NA-barcoded <u>LIN</u>eages – allows us to precisely track >100,000 strains at the cellular and subcellular levels, with each strain represented by thousands of individual cells studied for dozens of generations of growth in controllable environments, and then precisely calling point mutations in a genetic region of interest for every cell. This method can be applied to a wide range of genotype-to-phenotype mapping tasks ranging from mutational scans of DNA regulatory elements to systematic perturbations of gene networks. Here we will use it to identify unknown gene functions, and to mechanistically explain emergent physiological patterns and growth laws.

### Systematic Measurement of Cell Physiology in an Essentialome-wide Knockdown Library

In our first application of MARLIN, we chose to investigate the contribution of essential genes to cell growth and morphology in *Escherichia coli*, using a perturbative approach. Because loss of essential function by definition leads to nonviable cells, we decided to use CRISPRi-based transcriptional knockdowns, which we reasoned would allow us to measure phenotypes over time from the onset of knockdown, circumventing the need for cultivation to steady state. This also minimizes selection for compensatory mutations in our uninduced pre-cultures. To further maximize our retention of viable cells, we designed our CRISPRi library to include systematically “mismatched” sgRNAs with varying complementarity to their targets, manifesting as varying degrees of non-lethal phenotypes.^25–27^

We designed a CRISPRi library of around 134,000 strains, each with a unique barcode, covering 29,738 different sgRNAs that targeted 585 essential genes (Fig. 1d,e). Each barcode corresponds to a separate library transformant and each sgRNA is typically represented by 4-5 different clonal lineages (Extended Data Fig. 1f), to allow us to exclude any artifacts from random mutations in lineage founders. We then performed four replicate perturbation experiments, imaging growing lineages for roughly 16 consecutive generations and inducing CRISPRi after 3-4 generations to deplete essential gene products. Our basic technology would allow us to image orders of magnitude more generations,^15^ but 16 was chosen to allow genotyping each strain before all cells in a lineage died from essential gene depletion. In total, ∼2 million lineages passed quality controls over the four replicates. This aggregate MARLIN dataset contained at least 200 lineages for 581 of 585 essential gene targets, with ∼3,400 lineages observed on average per essential gene (Extended Data Fig. 1g,h).

To capture the global effects of knocking down essential genes, we extracted six phenotypic parameters (Fig. 2a and Methods) from each of the 1.6 billion cells analyzed in the MARLIN dataset – cell length (L), width (W), interdivision time (τ) and the related instantaneous single-cell growth rate (λ), error in division septum placement normalized by cell length (L*_s_*) and pRpsL fluorescence reporter intensity (I_rpsL_) as a measure of ribosome synthesis regulation. As a control, we imaged growing cells in the absence of CRISPRi for over 40 generations, showing that all measured parameters exhibit stable means and variances over the course of imaging (Extended Data Fig. 3a).

**Fig. 2:**
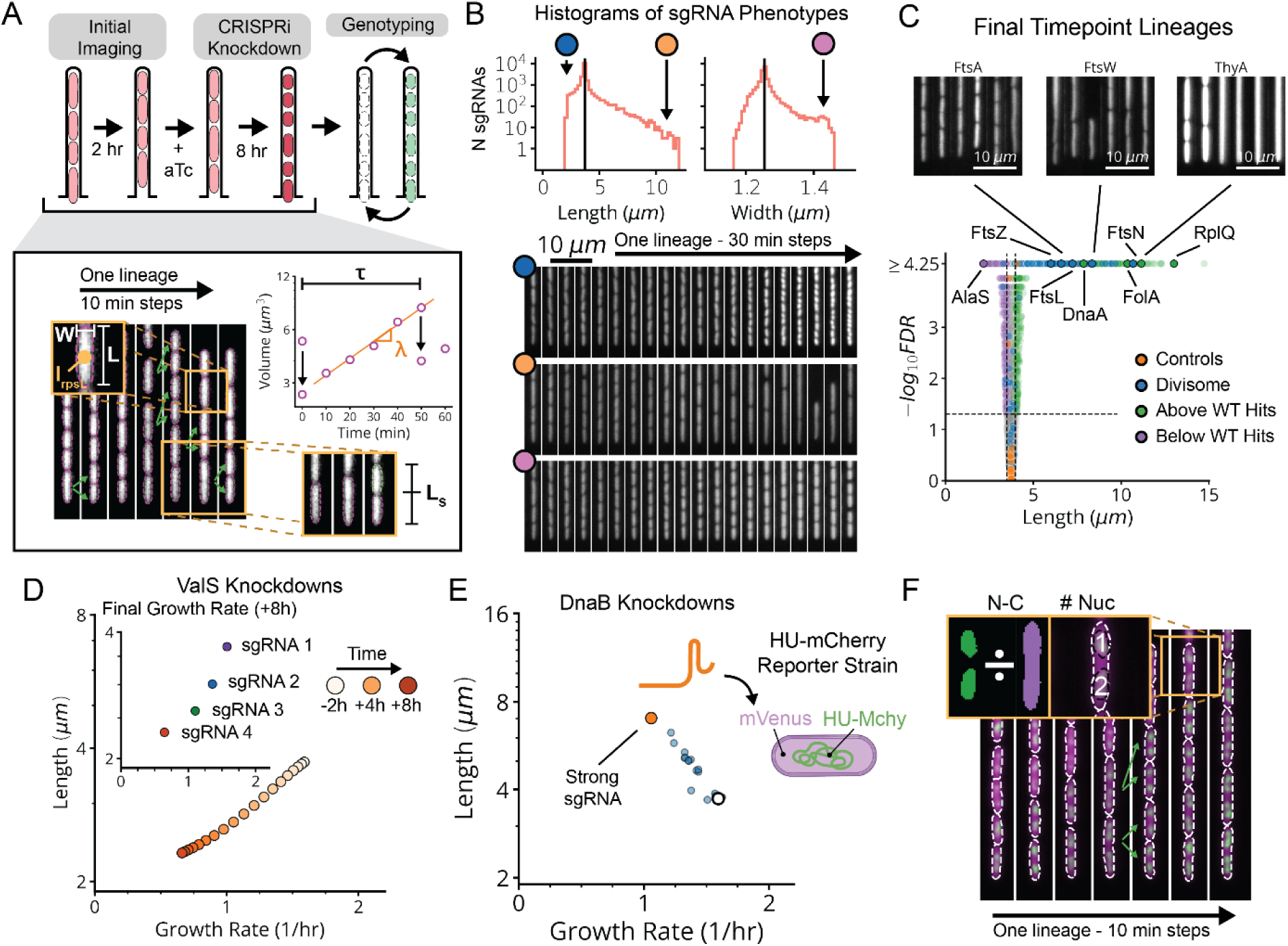
Essentialome-wide Mismatch-CRISPRi with Cell Cycle Measurements. **a,** Top, our CRISPRi library was monitored for over 2 hours of normal growth followed by 8 hours of CRISPRi induction before being genotyped. Bottom, kymograph of an example lineage from our library. Arrows show identified division events. Top-left zoom-in shows that we quantify length (L), width (W) and pRpsL reporter intensity (I_rpsL_) from single snapshots. Bottom-right zoom-in shows our terminal or “mother” cell growing over time, with its cell volume over time indicated on the graph above. From these data, we measure both the instantaneous growth rate (λ) and the interdivision time (τ), as well as the absolute error in septum placement (L_S_). **b,** Top, log histograms of sgRNA variant average phenotypes, between 5-8 hours of CRISPRi induction. Black line indicates the mean of control sgRNA mean phenotypes. Bottom, representative kymographs of lineages containing sgRNA variants with the indicated mean phenotypes. **c,** Volcano plot of cell lengths measured between 5-8 hours post-induction, for each sgRNA in the library. The horizontal line corresponds to FDR=0.05 and the vertical lines indicate two standard deviations around the distribution of control sgRNAs. Sample lineages are shown of representative and control strains, after 8 hours of CRISPRi induction. **d,** Length vs. growth rate plot of a single sgRNA targeting ValS over time. Inset indicates the growth-length scaling of representative ValS sgRNAs after 8 hours of CRISPRi induction. **e,** Left, length vs. growth rate plot for sgRNAs targeting DnaB after 8 hours of CRISPRi induction. The white circle indicates the mean length and growth rate of control strains. A strong sgRNA (orange dot) is selected from the set of DnaB-targeting sgRNAs and pooled with representative sgRNAs targeting all other genes in the library. Right, these sgRNAs were introduced into a strain bearing chromosomally integrated HU-mCherry and pRpsL-mVenus markers. **f,** Kymograph of an example lineage from our HU-mCherry library (lDE26, Extended Data Table 1). Top-left zoom-in shows that we may quantify the nucleoid to cell area ratio (NC) and the number of nucleoids (N_n_) in each cell.

Following CRISPRi induction, our library variants exhibited great phenotypic diversity in every measured parameter across the library (Fig. 2b and Extended Data Fig. 5), producing a spectrum of physiological impacts across different knockdown strengths (Fig. 2d and Extended Data Fig. 4b). Quantities were reproducible on repeated measurements of the same sgRNA, across regions of the same microfluidic chip, in replicate experiments (Extended Data Fig. 3d), and in select isolates moved into individual non-barcoded strains (Extended Data Fig. 3e). We also verified that well-known biology is captured by MARLIN. For example, knockdowns of members of the Divisome complex, responsible for carrying out cell division, showed significantly increased cell length (Fig. 2c, blue dots). Characteristic phenotypes were also obtained for depletions targeting the Elongasome, the MinCDE system and the ParCE system (Extended Data Fig. 5b-d).

For a subset of sgRNAs, we further used a DNA marker and higher imaging resolution to better distinguish division and replication defects. Specifically, we selected one or two sgRNAs per gene that had a strong effect on cell physiology and introduced these into a strain bearing a HU-mCherry fluorescent fusion protein that localizes to the nucleoid (Fig. 2e,f). This allowed us to quantify both the ratio of the nucleoid to cell area (NC ratio) and the number of nucleoids per cell (Fig. 2f). A reduced NC ratio appears to serve as a proxy for the failure to replicate, as exhibited by a DnaA knockdown (Extended Data Fig. 6a). Further, an elevation in the number of nucleoids indicates division failure: inhibition of division results in the formation of long, poly-nucleated cell filaments, as seen in FtsN knockdowns (Extended Data Fig. 6b).

Finally, for all genes we performed multiparametric clustering of the phenotypes observed.^4,8,28^ To retain information encoded over the time-course of CRISPRi knockdown, we aggregated measurements for each sgRNA into a multi-dimensional timeseries of Z-scores corresponding to each measured parameter (Fig. 3a and Supplemental Methods). We then generated clusters and a 2-dimensional UMAP based on a distance metric (softDTW)^29^ between those timeseries. Each cluster was assessed for statistical stability, measured by comparing cluster membership across jackknife resamples of the same data (Extended Data Fig. 7c). Finally, we collapsed redundant information from different sgRNAs targeting the same gene (Fig. 3b), which usually appear in the same cluster. As an initial validation of this approach, we found that members of multi-protein complexes clustered together with their complex partners (Fig. 3c,d) and reproduced phenotypes observed in previous studies.^30–32^

**Fig. 3:**
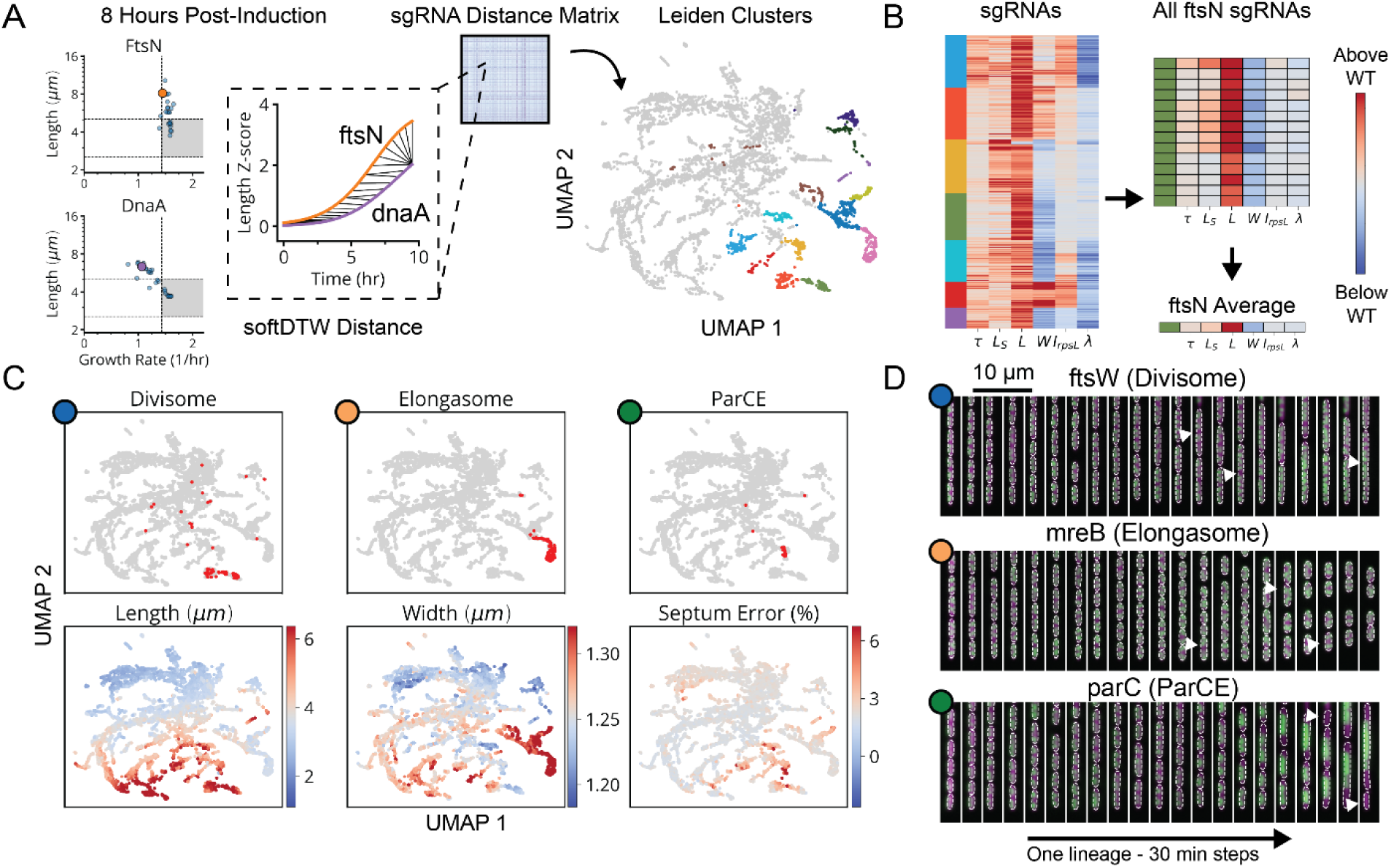
Clustering Reveals Phenotypic Signatures of Essential Processes. **a,** Left, each parameter, measured over time, is converted to a 20-point Z-score timeseries (Methods) and averaged over trenches belonging to the same sgRNA variant to produce an average timeseries. We subsequently filter out variants with weak phenotypes by excluding those which do not exhibit a Z-score > 1 for any timepoint, along any dimension (grey area). Right, distances between timeseries data for sgRNA variants are established using a softDTW distance metric, which produces a sgRNA vs sgRNA distance matrix that is then used to generate a 2-dimensional UMAP and clusters based on the Leiden clustering algorithm. Subset of clusters associated with a division/replication-like defect are indicated by color. **b,** Heatmap indicates average phenotypic quantities after 8 hours of CRISPRi induction for each sgRNA. Scale in Fig. 4c. sgRNA variants composing each of our clusters of interest are aggregated into single genes by taking the average of each phenotypic value. Single genes are assigned to the cluster in which the plurality of their targeting sgRNAs can be found, with two clusters assigned in the case of ties. **c,** Top, red dots correspond to sgRNAs targeting genes belonging to the indicated class, mapped onto a UMAP representation of each sgRNA variant. Bottom, endpoint values of L, W, and L_S_ for each sgRNA. Blue represents values below those of control sgRNAs and red represents values above the controls, with scale given by the right-side bar. **d,** Representative lineage kymographs from our HU-mCherry library, from the indicated classes in Fig. 3c.

These libraries and analysis tools allow us to comprehensively link systematic measurements of physiology and replication status to perturbations for essential function, laying the foundation for mechanistic insights, as we explore next.

### Systematic Dynamic Phenotyping Reveals Unknown Physiological Functions of Many Genes

Even in a well-studied model organisms such as *E. coli*, almost 20% of *essential* genes^1,2^ serve completely unknown functions. Homology-based approaches predict molecular-level properties for some of the corresponding proteins, such as enzymatic activity, but not the phenotypes or the with which cellular processes they are associated. By quantifying many single-cell properties and dynamically separating between initial responses to perturbations and secondary epistatic effects, we reasoned that our approach might map observed phenotypes to known molecular function with enough reliability and precision that unknown gene functions can be inferred.

Using the CRISPRi library above, our global map of physiology under partial loss of essential function revealed five functional supergroups for division, replication, translation, central metabolism and envelope synthesis. Strikingly, 100% of known Divisome genes (10/10) were assigned to the division supergroup, 100% of known Elongasome genes (6/6) were assigned to the envelope synthesis supergroup, and 75% of known essential Replisome genes (9/12) were assigned to the replication supergroup (Fig. 3c and Extended Data Fig. 7d). In addition, 93% of sgRNAs targeting r-proteins appeared in the translation supergroup, and 76% of sgRNAs targeting tRNA synthetases and 72% of sgRNAs targeting glycolysis/upstream glycolytic processes appeared in the central metabolism supergroup. Each supergroup further contains distinct sub-clusters reliably corresponding to specific physiological sub-functions, as follows:

The envelope synthesis supergroup includes six distinct sub-clusters, statistically stable under resampling (Extended Data Fig. 7c), that are all characterized by changes in cell width (Extended Data Fig. 8a). Based on the genes in each group, and consistent with previous data,^31,33^ increased width was associated with outer membrane (OM) or cell wall defects, while decreased width was associated with increased OM stiffness (Supplementary Note S1). Factors involved in modulating OM composition were found in both wide cells (*lpxABDH, lptAG, yejM*) and narrow cells (*waaC, gmhA, rfaE, ftsH*), implicating the OM as a major determinant of cell width homeostasis. We also identified a gene of previously unknown function, *ykgE*, that tightly co-clustered with the high-width “Envelope II” group (Extended Data Fig. 8a), suggesting an essential role in envelope synthesis.

The cell division supergroup included three distinct and highly stable sub-clusters, all uniquely characterized by an increase in cell length without any change in growth (Fig. 4a and Extended Data Fig. 8a), consistent with previous findings.^34^ Our nucleoid imaging data identified a distinct sub-cluster (“Division I”, Fig. 4a) with clear division-specific defects and filamentous, polynucleate phenotypes (Fig. 4a,b). This sub-cluster includes four genes with previously unknown functions, *yagH*, *ybcN*, *ycgB* and *yecM* (Fig. 4a), that impaired division but exhibited no replication defect. The other two sub-clusters exhibited complex nucleoid and morphological phenotypes in addition to the division defects (Extended Data Fig. 8a). As discussed in Supplementary Note S1, genes in these sub-clusters likely impair division indirectly through either reduced lipid synthesis (“Division II” cluster) or dysregulated division localization (“Septum Error” cluster). We identify several genes with previously unknown functions (*yeiH*, *yfaU*, and *yhcN*) that are part of these subclusters, implicating them as indirect but essential division factors.

**Fig. 4:**
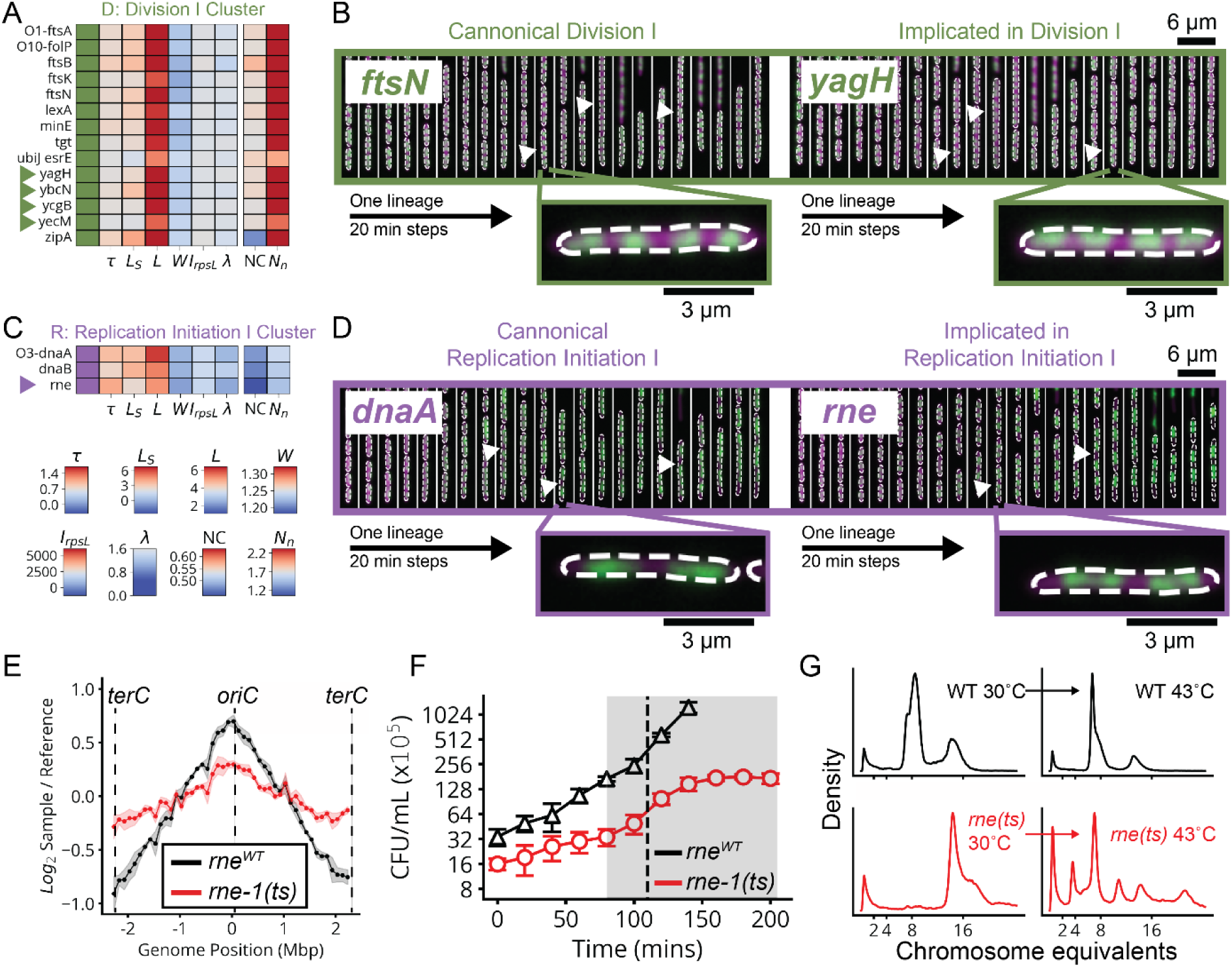
Clusters Implicate Unknown Genes in Division, Replication and Envelope Integrity. **a,** Heatmap of cluster phenotypes for the “Division I” cluster, belonging to the division “D” supergroup. Arrows indicate genes mentioned in the text. The leftmost column color denotes cluster membership, right columns display the average value of each quantity at the end of the experiment, for each gene. Same scale as in Fig. 3e. Entries beginning with O denote polycistronic operons, where the subsequent gene name indicates the last gene in a consecutive block all belonging to the indicated cluster. See Extended Data Fig. 8 for all curated clusters. **b,** Representative kymographs of genes from the “Division I” cluster, with the cytoplasm in magenta and nucleoid in green, as in the diagram in Fig. 3f. Arrows indicate clear manifestations of the cluster-associated phenotype. Left, kymograph of a *ftsN* knockdown, canonically associated with division. Right, kymograph of a *yagH* knockdown, not previously characterized as division-defective. **c,** Top, heatmap of cluster phenotypes for the “Replication Initiation I” cluster, belonging to the replication “R” supergroup. Same format as in Fig. 4a. Bottom, linear scale for variables in Fig. 3a,c, with red indicating quantities above the EV controls while blue indicates quantities below the controls. **d,** Representative kymographs of genes from the “Replication Initiation I” cluster, same as in Fig. 4c. Left, kymograph of a *dnaA* knockdown, canonically associated with replication initiation. Right, kymograph of a *rne* knockdown, not previously characterized as initiation-defective. **e,** Marker frequencies based on deep gRNA sequencing for 100kb bins across the *E. coli* genome, with the reference coordinate set to *oriC*. Frequencies normalized to reference measurements from non-replicating cells grown to stationary phase in MBM+0.2% Succinate. Measurements taken after shifting either *rne^WT^* (black) or *rne-1(ts)* (red) cells from 30 °C to 43 °C for 2 hours, with shaded area indicating ±2σ_SEM_ based on three biological replicates. **f,** CFU/mL over time for *rne^WT^* (triangles) and *rne-1(ts)* (circles) cells during a shift from 30 °C to 43 °C occurring at 80 minutes (red shading). The vertical dotted line indicates 30 minutes post-shift, the time at which post-treatment samples were taken for replication runout, dividing 1-2 times in this period. **g,** Flow cytometric measurement of PicoGreen-stained DNA before (left) and after (right) replication runout for cells grown first at 30 °C and shifted to 43 °C for 30 minutes, in strains bearing either *rne^WT^* (black) or *rne-1(ts)* (red) alleles. *rne-1(ts)* cells drop from 16 to 4-8 oriC copies after 30 minutes at 43 °C, indicating that initiation immediately halts on temperature shift and the original 16 oriC have simply partitioned into either 2 or 4 descendent cells.

The DNA replication supergroup also exhibited an increase in cell length, but unlike the division supergroup also exhibited a decreased growth rate, consistent with recent reports.^35^. It contained four distinct sub-clusters (Fig. 4c and Extended Data Fig. 8a) distinguished from each other by cell width and specific replication sub-functions: replication initiation, dTTP synthesis, or the Replisome (Supplementary Note S1). Genes with a direct role in replication exhibited a decrease in the NC ratio, consistent with their function, while those that indirectly affect replication via transcription exhibit an increase in the NC ratio, as has also been observed when treating cells with transcription-inhibiting drugs.^36,37^ This supergroup includes the gene of previously unknown function *yejF*. This essential gene shares characteristics of both the replication and envelope-related clusters (Extended Data Fig. 8a) and exhibits an expanded nucleoid signature suggestive of a role in transcription.

We further found a previously unappreciated player for replication initiation – a model molecular process of *E. coli* that has been extensively characterized both *in vivo* and *in vitro*. Three core factors, DnaA, DnaB and DnaC, participate in opening the replication bubble at the origin of replication (*oriC*).^38^ Strikingly, these three factors formed their own distinct cluster with only one other gene present: the single-strand endoribonuclease RNase E, encoded by *rne*, which was not known to play a role in this process (Fig. 4c,d). Interestingly, RNase E does not co-cluster with either of the 3’ to 5’ ssRNA exonucleases RNase II (*rnb*) and PNPase (*pnp*) (Extended Data Fig. 8e), indicating that its phenotype does not result from generic loss of mRNA degrading activity. To test whether RNase E activity is required for replication initiation, we used the fact that halting initiation reduces ploidy near *oriC*, using a temperature-sensitive RNase E allele and deep sequencing to confirm that the *oriC* copy numbers dropped after two hours of RNase E inactivation (Fig. 4e and Extended Data Fig. 10b). A replication runout assay (Fig. 4f-g and Extended Data Fig. 10c-e) in turn demonstrated immediate loss of initiation activity upon RNase E inactivation. RNase E is thus indeed required for the initiation of DNA replication *in vivo*, surprisingly demonstrating a new essential function for an exceptionally well-studied gene in one of the most central processes in *E. coli*.

Finally, the translation and central metabolism supergroups were each composed of highly interconnected clusters (Extended Data Fig. 9a,b), with lower statistical stability (Extended Data Fig. 7c) that populate a phenotypic continuum across these groups (Extended Data Fig. 7a). Though deeper phenotyping will likely reveal finer physiological distinctions in the future, many clusters were already enriched for specifically co-functional genes (Extended Data Fig. 9c). We also observed that the knockdowns targeting the tRNA synthetases for histidine and leucine (*hisS* and *leuS*) had unique phenotypes and occupied single-gene clusters (Extended Data Fig. 7e). In addition to tRNA synthetases (Extended Data Fig. 9b, orange), we found genes involved in amino acid and fatty acid synthesis, as well as glycolysis, within the central metabolism supergroup. We also found four essential genes of unknown function within this group – *ysaA* and *yfjT* (Extended Data Fig. 9d), as well as *ybcF* and *ylbF* that are consecutive genes on a putative catabolic operon,^39^ suggesting that this uncharacterized metabolism contributes significantly to overall growth.

Only five gene depletions in the whole data set did *not* cluster with others, and instead formed clusters composed of most or all redundant sgRNAs targeting the same gene. Such gene-unique phenotypes may be interesting for follow-up analysis (Extended Data Fig. 7e), especially for those which defy the typical phenotypes of co-functional genes (e.g., *hisS* and *leuS*), indicating non-canonical effects. In addition, a few genes previously assigned to specific molecular pathways reliably associated with different clusters (Extended Data Fig. 7d). For instance, knockdowns targeting fatty acid or lipid A synthesis enzymes appear in a number of clusters (Extended Data Fig. 7d) and showed a diversity of size scaling behaviors (Extended Data Fig. 8b-d). This suggests that metabolic intermediates in these pathways might have unknown regulatory activities distinct from those of end products (Supplementary Note S1).

### MARLIN Reveals Three Variants of the Nutrient and Ribosome Growth Laws

We next examined whether our approach could provide quantitative insights into the famous bacterial growth laws. Specifically, when cells are perturbed, either by external changes in conditions or by internal genetic modifications, some of the phenotypic responses follow distinct quantitative patterns. The classic ‘nutrient growth law’ of Schaechter, Maaløe, and Kjeldgaard (SMK) shows that cell size scales approximately exponentially with growth rate for a wide range of external nutrient conditions.^40^ Ribosomes in turn obey a well-known linear growth law where faster growth leads to proportionally more ribosomes.^41^ Both laws are enforced by the small molecule alarmone (p)ppGpp, which represses cell size and ribosome synthesis through regulatory circuits that are not well understood, despite playing a key role for such a central phenomenon in *E. coli*.^42,43^ We find that the systematic knockdowns above reveal three distinct variants of each growth law.

First, many of the gene knockdowns that have a clear growth defect (65 of 185) conform to classical SMK scaling, including tRNA synthetases, central metabolic enzymes and fatty acid biosynthesis genes (Fig. 5a,b and Extended Data Fig. 11b). This is consistent with our understanding of how translation affects (p)ppGpp: when the enzyme RelA binds to an uncharged tRNA at the A site of a stalled ribosome, (p)ppGpp synthesis is activated. Under conditions of low tRNA charge, there are more elongating ribosomes with uncharged tRNAs in their A-site,^44,45^ leading to increased (p)ppGpp synthesis. In a separately cloned knockdown of the tRNA synthetase PheT, we used LC-MS/MS to directly confirm that (p)ppGpp was indeed strongly elevated. We further measured the RNA/Protein ratio as a proxy for ribosomes, to confirm that ribosomes not only were correspondingly downregulated (Fig. 6b) but also depended linearly (with positive slope) on growth rate, as expected from the classical ribosome growth law (Fig. 5d).

**Fig. 5:**
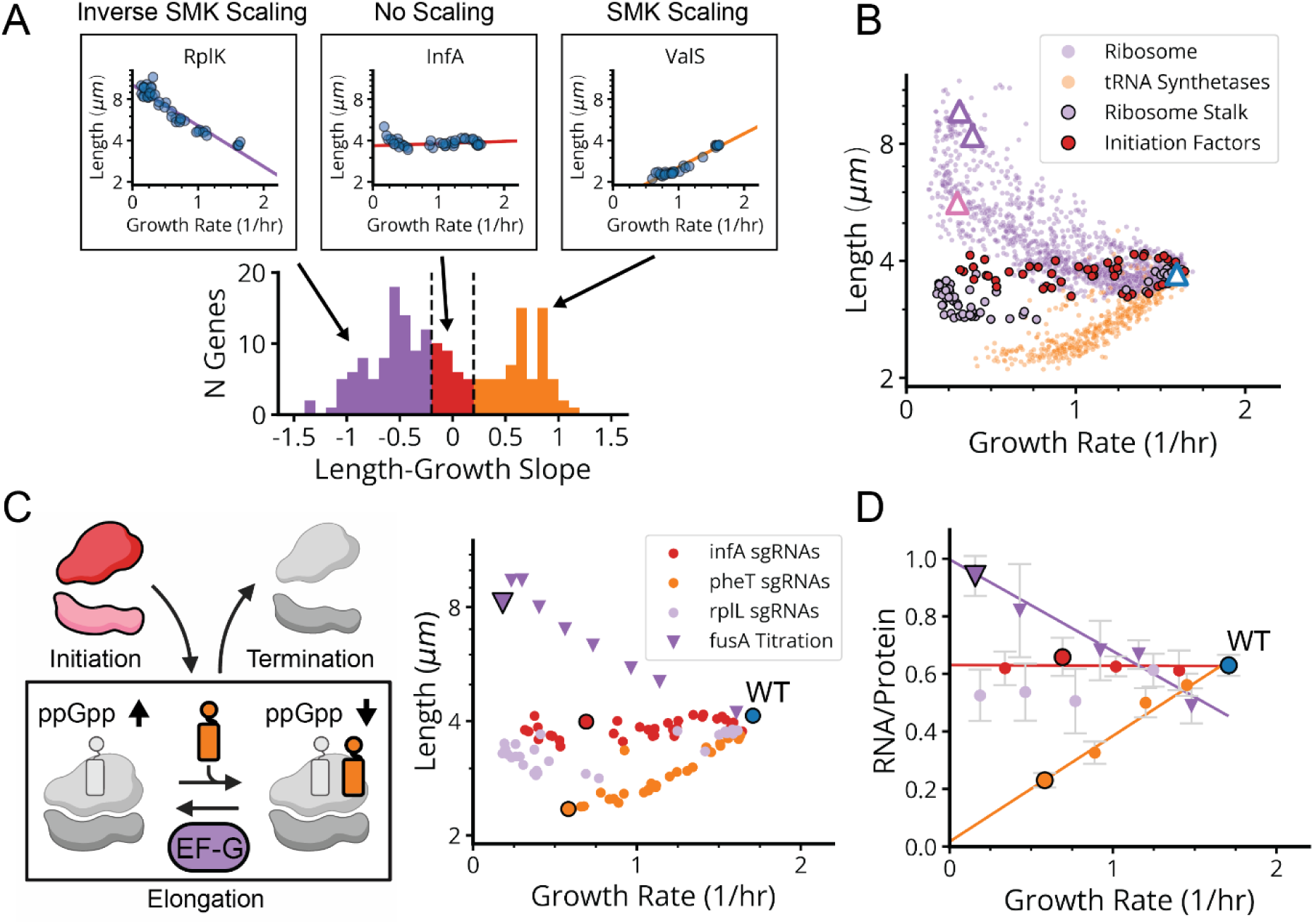
Three Size-Growth Scalings Emerge from Translation Knockdowns. **a,** Top, select log_2_ length vs. growth rate linear fits to groups of sgRNAs targeting single genes. Bottom, histogram of the log_2_ length vs. growth rate slope for genes exhibiting at least one sgRNA with a growth rate less than 0.9 doublings per hour and with sufficient points to fit a line (Extended Data Fig. 1i). Colors correspond to slopes less than −0.2 (purple), greater than 0.2 (orange) and between −0.2 and 0.2 (red). **b,** Length vs growth rate averaged between 5 and 8 hours after CRISPRi induction for variants corresponding to ribosomal proteins/rRNAs (purple circles), tRNA synthetases (orange circles), ribosome stalk proteins L7/L12 and L10 (black outlined purple circles) and initiation factors (black outlined red circles). Initiation factor InfC was omitted due to likely disruption of downstream gene expression. Note that ribosome stalk knockdowns exhibit exceptionally weak size scaling compared to other r-proteins (see Fig. 6c). Triangles correspond to points for which individual isolates were cloned in E, including 50S subunit proteins RplQ and RplA (purple), the 30S subunit protein RpsO (pink) and an EV control sgRNA (blue). **c,** Left, simplified kinetic model of translation, consisting of initiation, elongation and termination. Elongation kinetics are coarse-grained into an uncharged state (empty A-site) and a charged state (charged tRNA in the A-site). Colors indicate gene functions targeted in the corresponding knockdown strains. Right, small circles indicate average length vs average exponential growth rate for the indicated genes after 5-8 hours of CRISPRi induction (*pheT*, *infA*, *rplL*) in lDE20 (see library definitions in Extended Data Table 1). Inverted triangles indicate the same values for a strain with FusA under AHL induction (*P_AHL_-fusA*), 5-8 hours after stepping down from an AHL concentration of 1 µM to 0.2 µM, 0.16 µM, 0.13 µM, 0.1 µM, 0.07 µM, 0.05 µM, 0.03 or 0 µM. Outlined shapes represent the same values for indicated CRISPRi strains (DE428, DE830, DE831, and DE832) induced by aTc or the AHL induction strain (DE828) stepped down to 0 µM AHL. **d,** Average RNA/Protein ratio (n=5-6) after 6 hours of CRISPRi induction or removal of AHL vs average exponential growth rate from Fig. 5c (for CRISPRi strains) or measured in bulk (for the AHL strain). Final AHL concentrations used for bulk experiments were 1 µM, 0.05 µM, 0.03 µM, 0.01 µM, or 0 µM. Outlined shapes indicate the same strains and knockdown conditions as in the outlined shapes in Fig. 5c and Fig. 6b. Error bars denote 2σ_SEM_ range. Lines indicate linear fits for the *pheT* (orange), *infA* (red), and *fusA* (purple) points.

**Fig. 6:**
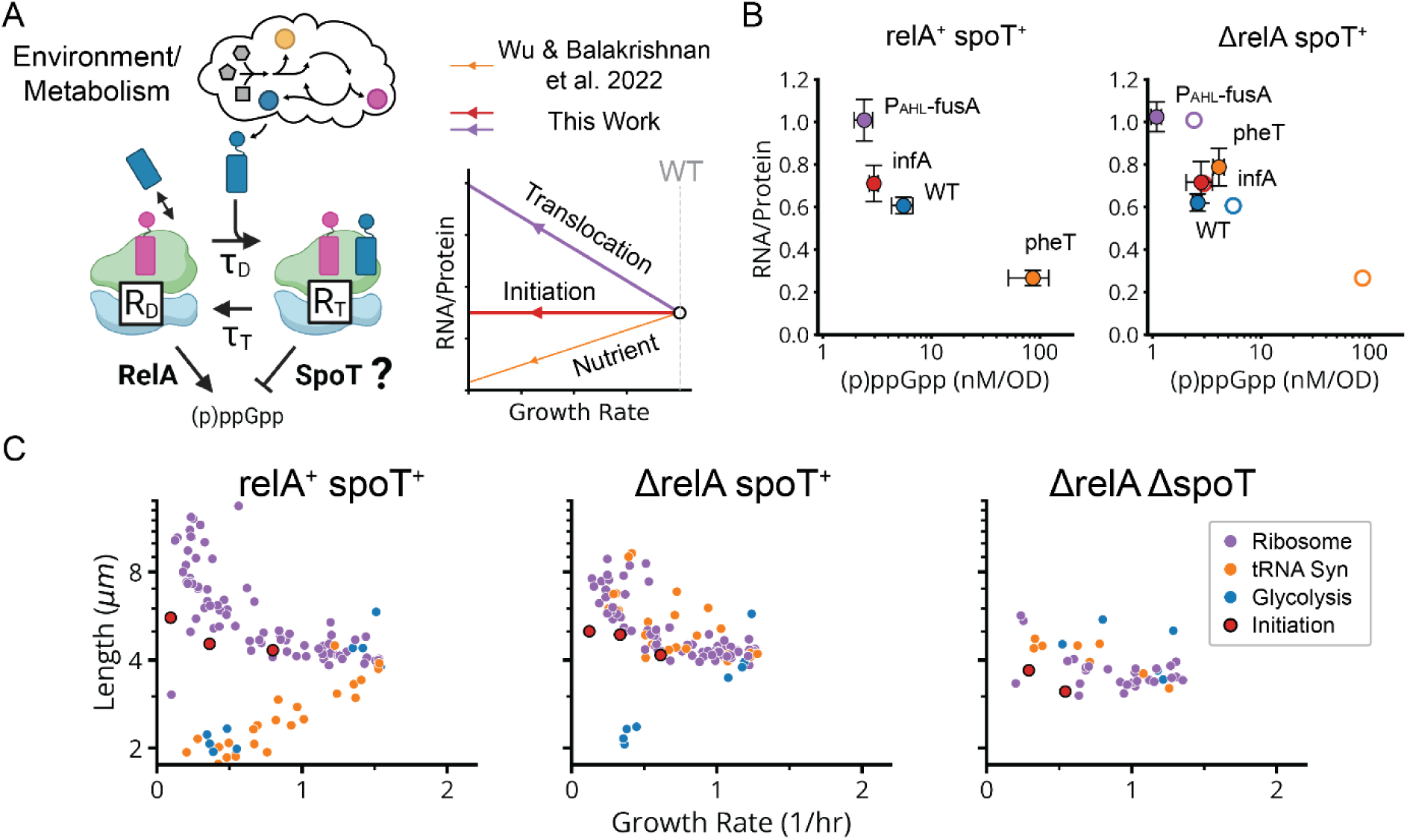
(p)ppGpp Degradation by Translocating Ribosomes Reconciles Diverse Translation Phenotypes. **a,** Left, illustration of a two-step model of elongation. Production of (p)ppGpp in proportion to uncharged or “dwelling” ribosomes (R_D_) is mediated by RelA, as previously described.^47^ Degradation of (p)ppGpp promoted by charged or “translocating” ribosomes (R_T_), in a SpoT-dependent manner, is proposed to reconcile our observations. Constitutive (p)ppGpp production by SpoT is assumed insignificant in WT cells, but considered as the only productive term in *ΔrelA spoT^+^*. Right, model-predicted changes in RNA/Protein vs growth rate under increased dwell time (orange), increased translocation time (purple) or reduced translation initiation (red). The model also accounts for RNA/Protein and size invariance under *rplL* knockdown, which interferes with initiation, translocation and charged tRNA capture (Theory Box and Supplementary Theory).^66^ **b,** Left, RNA/Protein and (p)ppGpp measurements of the indicated sgRNA knockdowns, P_AHL_-fusA and an EV sgRNA control (WT), in a *relA^+^ spoT^+^* background. All measurements made after 6 hours of CRISPRi induction or removal of AHL. Error bars denote 2σ_SEM_ range. Right, same plot in a *ΔrelA spoT^+^* background. Empty circles correspond to the positions of same-colored points in the *relA^+^ spoT^+^* plot. **c,** Average length vs average exponential growth rate for translation-related variants after 5-8 hours of CRISPRi induction in *relA^+^ spoT^+^, ΔrelA spoT^+^, ΔrelA ΔspoT* backgrounds (lDE26, lDE28 and lDE30, Extended Data Table 1). Ribosomal proteins are purple, tRNA synthetases are orange, glycolysis enzymes are blue, and initiation factors are red.

Second, 90 genes in our dataset – including 41 of the 45 strongly growth-inhibiting ribosomal protein (r-protein) knockdowns (Fig. 5a,b and Extended Data Fig. 11c) – *inverted* the SMK growth law and produced negative scaling between cell size and growth rate. This might also be rationalized using the tRNA charge explanation above: knockdowns of individual r-proteins often compromise ribosome elongation^46^ and decrease tRNA consumption, which increases tRNA charge and therefore reduces RelA activity and (p)ppGpp levels. Further supporting this explanation, we indeed observed the expected inversion in ribosome scaling, based on increased RNA/Protein at lower growth rates (Extended Data Fig. 11d). More specifically we observed a negative variant of the ribosome growth law, where ribosome levels again depend linearly on growth rate, but now with negative slope. We further found that ectopic expression of a hyperactive (p)ppGpp synthase (RelA*) largely restored size and ribosome content to wildtype levels (Extended Data Fig. 6g and 11d).

However, we then identified a third scaling class that shows *no discernable correlation* between size and growth (Fig. 5a,b and Extended Data Fig. 11a). This class included knockdowns targeting the few remaining r-proteins, located in the ribosome stalk, as well as translation initiators *infA* and *infB*. We measured (p)ppGpp and ribosome levels in an *infA* knockdown and found that they are indeed similar to wildtype (Fig. 6b). We further observed no change in ribosomes as a function of growth rate, creating yet another ‘linear’ ribosome growth law, now with zero slope. These observations stand in sharp contradiction to the tRNA charge explanation described above: knocking down initiation factors should mimic r-protein knockdowns and reduce the consumption of charged tRNA, reducing RelA activity. This led us to question our original rationalization that the inverse SMK scaling for r-protein knockdowns is due to changes in tRNA charge, and motivated us to search for some specific feature of translation that could discriminate in the response between the two classes of perturbations.

Since r-protein knockdowns have been reported to impact both elongation and initiation, we knocked down EF-G to compare the specific impact of impairing translocation but not initiation. The results closely mirrored the effects of knocking down r-protein genes: decreased growth, decreased (p)ppGpp, increased size and an increased RNA/Protein ratio (Fig. 5c, 6b)^47–49^. This demonstrates a clear difference in the control of (p)ppGpp levels by inhibiting initiation vs. inhibiting translocation, suggesting the existence of additional mechanisms for controlling (p)ppGpp levels beyond RelA.

### The Growth Laws are Upheld via Dual (p)ppGpp Control by Ribosome Translocation and Dwelling

A recent mathematical analysis by Wu et al.^47^, motivated by the quantitative scaling of (p)ppGpp with the elongation rate, posited that ribosomes may play a direct regulatory role not just in (p)ppGpp production via RelA, but also by making the (p)ppGpp degradation rate proportional to the number of A-site charged ribosomes primed to translocate. This proposal – at the time unsupported by direct experimental evidence – led to a model that could explain the classic positive ribosome growth law as a direct consequence of (p)ppGpp regulation. The model also predicts that (p)ppGpp levels should diminish in response to diminished translocation rates but not necessarily in response to diminished initiation rates, since in rich conditions translocation is limited by the ribosome’s own intrinsic kinetics rather than by initiation. Strikingly, these model predictions closely match our observations, prompting us to search for more direct evidence for translocation-driven (p)ppGpp regulation.

As noted in previous work^47^, the only known (p)ppGpp regulator in *E. coli* apart from RelA is the bifunctional enzyme SpoT which constitutively both synthesizes^50^ and degrades (p)ppGpp^51^ through largely unknown mechanisms that have appeared connected to ribosomes. We hypothesized that reduced translocation reduces (p)ppGpp via increased SpoT-dependent degradation (or possibly decreased SpoT-dependent synthesis), thereby inversing the SMK scaling (Fig. 6a). To test this, we sought to compare CRISPRi libraries with and without *spoT*. While the single *ΔspoT* knockout does not grow, due to the stabilization and accumulation of lethal amounts of (p)ppGpp^50^, the *spoT* deletion can be made in a *ΔrelA* strain. We therefore compared the growth/size scaling for three different CRISPRi libraries – the WT library above, but also new libraries built around the double knock-out *ΔrelA ΔspoT* and the single knockout *ΔrelA*.

In the *ΔrelA ΔspoT* background, which has no (p)ppGpp^15^, all scalings observed above collapsed to flat lines (Fig. 6c), confirming that they do rely on (p)ppGpp. In contrast, the single *ΔrelA* knockout, which should retain the putative SpoT-dependent reduction of (p)ppGpp under reduced translocation, retains the inverse SMK scaling under r-protein knockdowns that we observed in WT (Fig. 6c), showing that this effect does not depend on RelA. Similarly, when we directly target translocation by EF-G knockdowns in *ΔrelA*, we also observe a wildtype-like increase in ribosome levels and reduction in (p)ppGpp (Fig. 6b). Knocking down the initiation factor *infA* by contrast causes no change in size or ribosome levels in a *ΔrelA* background just as in WT (Fig. 6b,c).

In contrast to WT, however, in the *ΔrelA* strain, the tRNA synthetase knockdowns *do not* follow SMK scaling, but rather now show *inverse* SMK scaling (Fig. 6c) consistent with previous observations of increased cell size under single amino acid auxotrophy^51^. This supports the idea of a SpoT-dependent translocation response, as these knockdowns are predicted to impede translocation due to ribosome queuing upstream of codons with undercharged cognate tRNAs (Supplementary Theory). Consistent with this explanation, knockdowns which inhibit the synthesis of *multiple* amino-acids, which should not produce excessive queueing, do not exhibit this inverse scaling in *ΔrelA* (Fig. 6c). The presence of RelA appears to mask what would otherwise be an inverted, SpoT-mediated response to single amino acid depletion in WT. Thus, *spoT* is necessary for the reduction of (p)ppGpp under reduced translocation, while *relA* is required for standard SMK scaling under reduced tRNA synthetase activity.

These results support the Wu *et al* model^47^ by demonstrating translocation-specific control of (p)ppGpp. Determining whether that control is direct, via SpoT binding to ribosomes, or indirect, via other pathways, will require molecular and biochemical analyses. However, what is clear is that the SpoT-mediated control does distinguish changes in translocation from changes in tRNA charging. We find it remarkably insightful of Wu *et al.* to correctly raise such a specific yet likely correct hypothesis solely through kinetic modeling, in the absence at the time of genetic data.

We next evaluated whether the control mechanisms observed above can *quantitatively* explain all three variants of the ribosome growth laws (Fig. 5c), where the total ribosome level *R* depends linearly on the growth rate *λ*.^41,52,53^ Overall growth is proportional to total translation, but the proportionality between *λ* and *R* depends on the elongation rate per ribosome which in turn controls (p)ppGpp that then controls ribosome expression. Strikingly, we find that a simple first-order kinetic model of the combined RelA-SpoT system fully reproduces the conserved linearity, with different slopes, between ribosomes and growth rate across diverse perturbations of translation (Fig. 6a and Theory Box).

### The Nutrient Growth Law is Negative Hyperbolic and Emerges from Passive (p)ppGpp Regulation

Having established and explained our linear ribosome growth laws, we return to the nutrient growth law relating growth to size, for which a direct mechanistic explanation is still lacking. Because cell volume responds to growth in a (p)ppGpp-dependent manner, and (p)ppGpp directly controls transcription of many genes, it is tempting to hypothesize that some central division protein is under direct transcriptional control of (p)ppGpp. However, to our knowledge no such candidates have been identified, and most known division proteins have been shown *not* to be regulated by (p)ppGpp.^52^ Here we propose that not only the classical SMK growth law, where volume seems to depend exponentially on growth, but all three variants of growth vs volume scaling observed above, precisely reflect this *absence* of direct (p)ppGpp control of division proteins.

First, and similar to previous work linking proteome composition to cell size,^53^ our model postulates that cells on average divide after accumulating enough of some unspecified set of ‘division proteins’ that are not directly controlled by (p)ppGpp, whether structurally involved in division or in some indirect regulatory process (Fig. 7a). This assumption is supported by the observation that most division-related promoters are indeed not regulated by (p)ppGpp^52,54^ and is consistent both with *E. coli*’s behavior as a division ‘adder’^55,12^ where cells correct size deviations regressively by adding constant mass regardless of deviations in birth size, and the fact that changes in the expression of division proteins correlate with changes in cell size.^56^ In the simplest model, the average volume *V* of cells is then inversely proportional to the fraction or ‘sector’ *ϕ_NC_* of the proteome not controlled by (p)ppGpp, which is complementary to the fraction *ϕ_C_* =1 – *ϕ_NC_* that *is* controlled by (p)ppGpp, i.e., *V* ∝ 1/(1–*ϕ_C_*).

**Fig. 7:**
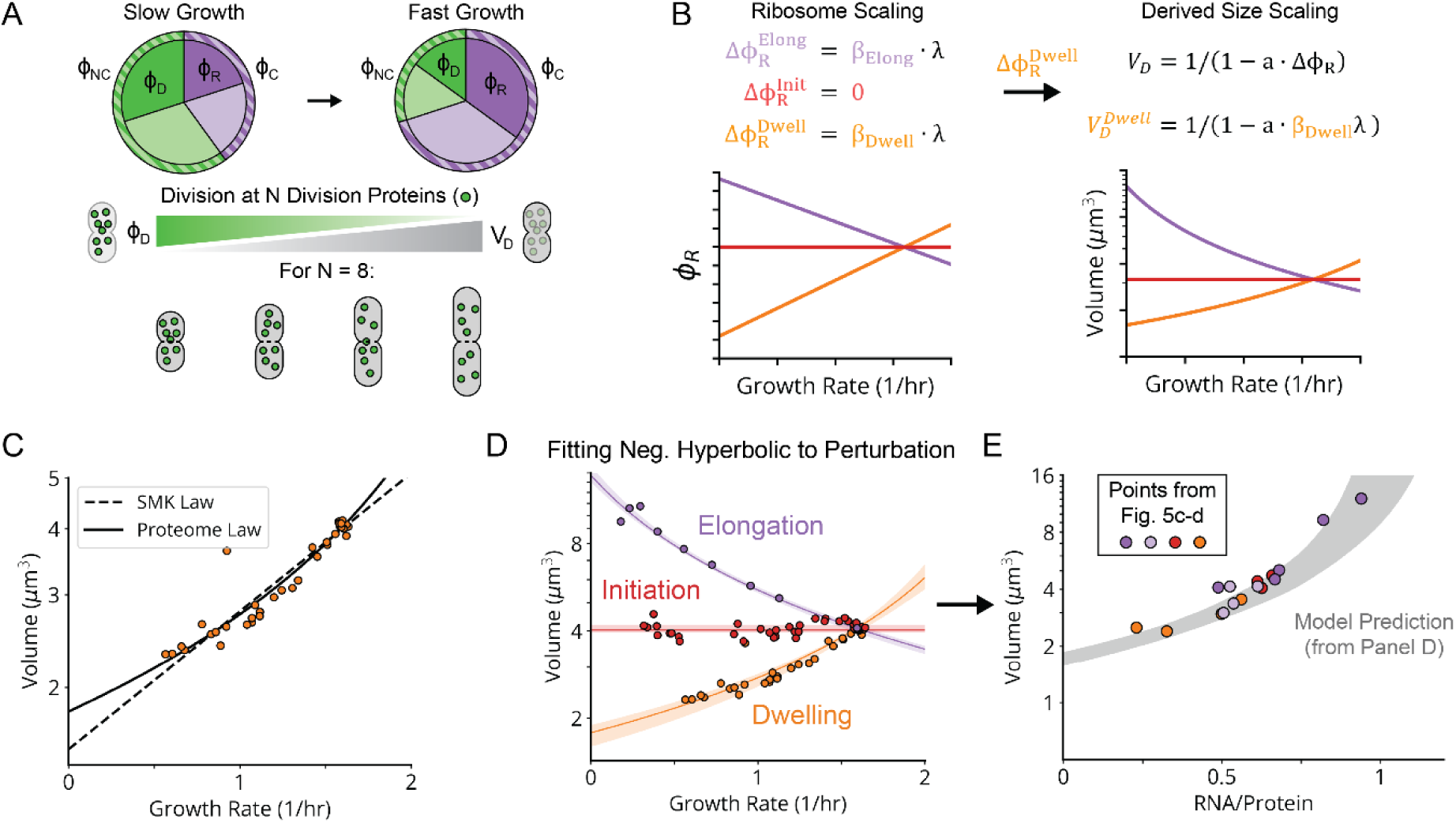
Size Scaling Law Emerges from Proteome Distortion by Ribosome Expression. **a,** Illustration of our size scaling model. The proteome is divided into two coarse-grained sectors, ϕ_C_ containing proteins repressed by (p)ppGpp and ϕ_NC_containing proteins that are not directly regulated by (p)ppGpp. ϕ_C_obeys the same regulation as ribosomal proteins and therefore the proteome fraction made up of ribosomes ϕ_R_ is a subset of this fraction. Division is assumed to occur when a subset of proteins in ϕ_NC_, with a fractional abundance of ϕ_𝐷𝐷_, reach a threshold abundance of N division proteins. **b,** Model-derived scaling of the ribosome proteome fraction ϕ_R_(left) and cell volume (right) with growth rate, under decreasing translocation rate (purple), initiation rate (red) and ternary complex capture rate (orange). **c,** Similar measurements as in Fig. 5c, for *pheT* knockdowns, with cell volume estimated from length and width assuming the cell has a spherocylindrical shape. Lines correspond to fits of the negative hyperbolic (i.e., Proteome) and exponential (i.e., SMK) size models. **d,** Same measurements as in Fig. 7c, for *pheT* (orange), *infA* (red) and *fusA*/EF-G (purple) knockdowns. Lines correspond to a three parameter, joint fit of the negative hyperbolic to *pheT* and *fusA* knockdowns, with the trendline for *infA* given by the point of intersection. 95% confidence intervals were determined by residual bootstrapping. **e,** Datapoints correspond to the empirical RNA/Protein vs volume relationship, derived from the data in Figs. 5c-d. Shaded area corresponds to the 95% confidence interval of the model-predicted relationship, given the fit in Fig. 7d.

Second, we further postulate that the sector *ϕ_C_* controlled by (p)ppGpp varies linearly with the ribosome sector, i.e., *ϕ_C_ = a*_1_ *ϕ_R_* + *b*_1_ with constant slope *a*_1_ and intercept *b*_1_ (Fig. 7a). This is approximately true simply because ribosomes constitute a large part of this sector. However, linearity also follows exactly if (p)ppGpp represses transcription from all (p)ppGpp-regulated promoters to the same degree, which likely is a good approximation since (p)ppGpp regulates expression by binding RNA polymerase far from the DNA binding domains, minimizing promoter-specific effects (Supplementary Information). By considering previous data^54,57,58^ across many growth conditions, we find that indeed the (p)ppGpp-repressed fraction of expression varies linearly with the ribosomal protein fraction (Extended Data Fig. 12). Connecting to the result above, this predicts a volume-ribosome relationship of the form *V* ∝ (1–*cϕ_R_*)^−1^ where *c* is a positive constant, as also directly supported by our measurements across different knockdowns (Fig. 7e).

Third and finally, we connect the ribosome sector *ϕ_R_* to growth rate *λ* with the linear ribosome growth law above, *ϕ_R_ = a*_2_ *λ* + *b*_2_, with different slopes *a*_2_ and intercepts *b*_2_ under knockdowns of tRNA charging, translocation and initiation (Fig. 7b).^59^

These three simple and experimentally supported assumptions predict a negative *hyperbolic* relationship between volume *V* and growth rate *λ*, i.e., of the form *V* ∼ (1–*kλ*)^−1^ rather than the previously proposed exponential curve *V ∼ e^kλ^*. However, the two curves can be very similar, and for the SMK perturbation class, the hyperbolic curve fits our data at least as well as the classical exponential, even though the exponential was chosen based on fitting (Fig. 7c) and the hyperbolic is independently predicted from experimentally supported mechanistic considerations. Strikingly, the hyperbolic function quantitatively captures all three perturbation classes (Fig. 7d), with class-specific size scaling constants proportional to their underlying ribosome scaling constants, consistent with our mechanistic expectations (Extended Data Fig. 13b). In fact, these fits require even fewer parameters than needed to fit three exponential curves. We believe that this for the first time explains the famous nutrient growth law of growing bacteria, and moreover generalizes to other classes of growth perturbations.

As noted above, this model is agnostic about the identity of the division proteins that mediate the inverse proportionality between division volume and division protein levels. However, we note that the assembly of the division-mediating Z-ring, which requires sufficient division protein to span the perimeter of the mid-cell cross-section, could mediate such a proportionality. Indeed, division volume exhibits a positive, roughly linear dependence on the mid-cell perimeter for the knockdowns that alter cell width (Extended Data Fig. 13a), and applying a perimeter-based correction to our size-growth scaling model results in the same or better quality of fit (Extended Data Fig. 13c).

It has been a long-standing mystery why so many bacteria follow the nutrient growth law with approximately exponential size-growth scaling.^12,40,60^ Our results show that hyperbolic size-growth scaling, which can easily be mistaken for exponential scaling, is a natural outcome in a model based on simple physiological observations. The linear relation between ribosome levels and growth rate, the complementary linear decrease in protein sectors not co-regulated with ribosomes, and the lack of strong co-regulation between division and ribosomes, may therefore all hold broadly. Many other cell processes could similarly respond to changes in conditions in a seemingly controlled and coordinated manner, simply by *not* being directly regulated by (p)ppGpp or its equivalents. In fact, properties that stay constant across conditions are more likely to be under (p)ppGpp control. For example, the initiation volume per origin of replication in *E. coli* changes very little between environments, requiring levels of the key regulator DnaA to be maintained at a constant level. DnaA is indeed controlled by (p)ppGpp.^54^

Evolutionarily, all these quantitative patterns, from division ‘adders’ to nutrient or ribosome growth laws, could just be spandrels – byproducts of selection on other traits, such as maximal ribosome efficiency. However, cell size matters for many reasons, including the surface-to-volume ratio needed to support nutrient intake, stochastic low-number effects, or transport rates across the cell. If other size-growth patterns were more beneficial, it is easy to imagine how they could be generated, for example, by adding or removing (p)ppGpp control elements to a few genes. We therefore view the simple passive control above as an elegant and highly efficient way for cells to ensure baseline size adaptation to different conditions.

## Conclusion

Our MARLIN platform enables multi-generational imaging of bacteria in stable, micro-controlled environments, followed by precise genotyping of every cell. The scale is currently at the level of a million parallel lineages – each of which could be a separate strain – that can grow and divide in precisely controlled environments for tens of consecutive generations while every cell is segmented and tracked. The phenotyping could go much deeper in future studies, using more imaging modalities and characterizing more cellular processes and subcellular structures^61,62^, including genome-wide reporting. Given the ease of building, sharing and storing pooled libraries, coupled with the recent explosion of computational approaches for genomics and model identification, we believe MARLIN will help transform our understanding of microbes, whether by allowing deep mutational scans of RNAs and proteins for systematic structure-function relationships, or by systematically perturbing expression to annotate unknown genes and dissecting regulatory networks.

### Theory Box: The Three Variants of the Ribosome Growth Laws can be Quantitatively Explained by the RelA/SpoT Mechanism

Here we demonstrate that the (p)ppGpp control mechanisms identified are quantitatively consistent also with the two new variants of the ribosome growth laws observed (Fig. 5c), where the total ribosome level *R* depends linearly on the growth rate *λ* with negative or zero slope. Because translation is proportional to cell growth, *λ* will be proportional to *R* multiplied by the per-ribosome efficiency *εf*, where *ε* is the elongation rate per ribosome and *f* is the fraction of ribosomes involved in translation, *λ* ∝ *εfR*. The challenge in determining the actual relation between *R* and *λ* is that all these variables are interconnected, where changes that vary *ε* then impact (p)ppGpp which in turn impacts ribosomes and – for combinations of slow growth and high (p)ppGpp – even can reduce *f*.

The classical positive linear scaling for perturbations in nutrients and tRNA charge follows directly in the regime of fast growth rates, because most ribosomes are involved in translation and *ε* saturates to a constant value – ultimately limited by either intrinsic translocation kinetics or ternary complex diffusion^63^ – and so *λ* ∝ *εR* ∝ *R*, ignoring the small constant offset due to basal ribosome expression at zero growth. At slower growth, changes in *ε* and *f* can no longer be neglected,^64^ yet *R* remains on the same linear curve with *λ*, now with the small basal offset no longer being negligible. This was previously explained^47^ as a consequence of the proposed (p)ppGpp regulation establishing a correspondence between *ε*, *f* and *R* that maintains linearity in *λ* vs. *R* and also explains the maintenance of a basal ribosome pool at zero growth (Supplementary Information).

The negative linear growth-ribosome scaling for perturbations to translocation cannot be explained by changes in *f* since, for these perturbations, (p)ppGpp levels are lower overall and most ribosomes should therefore be active, such that *λ* ∝ *εR*. However, perturbing translocation will still impact *ε* which in turn impacts (p)ppGpp and thereby *R*. Specifically, the elongation rate is set by the average ribosome dwell (*τ_D_*) and translocation (*τ_T_*) times of each elongation step, *ε* =1/(*τ_D_* +*τ_T_*). In the regime where perturbing translocation starts to impact growth, the second term is expected to dominate, such that *ε* ≈ 1/*τ_T_*.

Dwelling ribosomes with bound RelA produce (p)ppGpp,^44,65^ while, as observed above, translocating ribosomes directly or indirectly seem to cause (p)ppGpp hydrolysis (see Supplementary Information for alternative possibility of reduced synthesis). While we don’t explicitly model parallel, ribosome-independent (p)ppGpp production and degradation pathways, the impact of our translocation perturbations to increase both ribosome abundance and per-ribosome (p)ppGpp hydrolysis sufficiently to reduce (p)ppGpp levels indicates that this ribosome-specific activity likely dominates. Assuming first-order kinetics, changes in dwell or translocation times then approximately and proportionally increase or decrease the (p)ppGpp level *G*, such that *G* ∝*τ_D_*/*τ_T_* (Supplementary Information). For perturbations to translocation, *τ_D_* stays constant and *G* ∝1/*τ_T_*, which combined with the results above means that *ε* ∝ *G* and thus approximately *λ* ∝ *GR*. For a closed expression in terms of *R,* we next express *G* in terms of *R* by considering how (p)ppGpp levels control ribosome production.

Like all cellular components, ribosomes are diluted through first-order kinetics by cell growth, i.e., with total rate *λR*. With (p)ppGpp inhibiting ribosome production without strong cooperativity (Supplementary Theory),^47,49^ we approximately expect total ribosome production to be proportional to *λR_max_*/(1+*G*/*K*_1_) where *R_max_* and *K*_1_ are constants. At steady state this leads to *R* ∝ *R_max_*/(1+*G*/*K*_1_), similar to what has been observed experimentally and used in previous models^47^, and allows us to write (p)ppGpp in terms of the ribosome levels they control, *G* ∝ *R_max_*/*R –* 1. Combined with the results above this directly predicts the observed negative linear scaling *λ* ∝ *GR* ∝ *R_max_ – R*. Similar results arise when accounting for the small level of inactive ribosomes observed^64^ (Supplementary Information). The negative linear observation thus follows from first order control of (p)ppGpp decay via translocation, and first order control of ribosomes via (p)ppGpp.

Finally, for the flat scaling for perturbed initiation of translation, we expect no changes to *τ_D_* and *τ_T_*, and therefore predict and observe no change in *G* and thus no change in (p)ppGpp-controlled ribosome production, despite reductions in growth rate from reduced translation initiation rates.

In summary, a minimal, well-supported RelA-SpoT model can capture the ribosome-growth linearity, with different slopes, that we observe across our diverse translation perturbations.

## Supporting information

Supplementary Theory

---

**Extended Data Fig. 1:**
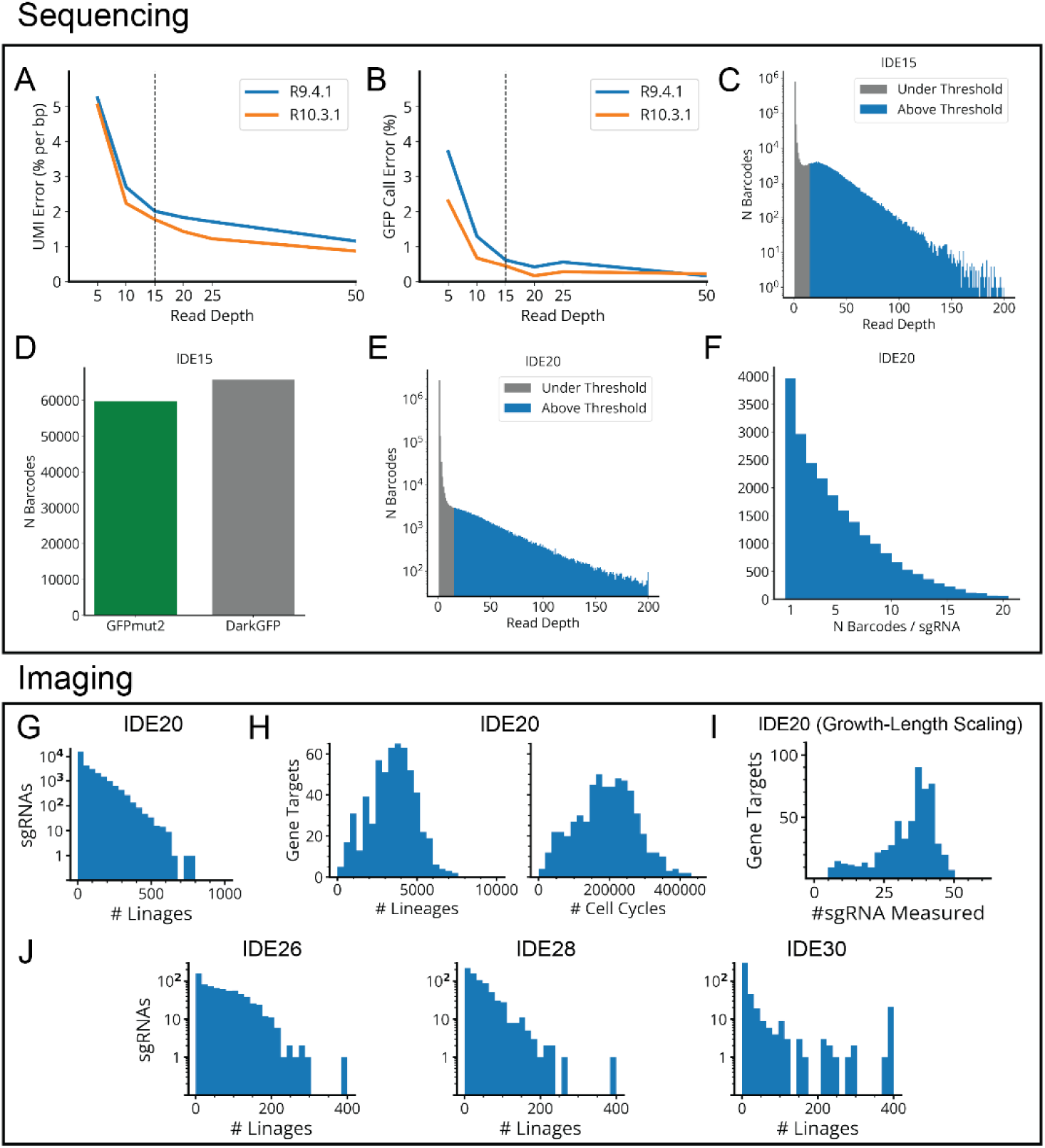
Sequencing Error Rates and Sampling Distributions for MARLIN Experiments. **a,** Error rate (per bp) in the random 15-bp UMI vs read depth, for synthetically down-sampled data. lDE11 (small *gfp* variant library) variants with read depth >200 were sequenced and their consensus UMIs taken as ground truth. Reads assigned to these variants were randomly downsampled to a fixed read depth, re-run through our consensus calling algorithm and compared to the high depth ground truth to estimate the error rate. **b,** Error rate in calling GFPmut2 vs. DarkGFP vs read depth, for synthetically down-sampled data generated as in A. **c,** Log scale histogram of the read depth distribution for each lDE15 (large *gfp* variant library) barcode, with a depth cut-off of 15 reads. Grey bars are below the depth threshold and do not contribute to the codebook, while blue bars above the threshold do. **d,** Bar chart showing the number of sequenced barcodes assigned to the GFPmut2 and DarkGFP variants. **e,** Log scale histogram of the read depth distribution for each lDE20 (mismatch-CRISPRi library) barcode, with a depth cut-off of 15 reads. **f,** Histogram of the number of MARLIN barcodes associated with each lDE20 sgRNA sequence. **g,** Histograms of the number of lineages measured for each designed sgRNA in lDE20. **h,** Histograms of the number of lineages or cell cycles observed per gene target in lDE20. **i,** Histogram of the number of sgRNAs measured per gene target, for sgRNAs passing the quality threshold (>7 lineages, CV_SEM_< 0.2) for inclusion in our growth-length scaling analysis (Fig. 5-6). **j,** Same histogram as panel g for lDE26, lDE28 and lDE30 (HU-mCherry libraries).

**Extended Data Fig. 2:**
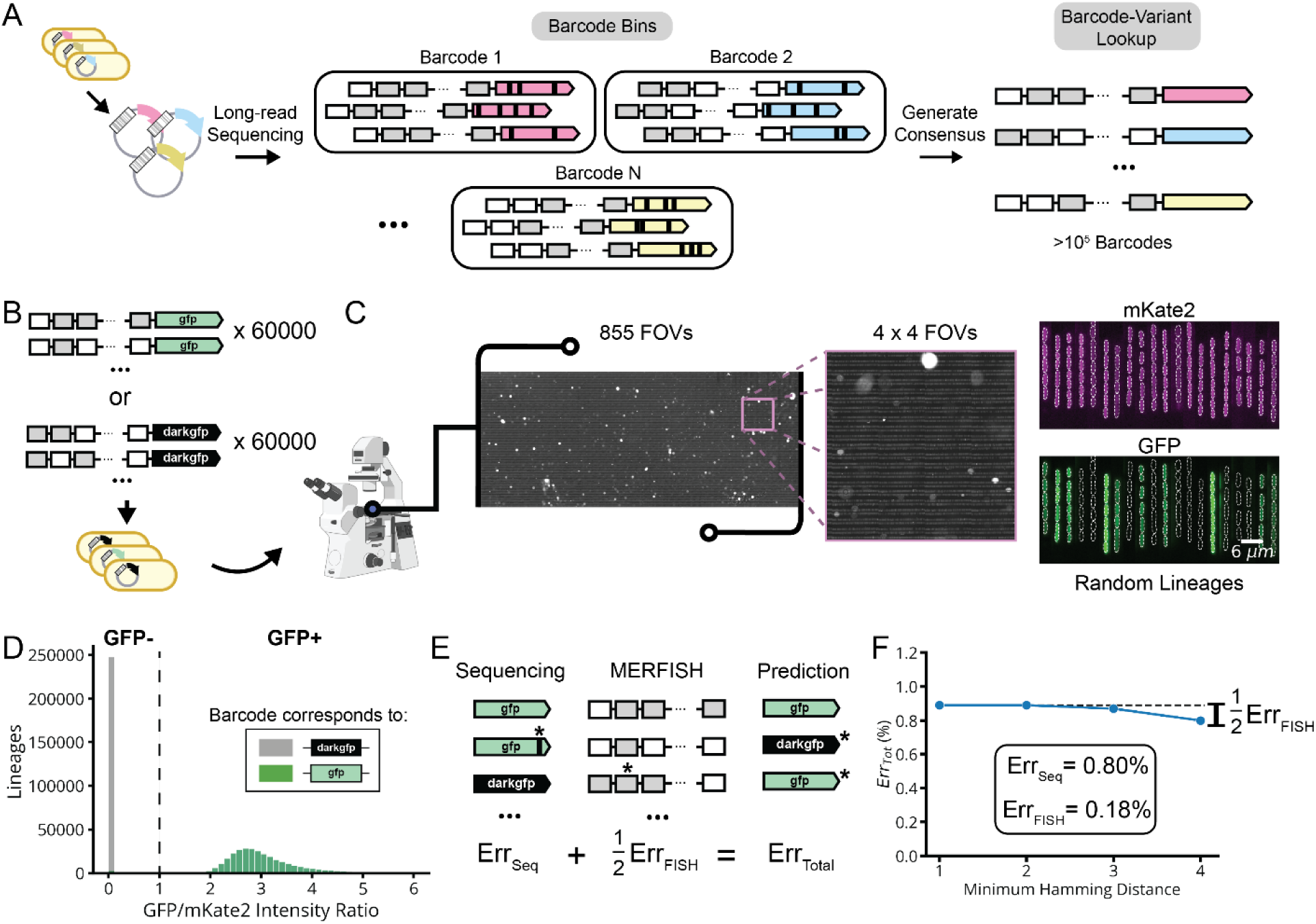
Barcode Error Rate Measurement. **a,** The barcode-library variant lookup table is established using nanopore sequencing of our pooled plasmid library. Reads are robustly grouped by FISH barcode in the presence of sequencing-associated error, which is subsequently corrected by consensus averaging for each group. **b,** A barcoded library (lDE15) of two fluorescent protein variants, GFP (i.e., GFPmut2) and DarkGFP, was measured by MARLIN to estimate our barcode error rates. **c,** During imaging of lDE15, we measured the fluorescent signal from both a constitutively expressed fluorescent protein encoded on the chromosome (pRpsL-mKate2Hyb) and the constitutively expressed GFP variant expressed on the lDE15 plasmid, every 15 minutes for 6 hours. **d,** Histogram of the average GFP/mKate2 intensity ratio for each lineage in the experiment. We called the lineage as being GFP positive if this ratio was above 1 (dotted line). **e,** Decomposition of total experimental error into contributions from library sequencing and MERFISH readout. Note that the FISH contribution is halved since barcode assignment errors will predict an incorrect GFP variant roughly half the time. **f,** Estimating error contributions by increasing error robustness of barcodes. Computationally filtering lDE15 data for sub-libraries composed of well-separated barcodes with hamming distances of up to four should eliminate the FISH contribution to error. The marginal difference of ∼0.09 indicates that the FISH error contribution is ∼0.18%. See Supplementary Methods for error estimation details.

**Extended Data Fig. 3:**
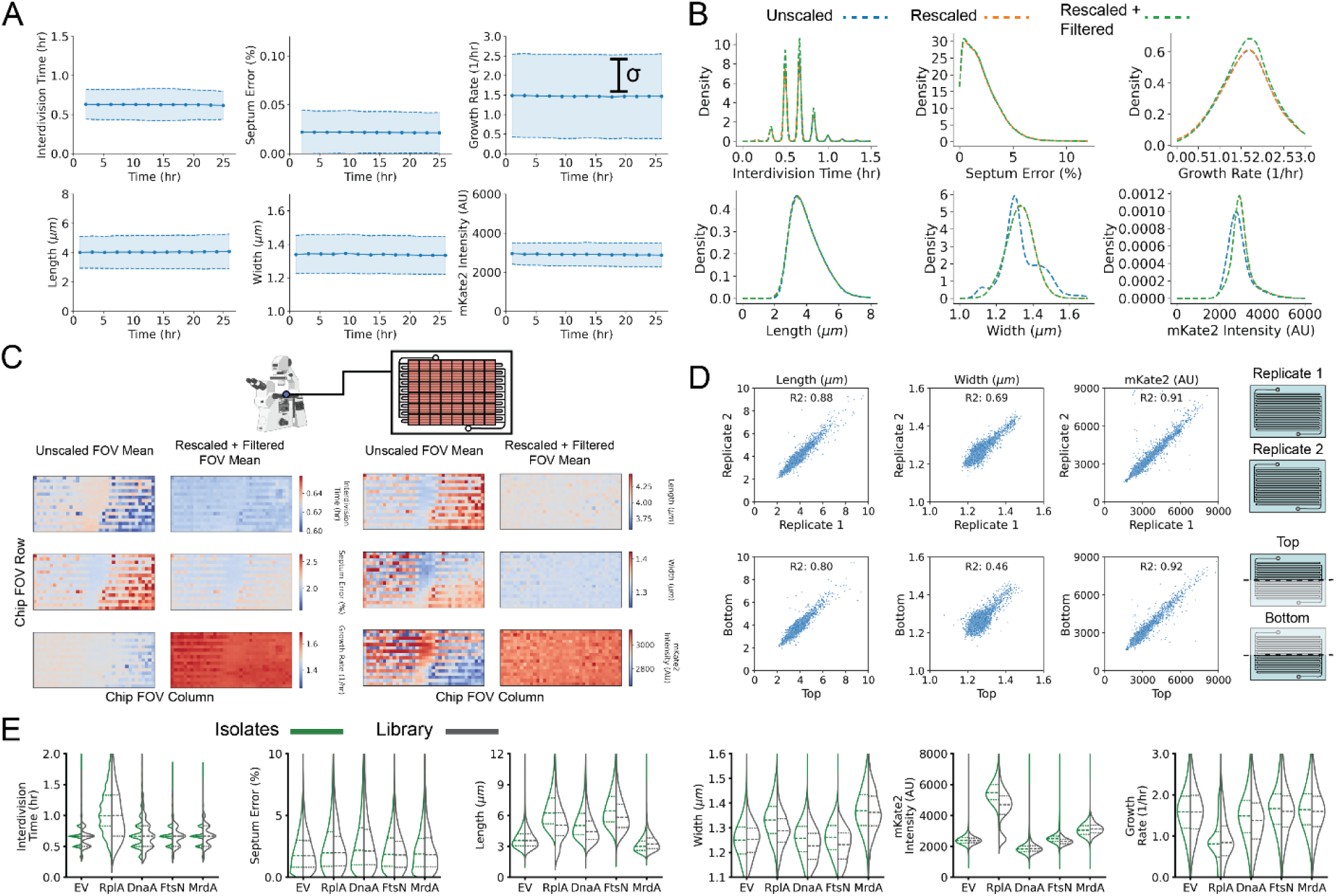
Phenotypic Quantities in MARLIN-CRISPRi Experiments are Highly Reproducible. **a,** Mean and standard deviation estimates for physiological parameters over time for 26 hours of growth of DE32. Values are computed for each 2-hour time window. **b,** Kernel density estimates for distributions of measured physiological parameters without correction (blue line), with trench-wise baseline rescaling (orange line) and with both rescaling and filtering based on growth characteristics (green line). See Supplementary Methods for details. **c,** Mean values of physiological parameters within each FOV across the chip using uncorrected or rescaled and filtered values. **d,** Top, relationship between cell length, width or mKate2 intensity, for each sgRNA in replicate lDE20 experiments. Points correspond to average values in the period from 5 to 8 hours post-induction, for the same sgRNA in each experiment. Bottom, relationship between cell length, width or mKate2 intensity, for each replicate sgRNA in the top or bottom half of the microfluidic. **e,** Distributions of measured physiological parameters in the period from 5 to 8 hours post-induction, for the indicated CRISPRi knockdowns. Distributions correspond to either values determined from the library experiment (right, grey) or from corresponding clonal isolates (left, green). Dotted lines indicate the quartiles of the distributions.

**Extended Data Fig. 4:**
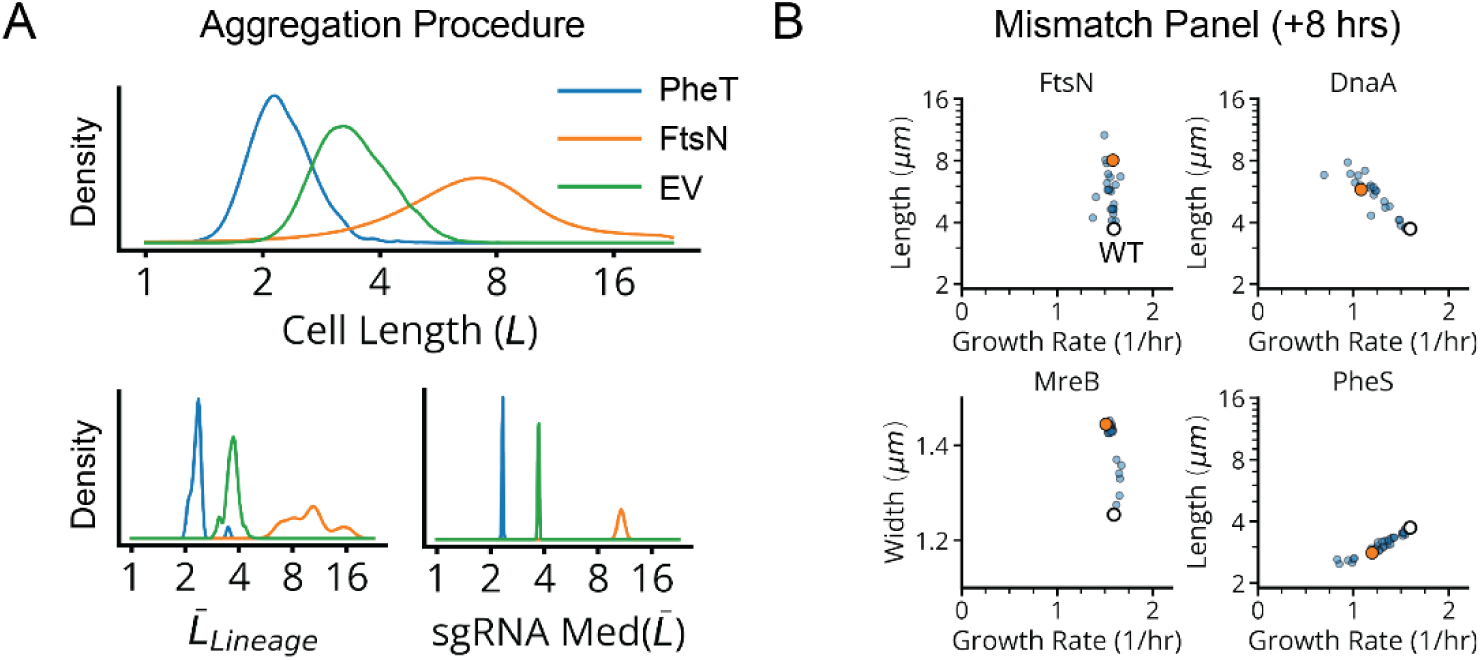
Data Aggregation Procedure for CRISPRi Libraries. **a,** Top, single-cell length distribution for sgRNA variants targeting the indicated genes, between 5-8 hours of CRISPRi induction. Bottom, bootstrap estimator distributions for the length averaged within single lineages (left) and the median of these average lengths, across trenches containing the same sgRNA (right). **b,** Length vs. growth rate plots for sgRNAs targeting the indicated gene after 8 hours of CRISPRi induction. The white circle indicates the mean length and growth rate of control strains, while the orange dots indicate sgRNAs used in the HU-mCherry reporter library (lDE26).

**Extended Data Fig. 5:**
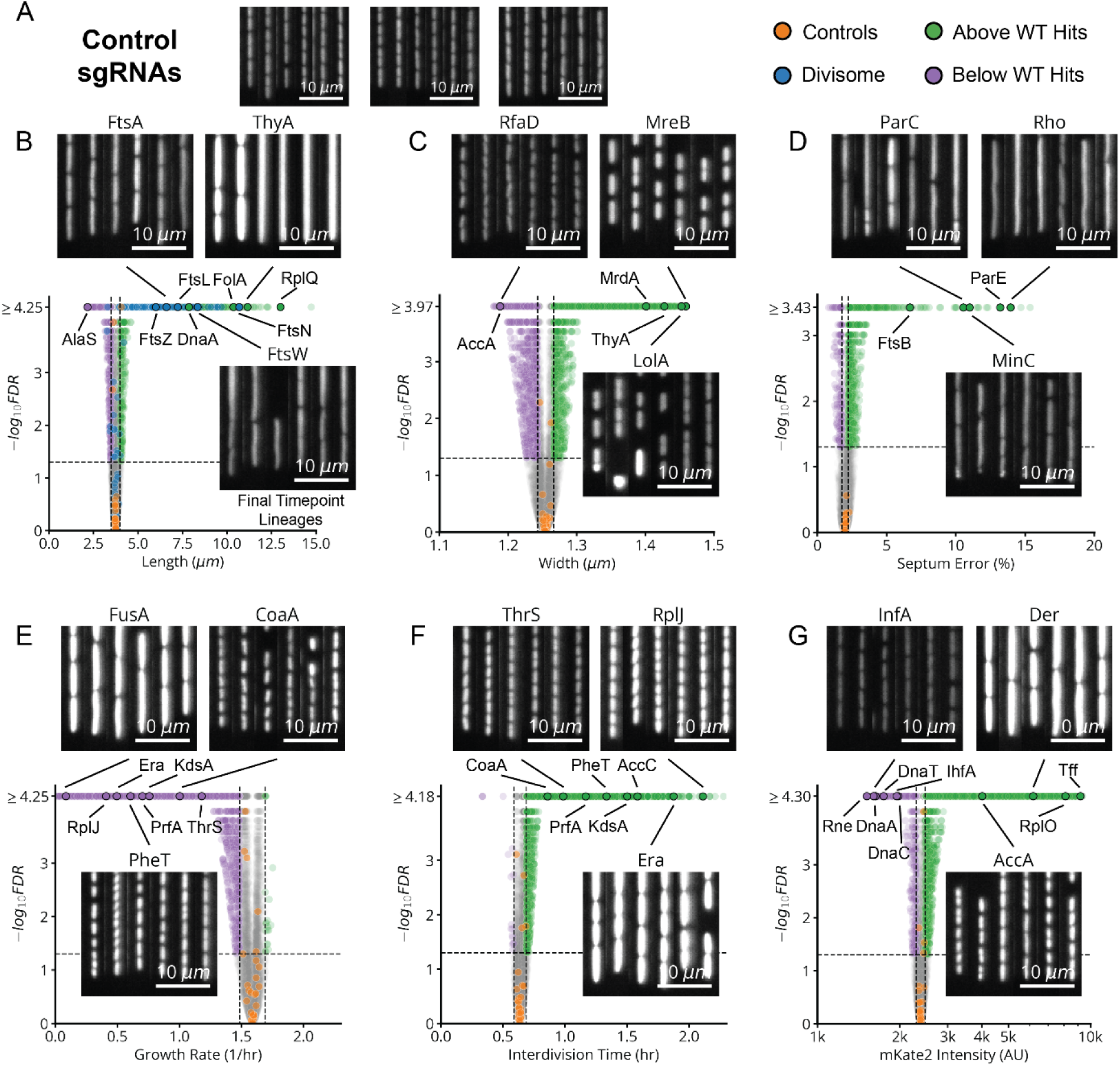
Volcano Plots of 6 Physiological Quantities After CRISPRi Knockdown. **a,** Sample lineages of EV control strains, after 8 hours of CRISPRi induction in our mismatch-CRISPRi library (lDE20). **b,** Volcano plot of cell lengths measured between 5-and 8-hours post-induction, for each sgRNA in lDE20. The horizontal line corresponds to FDR=0.05 and the vertical lines indicate three standard deviations above and below the distribution of control sgRNAs. Sample lineages are shown of representative strains, imaging the cytoplasmic pRpsL-mKate2Hyb marker. **c-g,** Similar volcano plots also included for cell width (**c**), septum placement error (**d**), growth rate (**e**), interdivision time (**f**), and mKate2 intensity (**g**). Consistent with previous reports, depletions targeting members of the cell wall synthesizing Elongasome (*mrdAB*, *mreBCD*, *rodZ*) exhibit an increase in width of >0.1 µm relative to EV controls.^31,67,68^ Also similar to previous reports^32,69^, depletions targeting the division localization factor *minC* and nucleoid partitioning *parCE* system exhibit an increase in septum placement error of >10% relative to EV controls.

**Extended Data Fig. 6:**
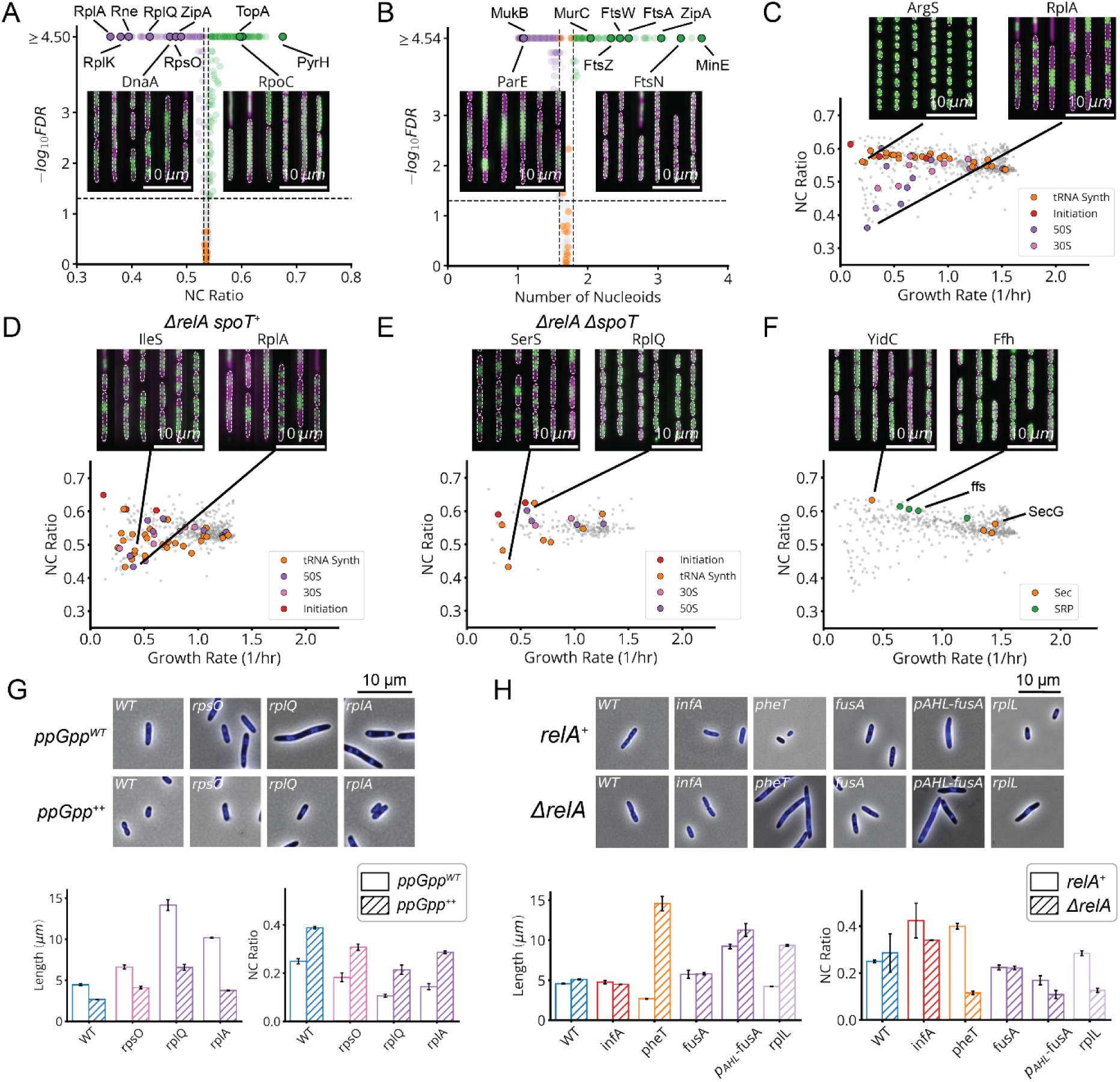
Nucleoid Compaction is Associated with Translation Elongation Defects, without RelA Activation. **a,** Volcano plot of the nucleoid to cell area (NC) ratio, measured between 5- and 8-hours post-induction, for each sgRNA in our HU-mCherry library (lDE26). The horizontal line corresponds to FDR=0.05 and the vertical lines indicate two standard deviations around the distribution of control sgRNAs. Sample lineages are shown of representative strains, where indicated. The cytoplasmic pRpsL-mVenus marker is shown in magenta and the nucleoid-localized HU-mCherry marker is shown in green. **b,** Volcano plot of the number of nucleoids in the cell. **c,** Relationship of NC ratio to growth rate, for all sgRNAs in lDE26. Indicated dots are genes coding for tRNA synthetases (tRNA Synth), translation initiation factors (Initiation), the 50S ribosomal subunit (50S) or the 30S ribosomal subunit (30S). **d,** Same measurement as in panel c, for each sgRNA in our *ΔrelA* library (lDE28). **e,** Same measurement as in panel c, for each sgRNA in our *ΔrelA ΔspoT* library (lDE30). **f,** Same measurement as in panel c. Indicated dots are genes coding for the Sec translocation complex (Sec) or the signal recognition particle (SRP). **g,** Cell length and NC ratio, as measured by DAPI staining, in CRISPRi strains depleted for the indicated gene products, WT denoting the EV sgRNA control. Top, snapshots of the indicated isolates imaged by phase contrast (greyscale) and for fluorescent DAPI stain (blue). *ppGpp^WT^* indicates cells uninduced for the hyperactive (p)ppGpp synthase RelA* and *ppGpp^++^* indicates cell cultured with +25uM IPTG to induce RelA* expression, resulting in ectopic synthesis of (p)ppGpp. Bottom, length and NC ratio measurements of the same strains. Error bars denote 2σ_SEM_ range. **h,** Same measurements as panel g. *ΔrelA* indicates the knockdown was performed in a *ΔrelA* background, while *relA^+^* indicates an intact locus. pAHL-fusA denotes the AHL induction strain introduced in Fig. 5c, in which *fusA* was depleted according to the same protocol as in Fig. 5d. Note the difference between this strain and *fusA*, which likely owes to disrupted expression of the downstream *tufA* gene.

**Extended Data Fig. 7:**
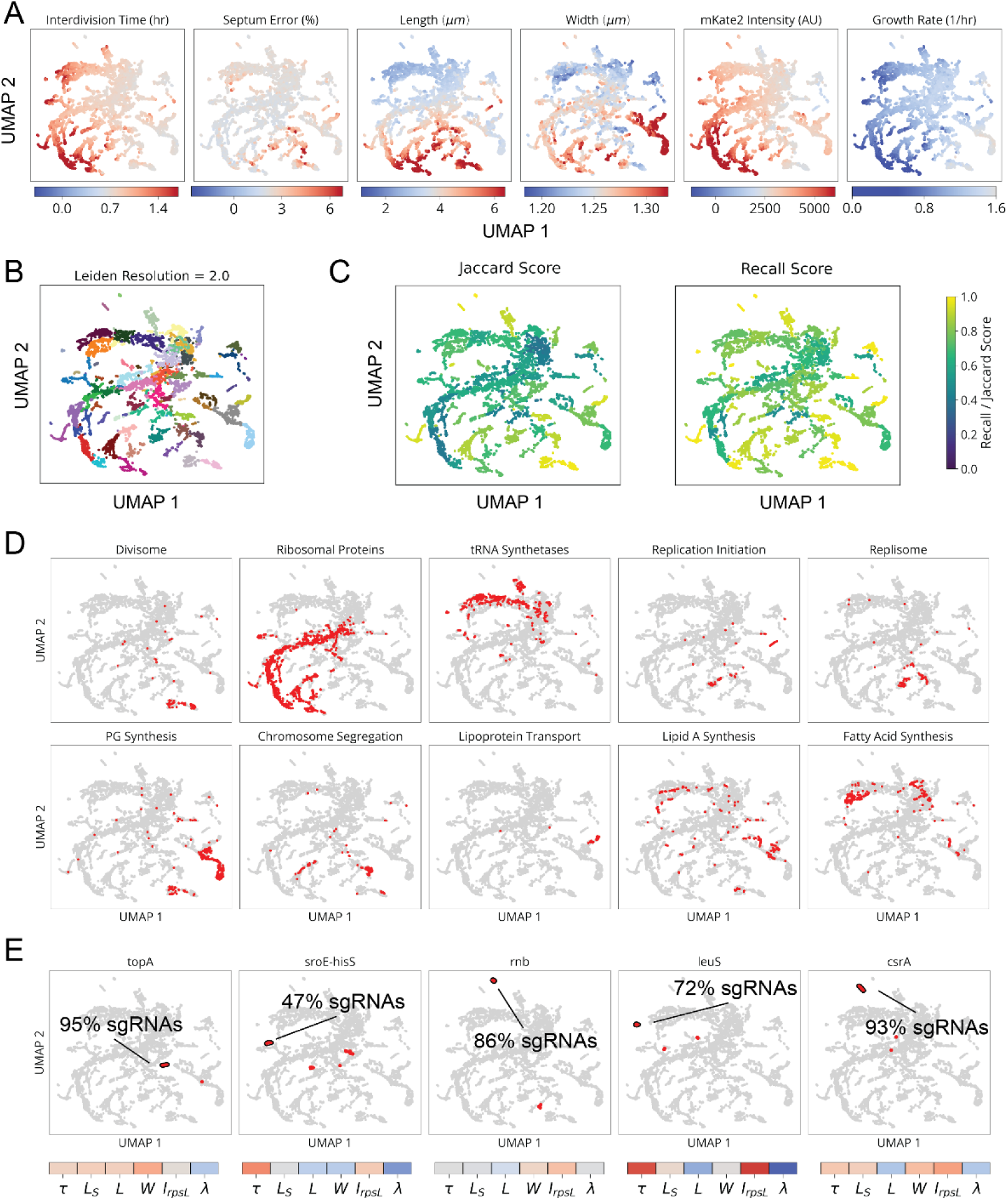
Clustering and Phenotype Distributions in MARLIN-CRISPRi Library. **a,** Distribution of measured parameters in our mismatch-CRISPRi library (lDE20). Colors on UMAPs indicate whether sgRNA variants on the UMAP take an average parameter value above (red), below (blue) or at the value of the sgRNA EV controls (white). **b,** Clusters determined by the Leiden community detection algorithm^70^, indicated by color, with a resolution of 2. **c,** Cluster robustness at different resolutions, determined from 10% jackknife resampling of Leiden clusters and expressed as either the average Jaccard score or the average recall score between a cluster and the most similar cluster in each resampling. **d,** Distribution of different gene groups across lDE20 UMAP, indicated in red. **e,** Top, distribution of different genes or operons across lDE20 UMAP, indicated in red. Points outlined in black correspond to Leiden clusters containing primarily the indicated gene or operon. The percentage corresponds to the fraction of sgRNAs targeting the indicated gene/operon that appear in the selected cluster. Bottom, the average phenotype of the indicated cluster, using the same scaling as in panel a.

**Extended Data Fig. 8:**
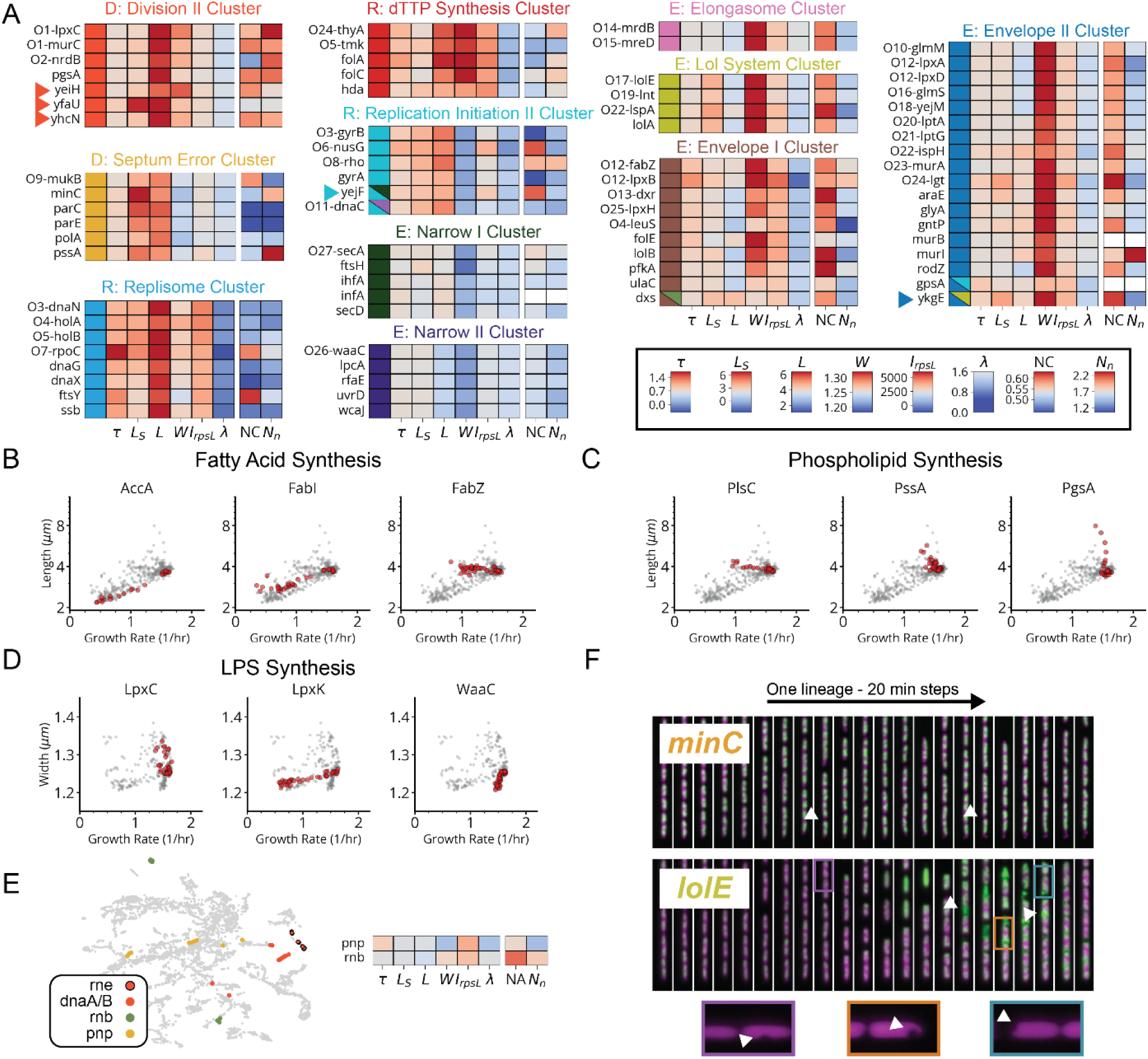
Extended Cluster Heatmaps and Select Scaling Phenotypes for Genes Involved in Fatty Acid, Phospholipid and LPS Synthesis. **a,** Heatmap of cluster phenotypes. Clusters belong to the division “D”, replication “R” or envelope “E” supergroups, as indicated by the title. Arrows indicate genes mentioned in the text. The leftmost column color denotes cluster membership, right columns display the average value of each quantity at the end of the experiment, for each gene. Same scale as in Fig. 4c. Entries beginning with O denote polycistronic operons, where the subsequent gene name indicates the last gene in a consecutive block all belonging to the indicated cluster. **b-c,** Semi-log plots of average growth rate vs length, measured between 5- and 8-hours post-induction, for sgRNAs in our mismatch-CRISPRi library (lDE20) targeting the indicated genes (red) involved in phospholipid and fatty acid synthesis (grey). **d,** Semi-log plots of average growth rate vs. width, measured between 5- and 8-hours post-induction, for sgRNAs in our mismatch-CRISPRi library (lDE20) targeting the indicated genes (red) involved in LPS synthesis (grey). **e,** Left, colored dots correspond to sgRNAs targeting the indicated genes involved in RNA degradation, mapped onto a UMAP representation of each sgRNA variant. Right, heatmap of gene phenotypes for *rnb* and *pnp*. **f.** Representative kymographs of *minC* and *lolE* knockdowns, with the cytoplasm in magenta and nucleoid in green. Top, knockdown, arrows indicating asymmetric divisions. Bottom, kymograph of a *lolE* knockdown, arrows indicating extracellular HU-mCherry resulting from lysis. Insets highlight instances of cytoplasmic invagination and plasmolysis.

**Extended Data Fig. 9:**
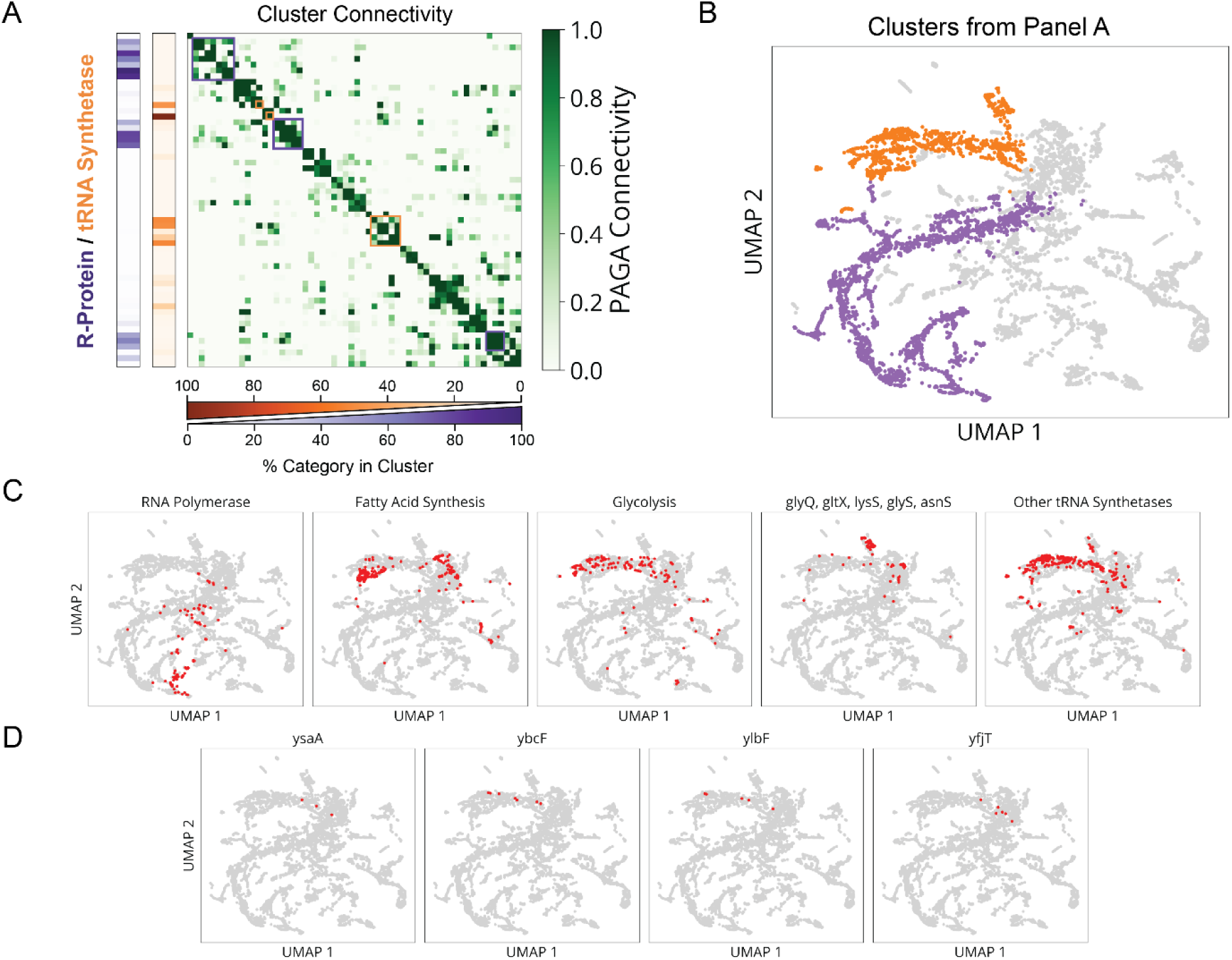
Ribosomal Protein and tRNA Synthetase Clusters are Highly Interconnected. **a,** Left, heatmap indicates the percentage of each cluster composed of either ribosomal proteins (purple) or tRNA synthetases (orange), with each row corresponding to a cluster in our mismatch-CRISPRi library (lDE20). Right, heatmap values indicate statistical connectivity between clusters using the PAGA connectivity metric^71^. Rows are matched to the same clusters as in the left heatmap. Boxed cluster groups indicated highly inter-connected groups enriched for either ribosomal proteins (purple) or tRNA synthetases (orange). **b,** Highlighted UMAP indicating the inter-connected cluster groups corresponding to the boxes in panel a, with the same color scheme. **c,** Distribution of different gene groups across lDE20 UMAP, indicated in red. **d,** Distribution of different genes of unknown function across lDE20 UMAP, indicated in red.

**Extended Data Fig. 10:**
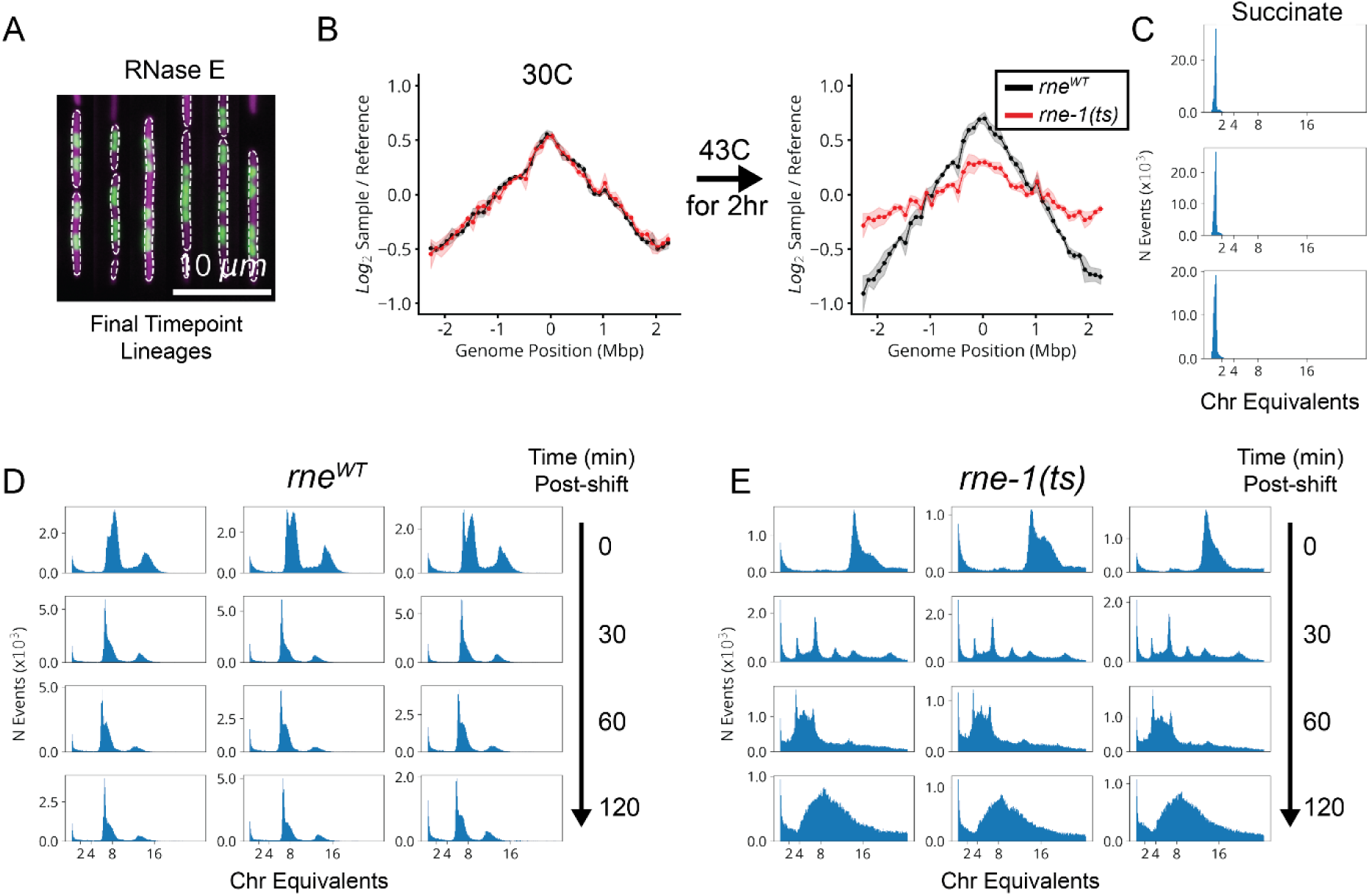
*rne-1(ts)* Cells Rapidly Lose Initiation Activity at a Restrictive Temperature. **a,** Representative lineages depleted for *rne* after 8 hours of CRISPRi induction in our HU-mCherry library (lDE26). The cytoplasmic pRpsL-mVenus marker is shown in magenta and the nucleoid-localized HU-mCherry marker is shown in green. **b,** Marker frequencies based on deep gRNA sequencing for 100kb bins across the *E. coli* genome, with the reference coordinate set to *oriC*. Frequencies normalized to reference measurements from non-replicating cells grown to stationary phase in MBM+0.2% Succinate at 37 °C. Measurements before (left) and after (right) shifting either *rne^WT^* (black) or *rne-1(ts)* (red) cells from 30 °C to 43 °C for 2 hours, with shaded area indicating ±2σSEM based on three biological replicates. **c,** Triplicate flow cytometric measurement of PicoGreen-stained DNA after replication runout for *rne^WT^* cells grown to stationary phase in MBM+0.2% Succinate at 37 °C, used to determine fluorescence intensity of a single chromosome. Chromosome equivalents reported as various multiples (2,4,8,16) of this value. **d,** Same as panel c, for *rne^WT^*cells before (0 mins) and at various times after a shift from 30 °C to 43 °C. **e,** Same as panel d, for *rne-1(ts)* cells.

**Extended Data Fig. 11:**
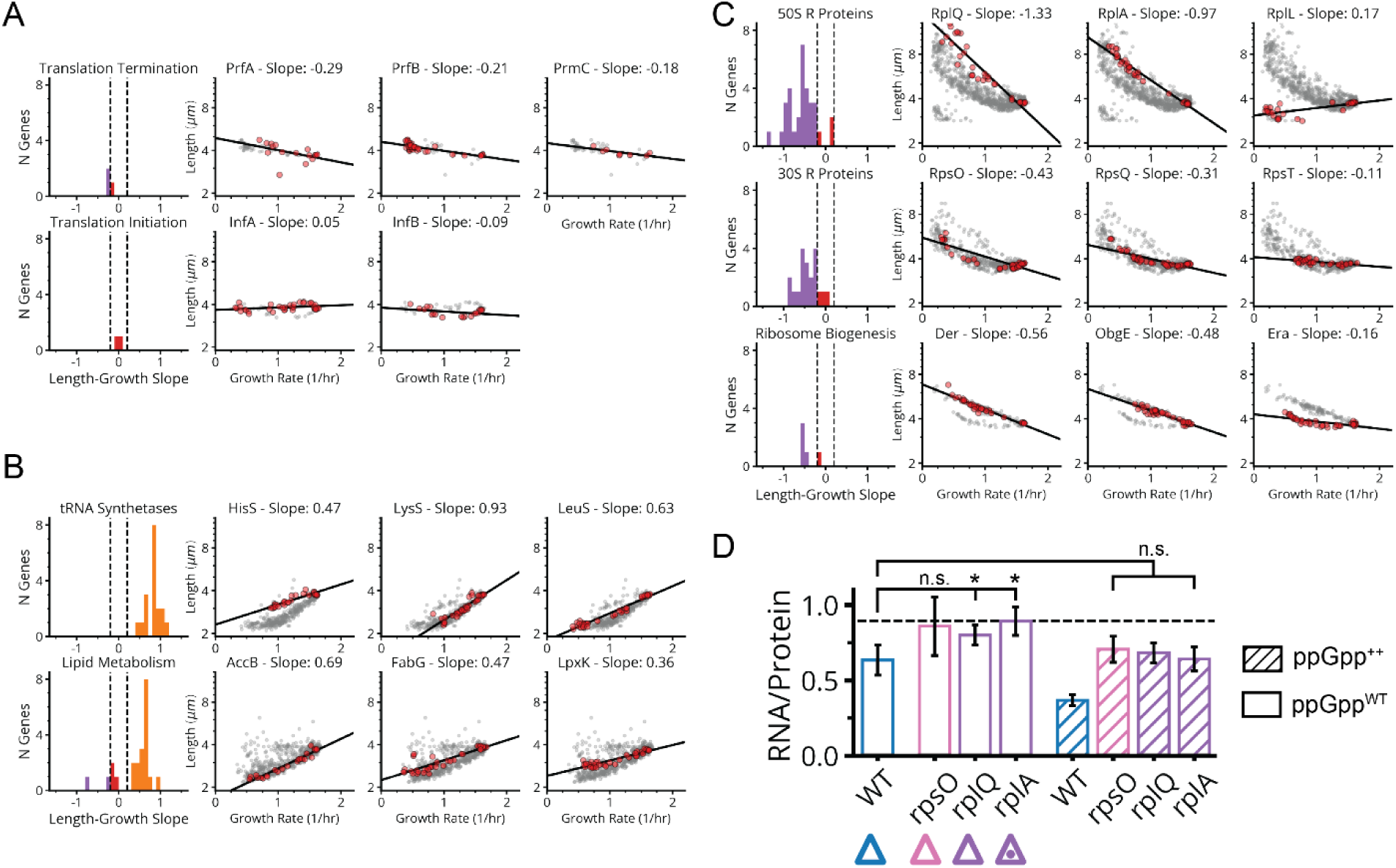
Growth-Size Scaling Analysis for Translation-Related Knockdowns. **a,** Left, histograms of log2 length vs growth rate slope for indicated gene groups including translation termination and initiation factors. Same color scheme for slopes as in Fig. 5a. Right, select log2 length vs growth rate linear fits to groups of sgRNAs targeting single genes belonging to the same gene class. **b,** Same as in panel a, with indicated gene groups including tRNA synthetases and lipid synthesis factors. **c,** Same as in panel a, with indicated gene groups including ribosomal proteins and ribosome biogenesis factors. **d,** RNA/Protein ratio after 6 hours CRISPRi depletion of indicated genes, with triangles matching library measurements indicated in Fig. 5b. WT denotes the EV sgRNA control. Unhatched bars indicate cells uninduced for the hyperactive (p)ppGpp synthase RelA* and hatched lines indicate +25uM IPTG to induce RelA* expression, resulting in ectopic synthesis of (p)ppGpp. Error bars denote 2σ_SEM_ range. Bars indicated with “*” exhibit a significant increase in RNA/Protein relative to WT (p<0.05). Dotted line indicates the maximum RNA/Protein ratio (∼0.89) reported in a previous study limiting protein synthesis by chloramphenicol.^72^

**Extended Data Fig. 12:**
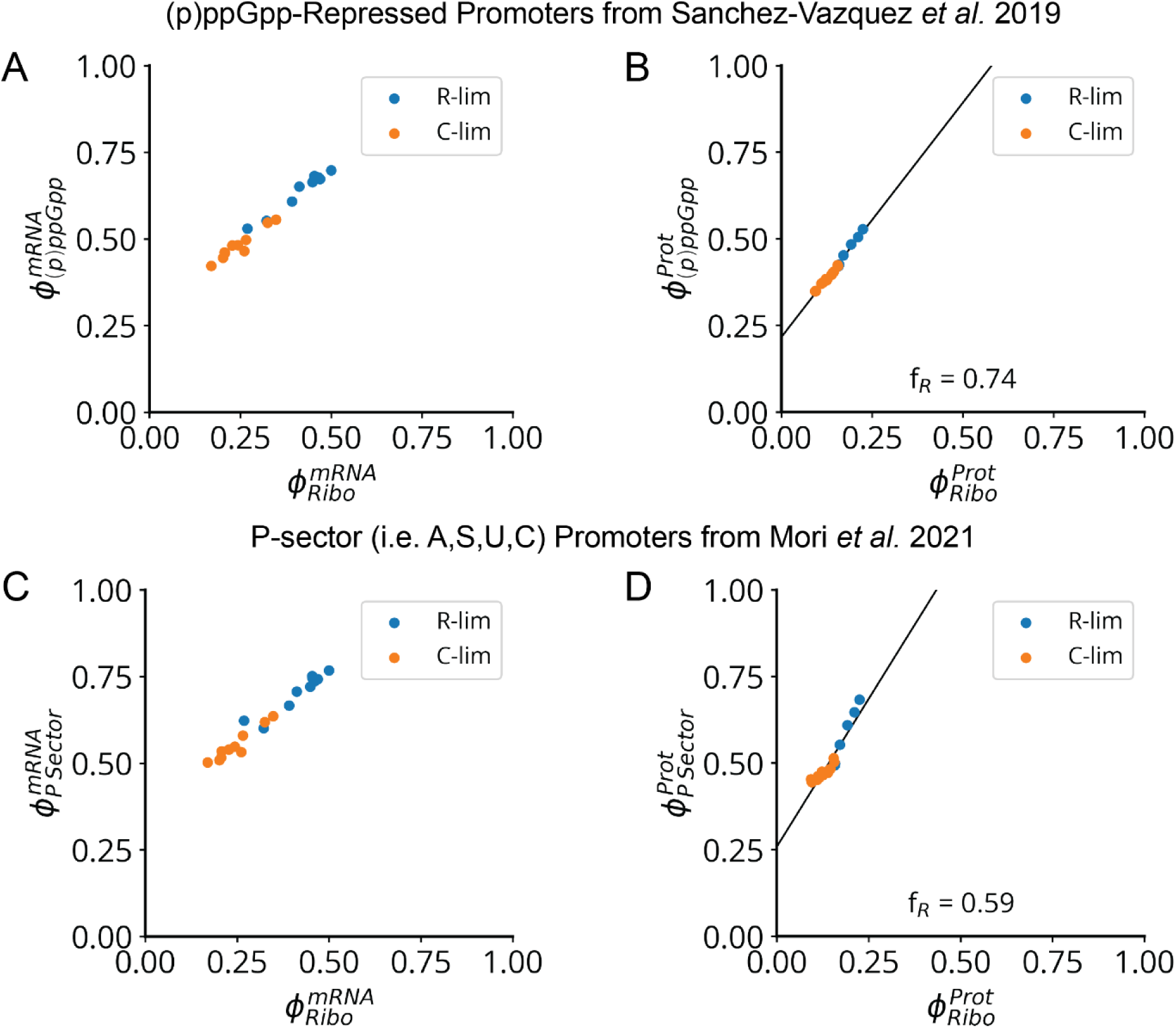
Ribosomal Protein Expression Varies Linearly with the (p)ppGpp Regulon. **a,** Measured ribosomal protein transcriptome fraction vs (p)ppGpp-repressed transcriptome fraction, under limitation of glucose import by PtsG titration (orange points) or translation inhibition by sub-lethal chloramphenicol (blue points). Transcriptome data from Balakrishnan and Mori et al. 2022.^58^ List of (p)ppGpp repressed promoters from Sanchez-Vazquez et al. 2019.^54^ **b,** Measured ribosomal proteome fraction vs (p)ppGpp-repressed proteome fraction, with the same conditions as in panel a. The ribosome fraction of the variable component of the (p)ppGpp regulon f_R_ – defined in Supplementary Note S2 – is measured as the slope of this relationship. Proteome data from Mori et al. 2021.^57^ **c-d,** Same plots as panels a-b, but for genes belonging to proteome sectors – identified in Mori et al. 2021^57^ – which are co-regulated with ribosomal proteins (i.e., the A,S,U,C sectors), which are collectively referred to as the (p)ppGpp-repressed or “P-sector”.

**Extended Data Fig. 13:**
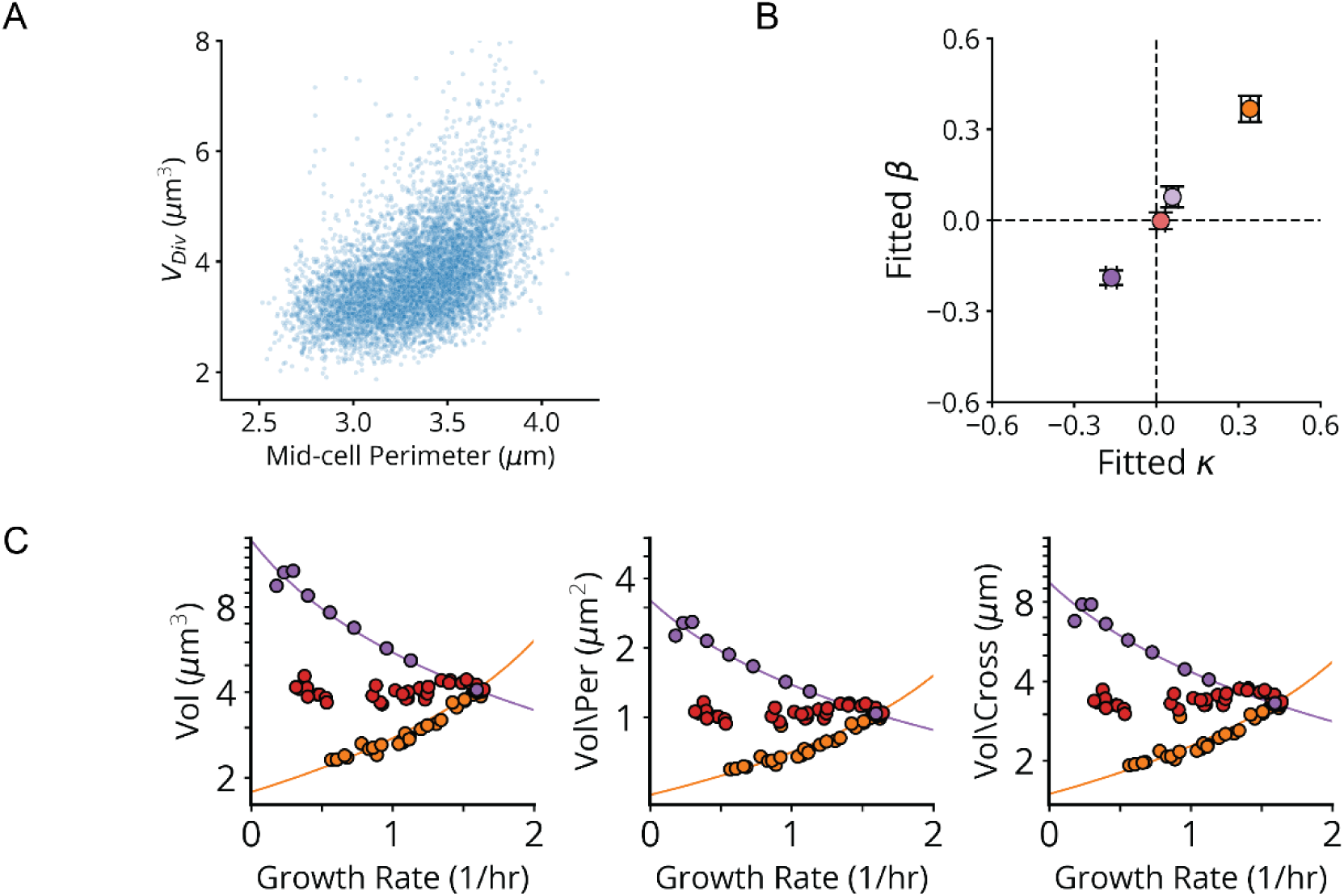
Strict Proportionality of Ribosome and Volume Scaling Constants. **a,** Single-cell measurements of division volume vs mid-cell perimeter at division, between 6-8 hours post-induction in lDE26. Points are only shown for library variants targeting members of the Rod complex (*mrdA*, *mrdB*, *mreB, mreC, mreD, rodZ*) or UDP-GlcNAc synthesis (*glmS*, *glmM*, *glmU*), which primarily impact cell width through reduced cell wall synthesis. **b,** Measured ribosome-growth scaling constants β vs volume-growth scaling constants κ, for *pheT* (orange), *infA* (red), *rplL* (light purple) and *fusA* (dark purple) knockdown series. Derived from scaling data in Fig. 5c,d. See Supplementary Note S2 for definitions of β and κ. **c,** Same fit of volume vs. growth rate as in Fig. 7d (left) as well as equivalent fits where volume is normalized by the measured mid-cell perimeter (center) or the mid-cell cross section (right).

**Extended Data Table 1:** Measured Library Statistics. Genotype and imaging information for each named MARLIN library. For extended library genotype information, see SI. Libraries include either pooled GFP variants or CRISPRi sgRNAs (“Library Type”). Libraries may include a fluorescent nucleoid marker (“*hupA-mCherry*”) or *relA*/*spoT* deletions (“*relA*” and “*spoT*”). Imaging was conducted with either 20x or 40x air objectives (“Magnification”). The number of measured lineages and cells for each library is indicated, except for lDE11 which was not measured with microscopy.

| Library | Library Type | *hupA-mcherry*? | *relA* | *spoT* | Magnification | # Measured Lineages | # Measured Cells |
| --- | --- | --- | --- | --- | --- | --- | --- |
| IDE11 | GFP | No | *WT* | *WT* | NA | NA | NA |
| IDE15 | GFP | No | *WT* | *WT* | 20x | 1102828 | 55133453 |
| IDE20 | CRISPRi | No | *WT* | *WT* | 20x | 4067090 | 1572367044 |
| IDE26 | CRISPRi | Yes | *WT* | *WT* | 40x | 169479 | 57120927 |
| IDE28 | CRISPRi | Yes | $\Delta relA782$ | *WT* | 40x | 119231 | 37559807 |
| IDE30 | CRISPRi | Yes | $\Delta relA782$ | $\Delta spoT(2-696)$ | 40x | 133602 | 51651391 |

## Methods

### Barcode Assembly

Our barcode library consists of a set of plasmids, with each plasmid containing a DNA sequence encoding an RNA barcode. Each barcode contains a set of 30 unique binding sites representing a 30-bit word. Each binding site is 20 bp long and consists of one of two possible sequences, “0” or “1”. If the site consists of sequence “1”, it will be complementary to a cognate FISH probe for that site, while if it consists of sequence “0” it will not bind that cognate probe. These readout sequences were designed to maximize hybridization rates and minimize crosstalk as described previously.^3,24^

Barcodes were generated by cycled ligation assembly (CLA) of pools of 3-bit blocks, each consisting of all 2^3^ possible block sequences.^73^ Both strands of dsDNA coding for each 3-bit block were synthesized as single stranded oligos and pooled to a final concentration of 50 nM for each oligo. These oligos were then phosphorylated by T4 PNK (44 µL oligo pool, 5 µL T4 ligase buffer, 2 µL PNK (NEB, M0201S)). Following this, the phosphorylated oligo blocks were assembled, in proper order, using a mix of “Scaffold Oligonucleotide Connectors” or SOCs, which are unphosphorylated oligos designed with scaffold sequences complementary to the 3 and 5 prime ends of consecutive blocks (4 µL phosphorylation mix, 2 µL 100 nM SOC mix, 2 µL Ampligase buffer, 2 µL 5 µM/µL Ampligase, 10 water). Thermocycling protocol was 95 °C for 2 min, then 60 cycles of 95 °C for 10 s, 60 °C for 30s and 66 °C for 60 s.

To generate barcodes for insertion into plasmids, we diluted the CLA reaction five-fold and amplified full length barcodes using a 100 µL Phusion polymerase (NEB, M0531S0) reaction with 2 µL of diluted CLR template, for 20 cycles, using primers oDE311 and oDE201. An additional 10 µL of 100 µm oDE201 primer (10X) was added to the reaction, which was followed by a single additional thermocycle. Notably, oDE311 contained a random 20bp N-mer sequence such that the final barcode library also included a (20-mer) unique molecular identifier (UMI). 5 µL of this reaction was inspected on an agarose gel for products of the expected length. If non-specific bands were present, reactions would be repeated with fewer cycles. Acceptable products were purified (Monarch PCR & DNA Cleanup Kit, T1030S) and inserted into a oDE202 and oDE253 primer-amplified pDE47 backbone using isothermal assembly (NEB NEBuilder HiFi DNA Assembly Master Mix, E2621L). Assembled barcode plasmids contained a pBR322 origin, a kanamycin resistance gene, and a T7 promoter controlling barcode expression. These assemblies were then purified (Monarch PCR & DNA Cleanup Kit, T1030S), eluted in 6 µL water of which 2 µL was transformed by electroporation into *E. coli* (DH5G Gold Electrocompetent *E.coli*, Genlantis or NEB 10-beta Electrocompetent *E. coli*). 1 mL SOC was added after electroporation and cells were recovered at 37 °C with shaking for 1 hour. The SOC culture was then diluted in 50 mL of LB broth with 50 µg/mL Kanamycin and cultured at 37 °C 220 rpm in a shaking incubator overnight. In parallel, samples of the SOC culture were dilution plated on LB plates with 50 µg/mL Kanamycin to check for adequate transformation efficiency (>10 million CFU/mL). The following day, DNA from the liquid culture was extracted (GeneJET Plasmid Midiprep Kit, K0482) producing the barcode library.

### sgRNA Cloning and Library Assembly

To create a mismatch-CRISPRi library targeting essential genes, we began by selecting all sgRNA sequences in a previous genome-wide CRISPRi screen in *E. coli*.^2^ We considered as essential all genes which exhibited a log fold change < −5.57 at the final timepoint in the original screen or were annotated as essential in a separate TraDIS screen.^1^ For each initial sgRNA, the first member of a mismatch series was generated by mutating the 5’ base pair of the targeting sequence. The second member was then made by further mutating this single-mismatch sequence at the base pair just 3’ to the original mutation. This was continued up to 10 consecutive mutations to generate a total of 11 sgRNAs per target site. 40 control sequences following the design rules of the original CRISPRi screen^2^ and containing less than 8bp of complementarity to any sites in the genome were also included, as well as a handful of sgRNAs from the previous screen targeting non-essential genes.

sgRNA targeting sequences were ordered as an oligo pool from Twist, resuspended to a final (total) concentration of 100 nM in TE buffer. The pool was amplified using a 50 µL Phusion polymerase (NEB, M0531S0) reaction with 1 µL of a 1:250 dilution of the 100 nM oligo pool and oDE418/oDE433 primers, for 8 cycles. The PCR product was purified (Monarch PCR & DNA Cleanup Kit, T1030S) and subsequently digested using BbsI-HF (NEB, R3539S) for 30 mins at 37 °C. 1 µL of 5M NaCl to condition the buffer for a subsequent digestion with BsmBI-v2 (NEB, R0739S) for 30 mins at 55 °C. After heat inactivation at 80 °C for 20 mins, the digested product was purified with a low molecular weight protocol (Monarch PCR & DNA Cleanup Kit, T1030S).

The sgRNA plasmid backbone (pDE93) contains a constitutive sgRNA expression cassette, a pBR322 origin and a Kanamycin resistance gene. This plasmid was digested with BbsI-HF (NEB, R3539S) and CIP treated (NEB, M0525S) to prevent backbone relegation at 37 °C for 1 hour. After heat inactivation at 80 °C for 20 mins, the digested product was run on an agarose gel and the band at the expected backbone size gel purified (Monarch DNA Gel Extraction Kit, T1020L). The digested sgRNA library was then ligated into the plasmid backbone using a 50-100 µM T4 DNA ligase reaction (NEB, M0202S) at RT for 30 mins. The reaction was then purified (Monarch PCR & DNA Cleanup Kit, T1030S) and eluted in 7 µL of water, of which 2 µL was transformed into electrocompetent *E. coli*, recovered in SOC, cultured overnight, and the library was extracted according to the same protocol as with the barcode library.

### Barcoding Library

To assemble our variant libraries with barcodes, we digested both our variant library, barcode plasmid library and a destination vector with BsaI-HFv2 (NEB, R3733S) for 1 hour at 37 °C. The digests were then run on an agarose gel and bands were gel purified corresponding to the cassettes of the two libraries and the backbone of the destination plasmid (p15A ori, Chlor-R). Finally, the two libraries were combinatorially ligated using a 40 µL T4 DNA ligase reaction at RT for 30 mins, purified and eluted in 10 µL. 1 µL of this elution was transformed into electrocompetent *E. coli* with the relevant genetic background and recovered in SOC according to the same protocol as with the barcode library.

To test transformation efficiency, the recovery culture was dilution plated on LB plates supplemented with 25 µg/mL Chloramphenicol, cultured overnight at 37 °C and plating efficiency (CFU/mL recovery mix) measured the next day. Transformation and recovery were then repeated, followed by dilution and plating of the recovery mixture with a target of ∼10000 CFU/plate, onto 20-30 plates. An additional dilution series was also performed on this day to measure the efficiency of this transformation. Plates were cultured overnight at 37 °C and the library bottlenecked to the desired complexity by solubilizing colonies on a number of plates corresponding to the desired CFUs, in EZ Rich Defined Medium (EZRDM) (Teknova, M2105) supplemented with 25 µg/mL Chloramphenicol. The OD600 of the resulting culture was measured and further diluted in media to desired OD600 of 5. Finally, the diluted culture was diluted 1:1 in 50% glycerol, separated into 100 µL aliquots and stored at −80 °C for storage. The residual culture plasmid was extracted (GeneJET Plasmid Midiprep Kit, K0482) for later use in sequencing of the barcode to variant map.

### Library Sequencing

According to our procedure, barcodes are randomly ligated to variants, so we use sequencing to establish the lookup table from each barcode to its corresponding variant. Since the length of the variant-barcode cassette is long (>700bp), we used Nanopore long read sequencing to acquire reads spanning the entire construct and leveraged our barcodes to bin reads by library members.

Library plasmid cassettes were first amplified in a 400 µL Phusion polymerase PCR with primers oDE154 and oDE201 for 12-15 cycles, depending on how many cycles were needed to yield sufficient material. PCR products were then purified by AMPure bead cleanup (Beckman Coulter, A63881). Amplicons were then prepared for sequencing using a standard protocol from Oxford Nanopore Technologies (ONT) which included FFPE DNA Repair (NEB, M6630S), End Repair/dA-Tailing (NEB, E7546S), and ligation of adaptors from the ONT SQK-LSK109 ligation sequencing kit. Prepared amplicons were then run on a ONT MinION with either R9.4.3 or R10.3.1 pores. If necessary to achieve adequate depth, multiple sequencing runs were conducted on a single library and the reads pooled.

Raw reads were processed using high accuracy calling (HAC) with the Guppy basecaller. Following this, reads were aligned to a graph genome reference (gfa) specifying the combinatorial structure of our barcodes using GraphAligner.^74^ The alignment, corresponding to a traversal through this barcode graph, provided the barcode contained in each read, allowing us to subsequently group reads based on barcode. Grouped reads were then aligned to a reference using Minimap2^75^, also specified by the graph alignment, and finally used to generate a barcode-variant consensus using Medaka. This analysis was parallelized using the Snakemake^76^ module in Python (https://github.com/paulssonlab/marlin-nanopore), enabling us to scale this approach to large libraries.

### MARLIN Phenotyping

Prior to each experiment, 100 µL library aliquots of *E. coli* were inoculated into 5 mL cultures of EZRDM with 25 µg/mL Chloramphenicol and grown at 37 °C 220 rpm on a shaking incubator to saturation (6-8 hours). Prior to loading cells, the microfluidic device was primed by injecting ∼200 µL of faCellatite BIOFLOAT^TM^ solution (faCellitate, F202005) and allowing it to adsorb at RT for 30 mins. 3 mL of saturated library culture was concentrated 10X by spinning in a microcentrifuge at 4000 rcf for 1 min. Supernatant was then collected to leave a residual ∼300 µL of liquid before resuspension of the pellet. The concentrated culture was then injected into the microfluidic device to load the library. To ensure maximum loading, the microfluidic was spin loaded using a custom microfuge adaptor, at 650 rcf for 30s, loading the cells into chambers in the downward orientation. The orientation of the device in the adaptor was then reversed and spun at 650 rcf for 30 s, to load chambers in the upward orientation. Once loaded, the device was connected to tubing at the inlet carrying growth medium driven upstream of the device by a peristaltic pump (Longer Precision Pump Co. Ltd, T60-S2&WX10-14), while the outlet was connected to a waste line. The device and feeding lines were then transferred to the microscope for imaging.

Imaging was conducted on a Nikon Ti-2 microscope, with the entire stage enclosed in an incubator (Okolabs) to maintain a constant temperature of 30 °C (lDE20, lDE28 and lDE30) or 37 °C (lDE15). The device was imaged using either a Nikon Plan Apo Lambda 20x Ph2 DM air objective with 0.75 NA (lDE15 and lDE20) or a Nikon Plan Apo Lambda 40x Ph2 DM air objective with 0.95 NA (lDE26, lDE28 and lDE30). All images were acquired using a high pixel density (4.5 µm x 4.5 µm pixel size) sCMOS camera (Photometrics Iris 9) to enable adequate sampling of the 20x objective point spread function. We used an LED lamp (CoolLED pE-100) as a white-light source for phase-contrast imaging and a Spectra III mixed LED/Laser light source (Lumencor) for fluorescence imaging. For most experiments, our light path consisted of a multi-band dichroic (409/493/573/652/759 Brightline, Semrock) and a variable emission filter (519/26, 595/33, or no filter), mounted on an emission filter wheel for fast multi-channel imaging (Finger Lakes Instruments HS-625). For imaging the mKate2Hyb marker in lDE20, we used an alternative configuration with a multi-band dichroic (89403, Chroma) and an 642/80 emission filter.

Before imaging, cells in the microfluidic were allowed to proliferate on-scope for a period of 10-14 hours, under constant flow of media. The phenotyping portion of the experiment then proceeded, with all microscope control and acquisition handled by the Nikon ND Elements software. Depending on the phenotype being measured, different numbers of fields of view and image cadences were used. Imaging of GFPmut2 and mKate2Hyb in lDE15 was conducted every 40 mins for a period of 5 hours and 20 mins, over a grid of 855 fields of view (FOVs). Imaging of mKate2Hyb in lDE20 was conducted every 10 mins for a period of 10 hours, over 629 FOVs. Finally, imaging of mVenus and HU-mCherry in lDE26, lDE28 and lDE30 was conducted every 10 mins for a period of 10 hours, over 325 FOVs. In all cases, the Nikon perfect focus system (PFS) was engaged throughout the experiment to maintain the sample in the focal plane. In all 10 hour CRISPRi experiments, aTc (Sigma, #37919) was added to the flask containing the input growth medium to a final concentration of 0.2 µM at the 2 hour mark, in order to induce dCas9 expression.

### MARLIN Genotyping

Genotyping FISH probes complementary to the MERFISH barcode were ordered from IDT, with the probe DNA conjugated at the 5’ end with a six carbon (C6) linker to a di-thiol group followed by another C6 linker conjugated to a dye. Depending on the experiment, we used probe sets utilizing three of the following five dyes: Alexa Fluor 488, Alexa Fluor 555, Alexa Flour 647, Cy5 and Alexa Fluor 750. Probes were received as lyophilized pellets, resuspended to 100 µM using nuclease-free water, split into a number of strip tubes, and re-lyophilized in 2 nmol aliquots. The dry aliquots were stored at −80 °C for long term storage. 100 µM stocks were prepared from these aliquots by adding 20 µL of nuclease-free water and kept at −20 °C for less than six months or until exhaustion. Working stocks of probe combinations to be used in the same imaging cycle were prepared by adding 6.66 µL each of three 100 µM probe stocks to 180 µL of readout probe buffer (TE buffer with 0.1% Triton X-100) to a final concentration of 3.33 µM per probe. Working stocks were stored at −20 °C for less than two months or until exhaustion. Each round of FISH probing uses 15 µL of this working stalk in a 5 mL hybridization buffer (10% Ethylene Carbonate, 0.125% Triton X-100, 40 U/mL Murine RNAse inhibitor and 20 nM DNA probe in pH 7.0 2X SSC).

Genotyping was conducted immediately after the conclusion of phenotype imaging for each library experiment. Cells in the device were induced for barcode expression for a period of ∼75 mins by the addition of Cuminic acid to a final concentration of 100 µM in the input growth media. During this time, we performed setup of our automated liquid handling system (see extended methods), which enabled automated selection of FISH reagents in conjunction with image acquisition. We controlled this system, in conjunction with the microscope, from a Jupyter notebook making use of both a custom Python module for this purpose (https://github.com/paulssonlab/marlin) and the Python Micromanager API.^77^ During setup, all liquid lines in this system were cleaned using DI water and inspected for adequate flow. Once barcode induction was concluded, we loaded the liquid handler with our FISH reagents and replaced the growth medium line with the liquid handler line, just upstream of the device inlet. The liquid handler proceeded to perform fixation and FISH imaging over the next 16-20 hours.

During FISH genotyping, we first inject a 4:1:1:4 solution of methanol, acetone, 20X SSC and nuclease-free water (50% MeAc) into the device, followed immediately by a 4:1 solution of methanol and acetone (MeAc) to fix the cells. We allow fixation to proceed for 45 mins, then inject 50% MeAc, followed by the hybridization solution containing the first probe set. After allowing probes to hybridize for 30 mins, we inject an oxygen-scavenging imaging buffer solution consisting of 3mM PCA, 0.03% Trolox, ∼0.18 U/mL rPCO, and Trolox-Quinone (>30 µM, prepared according to a previously published protocol^78^) in pH 7.0 2X SSC. Imaging buffer is flowed for an additional 5 mins to clear any bubbles from the device and then FISH imaging proceeds. During each round of imaging, all FOVs phenotyped previously are imaged in phase-contrast for lineage-detection purposes and three additional fluorescence channels, corresponding to each probe dye. After all channels are imaged, we inject a cleavage buffer (33.3 mM TCEP in pH 7.0 2x SSC) containing a reducing agent to break the di-thiol bond linking dyes to DNA probes, eliminating the fluorescence signal from this round of imaging. To continue genotyping, we repeat this protocol from the hybridization step, for the next set of probes. We repeat 10 such cycles, reading out 3 DNA probes in each, for a total of 30 probe readouts, completing the experiment.

### Mother Machine Isolate Imaging

Mother machine experiments with single isolates were performed using the same imaging setup as described in “MARLIN Imaging”. Starter cultures for strains to be loaded into the device were generally prepared 12-16 hours in advance, in the same manner as described in “Strains and Growth Conditions”. All CRISPRi validation strains were imaged according to the same schedule and under the same conditions as described for lDE20.

### Strains and Growth Conditions

Cells were initially inoculated into 5 mL EZ Rich Defined Medium with appropriate antibiotics in a test tube from a single colony on an agar plate, struck within 2 weeks of the experiment. Cells were then cultured for 14-16 hrs at 37 °C in a shaking incubator (220 rpm), until saturation. Cells were then diluted 10^-7^ into 5 mL EZRDM with appropriate antibiotics, as a pre-culture, and incubated for an additional 10-12 hrs at 30 °C in a shaking incubator (220 rpm). We then measured OD600 for the pre-cultures and proceeded to dilute each into 25 mL of inductive media (EZRDM + 0.2 µM aTc) with or without 1000 µM IPTG, depending on the experiment. Cultures with OD600 over 0.5 were discarded. Induced cultures were subsequently incubated for 6 hrs at 30 °C in a shaking incubator (220 rpm). Cultures with OD600 between 0.1 and 0.4 at this time were further processed to measure the RNA/Protein ratio (see “RNA/Protein Measurement”) and the (p)ppGpp concentration (see “ppGpp/GTP Sample Harvest Protocol”).

To prepare fixed samples for agar pad imaging the culture protocol was similar, with starter cultures and pre-cultures prepared in the same manner. Pre-cultures were then diluted in 2.5 mL of inductive media (EZRDM + 0.4 µM aTc) and incubated at 30 °C in a shaking incubator (220 rpm) for 6 hours. The OD600 of the cultures were measured and the cultures processed if their OD600 was between 0.1 and 0.4. Subsequently, 360 µL of 32% PFA was added to each tube, which was then placed back into the incubator for a total of 30 minutes. Cells were then collected by centrifugation of the fixed culture at 4000 g for 5 mins, at RT in 10 mL falcon tubes, removing the supernatant and resuspending the pellet in 500 µL of PBS, this centrifugation step was repeated with a resuspension volume of 500 µL and then once more with a final resuspension volume of 250 µL. Samples were then stored at 4 °C until staining and imaging.

## Data availability

Raw sequencing reads for Nanopore and NGS libraries have been deposited in the NCBI BioProject database under accession number PRJNA1205775. Processed data are available on Zenodo (https://doi.org/10.5281/zenodo.14537796). An archived release of the TrenchRipper module version used for this publication is also available on Zenodo at https://doi.org/10.5281/zenodo.14552982. Other source code for image analysis, sequencing analysis, and figure generation are available on GitHub (https://github.com/paulssonlab/2025_Eaton_optical-pooled-screen). Raw or processed image data are available upon request.

## Acknowledgements

We are grateful for the gifts of bacterial strains and plasmids from the labs of Richard Murray, David Bikard, Christopher Voigt, Stanley Qi, Kathleen Matthews, Drew Endy, Keith Shearwin, Greg Bokinsky, Philippe Cluzel, Donald Court, James Keck, Michael Baym, and Thomas Bernhardt. Mass spectrometry measurements and consultation were performed at the Harvard Center for Mass Spectrometry. Fluid handling systems were built at the Research Instrumentation core at Harvard Medical School. All Computational work was performed on the O2 cluster supported by the Research Computing Group at Harvard Medical School. D.S.E. is grateful for formal and informal feedback on this work from Andrew Murray, Noah Olsman, Gene-Wei Li, Rebecca Ward, Thomas Bernhardt, and Maximillian Marin. This work was supported by National Institute of Health grant R01GM081563, Defense Advanced Research Projects Agency grant W911NF-19-2-0018, and Advanced Research Projects Agency for Health grant 1AY2AX000005-01. D.S.E., C.S., L.G.L., and J.Q.S. acknowledge support from the Systems, Synthetic and Quantitative Biology PhD program training award (T32GM135014). D.S.E. was supported in part by the NSF-Simons Center for Mathematical and Statistical Analysis Biology at Harvard (award number #1764269), and the Harvard Quantitative Biology Initiative.

## Author Contributions

D.S.E., J.R.M. and J.P. conceived of and designed the study. D.S.E., C.S., and L.G.L. carried out the principal experiments and data analysis. J.Q.S. and Y.G. designed and fabricated the microfluidic master molds. B.R.W. assisted with automation of liquid handling. D.S.E., C.S., L.G.L. and V.H. engineered the genetic strains made for this study. J.R.M., J.P. and E.C.G. supervised the study. D.S.E., Q.A.J., J.R.M. and J.P. wrote the paper.

