## Supplementary Information for "Essentialome-Wide Multigenerational Imaging Reveals Mechanistic Origins of Cell Growth Laws"

### Supplementary Information for Eaton et al. 2025

### Supplementary Methods

#### Design and Fabrication of High-throughput Mother Machine

*Design Principles and Details*

Here, we describe the design and fabrication of our 1.6 million lineage mother machine. Previous high-throughput mother machine designs scaled to ~100k lineages[1] in single contiguous feeding channels but suffered from problems preventing their use at the million-lineage scale. First, the entire device could be disabled by small defects in fabrication or contaminating particles in the flow path. In addition, the high pressure required to sustain flow across the long feeding channel led to high rates of failure in upstream fluid lines as well as debonding of the PDMS device from the coverslip. Both issues are ameliorated by adopting a parallel feeding channel design, facilitated by fan-out and fan-in manifolds at the inlet and outlet, respectively. The exact layout of this device required extensive considerations including computing flow uniformity across the parallel lanes[2], avoiding geometries that promote biofilm formation, and computing optimal packing of lineages within single FOVs based on our optical configuration. We implemented a software package, snakenbake, to perform this optimization and output mother machine designs under user-specified constraints.[3]

*Silicon Master Mold Fabrication*

The master mold for the microfluidic chip was patterned on a 3-inch silicon wafer using a combination of direct photolithography and etching techniques. The design has two basic features: feeding channels and growth trenches, which were prepared separately. The etching process was specifically employed to create the bacteria growth trenches within a SiO2 layer, whereas the feeding channels were formed by patterning SU-8 photoresist. Initially, a 2 µm thick layer of SiO2 was deposited on the silicon wafer using a dual-frequency powered parallel electrode reactor (STS PECVD). Following this, a 2.5 µm thick layer of SPR 220-3.0 (MEGAPOSIT™) photoresist was spin-coated onto the wafer. HMDS was pre-applied as an adhesion promoter. The wafer underwent a pre-exposure bake at 115°C for 1 minute to evaporate the solvent and stabilize the photoresist. A negative pattern of the chip design, specifically the bacteria growth trenches, was then directly exposed onto the photoresist using a maskless aligner MLA150 (Heidelberg Instruments) with a 405 nm wavelength light source and a dose of 195 mJ/cm$^2$. Post-exposure baking was conducted at 90°C for 1 minute to accelerate the photochemical reaction. The developed pattern was created by immersing the wafer in MF®319 developer (Microposit®). To etch the SiO2 layer, a ULVAC NLD-570 deep oxide etcher was employed, removing 1.4 µm of SiO2 from areas not protected by the photoresist, using a gas mixture of Ar (90 sccm) and C3F8 (10 sccm). Following the etching process, the photoresist was stripped by soaking the wafer in Remover PG (Kayaku™) at 80°C overnight. To make 70 µm thick feeding channels, a 70 µm thick layer of SU-8 2050 photoresist was spin-coated on the wafer. The pre-exposure bake consisted of three steps to prevent photoresist cracking: 65°C for 2 minutes, 95°C for 7 minutes, and then 65°C for 2 minutes with ramped temperature increases and decreases. The pattern for the feeding channels was directly exposed onto the photoresist using the MLA150, with a specified energy dose of 2100 mJ/cm2 and a wavelength of 375 nm. Post-exposure baking was performed under the same conditions as the pre-exposure steps. The SU-8 2050 photoresist was then developed using SU-8 developer (Kayaku™), and to further cross-link the photoresist, the master mold silicon wafer was hard-baked at 130°C for 20 minutes.

*PDMS Cast Fabrication and Coverslip Bonding*

To fabricate each PDMS device, we used the following protocol, along with the previously prepared master mold. First, a 10:1 mixture of silicon base to curing agent (Sylgard 184 elastomer) was homogenized using a planetary centrifuge (Thinky AR-100). The mixture was poured onto the master mold, degassed under vacuum for 15 min, and baked for 30 min to cast the device. Removing the cast PDMS from the master mold, we then used a 700 µm diameter coring needle to punch out inlet and outlet ports before cutting the device out of the cast with a razor blade. The device and coverslips were heated to 30°C or 37°C to match the temperature of the experiment, which prevents warping, and promptly plasma treated (10 s at 75 W with O2 pressure between 150-300 mTorr), bonded, and baked at 95°C for 1 hour. Chips were then stored at room temperature until use.

**MARLIN Library Construction Guidelines**

Here, we detail how to choose the number of barcoded library variants to bottleneck at the point of library creation. We include considerations related to genetic variant dropout as well as required sequencing depth.

*Minimum Complexity to Avoid Dropout*

When we ligate FISH barcodes to our underlying variants of interest, we must transform and recover sufficient colonies to ensure that every genetic variant is recovered in at least one colony. Should we want the frequency with which variants appear in the library to be more narrowly distributed, we would desire to harvest even more colonies than this. At the same time, the number of sequencing reads we require to characterize the map from barcode to genetic variant will scale with the number of recovered colonies. Given this trade-off with sequencing depth, we choose to bottleneck our libraries to the minimum number of colonies required to avoid significant dropout.

To determine the rate of dropout expected for a given bottleneck, we begin with a binomial model of variant frequency. If we randomly pick $N$ library plasmids, we obtain for a given variant $v_j$ in a library with $V$ variants:

$$v_j \sim Bin(n = N, p = V^{-1})$$

since $V^{-1}$ is typically very small and N is very large ($V^{-1} < 0.002$ and $N > 13000$, for all libraries in this paper), we may approximate this as a Poisson distribution:

$$v_j \sim Pois\left(\lambda = \frac{N}{V}\right)$$

We can use this form to bound the rate of dropout by bounding the probability of zero events under this distribution, and determining the corresponding constraint on the bottlenecking rate:

$$P\left(v_j = 0\right) < \epsilon \Rightarrow e^{-\frac{N}{V}} < \epsilon$$

Rearranging this, we now know the minimum number of colonies $N$ required to bound the dropout rate $\epsilon$:

$$N > -V \ln \epsilon$$

A reasonable choice for $\epsilon$ is 0.01 or a dropout rate of 1%, which works out to:

$$N > 4.6\mathrm{V}$$

Thus, a good rule of thumb for bottlenecking libraries after barcoding is to retain at least 5 times the number of underlying variants of interest in the library. The higher above this number we choose to be, the lower the CV of the resulting variant frequency distribution, owing to the scaling of the Poisson CV with $1/\sqrt{\lambda}$ or, equivalently, $1/\sqrt{\frac{N}{V}}$.

*Maximum Complexity to Sequence Deeply*

While increasing the number of transformants in our library minimizes dropout, it also increases the read depth required to accurately sequence each of our library variants. Suppose each library variant produces $r_i$ reads during sequencing. Then, given that these reads possess a degree of error, there will be a threshold read depth $R_{thr}$ which we will require to achieve a more accurate consensus sequence within our error tolerances. Assuming that variant sequences are sampled randomly, we expect that:

$$r_i \sim Bin(n = R_{tot}, p = N^{-1})$$

Since we need at least $R_{thr}$ reads to get an accurate consensus, we may define the dropout rate as the lower tail of this distribution $P(X < R_{thr})$. Empirically, we find that $R_{thr} = 15$ produces consensus sequences that are sufficiently accurate for both GFP and sgRNA variant assignment (Extended Data Fig. 1a-c,e). For an average MinION sequencing run, we produce $\sim 10^7$ reads for a 1 kb construct. Fixing $R_{tot}$ and $R_{thr}$ to these values, we can compute our dropout rate $P(X < R_{thr})$ for a given number of barcodes. Thus, given these parameters, $P(X < R_{thr}) < 0.001$ for libraries with <250k barcodes. Knowing this, we chose to bottleneck to ~128k colonies for lDE15, ~134k colonies for lDE20, and ~14k colonies for lDE26. In all cases, the resulting dropout from insufficient sequencing should be extremely low, though in practice variant frequency biases inherited from variability in array synthesis efficiency and fitness differences can confound this.

**GFPmut2/DarkGFP Error Rate Measurement**

Here, we detail two approaches to quantifying the FISH-attributable error rate in GFP variant calling for the library lDE15. In the first method, we extract the FISH-attributable error from the discrepancy between the total library error and that of a library subset containing only highly distinct barcodes. For the second method, we determine the expected rate of FISH decoding error from the complementary rate at which barcodes which do not belong to our library members are read. We show that both approaches suggest FISH barcode accuracy in the neighborhood of 99.9%. We further show from complementary characterization of our sequencing approach that the majority of our overall error rate (0.89%) is likely inherited from sequencing errors.

*Error Rate Decomposition*

We measured the overall error rate of GFPmut2/DarkGFP variant calling by first measuring the GFP signal from each lineage during phenotyping, establishing a signal threshold to distinguish the dim DarkGFP from the bright GFPmut2 (Extended Data Fig. 2d). We compared this determination to the GFP variant inferred from the subsequently measured FISH barcode and considered any case where these assignments disagreed to be an error. We consider this overall error rate to be composed of instances in which the FISH barcode is incorrectly read ($Err_{FISH}$) and instances in which the GFP variant is incorrectly determined from sequencing ($Err_{Seq}$):

$$Err_{Total} = Err_{Seq} + \frac{1}{2} Err_{FISH}$$

While other effects such as poor quantification of GFP signal during imaging and sequencing errors in barcode identification may also contribute to the overall error, we will show that the preceding two effects account for the vast majority of our error. Importantly, we halve the contribution of $Err_{FISH}$ to the overall error rate, reflecting our assumption that a mistaken barcode assignment results in a change in the GFP variant assignment only half of the time (Extended Data Fig. 2e).

*Error Based on Highly Distinct Barcodes*

Because $Err_{FISH}$ reflects the underlying error of our barcoding approach, while $Err_{Seq}$ is an application-dependent function of sequencing depth, we were interested in decomposing these two error types from each other. We reasoned that if we only considered FISH barcodes that were highly distinct from each other (i.e., with a large Hamming distance), we could virtually eliminate the possibility for errors in FISH decoding. For instance, for a barcode library with each member separated by a Hamming distance of 4, even a massive bit-wise error rate of 10% would produce an overall FISH error of 0.01%. When we computationally subsample our library to include only barcodes separated by a minimum Hamming distance of 4, we decrease the error rate from 0.89% to 0.8% (Extended Data Fig. 2f). If we assume that at a Hamming distance of 4, $Err_{FISH} = 0$, it follows that $Err_{Seq} = 0.8$. Since $Err_{Seq}$ should be invariant under different Hamming distance bottlenecks, it follows from our error decomposition that $Err_{FISH} = 0.18$ for the overall library. So, this first procedure estimates the accuracy of FISH decoding to be ~99.8%.

*Error Based on Out of Library Barcodes*

We adapted a complementary approach to estimating the FISH decoding error rate, based on a similar argument made in Emanuel *et al.* 2017.[4] Briefly, this technique leverages the relationship between the rate of FISH misidentification and the rate of detecting barcodes that are not present in the library. First, we assume that a misread barcode will map back into the library with a probability corresponding to the fraction of all possible barcodes occupied by the library. That is, for a library of size $|I|$:

$$P(L|E=k)=\frac{|I|}{|O|+|I|}$$

where $|O|$ is the number of possible barcodes not represented in the library and $P(L|E=k)$ is the probability of mapping back into the library for FISH readouts with $k$ bit errors, with $k>0$. It follows that the probability of reading a barcode that is not in the library $P(NL|E=k)$ is just $1-P(L|E=k)$ or, equivalently, $\frac{|O|}{|O|+|I|}$. We can express the total probability of reading a barcode not in the library as the sum of probabilities with particular numbers of bit errors $k$:

$$P(NL)=\sum_{k=1}^{N}P(NL|E=k)P(E=k)=\sum_{k=1}^{N}\left(\frac{|O|}{|O|+|I|}\right)P(E=k)=\left(\frac{|O|}{|O|+|I|}\right)P(E\geq 1)$$

We can then rearrange this relationship to ascertain the probability of possessing any number of FISH errors, in terms of the probability of detecting barcodes that are not in the library:

$$P(E\geq 1)=\left(\frac{|O|+|I|}{|O|}\right)P(NL)$$

Finally, we can express the FISH error rate, $P(L,E\geq 1)$, in terms of FISH detection of barcodes absent from the library:

$$P(L,E\geq 1)=P(L|E\geq k)P(E\geq 1)=\frac{|I|}{|O|+|I|}P(E\geq 1)=\frac{|I|}{|O|}P(NL)$$

Thus, for an N bit barcode:

$$P(L,E\geq 1)=\left(\frac{|I|}{2^N-|I|}\right)P(NL) \qquad (1)$$

To determine our FISH-attributable error rate, we then plug in the values from our lDE15 experiment:

$$P(NL)=0.42 \quad N=30 \quad I=125500$$

which gives $P(L,E\geq 1)=0.005\%$ corresponding to $Err_{FISH}$ in the case of assigning only barcodes with a perfect match. While this results in a very low FISH error rate, the reduction in overall throughput resulting from this strict cutoff is undesirable. As such, in our experiments, we assign decoded barcodes to their closest match in the library, up to a Hamming distance of 1 away from the true barcode sequence. We examine the resulting form of $P(L, E \geq 1)$ for this case in the next section.

*Error Rate for H(1) Assignments*

Since most of our barcodes are separated by 2 or more bits, we can tolerate some error in our FISH decoding without high risk of incorrectly assigning a barcode to an existing variant in the library. Because of this, called barcodes that do not match a member of the codebook but are within one bit of an existing member, are assigned to that member. We can modify equation (1) for that case, through a similar argument:

$$P(L, E \geq 2) = \frac{|I_H|}{|O_H| + |I_H|} P(E \geq 2) = \frac{|I_H|}{|O_H|} P(NL)$$

where $I_H$ and $O_H$ are modified to represent that the codebook effectively contains both true barcodes and all barcodes H bit flips away from those barcodes. Note that we are assuming that in all cases $P(L, E = 1)$ corresponds to a true barcode assignment, which will be the case if all barcodes are at a Hamming distance of 2 or greater, which is the case for \textgreater 99% of barcodes in lDE15. For an N bit barcode this formulation results in:

$$P(L, E \geq 2) = \frac{|I|N^H}{2^N - |I|N^H} P(NL) \qquad (2)$$

Plugging in the values from the lDE15 experiment we have:

$$P(NL) = 0.29 \quad N = 30 \quad I = 125500 \quad H = 1$$

which gives $P(L, E \geq 2) = 0.001$ corresponding to $Err_{FISH}$. Thus, this approach to characterizing the FISH error rate suggests that the accuracy of FISH decoding is ~99.9%, which is quite similar to the value of ~99.8% computed using the first approach.

*Error from Sequencing*

We estimated that FISH decoding contributed between 0.1 and 0.18% to the overall error rate, based on our previous calculations. We were interested in determining whether sequencing errors explained a large fraction of the residual error rate. To this end, we deeply sequenced a lower complexity GFPmut2/DarkGFP library lDE11, such that a substantial number of variants were sequenced to a depth of at least 200 reads, using two nanopore flow cell variants, R9.4.1 and R10.3.1. Taking the consensus determined for these deeply sequenced variants as ground truth, we then computationally sub-sampled the number of reads available for each barcode and recomputed a lower depth consensus. We compared our GFP variant call and the barcode UMI sequence from these consensuses to our ground truth to compute UMI (Extended Data Fig. 1a) and GFP call error rates (Extended Data Fig. 1b) at different sequencing depths, for both of our nanopore flow cells. Since we used a R9.4.1 flow cell to sequence the library lDE15, with a minimum depth of 15 reads, we estimated from these results that the GFP calling error we can attribute to sequencing, $Err_{Seq}$, is roughly 0.62%, representing the majority of our residual error. We therefore anticipate that deeper sequencing could increase our overall barcoding accuracy to 99.7%.

**CRISPRi Library Design**

The design of our CRISPRi library lDE20 followed the following procedure, as implemented in lDE20_Design.ipynb in the accompanying data.[5] First, to target essential genes, we began by selecting all sgRNA sequences in a previous genome-wide CRISPRi screen in *E. coli*.[6] We considered as essential all genes that exhibited a log-fold change < -5.57 at the final timepoint in the original screen or were annotated as essential in a separate TraDIS screen.[7] For each initial sgRNA, the first member of a mismatch series was generated by mutating the 5' base pair of the targeting sequence. The second member was then made by further mutating this single-mismatch sequence at the base pair just 3' to the original mutation. This was continued up to 10 consecutive mutations to generate a total of 11 sgRNAs per target site.

We also generated 40 control sequences following the design rules of the original CRISPRi screen.[6] Specifically, we disallowed control sequences with so-called "bad seed" sequences that produce off-target toxicity.[8] We also excluded sequences matching off-target NGG PAM sites by 9 or more nucleotides of perfect identity or more in the seed sequence. In addition, we ensured that these control sequences contain fewer than 8 bp of complementarity to any sites in the *E. coli* MG1655 genome (accession: U00096.3). Finally, we added a handful of sgRNAs from the previous screen targeting non-essential genes.

The design of the CRISPRi library lDE26 followed the following procedure, as implemented in lDE26_Design.ipynb in the accompanying data.[5] To create a low complexity library targeting the same genes in lDE20, we selected sgRNAs for which we observed strong phenotypic effects in our original MARLIN experiments, based on our time-series clustering data. First, we labeled each sgRNA according to its cluster at the low Leiden cluster resolution of 0.2. Then, for each NGG PAM target site, we determined the mismatched sgRNA with the strongest phenotypic effect according to the maximum absolute Z-score (MAZ), over all dimensions and timepoints. With the strongest sgRNA per PAM site selected, we then narrowed down our library further by performing the following algorithm on each set of strong sgRNAs targeting the same gene: (1) group candidate sgRNAs by cluster assignment, (2) for each group of candidates, pick the sgRNA with the highest MAZ. We further included all of the control sgRNAs present in lDE20. This led to a library with a total of 745 sgRNAs.

**Image Analysis and TrenchRipper Pipeline**

All image data in this work was analyzed using our custom Python module, TrenchRipper, which performs cell trap detection, cell segmentation, and lineage tracing.[9] TrenchRipper is optimized for use on high performance computing (HPC), systems and all of our analysis was conducted on the O2 High Performance Compute Cluster at Harvard Medical School.

*Growth and Division Phenotyping*

Briefly, we analyzed our cell phenotype data by first detecting lineage bounding boxes based on the fluorescent segmentation marker in our imaging data. We then composed a local Otsu threshold along with a triangle threshold on this fluorescence data to roughly segment cells. Edges between contacting cells were detected by thresholding a Hessian-transformed image[10], and the resulting mask was used to guide a watershed of the previously determined cell segments. Finally, cell lineages were determined from the segmentation using a modified version of a previously described linear program for use in mother machine imaging.[11] For experiments with a HU-mCherry nucleoid marker, nucleoid segments were determined by performing an Otsu threshold on only the pixels within each cell, using the previously determined cell segment.

Cell length and width were determined from the measured cell area and perimeter, under the assumption that the cell mask takes a stadium (i.e., disco-rectangular) geometry. Cell volume was extrapolated from length and width assuming a spherocylinder geometry. The instantaneous growth rate was subsequently determined from volume measurements by regressing a line to each three timepoint window available within each cell cycle, including extrapolated volumes at cell birth and division in between timepoints.

*MERFISH Genotyping*

For FISH genotyping, we first detected lineage bounding boxes, using the cell traps visible in phase contrast images. FISH signal was then aggregated for each channel and cycle, using the 98th percentile signal within each bounding box, resulting in a single aggregated intensity measurement for each combination of trench, channel, and FISH cycle. We estimated a normalized signal by dividing each aggregated intensity by the median intensity across all trenches for each channel-timepoint combination. We proceeded to sum the normalized intensity over all channels and timepoints to produce a total normalized intensity. Empty cell traps were discarded by manual thresholding of the total normalized intensity, typically taking a value of at least ~30 for loaded cell traps. Finally, manual thresholds were set for positive and negative signals, for each cycle and channel, based on inspection of the bimodal intensity histogram. The thresholded signal was converted to a binary representation of the underlying RNA barcode and compared to the codebook generated from sequencing. For our CRISPRi experiments, we assigned barcode readouts to codebook barcodes if they were separated by no more than a Hamming distance of 1.

*Data Normalization and Filtering*

For our CRISPRi experiments, we accounted for measurement bias arising from differences in focus across single fields of view (FOVs) by linearly normalizing length (L), width (W) and mKate2Hyb intensity ($I_{rpsL}$), to their trench-wise baseline values, prior to dCas9 induction. These normalizations were subsequently propagated to the derived values of growth rate (λ), septum error ($L_s$), and inter-division time (τ). Examining the distributions of each parameter, with and without normalization, we observed that baseline normalization most strongly affects the distribution of widths (Extended Data Fig. 3b,c), likely because it is the parameter most strongly impacted in out-of-focus lineages.

After measuring our parameters, we filter our data to eliminate poorly quantified time-series phenotypes (growth rate, inter-division time, and septum error). First, we filtered out any time-series phenotypes measured from cell cycles that occur within a single timepoint (i.e., <20 min). Then, for each cell cycle, we fitted the growth rate across the cycle and eliminated measurements (inter-division time and septum error) from cell cycles with a poor growth fit (R2 < 0.9). Similarly, we filtered out instantaneous growth rate measurements with poor fits to the data (R2 < 0.9). We additionally removed instantaneous growth rate measurements with values less than 0 and greater than 4σ above the median growth rate, which is typically WT-like in these experiments. Finally, we filtered out measurements (inter-division time and septum error) from cell cycles with two timepoints (i.e., <30 min) that also exhibit an unrealistic added length of 4σ above the median (WT-like) added length.

After applying these filters based on cell growth quantification, we additionally filtered trenches exhibiting abnormal parameter values during the pre-induction period. To determine this, we fit a normal distribution to each measurable of interest and used this distribution to compute a sample-normalized likelihood for the pre-induction values from each trench. Following this, we remove data belonging to trenches with normalized likelihoods below the $10^{th}$ percentile for any of these quantities. For our data, we applied this filter to length, width, division length, mKate2Hyb intensity, and the instantaneous growth rate.

*Data Aggregation and Clustering*

To measure the post-induction parameter values of each sgRNA knockdown, we computed the parameter mean for each trench after 5 hours of dCas9 induction and then aggregated these values with a median across all trenches containing the same sgRNA (Extended Data Fig. 4a). Using a bootstrap reflecting this aggregation procedure, we computed the FDR of these values, using control sgRNAs to build the distribution under the null hypothesis. To account for differences in the number of trenches sampled for each variant, a bootstrapped null distribution was determined for the specific number of trenches observed in the case of the sgRNA being assessed (Supplementary Protocols).

To determine parameter time-series for each sgRNA, we first computed a time-series for each trench using a Gaussian kernel regression and proceeded to take the mean over all such time-series for the same sgRNA variant. The resultant time-series were optionally normalized by a Yeo-Johnson transform and subsequently transformed into Z-scores using a modified Z-score calculation where $\sigma$ is determined from the non-WT-like population of lineages (see supplementary protocol for details). Comparing these time-series for different sgRNAs, a soft-DTW (gamma = 1) distance matrix[12] was constructed and used to build a KNN graph (with 10 nearest neighbors). Leiden clustering at a resolution of 2 was used on this graph to determine phenotypic clusters, and a UMAP was generated (with PAGA initialization) for visualization purposes.[13]

To determine cluster stability, we used a jackknifing approach to resample our data with a dropout rate of 10%, generating 10 resampled cluster proposals. We then measured the similarity between each original cluster and its resampled derivatives, using the Jaccard index, taking the mean of this value to be a measure of cluster stability (Extended Data Fig. 7c). While some of these groupings are sensitive to sample noise and therefore spurious, many clusters were in fact stable to resampling, and we focused on these groups for our subsequent interpretation (compare highlighted clusters in Fig. 3a to Extended Data Fig. 7b).

*GFPmut2/DarkGFP Quantification and Analysis*

Quantification of GFPmut2 and mKate2Hyb signal from lDE15, to compute the GFP variant error rate (Extended Data Fig. 2), followed a similar procedure as in our CRISPRi measurement. We limited this quantification to measuring the mean intensity of the GFPmut2 and mKate2Hyb signals within cell masks. Unlike in the CRISPRi analysis, where normalization leveraged pre-induction phenotypes, systematic differences in intensity across different FOVs were corrected by measuring the modal values of positive GFPmut2 and mKate2Hyb signal within each field. Intensities from individual trenches were then linearly normalized based on these FOV-wise modal values. Following this, we performed background subtraction on the mean intensity in each cell mask, using the mean intensity of the area in each lineage bounding box without cell masks. Following this, we additionally filtered out trenches with abnormal mKate2Hyb signal intensity by a likelihood fitting procedure similar to that of the CRISPRi experiment, with a 2$^{nd}$ percentile cutoff. Finally, we computed the ratio of the GFPmut2 and mKate2Hyb signals for each cell mask and subsequently computed the median ratio for each trench, which was used to determine the overall GFP fluorescence call (Extended Data Fig. 2).

**Off-target Detection and sgRNA Curation**

For our 13 clusters of interest, we performed additional manual curation to account for possible off-target and polar effects that could result in misclassification of gene function. First, using the RegulonDB[14] annotations on the EcoCyc[15] web portal, we inspected each sgRNA for polar effects. We annotated sgRNAs as targeting either (1) monocistronic genes or genes at the promoter-distal end of polycistronic operons (monocistronic or "M"); (2) genes in polycistronic operons with no transcriptionally downstream genes belonging to the same cluster (non-polar or "NP"); (3) genes in polycistronic operons with downstream genes belonging to the same cluster (polar or "P"); or (4) genes in polycistronic operons with downstream genes suspected as exhibiting a related phenotype, based on previous literature (unclear or "UNC"). Then, if the sgRNA targets a PAM site which, in a previous bulk screen[6] showed more than a -2 log-fold fitness difference from all other PAM targets in the same gene, we annotated the sgRNA as a fitness outlier. Since this property is indicative of a potential off-target effect, we checked these sgRNAs for off-target sites by searching the chromosome for any 10 bp sequences adjacent to a valid PAM that perfectly match the PAM-adjacent sequence of the sgRNA. If any of the off-target sites belong to genes appearing in the same phenotypic cluster or are annotated in the literature with a similar phenotype, the sgRNA was annotated as potentially off-target. Finally, genes were assigned to clusters if at least 2 sgRNAs targeting those genes (1) did not lead to an apparent polar effect and (2) had no potential off-target effects.

**RNA/Protein Measurement**

First, 5 mL of culture was collected in a pre-chilled falcon tube, pelleted by centrifugation (4000 rcf) at 4°C, and resuspended in 1 mL of ice-cold 0.9% NaCl. This wash was then repeated once. Following this, 800 µL of the final resuspension was added to 200 µL of 0.2% SDS and 50 mM EDTA at 100°C to lyse the cells. The lysed sample then rested for 15 s at this temperature and was subsequently transferred to -20°C for storage.

To measure the protein concentration, 50 µL of these samples were processed and quantified using a Pierce BCA Protein Assay (Thermo Fisher Scientific #23227) according to the manufacturer's instructions. Briefly, each 50 µL sample was loaded onto a 96-well plate, with BCA kit standards, and mixed with 200 µL of BCA working reagent. After plates were incubated at 37°C for 30 min, absorbance from the sample was quantified using a plate reader (Synergy H1MF BioTek) with the monochromator set to 562 nm. Absorbance values were converted into protein concentration using a linear curve fit to triplicate measurements of the kit's protein standard curve.

To measure the RNA concentration in the sample, we followed a previously described sample processing protocol[16] and quantified RNA using a Quant-iT RNA Reagent and Assay Kit (Thermo Fisher Scientific Q33140) according to the manufacturer's instructions. Briefly, we diluted the sample lysate 1:20 in water and loaded 20 µL of each diluted sample on a 96-well plate, with Quant-iT RNA standards. We then mixed each sample with 200 µL of BR working solution and quantified sample fluorescence using a plate reader (Synergy H1MF BioTek) with monochromators set to 644 nm for excitation and 673 nm for emission. Fluorescence values were converted into RNA concentration using a linear curve fit to triplicate measurements of the kit's RNA standard curve. The RNA/Protein ratio was then calculated in terms of the measured abundance (by mass) of each.

**(p)ppGpp Measurement**

Prior to collection, we prepared 1.5 mL of lysis solvent for each sample, consisting of a 70:30 by volume ACN/H2O mixture, with 200 pmol/1.5 mL of a $^{13}C_{10}$,$^{15}N_5$-GTP isotope (CNLM-4269-CA-20, Cambridge Isotope Laboratories) as an internal standard. During collection, 10 mL of culture was run through a 0.45 μm PVDF filter (47 mm diameter) on a fritted glass funnel under vacuum over a period of ~10 s. Cells captured on the filter were immediately submerged in 1.5 mL of dry-ice chilled lysis solvent in a glass petri dish and moved to dry ice for a period of 10 minutes. After this, cells were gently dislodged by pipetting the solvent over the filter. Lysed cell solutions were then moved to microfuge tubes and stored at -80°C.

Our sample prep was derived from a previous approach to quantifying (p)ppGpp[17], with minor modifications. Samples were prepared for solid phase extraction (SPE) by retrieving them from the -80°C and thawing them on a heat block set to 40°C for 2 min. Samples were then immediately transferred, on ice, to a sonicator containing an ice-water slurry and sonicated for 10 min. Samples were then centrifuged at 20,000 rcf for 10 min at 4°C, to sediment cell debris. Then, samples were subject to SPE using Oasis WAX Cartridges (186002489, Waters). Each cartridge was pre-conditioned by adding 1 mL of methanol, followed by 1 mL of 50 mM ammonium acetate buffer (pH 4.5). Samples were then individually flowed through their corresponding cartridge, followed by a wash with 1 mL of the ammonium acetate buffer. Samples were then eluted into microfuge tubes by the addition of 200 μL of 2.8% ammonium hydroxide–MeOH/ACN/H2O (50:30:20 by volume) buffer. SPE-cleaned samples were stored at -80°C until further processing.

SPE-cleaned samples were finally prepared for HPLC-MS/MS analysis by retrieving them from the -80°C and thawing them on a heat block set to 40°C for 1 min, before putting them on ice. Samples were then centrifuged at 4000 rcf for 15 s. Following this, 10 μL of a 5% trehalose solution was added to each sample, which was subsequently vortexed. Each sample was then centrifuged at 4000 rcf for 15 s and transferred to a SpeedVac (Thermo Fisher) for drying at room temperature overnight. The next day, samples were resuspended in a 20 μL solution of ice-cold ACN/H2O (30:70 by volume). Samples were returned to -80°C until they were run on the HPLC-MS/MS. Just prior to analysis, 1 μL of medronic acid (200 mM in water) was added to each sample to reduce phosphate-group complexation to metal surfaces.

For quantification, we prepared triplicate nucleotide standard curves consisting of a 7-level, 1:5 dilution series, starting with a mixture of 500 μM GDP and GTP, and 50 μM ppGpp and pppGpp in water as well as triplicate blanks. 20 μL of each standard was added to 1.5 mL of lysis solvent and briefly vortexed before the addition of 10 µL of 5% trehalose. Standards were then dried and resuspended in the same manner as biological samples. Samples were analyzed on an Agilent 1290 liquid chromatography (LC) system (Agilent, Lexington, MA, USA) coupled with a 7500 triple quad mass spectrometer (MS) (AB Sciex, Framingham, MA, USA). 4 µL of sample was injected on a ZIC-pHILIC column (150x2.1 mm, Sigma-Aldrich, Saint Louis, MO, USA) kept at 40°C. Mobile phase A (MA) consisted of 20 mM ammonium carbonate in LCMS grade water with 0.1% ammonium hydroxide (all from Sigma-Aldrich, Saint Louis, MO, USA). Mobile phase B (MB) consisted of LC-MS grade acetonitrile (Sigma-Aldrich, Saint Louis, MO, USA). The gradient for the LC was as follows: starting at 80% B; to 0% B in 15 min; then 5 min isocratic at 0% B; then back to 80% B in 2 min; and a 10 min re-equilibration at 80% B. Data on the MS was acquired in negative mode with the following source parameters: source temperature 350°C, curtain gas at 42 psi, CAD gas at 8, source gas 1 at 25 psi and gas 2 at 60 psi, spray voltage at 3500 V. Both quadrupoles were operated in high resolution mode, with a 5 ms pause time. The following transitions were monitored: pppGpp quantifier: 681.97 to 583.89 (250 ms dwell time, EP -5 V, CE -33 V, CXP -36 V, Q0D -50 V). pppGpp qualifier: 681.97 to 503.99 (30 ms dwell time, EP -5 V, CE -37 V, CXP -14 V, Q0D -50 V). ppGpp quantifier: 601.99 to 503.99 (250 ms dwell time, EP -5 V, CE -35 V, CXP -26 V, Q0D -50 V). ppGpp qualifier: 601.99 to 423.95 (30 ms dwell time, EP -5 V, CE -47 V, CXP -26 V, Q0D -50 V). GTP quantifier: 522.02 to 158.9 (30 ms dwell time, EP -5 V, CE -39 V, CXP -14 V, Q0D -50 V). GTP qualifier: 522.02 to 424.09 (20 ms dwell time, EP -5 V, CE -31 V, CXP -12 V, Q0D -50 V). GDP quantifier: 442.07 to 158.9 (30 ms dwell time, EP -5 V, CE -33 V, CXP -20 V, Q0D 10 V). GDP qualifier: 442.07 to 344.06 (20 ms dwell time, EP -5 V, CE -39 V, CXP -22 V, Q0D 10 V). Internal standard quantifier: 537 to 457 (50 ms dwell time, EP -5 V, CE -35 V, CXP -26 V, Q0D -50 V). Quantification was performed on the quantifier areas divided by the IS quantifier area in Sciex OS (AB Sciex, Framingham, MA, USA). All LC-MS/MS steps downstream of sample prep were conducted by staff at the Harvard Center for Mass Spectrometry.

**Measurement of Length and Nucleoid Area in Agar Pads**

Prior to sample staining and imaging, we prepared a staining solution of 10 µg/mL DAPI in PBS. Then, fixed samples were retrieved from 4°C, pelleted by centrifugation at 4000 rcf for 5 min, and resuspended in 250 µL of the staining solution. Samples were incubated for 15 min at room temperature, pelleted by centrifugation at 4000 rcf for 5 min, and resuspended in 1 mL PBS.

At this point, samples were ready for imaging and kept under dark conditions. 22x22 mm 1.5% ultra-pure agarose (SeaKem LE Agarose) pads were prepared by heat dissolving the agarose in filtered PBS and then sandwiching the molten agarose solution between two 22x22 mm glass coverslips. To prepare each imaging sample, 1.5 µL of stained cells was pipetted onto an agar pad which was subsequently placed, sample-down, onto a 60x24 mm No. 1.5 coverslip. An evenly spaced grid of 36 FOVs was then imaged using largely the same microscopy set-up as described in "Mother Machine Phenotyping", with a 100x oil objective (CFI Plan Apo DM Lambda 100X Oil NA 1.45). For imaging the DAPI channel, our light path consisted of a multi-band dichroic (409/493/573/652/759 BrightLine, Semrock) and no emission filter.

**Replication Runout Assay**

Starter cultures for DE19 and DE502 were prepared as described in previous sections, in triplicate, and conducted at 30°C to avoid excessive growth defects in the *rne-1(ts)* strain DE502. On the day of the experiment, we measured OD600 for replicate starter cultures. Starter cultures for DE19 and DE502 were then diluted into 5 mL of EZRDM, pre-warmed in a 30°C shaking incubator, to a final OD600 of 0.0008 for DE19 and 0.0014 for DE502. We then serially diluted each of these cultures ten (DE19) or five (DE502) times at 1:2 into replicate 5 mL cultures. All cultures were then incubated at 30°C in a shaking incubator (220 rpm) for 4 h 10 min and subsequently transferred to a water bath with shaking (220 rpm) at 43°C (i.e., the restrictive temperature for *rne-1*). Just prior to transferring the cultures to 43°C, we measured the OD600 of each and collected the densest culture, for each replicate, with an OD between 0.125 and 0.25 for replication runout at 30°C. The cultures moved to the 43°C water bath continued growing for an additional 2 hours, with samples collected for runout at 43°C, at the 30-minute, 1-hour, and 2-hour post-shift timepoints.

Runouts consisted of inoculating 1 mL of selected cultures into 4mL of EZRDM supplemented with rifampicin (USBiological #13292-46-1) and cephalexin (Sigma C4895) such that the final concentrations at 5 mL were 300 µg/mL and 30 µg/mL, respectively. Runout cultures were kept in a shaking incubator at 30°C or 43°C and cultured for an additional 3 hours, before being fixed by pipetting 1 mL of culture into 9mL of filtered 70% EtOH in conical tubes, which were then stored at 4°C. The timings and concentrations used for the runout protocol were derived from previous work.[18] To generate reference populations of cells possessing single chromosomes, we performed a triplicate starter and pre-culture protocol like that described in "Strains and Growth

Conditions" for DE19 grown in MBM + 0.2% Succinate medium at 37°C. We then allowed pre-cultures to grow to early stationary phase before collecting samples by the same EtOH fixation protocol used for runouts, ensuring that all cells in the population bore a single, fully replicated chromosome.

Fixed samples were stained by first pelleting the sample (4000 rcf for 10 min), aspirating out the supernatant down to ~50 µL, and allowing the sample to dry at 37°C for 30 min. Following this, the pellet was resuspended in 1 mL staining solution (4X Pico488 in filtered TBS). Samples were then stained at room temperature for 30 min, pelleted (4000 rcf for 10 min), and the supernatant subsequently aspirated. Stained pellets were then carefully resuspended in 1 mL of filtered PBS by pipetting. Finally, the 1 mL sample was run through a 35 µm Cell Strainer Cap into a Round-Bottom Polystyrene 12 x 75 mm Tube (VWR, #21008-948).

To measure Pico488 fluorescence in single cells, we used a BD LSR II Flow Cytometer with a 488nm laser. We also measured forward and side scatter; however these data were not used to gate cells. For each sample, we collected at least 100,000 events, with a detection rate between 200 and 1000 events per second, which was achieved by diluting samples in additional PBS when necessary. All flow cytometry was performed at the Immunology Flow Cytometry Core Facility at Harvard Medical School.

**Time-course CFU Measurements**

Replicate starter cultures for CFU measurements were treated with the same protocol as in the "Replication Runout Assay" and then diluted into 25 mL of EZRDM, pre-warmed in a 30°C shaking incubator, to a final OD600 of 0.0008 for DE19 and 0.0014 for DE502. All cultures were then incubated at 30°C in a shaking incubator (220 rpm) for 2 h 50 min. Following this, cultures were grown under these conditions for an additional 80 minutes, with samples taken every 20 minutes, serially diluted by a factor of 1:20,000, and plated by bead spreading of 100 µL of the dilute sample onto LB Lennox plates. After these 80 minutes, cultures were transferred to a water bath with shaking (220 rpm) at 43°C (i.e., the restrictive temperature for *rne-1*). Subsequently, cultures were grown for an additional 1 or 2 hours, for $rne^{WT}$ and *rne-1(ts)* cultures respectively, with samples for plating taken every 20 minutes. Following sample collection, plates were incubated at 30°C overnight and CFUs were assessed by counting visible colonies.

**NGS Profiling of Chromosome Ploidy**

Replicate starter cultures for CFU measurements were treated with the same protocol as in the "Replication Runout Assay" and then diluted into 5 mL of EZRDM, pre-warmed in a 30°C shaking incubator, to a final OD600 of 0.04 for DE19 and 0.06 for DE502. We then serially diluted each of these cultures ten (DE19) or five (DE502) times at 1:2 into replicate 5 mL cultures. All cultures were then incubated at 30°C in a shaking incubator (220 rpm) for 1 h 30 min and subsequently transferred to a water bath with shaking (220 rpm) at 43°C (i.e., the restrictive temperature for *rne-1*). Just prior to transferring the cultures to 43°C, we measured the OD600 of each and selected the densest culture, for each replicate, with an OD between 0.125 and 0.25. 1.9 mL of this culture was flash frozen in liquid nitrogen for NGS-based marker frequency analysis. The cultures moved to the 43°C water bath continued growing for an additional 2 h, after which point the same sample collection was performed on the post-treatment cultures. Triplicate stationary phase MBM + 0.2% Succinate cultures were also prepared, similarly to the "Replication Runout Assay", to provide single chromosome reference samples.

Flash-frozen samples were later thawed at 4°C and subsequently pelleted at 4°C (4000 rcf for 5 min). gDNA was subsequently extracted from cells using NEBExpress® T4 Lysozyme (NEB #P8115) and the Monarch® Genomic DNA Purification Kit (NEB #T3010), according to the manufacturer's protocol for Gram-negative bacteria, and eluted in nuclease-free water. gDNA concentration was subsequently measured by a Qubit High Sensitivity dsDNA Quantification Assay (Thermo Fisher #Q32851). These samples were subsequently prepared for multiplexed sequencing by tagmentation using a Nextera XT Kit (Illumina). Following library prep, samples were subjected to quality control by TapeStation (Agilent 2200 TapeStation D1000 HS ScreenTape). Finally, libraries were pooled with equal stoichiometry and sequenced on a NextSeq 500 using a high output, 75-cycle kit (Illumina # 20024906). All library prep and sequencing in this section was performed by staff at the Biopolymers Facility at Harvard Medical School.

Raw reads were mapped to a reference genome (Accession: U00096.3) using Minimap2[19] and then filtered for a Mapping Quality Score of at least 22. The reference genome was split into 100kb bins, with the 0-coordinate set to *oriC*. Read counts within each of these bins were quantified based on the mapping positions of the filtered reads. Counts were converted into frequencies by normalizing to the total number of reads per sample, and, subsequently, the mean and standard error of read frequencies for each bin were computed from triplicate biological replicates. These values were then normalized to the single-chromosome succinate reference, to account for mapping biases, with the SEM of the final ratio determined by error propagation.

**Cloning Strains**

*pRpsL Fluorescent Reporters*

To perform library-scale screens on *E. coli*, we engineered various background strains with combinations of fluorescence reporters, a dCas9 expression system, and a T7 RNA polymerase expression system. To integrate a *pRpsL-mKate2Hyb* fluorescent reporter, which consists of a translational fusion replacing the N-terminus of mKate2 (i.e., MVSE) with the N-terminus of mCherry (i.e., MVSKGEENNMA),[20] a dsDNA gBlock (gbMI4) was inserted into the cassette of pNDL-1 using BP Clonase, and subsequently transformed into CGSC6300 and grown up at 37°C to induce integration into the *attTN7* site, resulting in MI13/JP1458. MI13 was then transformed with pE-FLP to cure Kan-R and subsequently cultivated at 43°C to remove pE-FLP, resulting in DE32. The same strategy was used to integrate a *pRpsL-mVenus* fluorescent reporter from another dsDNA gBlock (gbMI3) into CGSC6300 at the *attTN7* site, resulting in MI12/JP1457.

*T7 RNA Polymerase Expression System*

Construction of an appropriate T7 RNA polymerase expression system went through several iterations. We first integrated an Arabinose-inducible T7 expression system by isothermal assembly of a pBAD-T7RNAPol fragment amplified from pTARA (oDE71/oDE72) and the pOSIP-KT (kan) backbone after BamHI, XhoI restriction digestion. Transformation, subsequent integration, and curing of kan resistance by pE-FLP resulted in DE43. Switching to a Cuminic acid-inducible system, we isothermally assembled a T7RNAPol fragment amplified from DE43 (oDE284/ oDE285) and a fragment amplified from pAJM.657 (oDE282, oDE283) to yield the plasmid pDE57. The expression cassette of pDE57 was amplified (oDE286, oDE294) and assembled with the pOSIP-KT (kan) backbone after XhoI, KpnI restriction digestion, yielding the integration plasmid pDE91. The rbs in pDE91 was weakened by swapping it for the BCD24 rbs by overhang PCR of pDE91 (oDE397, oDE398), blunt end ligation, and transformation into PIR2 cells, making pDE112. We subsequently used pDE112 to integrate the T7RNAPol expression system into various strains.

*dCas9 Expression System*

We integrated an aTc-inducible dCas9 expression system by first constructing an isothermal assembly of a PaTc fragment (oDE137, oDE138) and a dCas9 fragment (oDE139, oDE140) amplified from pdCas9-bacteria, into a pOSIP-KT (kan) backbone after PstI, EcoRI restriction digestion. The assembly was then transformed into CGSC6300 and cultured at 37°C to induce integration into the *att186(O)* site. This intermediate strain was then transformed with pE-FLP to cure Kan-R as in DE32, resulting in DE65. Note that the weaker rbs (LC-E75) from Cui et al 2018[8] was included in the overhang of oDE139 to override the pdCas9-bacteria rbs in the final DE65 strain. This step was taken to reduce the severity of "bad seed" off-target effects.

*CRISPRi and Barcoding Background Strains*

We used P1 transduction to transfer *pRpsL-mKate2Hyb* from MI13/JP1458 into DE65, subsequently curing Kan-R with pE-FLP, resulting in DE122. Following this, we generated MARLIN-compatible backgrounds by transforming pDE112 (containing the Cuminic acid-inducible T7RNAPol system) into several base strains, integrating the pDE112 cassette by growth at 37°C, and subsequently curing Kan-R with pE-FLP. This procedure was applied to DE32, DE65, and DE122, resulting in DE344, DE346, and DE348. We additionally transferred the *hupA::hupA-mCherry* construct present in SS6279 into DE346 by P1 transduction, followed by curing Kan-R by pE-FLP, resulting in DE566. Subsequently, we introduced a *pRpsL-mVenus* reporter into this strain by P1 transduction from MI12/JP1457 and Kan-R curing by pE-FLP, producing LAG436.

In addition to the main barcoding strains, we also generated the background DE120 by P1 transducing *pRpsL-mKate2Hyb* from MI13/JP1458 into DE43, followed by curing Kan-R with pE-FLP. While this strain was not used during a full MARLIN experiment, it was used as the genetic background for the library lDE11 (see "GFPmut2/DarkGFP Library Cloning"), which was used to benchmark nanopore sequencing performance. This procedure was also used to P1 transduce *pRpsL-mKate2Hyb* into DE806, DE814, DE816, and DE817, to produce DE828, DE830, DE831, and DE832, respectively.

*ppGpp Deletion Series Cloning*

We introduced deletions of *relA* and *spoT* into several strains to modulate (p)ppGpp control. To delete *relA*, we transferred the *ΔrelA782::frt-kan-frt* allele from CGSC10159 into several strains by P1 transduction and subsequently cured Kan-R by pE-FLP. We performed this procedure on DE65, DE122, LAG436, and CGSC6300, resulting in the strains DE419, DE420, DE683 and DE792. In order to delete *spoT*, we amplified the *ΔspoT696::frt-kan-frt* allele from TU244 (oDE659, oDE660) and integrated it into the native locus in DE420 and DE683 by Lambda Red recombineering, resulting in DE730 and DE735. Briefly, pSIM-Tet was transformed into the target strain to introduce the Lambda Red genes, cells were recovered, and single colonies were isolated at 30°C. Cultures were then inoculated with the selected colonies, cultivated to OD600 ~0.4, and recombineering functions were induced by incubation at 42°C for 15 min. Cells were then made electrocompetent, transformed with the *ΔspoT696::frt-kan-frt* fragment, and plated on selective (kan) medium at 37°C. Colonies were subsequently screened for Tet sensitivity to confirm loss of pSIM-Tet as well as sequenced at the *spoT* locus before being archived.

*Cloning MARLIN Plasmids*

All MARLIN libraries were cloned using a three-part combinatorial ligation composed of a barcode expression cassette, a variant (sgRNA or GFP) cassette, and a p15a plasmid backbone. Here we describe the cloning of the various plasmids generated for this purpose. To make pDE29, which constitutively expresses a GFPmut2 protein, we first used isothermal assembly to join a GFPmut2 fragment amplified (oDE197, oDE94) from pEB1-mGFPmut2, a backbone amplified (oDE195, oDE196) from LPT41, and two ssDNA oligos (oDE199, oDE205) containing a J23116 promoter and a T1 terminator. The assembly was transformed, and the resulting plasmid was miniprepped. This intermediate plasmid was subsequently amplified (oDE207, oDE208) and blunt-end ligated to remove a BsaI site from the mGFPmut2 CDS. The ligation was then transformed into DH5α cells, yielding DE93/pDE29. A subsequent round of amplification (oDE209, oDE210) of pDE29 (introducing a Y66L mutation into GFPmut2), blunt-end ligation, and transformation into DH5α resulted in DE98/pDE32.

We produced two versions of a sgRNA expression plasmid compatible with our barcode ligation procedure, pDE93 and pDE170. pDE93 was produced in two steps. First, we performed an isothermal assembly of a fragment amplified (oDE276, oDE277) from pDE29 and a ssDNA oligo oDE278, transforming the assembly into DH5α cells to yield the intermediate plasmid pDE56. Following this, a second isothermal assembly was performed on a fragment amplified (oDE338, oDE339) from pDE56 and a fragment amplified (oDE342, oDE336) from V37m. Transformation of this assembly into DH5α resulted in DE237/pDE93. In the course of this work, we found that the ligation of the small 28 bp library dsDNA into pDE93 was inefficient, making CRISPRi library cloning difficult. pDE170 was designed as an alternate version of pDE93 where part of the sgRNA promoter is omitted and instead included on the array-synthesized library ssDNA, making the final insert substantially larger and more efficient to ligate. To make pDE170, we performed an isothermal assembly of a fragment amplified (oDE553, oDE554) from pDE93 and a fragment amplified (oDE342, oDE336) from V37m. This assembly was transformed into DH5α cells, producing DE562/pDE170.

The barcode expression plasmid (pDE47), into which barcodes are assembled as described in the main methods, was made in two steps. First, we performed an isothermal assembly of a fragment amplified (oDE195, oDE196) from LPT41 and the ssDNA oligo oDE204, transforming this assembly into DH5α cells, resulting in the intermediate plasmid DE94/pDE30. Then, we performed a subsequent assembly of a fragment amplified (oDE252, oDE196) from pDE30 and the ssDNA oligo oDE254, transforming the assembly into DH5α cells to yield DE134/pDE47. To assemble combinatorial FISH barcodes into this expression cassette, we used the approach described in the main methods. In this work, we used two different oligo pools to assemble barcodes, oDEPool2 (lDE15, 20, 26, 28, and 30) and oDEPool3 (lDE11). These two barcode designs are the same in most respects; however, oDEPool3 contains more constant "spacer" binding sites interspersed within the barcode sequence, which can help facilitate short-read sequencing, though we did not use this feature.

Finally, we made a vector (pDE36) with a digestible backbone providing the resistance marker (chlor) and a p15a origin of replication. To do this, we performed an isothermal assembly of a fragment amplified (oDE225, oDE226) from pDHL981 and a fragment amplified (oDE227, oDE228) from pDHL138. The assembly was then transformed into NEB Turbo cells, resulting in DE109/pDE36. We note that the BsaI restriction sites in each of our variant, barcode, and backbone vectors are also compatible with golden gate assembly, though we used standard restriction-ligation throughout this work.

*CRISPRi Library Cloning*

The CRISPRi library cloning procedure is described at length in the main methods section and the supplementary protocol. For the MARLIN library lDE20, we ligated a BbsI-HF digested backbone fragment of pDE93 with a BbsI-HF and BsmBI-v2 digested amplicon (oDE418/oDE433) of the array-synthesized sgRNA pool oDEPool7. After ligating the resulting library cassette to barcodes, according to the barcoding procedure in the main methods section, we transformed the resulting library into DE348 to produce plDE20 in lDE20. We followed a similar procedure for lDE26, where instead we ligated a BbsI-HF digested amplicon (oDE418/oDE550) of our sgRNA library oDEPool13 into a BbsI-HF digested backbone fragment of the alternative sgRNA expression plasmid, pDE170. This library was then barcoded, as before, and transformed into LAG436 to produce plDE26 in lDE26. lDE28 and lDE30 were produced by transforming plDE26 into DE683 and DE735, respectively.

*GFPmut2/DarkGFP Library Cloning*

The GFPmut2/DarkGFP library cloning procedure proceeded similarly to that of the CRISPRi libraries, with the fluorescent protein expression plasmid (pDE29 or pDE32) taking the place of the sgRNA expression plasmid. Barcode ligation and transformation into the DE344 background strain proceeded independently for the GFPmut2 (pDE29) plasmid and the DarkGFP plasmid (pDE32). After transformation efficiencies were estimated, these two parts of the library were pooled to make lDE15 such that the same number of each GFP variant was similar (Extended Data Fig. 1d).

The lower complexity GFP library lDE11, which was used to characterize the nanopore error in GFP variant calling, was generated using the same strategy. However, while all other libraries in this work were labeled with FISH barcodes assembled from the oDEPool2 oligo pool, this library was labeled with barcodes derived from the oDEPool3 oligo pool (see "Cloning MARLIN Plasmids"). In addition, this plasmid library was transformed into a DE120 background, containing a legacy Arabinose-inducible expression system which was not used in this work.

*Recombineering DE502*

To introduce the *rne-1(ts)* allele, consisting of a G66S missense mutation, into CGSC6300, we used a version of ssDNA recombineering.[21] The recombineering protocol requires the plasmid pAV203 (courtesy of Antoine Vigouroux), a derivative of the pORTMAGE-Ec1 vector,[21] which expresses CspRecT, a single-stranded-DNA-annealing protein which improves recombineering efficiency, as well as MutL(E32K), a dominant-negative protein which interferes with mismatch repair. In pAV203, both CspRecT and MutL(E32K) are under m-toluate-inducible control, while the vector itself possesses a pSC101ts origin, enabling heat-curing. pAV203 was made by an isothermal assembly of a fragment amplified (oAV_V1/ oAV_V2) from the pORTMAGE-Ec1 plasmid and a fragment amplified (oAV_V3/ oAV_V4) from pE-FLP and transformation into VH1000 cells.

Briefly, recombineering consists of first transforming the target strain with pAV203, culturing the resultant isolate under m-toluate induction for 1 hour, and making the induced culture electrocompetent. Competent cells should then be transformed with a ssDNA oligo containing a sequence complementary to the lagging strand at the target site, with the intended substitution, and usually ~90 bp in length with the substitution in the center. It is recommended to include phosphorothioate modifications in between the first two bps at the 5' end of the oligo. We performed ssDNA recombineering with oDE516 to introduce the *rne-1* allele into CGSC6300 and subsequently cured pAV203 by growth at 37°C, yielding DE502.

*Inducible FusA*

To put EF-G/FusA expression under AHL-inducible control, we first constructed an isothermal assembly of a PLux fragment amplified (oDE382, oDE383) from pAJM.713, a LuxR fragment amplified (oDE386, oDE387) from sAJM.1506, and the pOSIP-KT (kan) backbone after XhoI, KpnI restriction digestion. The assembly was transformed into PIR2 to yield pDE104. We then replaced the YFP CDS in pDE104 with the native FusA CDS by isothermal assembly of a fragment amplified (oDE282, oDE583) from pDE104 and a fragment amplified (oDE736, oDE737) from CGSC6300, which was transformed into PIR2 to yield pDE237. Following this, we transformed pDE237 (containing AHL-inducible FusA) into several strains, integrating the pDE237 cassette by growth at 37°C, and subsequently curing Kan-R with pE-FLP. We performed this procedure on CGSC6300 and DE792, resulting in the strains DE794 and DE796. To then delete the native copy of *fusA* in each of these strains, we performed Lambda red recombineering with pSIM-Tet (see "ppGpp Deletion Series Cloning") using a fragment amplified (oDE744, oDE745) from CGSC10818, containing the KEIO Kan cassette with added overhangs to integrate at the native *fusA* locus. The Kan marker was then cured by pE-FLP, resulting in strains DE806 and DE808.

*sgRNA Isolate Cloning*

As part of validating and following up on our screen, we cloned several strains expressing particular sgRNAs in isolation. For each of these strains, sgRNA targeting sequences were cloned into psgRNAc by annealing paired oligos designed with the 20 bp targeting sequence on the F oligo and its reverse complement on the R oligo. Each oligo was also designed with overhang sequences to facilitate ligation, with the sequence "TAGT" on the 5' end of the F oligo and the sequence "AAAC" on the 5' end of the R oligo. After annealing these oligo pairs, the resulting sticky-ended dsDNA was ligated into the psgRNAc backbone generated by BsaI-HF2 restriction digestion. The ligated plasmid was finally transformed into the relevant target strain, selected for on chlor, and the plasmid cassette was confirmed by Sanger sequencing.

pDE223/DE751 resulted from a ligation of oligos oDE690, oDE699 and transformation into DE65. pDE224/DE752 resulted from a ligation of oligos oDE691, oDE700 and transformation into DE65. pDE235/DE772 resulted from a ligation of oligos oDE716, oDE717 and transformation into DE65. pDE236/DE428 resulted from a ligation of oligos oDE754, oDE760 and transformation into DE122. Subsequently, pDE236 was miniprepped and transformed into DE65, DE65(co-transformed with pDE214), and DE419, resulting in DE787, DE773, and DE825. pDE239/DE814 and pDE239/DE818 resulted from a ligation of oligos oDE718, oDE722 and transformation into DE65 and DE419. pDE240/DE815 and pDE240/DE819 resulted from a ligation of oligos oDE719, oDE723 and transformation into DE65 and DE419. pDE241/DE816 and pDE241/DE820 resulted from a ligation of oligos oDE720, oDE724 and transformation into DE65 and DE419. pDE242/DE817 and pDE242/DE821 resulted from a ligation of oligos oDE721, oDE725 and transformation into DE65 and DE419. pDE244/DE423 resulted from a ligation of oligos oDE749, oDE755 and transformation into DE122. pDE246/DE425 resulted from a ligation of oligos oDE751, oDE757 and transformation into DE122. pDE247/DE426 resulted from a ligation of oligos oDE752, oDE758 and transformation into DE122. pDE248/DE427 resulted from a ligation of oligos oDE753, oDE759 and transformation into DE122. pDE260/DE847 resulted from a ligation of oligos oDE775, oDE784 and transformation into DE65. pDE261/DE848 resulted from a ligation of oligos oDE776, oDE785 and transformation into DE65. pDE262/DE849 resulted from a ligation of oligos oDE777, oDE786 and transformation into DE65. pDE263/DE850 resulted from a ligation of oligos oDE778, oDE787 and transformation into DE65. pDE264/DE851 resulted from a ligation of oligos oDE779, oDE788 and transformation into DE65. pDE265/DE852 resulted from a ligation of oligos oDE780, oDE789 and transformation into DE65. pDE266/DE853 resulted from a ligation of oligos oDE781, oDE790 and transformation into DE65. pDE267/DE854 resulted from a ligation of oligos oDE782, oDE791 and transformation into DE65. pDE268/DE855 resulted from a ligation of oligos oDE783, oDE792 and transformation into DE65.

*Cloning RelA* Expression Plasmid*

In order to express the hyperactive *relA* allele, *relA**, we placed RelA* expression under the control of IPTG, in a plasmid (pDE214) compatible with psgRNAc in a two-plasmid system. We generated this plasmid by performing an isothermal assembly of a fragment amplified (oDE672, oDE673) from pRelA' and a fragment amplified (oDE282, oDE583) from pDE211. This assembly was transformed into DH5α cells, yielding DE737/pDE214.

### Supplementary Tables

**Supplementary Table 1: DNA probes used in this study.**

| Probe Name | Dye | Barcode Site | Sequence | FISH Cycle |
|---|---|---|---|---|
| DE-A1 | AF555 | 1 | ACACTACCACCATTTCCTAT | 1 |
| DE-A2 | AF555 | 2 | AAACACACACTAAACCACCC | 2 |
| DE-A3 | Cy5 | 3 | ATCCTCCTTCAATACATCCC | 1 |
| DE-A4 | Cy5 | 4 | TATCTCATCAATCCCACACT | 2 |
| DE-A5 | Alexa750 | 5 | ACTCCACTACTACTCACTCT | 1 |
| DE-A6 | Alexa750 | 6 | AACTCATCTCAATCCTCCCA | 2 |
| DE-A7 | AF555 | 7 | ACCACAACCCATTCCTTTCA | 3 |
| DE-A8 | AF555 | 8 | TCTATCATCTCCAAACCACA | 4 |
| DE-A9 | Cy5 | 9 | ACCCTCTAACTTCCATCACA | 3 |
| DE-A10 | Cy5 | 10 | AATACTCTCCCACCTCAACT | 4 |
| DE-A11 | Alexa750 | 11 | TTTCTACCACTAATCAACCC | 3 |
| DE-A12 | Alexa750 | 12 | TCCAACTCATCTCTAATCTC | 4 |
| DE-A13 | AF555 | 13 | TCCTATTCTCAACCTAACCT | 5 |
| DE-A14 | AF555 | 14 | ATAAATCATTCCCACTACCC | 6 |
| DE-A15 | Cy5 | 15 | ACCCTTTACAAACACACCCT | 5 |
| DE-A16 | Cy5 | 16 | TTCCTAACAAATCACATCCC | 6 |
| DE-A17 | Alexa750 | 17 | TATCCTTCAATCCCTCCACA | 5 |
| DE-A18 | Alexa750 | 18 | ACCCAACACTCATAACATCC | 6 |
| DE-A19 | AF555 | 19 | TTTACTCCCTACACCTCCAA | 7 |
| DE-A20 | AF555 | 20 | ACTTTCCACATACTATCCCA | 8 |
| DE-A21 | Cy5 | 21 | ACATTACACCTCATTCTCCC | 7 |
| DE-A22 | Cy5 | 22 | TACTACAAACCCATAATCCC | 8 |
| DE-A23 | Alexa750 | 23 | TTCTCCCTCTATCAACTCTA | 7 |
| DE-A24 | Alexa750 | 24 | TTCTTCCCTCAATCTTCATC | 8 |
| DE-A25 | AF555 | 25 | TCCTAACAACCAACTACTCC | 9 |
| DE-A26 | AF555 | 26 | ACCTTTCTCCATACCCAACT | 10 |
| DE-A27 | Cy5 | 27 | ACCCTTACTACTACATCATC | 9 |
| DE-A28 | Cy5 | 28 | AATCTCACCTTCCACTTCAC | 10 |
| DE-A29 | Alexa750 | 29 | TCTATCATTACCCTCCTCCT | 9 |
| DE-A30 | Alexa750 | 30 | TCCTCATCTTACTCCCTCTA | 10 |
| DE-A61 | AF647 | 3 | ATCCTCCTTCAATACATCCC | 1 |
| DE-A62 | AF647 | 4 | TATCTCATCAATCCCACACT | 2 |
| DE-A63 | AF647 | 9 | ACCCTCTAACTTCCATCACA | 3 |
| DE-A64 | AF647 | 10 | AATACTCTCCCACCTCAACT | 4 |
| DE-A65 | AF647 | 15 | ACCCTTTACAAACACACCCT | 5 |
| DE-A66 | AF647 | 16 | TTCCTAACAAATCACATCCC | 6 |
| DE-A67 | AF647 | 21 | ACATTACACCTCATTCTCCC | 7 |
| DE-A68 | AF647 | 22 | TACTACAAACCCATAATCCC | 8 |
| DE-A69 | AF647 | 27 | ACCCTTACTACTACATCATC | 9 |
| DE-A70 | AF647 | 28 | AATCTCACCTTCCACTTCAC | 10 |

| | | | | |
|---|---|---|---|---|
| DE-A71 | AF488 | 1 | ACACTACCACCATTTCCTAT | 1 |
| DE-A72 | AF488 | 2 | AAACACACACTAAACCACCC | 2 |
| DE-A73 | AF488 | 7 | ACCACAACCCATTCCTTTCA | 3 |
| DE-A74 | AF488 | 8 | TCTATCATCTCCAAACCACA | 4 |
| DE-A75 | AF488 | 13 | TCCTATTCTCAACCTAACCT | 5 |
| DE-A76 | AF488 | 14 | ATAAATCATTCCCACTACCC | 6 |
| DE-A77 | AF488 | 19 | TTTACTCCCTACACCTCCAA | 7 |
| DE-A78 | AF488 | 20 | ACTTTCCACATACTATCCCA | 8 |
| DE-A79 | AF488 | 25 | TCCTAACAACCAACTACTCC | 9 |
| DE-A80 | AF488 | 26 | ACCTTTCTCCATACCCAACT | 10 |

**Supplementary Table 2: Genetic libraries used in this study.**

| Library | Parent Strain | *Genotype* | Figure |
|---|---|---|---|
| IDE11 | DE120 | *attP21::pAra-T7RNAPol*<br>*attTN7::pRpsL-mKate2Hyb*<br>*pIDE11* | S1a-b |
| IDE15 | DE344 | *attTN7::pRpsL-mKate2Hyb*<br>*attP21::pCymRC-BCD24-T7RNAPol*<br>*pIDE15* | S1c-d; S2c,d,f |
| IDE20 | DE348 | *att186 (primary)::paTc-rbs*-dCas9*<br>*attTN7::pRpsL-mKate2Hyb*<br>*attP21::pCymRC-BCD24-T7RNAPol*<br>*pIDE20* | 1b; 2a-e; 3a-c; 4a,c; 5a-d; 7c-e; S1e-i; S3d,e; S4; S5; S7; S8a-e; S9; S11a-c; S13b,c |
| IDE26 | LAG436 | *att186 (primary)::paTc-rbs*-dCas9*<br>*attP21::pCymRC-BCD24-T7RNAPol*<br>*hupA::hupA-mCherry*<br>*attTN7::pRpsL-mVenus*<br>*pIDE26* | 2f; 3d; 4a-d; 6c; S1j; S6a-c,f; S8a,e,f; S10a; S13a |
| IDE28 | DE683 | *att186 (primary)::paTc-rbs*-dCas9*<br>*attP21::pCymRC-BCD24-T7RNAPol*<br>*hupA::hupA-mCherry*<br>*attTN7::pRpsL-mVenus*<br>*ΔrelA782*<br>*pIDE26* | 6c; S6d |
| IDE30 | DE735 | *att186 (primary)::paTc-rbs*-dCas9*<br>*attP21::pCymRC-BCD24-T7RNAPol*<br>*hupA::hupA-mCherry*<br>*attTN7::pRpsL-mVenus*<br>*ΔrelA782*<br>*ΔspoT(2-696)::frt-kan-frt*<br>*pIDE26* | 6c; S6e |

**Supplementary Table 3: Strains used in this study.**

| Strain | Background | *Genotype* | Source | Figure |
|---|---|---|---|---|
| CGSC6300 | *E. coli* MG1655 | *wild-type* | [22] | 4f-h; S9b-d |
| CGSC10818 | *E. coli* K12 BW25113 | *ΔhslV720::kan* | [23] | N/A |
| CGSC10159 | *E. coli* K12 BW25113 | *ΔrelA782::kan* | [23] | N/A |
| AddGene_121043 | *E. coli* JM109 | *V37m (kan)* | V37m was a gift from Richard Murray, Part of CIDAR MoClo Extension | N/A |
| AddGene_114006 | *E. coli* DH5α | *psgRNAc (chlor)* | [8] | N/A |
| AddGene_108528 | *E. coli NEB Stable* | *pAJM.336 (kan)* | [24] | N/A |
| AddGene_108525 | *E. coli NEB Stable* | *pAJM.657 (kan)* | [24] | N/A |
| AddGene_108514 | *E. coli NEB Stable* | *pAJM.713 (kan)* | [24] | N/A |
| AddGene_44249 | *E. coli Top10* | *pdCas9-bacteria (chlor)* | [25] | N/A |
| AddGene_31491 | *E. coli GC5* | *pTARA (chlor)* | [26] | N/A |
| AddGene_45985 | *E. coli DB3.1* | *pOSIP-KO (kan)* | [27] | N/A |
| AddGene_45987 | *E. coli DB3.1* | *pOSIP-KT (kan)* | [27] | N/A |
| AddGene_45978 | *E. coli E811 (P2 lysogen)* | *pE-FLP (amp)* | [27] | N/A |
| AddGene_175595 | *E. coli* DH5α | *pRelA' (kan)* | [28] | N/A |
| AddGene_103980 | *E. coli* MG1655 | *pEB1-mGFPmut2 (kan)* | [29] | N/A |
| LPT41 | *E. coli MC1061* | *pLPT41 (kan)* | This Study | N/A |
| DHL138 | *E. coli* DH5α | *pDHL138 (kan)* | [20] | N/A |
| DHL981 | *E. coli* Turbo | *pDHL981 (chlor)* | This Study | N/A |

| | | | | |
|---|---|---|---|---|
| pSIM5-Tet | *E. coli W3110* | *pSIM5-Tet (tet)* | [30] | N/A |
| pNDL-1 | *E. coli* DH5α | *pNDL-1 (amp)* | [20] | N/A |
| pAV203 | *E. coli* VH1000 | *pAV203 (amp)* | This Study | N/A |
| AddGeneStrain_108254/sAJM.1506 | *E. coli* MG1655 | *position_3,860,010::Marionette_Cluster(see ref)* | [24] | N/A |
| SS6279 | *E. coli JC13509* | *hupA::hupA-mCherry-frt-kan-frt* | [31] | N/A |
| TU244 | *E. coli* MG1655 | *rph1 ilvG rfb-50 lacIZYA(del)-FRT relA(del)-FRT attLambda::cro-bio(del) nad::Tn10 spoT::kan* | TU244 was a gift from Tom Bernhardt | N/A |
| MI12/JP1457 | *E. coli* MG1655 | *attTN7::pRpsL-mVenus-frt-kan-frt* | This Study | N/A |
| MI13/JP1458 | *E. coli* MG1655 | *attTN7::pRpsL-mKate2Hyb-frt-kan-frt* | This Study | N/A |
| DE32 | *E. coli* MG1655 | *attTN7::pRpsL-mKate2Hyb* | This Study | S3a-c |
| DE43 | *E. coli* MG1655 | *attP21::pAra-T7RNAPol:N823D* | This Study | N/A |
| DE65 | *E. coli* MG1655 | *att186 (primary)::paTc-rbs*-dCas9* | This Study | N/A |
| DE93 | *E. coli* DH5α | *pDE29 (kan)* | This Study | N/A |
| DE94 | *E. coli* DH5α | *pDE30 (kan)* | This Study | N/A |
| DE98 | *E. coli* DH5α | *pDE32 (kan)* | This Study | N/A |
| DE109 | *E. coli* NEB Turbo | *pDE36 (chlor)* | This Study | N/A |
| DE120 | *E. coli* MG1655 | *attP21::pAra-T7RNAPol attTN7::pRpsL-mKate2Hyb* | This Study | N/A |
| DE122 | *E. coli* MG1655 | *att186 (primary)::paTc-rbs*-dCas9 attTN7::pRpsL-mKate2Hyb* | This Study | N/A |
| DE134 | *E. coli* DH5α | *pDE47 (kan)* | This Study | N/A |
| DE138 | *E. coli* DH5α | *pDE56 (kan)* | This Study | N/A |
| DE150 | *E. coli* DH5α | *pDE57 (kan)* | This Study | N/A |
| DE223 | *E. coli* One-shot PIR1 | *pDE91 (kan)* | This Study | N/A |

| | | | | |
|---|---|---|---|---|
| DE237 | *E. coli* DH5α | *pDE93 (kan)* | This Study | N/A |
| DE307 | *E. coli* One-shot PIR2 | *pDE104 (kan)* | This Study | N/A |
| DE326 | *E. coli* One-shot PIR2 | *pDE112 (kan)* | This Study | N/A |
| DE344 | *E. coli* MG1655 | *attTN7::pRpsL-mKate2Hyb attP21::pCymRC-BCD24-T7RNAPol* | This Study | N/A |
| DE346 | *E. coli* MG1655 | *att186 (primary)::paTc-rbs*-dCas9 attP21::pCymRC-BCD24-T7RNAPol* | This Study | N/A |
| DE348 | *E. coli* MG1655 | *att186 (primary)::paTc-rbs*-dCas9 attTN7::pRpsL-mKate2Hyb attP21::pCymRC-BCD24-T7RNAPol* | This Study | N/A |
| DE419 | *E. coli* MG1655 | *att186 (primary)::paTc-rbs*-dCas9 ΔrelA782* | This Study | N/A |
| DE420 | *E. coli* MG1655 | *att186 (primary)::paTc-rbs*-dCas9 attTN7::pRpsL-mKate2Hyb ΔrelA782* | This Study | N/A |
| DE423 | *E. coli* MG1655 | *att186 (primary)::paTc-rbs*-dCas9 attTN7::pRpsL-mKate2Hyb pDE244 (chlor)* | This Study | S3e |
| DE425 | *E. coli* MG1655 | *att186 (primary)::paTc-rbs*-dCas9 attTN7::pRpsL-mKate2Hyb pDE246 (chlor)* | This Study | S3e |
| DE426 | *E. coli* MG1655 | *att186 (primary)::paTc-rbs*-dCas9 attTN7::pRpsL-mKate2Hyb pDE247 (chlor)* | This Study | S3e |
| DE427 | *E. coli* MG1655 | *att186 (primary)::paTc-rbs*-dCas9 attTN7::pRpsL-mKate2Hyb pDE248 (chlor)* | This Study | S3e |
| DE428 | *E. coli* MG1655 | *att186 (primary)::paTc-rbs*-dCas9 attTN7::pRpsL-mKate2Hyb pDE236 (chlor)* | This Study | S3e |

| | | | | |
|---|---|---|---|---|
| DE502 | *E. coli* MG1655 | *rne-1(substitution; ts)* | This Study | 4f-h; S9b,e |
| DE562 | *E. coli* DH5α | *pDE170 (kan)* | This Study | N/A |
| DE566 | *E. coli* MG1655 | *att186 (primary)::paTc-rbs*-dCas9 attP21::pCymRC-BCD24-T7RNAPol hupA::hupA-mCherry attTN7::pRpsL-mVenus* | This Study | N/A |
| LAG436 | *E. coli* MG1655 | *att186 (primary)::paTc-rbs*-dCas9 attP21::pCymRC-BCD24-T7RNAPol hupA::hupA-mCherry attTN7::pRpsL-mVenus* | This Study | N/A |
| DE683 | *E. coli* MG1655 | *att186 (primary)::paTc-rbs*-dCas9 attP21::pCymRC-BCD24-T7RNAPol hupA::hupA-mCherry attTN7::pRpsL-mVenus ΔrelA782* | This Study | N/A |
| DE730 | *E. coli* MG1655 | *att186 (primary)::paTc-rbs*-dCas9 attTN7::pRpsL-mKate2Hyb ΔrelA782 ΔspoT(2-696)::frt-kan-frt* | This Study | N/A |
| DE735 | *E. coli* MG1655 | *att186 (primary)::paTc-rbs*-dCas9 attP21::pCymRC-BCD24-T7RNAPol hupA::hupA-mCherry attTN7::pRpsL-mVenus ΔrelA782 ΔspoT(2-696)::frt-kan-frt* | This Study | N/A |
| DE737 | *E. coli* DH5α | *pDE214 (kan)* | This Study | N/A |
| DE751 | *E. coli* MG1655 | *att186 (primary)::paTc-rbs*-dCas9 pDE223 (chlor)* | This Study | N/A |
| DE752 | *E. coli* MG1655 | *att186 (primary)::paTc-rbs*-dCas9 pDE224 (chlor)* | This Study | N/A |
| DE760 | *E. coli* MG1655 | *att186 (primary)::paTc-rbs*-dCas9* | This Study | 6a; S5g |

| | | | | |
|---|---|---|---|---|
| | | *pDE223 (chlor)*<br>*pDE214 (kan)* | | |
| DE761 | *E. coli* MG1655 | *att186 (primary)::paTc-rbs*-dCas9*<br>*pDE224 (chlor)*<br>*pDE214 (kan)* | This Study | 6a; S5g |
| DE772 | *E. coli* MG1655 | *att186 (primary)::paTc-rbs*-dCas9*<br>*pDE235 (chlor)* | This Study | N/A |
| DE773 | *E. coli* MG1655 | *att186 (primary)::paTc-rbs*-dCas9*<br>*pDE236 (chlor)*<br>*pDE214 (kan)* | This Study | 6a; S5g |
| DE785 | *E. coli* MG1655 | *att186 (primary)::paTc-rbs*-dCas9*<br>*pDE235 (chlor)*<br>*pDE214 (kan)* | This Study | 6a; S5g |
| DE787 | *E. coli* MG1655 | *att186 (primary)::paTc-rbs*-dCas9*<br>*pDE236 (chlor)* | This Study | 5f; 6b; S5h; S12b |
| DE790 | *E. coli* One-shot PIR2 | *pDE237 (kan)* | This Study | N/A |
| DE792 | *E. coli* MG1655 | *ΔrelA782* | This Study | N/A |
| DE794 | *E. coli* MG1655 | *P21::pLuxB-fusA-luxR_marionettewild* | This Study | N/A |
| DE796 | *E. coli* MG1655 | *ΔrelA782*<br>*P21::pLuxB-fusA-luxR_marionettewild* | This Study | N/A |
| DE806 | *E. coli* MG1655 | *P21::pLuxB-fusA-luxR_marionettewild*<br>*ΔfusA698* | This Study | 5f; 6b; 7e; S5h; S12b |
| DE808 | *E. coli* MG1655 | *ΔrelA782*<br>*P21::pLuxB-fusA-luxR_marionettewild*<br>*ΔfusA698* | This Study | 6e; S5h |
| DE814 | *E. coli* MG1655 | *att186 (primary)::paTc-rbs*-dCas9*<br>*pDE239 (chlor)* | This Study | 5f; 6b; 7e; S5h; S12b |
| DE815 | *E. coli* MG1655 | *att186 (primary)::paTc-rbs*-dCas9*<br>*pDE240 (chlor)* | This Study | S5h |
| DE816 | *E. coli* MG1655 | *att186 (primary)::paTc-rbs*-dCas9*<br>*pDE241 (chlor)* | This Study | 5f; 6b; 7e; S5h; S12b |

| | | | | |
|---|---|---|---|---|
| DE817 | *E. coli* MG1655 | *att186 (primary)::paTc-rbs*-dCas9*<br>*pDE242 (chlor)* | This Study | 5f; 6b; 7e; S5h; S12b |
| DE818 | *E. coli* MG1655 | *att186 (primary)::paTc-rbs*-dCas9*<br>*ΔrelA782*<br>*pDE239 (chlor)* | This Study | 6e; S5h |
| DE819 | *E. coli* MG1655 | *att186 (primary)::paTc-rbs*-dCas9*<br>*ΔrelA782*<br>*pDE240 (chlor)* | This Study | S5h |
| DE820 | *E. coli* MG1655 | *att186 (primary)::paTc-rbs*-dCas9*<br>*ΔrelA782*<br>*pDE241 (chlor)* | This Study | 6e; S5h |
| DE821 | *E. coli* MG1655 | *att186 (primary)::paTc-rbs*-dCas9*<br>*ΔrelA782*<br>*pDE242 (chlor)* | This Study | 6e; S5h |
| DE825 | *E. coli* MG1655 | *att186 (primary)::paTc-rbs*-dCas9*<br>*ΔrelA782*<br>*pDE236 (chlor)* | This Study | 6e; S5h |
| DE828 | *E. coli* MG1655 | *P21::pLuxB-fusA-luxR_marionettewild*<br>*ΔfusA698*<br>*attTN7::pRpsL-mKate2Hyb* | This Study | 5e; 7e; S12b |
| DE830 | *E. coli* MG1655 | *att186 (primary)::paTc-rbs*-dCas9*<br>*attTN7::pRpsL-mKate2Hyb*<br>*pDE239 (chlor)* | This Study | 5e; 7e; S12b |
| DE831 | *E. coli* MG1655 | *att186 (primary)::paTc-rbs*-dCas9*<br>*attTN7::pRpsL-mKate2Hyb*<br>*pDE241 (chlor)* | This Study | 5e; 7e; S12b |
| DE832 | *E. coli* MG1655 | *att186 (primary)::paTc-rbs*-dCas9*<br>*attTN7::pRpsL-mKate2Hyb*<br>*pDE242 (chlor)* | This Study | 5e; 7e; S12b |
| DE847 | *E. coli MG1655* | *att186 (primary)::paTc-rbs*-dCas9*<br>*pDE260 (chlor)* | This Study | 5f; 7e; S12b |
| DE848 | *E. coli MG1655* | *att186 (primary)::paTc-rbs*-dCas9*<br>*pDE261 (chlor)* | This Study | 5f; 7e; S12b |

| | | | | |
|---|---|---|---|---|
| DE849 | *E. coli MG1655* | *att186 (primary)::paTc-rbs*-dCas9 pDE262 (chlor)* | This Study | 5f; 7e; S12b |
| DE850 | *E. coli MG1655* | *att186 (primary)::paTc-rbs*-dCas9 pDE263 (chlor)* | This Study | 5f; 7e; S12b |
| DE851 | *E. coli MG1655* | *att186 (primary)::paTc-rbs*-dCas9 pDE264 (chlor)* | This Study | 5f; 7e; S12b |
| DE852 | *E. coli MG1655* | *att186 (primary)::paTc-rbs*-dCas9 pDE265 (chlor)* | This Study | 5f; 7e; S12b |
| DE853 | *E. coli MG1655* | *att186 (primary)::paTc-rbs*-dCas9 pDE266 (chlor)* | This Study | 5f; 7e; S12b |
| DE854 | *E. coli MG1655* | *att186 (primary)::paTc-rbs*-dCas9 pDE267 (chlor)* | This Study | 5f; 7e; S12b |
| DE855 | *E. coli MG1655* | *att186 (primary)::paTc-rbs*-dCas9 pDE268 (chlor)* | This Study | 5f; 7e; S12b |

**Supplementary Table 4: Plasmids used in this study.**

| Plasmid | Description | Source | Notes |
|---|---|---|---|
| pAJM.336 | p15a-PTac-EYFP-LacIAM (kan) | [24] | |
| pAJM.657 | p15a-PCymRC-EYFP-CymRAM (kan) | [24] | |
| pAJM.713 | p15a-PLuxB-EYFP (kan) | [24] | |
| psgRNAc | p15a-PJ23119-sgRNA_scaffold (chlor) | [8] | |
| pRelA' | pSC101-PTet-relA*-mVenus (kan) | [28] | |
| pOSIP-KT | pR6K-φ21attP-ccdB-pUC-Pλ(ts)-φ21Integrase (kan) | [27] | |
| pOSIP-KO | pR6K-phage186attP-ccdB-pUC-Pλ(ts)-phage186Integrase (kan) | [27] | |
| pEB1-mGFPmut2 | pSC101-proC-GFPmut2 (kan) | [29] | |
| pLPT41 | pBR322-pZE21-GFPaav (kan) | This Study | |
| pDHL138 | pSC101-PnlpD-nlpD-PrpoS-GFPmut2 (kan) | [20] | |
| pDHL981 | p15a-PClpP-clpP-linker-mGFPmut3 (chlor) | This Study | |
| V37m | ColE1-mScarlet (kan) | V37m was a gift from Richard | |

| | | | |
|---|---|---|---|
| | | Murray, Part of CIDAR MoClo Extension | |
| pTARA | p15a-pBAD-T7RNAPol (chlor) | [26] | |
| pdCas9-bacteria | p15a-Ptet-dCas9 (chlor) | [25] | |
| pE-FLP | pSC101ts-PE-FLP (amp) | [27] | |
| pSIM5-Tet | pSC101-Pλ(ts)-RedCassette (tet) | [30] | |
| pNDL-1 | pSC101ts-attClonase1-mTn7-attClonase2-pBAD-tnsABCD (amp) | [20] | |
| pAV203 | pSC101ts-CspRec-MutL(E32K) (amp) | This Study | |
| pDE29 | pBR322-PJ23116-gfpmut2 (kan) | This Study | |
| pDE30 | pBR322-PT7 (kan) | This Study | |
| pDE32 | pBR322-PJ23116-gfpmut2(Y66L) (kan) | This Study | |
| pDE36 | p15a-Plac-lacZalpha (chlor) | This Study | |
| pDE47 | pBR322-PT7 (kan) | This Study | Similar to pDE30, with restriction site overhangs compatible with pDE56 and other MARLIN plasmids |
| pDE56 | pBR322-PJ23119-sgRNA_scaffold (kan) | This Study | |
| pDE57 | p15a-PcymRC-t7RNAPol-CymRAM (kan) | This Study | |
| pDE91 | pOSIP-KT-PCymRC-t7RNAPol (kan) | This Study | |
| pDE93 | pBR322-PJ23119-sgRNA_scaffold-PJ23102-mScarlet (kan) | This Study | |
| pDE104 | pOSIP-KT-Plux-EYFP-LuxR_marionette_wild (kan) | This Study | |
| pDE112 | pOSIP-KT-PCymRC-BCD24-t7RNAPol (kan) | This Study | |
| pDE170 | pBR322-PJ23119-sgRNA_scaffold*-PJ23102-mScarlet (kan) | This Study | |
| pDE214 | p15a-Plac_marionette-relA* (kan) | This Study | |

| | | | |
|---|---|---|---|
| pDE223 | p15a-PJ23119-sgRNA_rplA (chlor) | This Study | sgRNA derived from oDEPool7id 4737 |
| pDE224 | p15a-PJ23119-sgRNA_rplQ (chlor) | This Study | sgRNA derived from oDEPool7id 6217 |
| pDE235 | p15a-PJ23119-sgRNA_rpsO (chlor) | This Study | sgRNA derived from oDEPool7id 18952 |
| pDE236 | p15a-PJ23119-sgRNA_control (chlor) | This Study | sgRNA derived from oDEPool7id 29672 |
| pDE237 | pOSIP-KT-Plux-fusA-PJ23119-luxR_marionettewild | This Study | |
| pDE239 | p15a-PJ23119-sgRNA_rplL (chlor) | This Study | sgRNA derived from oDEPool7id 702 |
| pDE240 | p15a-PJ23119-sgRNA_fusA (chlor) | This Study | sgRNA derived from oDEPool7id 20318 |
| pDE241 | p15a-PJ23119-sgRNA_infA (chlor) | This Study | sgRNA derived from oDEPool7id 20899 |
| pDE242 | p15a-PJ23119-sgRNA_pheT (chlor) | This Study | sgRNA derived from oDEPool7id 23961 |
| pDE244 | p15a-PJ23119-sgRNA_ftsN (chlor) | This Study | sgRNA derived from oDEPool7id 9586 |
| pDE246 | p15a-PJ23119-sgRNA_rplA (chlor) | This Study | sgRNA derived from oDEPool7id 4754 |
| pDE247 | p15a-PJ23119-sgRNA_dnaA (chlor) | This Study | sgRNA derived from oDEPool7id 8869 |
| pDE248 | p15a-PJ23119-sgRNA_mrdA (chlor) | This Study | sgRNA derived from oDEPool7id 9865 |
| pDE260 | p15a-PJ23119-sgRNA_rplL_2 (chlor) | This Study | sgRNA derived from oDEPool7id 694 |
| pDE261 | p15a-PJ23119-sgRNA_rplL_3 (chlor) | This Study | sgRNA derived from oDEPool7id 706 |

| | | | |
|---|---|---|---|
| pDE262 | p15a-PJ23119-sgRNA_rplL_4 (chlor) | This Study | sgRNA derived from oDEPool7id 711 |
| pDE263 | p15a-PJ23119-sgRNA_infA_2 (chlor) | This Study | sgRNA derived from oDEPool7id 4925 |
| pDE264 | p15a-PJ23119-sgRNA_infA_3 (chlor) | This Study | sgRNA derived from oDEPool7id 4926 |
| pDE265 | p15a-PJ23119-sgRNA_pheT_2 (chlor) | This Study | sgRNA derived from oDEPool7id 9220 |
| pDE266 | p15a-PJ23119-sgRNA_infA_4 (chlor) | This Study | sgRNA derived from oDEPool7id 20903 |
| pDE267 | p15a-PJ23119-sgRNA_pheT_3 (chlor) | This Study | sgRNA derived from oDEPool7id 23953 |
| pDE268 | p15a-PJ23119-sgRNA_pheT_4 (chlor) | This Study | sgRNA derived from oDEPool7id 23957 |
| pIDE11 | p15a-PBBa_J23116-GFPmut2lib-PT7-Barcode (chlor) | This Study | |
| pIDE15 | p15a-PBBa_J23116-GFPmut2lib-PT7-Barcode (chlor) | This Study | |
| pIDE20 | p15a-PJ23119-oDEPool7-PT7-Barcode (chlor) | This Study | |
| pIDE26 | p15a-PJ23119-oDEPool13-PT7-Barcode (chlor) | This Study | |

**Supplementary Table 5: Oligonucleotides used in this study.**

| Name | Sequence |
|---|---|
| gbMI3 | CACCGAATTCCCCGGGGGGGACAAGTTTGTACAAAAAAGCAGGCTTAGGATCCTTAAGCACCCC<br>AGCCAGATGGCCTGGTGATGGCGGGATCGTTGTATATTTCTTGACACCTTTTCGGCATCGCCC<br>TAAAATTCGGCGTCCTCATATTGTGTGAGGACGTTTTATTACGTGTTTACGAAGCAAAAGCTAAA<br>ACCAGGAGCTATTTAATGGCAACAGTTAACCAGCTGGTACGCAAACCACGTGCTCGCAAAGTTG<br>CGAAAAGCAACGTGCCTGCGCTGGAAGCATGCCCGCAAAAACGTGGCGTATCTCGAGAGATCC<br>TCTAGATTTAAGAAGGAGATATACATATGAGTAAAGGAGAAGAACTTTTCACTGGAGTTGTCCCA<br>ATTCTTGTTGAATTAGATGGTGATGTTAATGGGCACAAATTTTCTGTCAGTGGAGAGGGTGAAG<br>GTGATGCAACATACGGAAAACTTACCCTTAAATTGATTTGCACTACTGGAAAACTACCTGTTCCA<br>TGGCCAACACTTGTCACTACTTTGGGTTATGGTGTTCAATGCTTTGCGAGATACCCAGATCATA<br>TGAAACAGCATGACTTTTTCAAGAGTGCCATGCCCGAAGGTTATGTACAGGAAAGAACTATATTT<br>TTCAAAGATGACGGGAACTACAAGACACGTGCTGAAGTCAAGTTTGAAGGTGATACCCTTGTTAA<br>TAGAATCGAGTTAAAAGGTATTGATTTTAAAGAAGATGGAAACATTCTTGGACACAAATTGGAATA<br>CAACTATAACTCACACAATGTATACATCACGGCAGACAAACAAAAGAATGGAATCAAAGCGAACTT<br>CAAAATTAGACACAACATTGAAGATGGAGGTGTTCAACTAGCAGACCATTATCAACAAAATACT<br>CCAATTGGCGATGGCCCTGTCCTTTTACCAGACAACCATTACCTGTCCTACCAATCTAAGCTTT<br>CGAAAGATCCCAACGAAAAGAGAGACCACATGGTCCTTCTTGAGTTTGTAACAGCTGCTGGGAT<br>TACACATGGCATGGATGAACTATACAAATAAATGTCCAGACCTGCAGGCATGCAAGCTCTAGAGG<br>CATCAAATAAAACGAAAGGCTCAGTCGAAAGACTGGGCCTTTCGTTTTATCTGTTGTTTGTCGG<br>TGAACGCTCTCCTGAGTAGGACAAATCCGCCGCGAATTCTGCAGGTGTTAATTCAGGAGCATTG<br>TTATCAGACCAAATATGTGTAGGCTGGAGCTGCTTCGAAGTTCCTATACTTTCTAGAGAATAGGA<br>ACTTCGGAATAGGAACTTCAAGATCCCCTTATTAGAAGAACTCGTCAAGAAGGCGATAGAAGGCG<br>ATGCGCTGCGAATCGGGAGCGGCGATACCGTAAAGCACGAGGAAGCGGTCAGCCCATTCGCCG<br>CCAAGCTCTTCAGCAATATCACGGGTAGCCAACGCTATGTCCTGATAGCGGTCCGCCACACCCA<br>GCCGGCCACAGTCGATGAATCCAGAAAAGCGGCCATTTTCCACCATGATATTCGGCAAGCAGGC<br>ATCGCCATGGGTCACGACGAGATCCTCGCCGTCGGGCATGCGCGCCTTGAGCCTGGCGAACA<br>GTTCGGCTGGCGCGAGCCCCTGATGCTCTTCGTCCAGATCATCCTGATCGACAAGACCGGCTT<br>CCATCCGAGTACGTGCTCGCTCGATGCGATGTTTCGCTTGGTGGTCGAATGGGCAGGTAGCCG<br>GAT |

| | |
|---|---|
| gbMI4 | CACCGAATTCCCCGGGGGGGACAAGTTTGTACAAAAAAGCAGGCTTAGGATCCTTAAGCACCCC<br>AGCCAGATGGCCTGGTGATGGCGGGATCGTTGTATATTTCTTGACACCTTTTCGGCATCGCCC<br>TAAAATTCGGCGTCCTCATATTGTGTGAGGACGTTTTATTACGTGTTTACGAAGCAAAAGCTAAA<br>ACCAGGAGCTATTTAATGGCAACAGTTAACCAGCTGGTACGCAAACCACGTGCTCGCAAAGTTG<br>CGAAAAGCAACGTGCCTGCGCTGGAAGCATGCCCGCAAAAACGTGGCGTATCTCGAGAGATCC<br>TCTAGATTTAAGAAGGAGATATACATATGGTTAGTAAAGGAGAAGAAAATAACATGGCACTGATTA<br>AGGAGAACATGCACATGAAGCTGTACATGGAGGGCACCGTGAACAACCACCACTTCAAGTGC<br>ACATCCGAGGGCGAAGGCAAGCCCTACGAGGGCACCCAGACCATGAGAATCAAGGCCGTCGAG<br>GGCGGCCCTCTCCCCTTCGCCTTCGACATCCTGGCTACCAGCTTCATGTACGGCAGCAAAACC<br>TTCATCAACCACACCCAGGGCATCCCCGACTTCTTTAAGCAGTCCTTCCCTGAGGGCTTCACAT<br>GGGAGAGAGTCACCACATACGAAGACGGGGGCGTGCTGACCGCTACCCAGGACACCAGCCTCC<br>AGGACGGCTGCCTCATCTACAACGTCAAGATCAGAGGGGTGAACTTCCCATCCAACGGCCCTGT<br>GATGCAGAAGAAAACACTCGGCTGGGAGGCCTCCACCGAGACCCTGTACCCCGCTGACGGCGG<br>CCTGGAAGGCAGAGCCGACATGGCCCTGAAGCTCGTGGGCGGGGGCCACCTGATCTGCAA<br>CTTGAAGACCACATACAGATCCAAGAAACCCGCTAAGAACCTCAAGATGCCCGGCGTCTACTATG<br>TGGACAGAAGACTGGAAAGAATCAAGGAGGCCGACAAAGAGACCTACGTCGAGCAGCACGAGGT<br>GGCTGTGGCCAGATACTGCGACCTCCCTAGCAAACTGGGGCACAGATAAATGTCCAGACCTGC<br>AGGCATGCAAGCTCTAGAGGCATCAAATAAAACGAAAGGCTCAGTCGAAAGACTGGGCCTTTCG<br>TTTTATCTGTTGTTTGTCGGTGAACGCTCTCCTGAGTAGGACAAATCCGCCGCGAATTCTGCAG<br>GTGTTAATTCAGGAGCATTGTTATCAGACCAAATATGTGTAGGCTGGAGCTGCTTCGAAGTTCC<br>TATACTTTCTAGAGAATAGGAACTTCGGAATAGGAACTTCAAGATCCCCTTATTAGAAGAACTC<br>GTCAAGAAGGCGATAGAAGGCGATGCGCTGCGAATCGGGAGCGGCGATACCGTAAAGCACGAG<br>GAAGCGGTCAGCCCATTCGCCGCCAAGCTCTTCAGCAATATCACGGGTAGCCAACGCTATGTC<br>CTGATAGCGGTCCGCCACACCCAGCCGGCCACAGTCGATGAATCCAGAAAAGCGGCCATTTTC<br>CACCATGATATTCGGCAAGCAGGCATCGCCATGGGTCACGACGAGATCCTCGCCGTCGGGCA<br>TGCGCGCCTTGAGCCTGGCGAACAGTTCGGCTGGCGCGAGCCCCTGATGCTCTTCGTCCAGA<br>TCATCCTGATCGACAAGACCGGCTTCCATCCGAGTACGTGCTCGCTCGATGCGATGTTTCGCT<br>TGGTGGTCGAATGGGCAGGTAGCCGGATCAAGCGTATGCAGCCGCCGCATTGCATCAGCCAT<br>GATGGATACTTTCTCGGCAGGAGCAAGGTGAGATGACAGGAGATCCTGCCCCGGCACTTCGCC<br>CAATAGCAGCCAGTCCCTTCCCGCTTCAGTGACAACGTCGAGCACAGCTGCGCAAGGAACGCC<br>CGTCGTGGCCAGCCACGATAGCCGCGCTGCCTCGTCCTGCAGTTCATTCAGGGCACCGGACA<br>GGTCGGTCTTGACAAAAAGAACCGGGCGCCCCTGCGCTGACAGCCGGAACACGGCGGCATCA<br>GAGCAGCCGATTGTCTGTTGTGCCCAGTCATAGCCGAATAGCCTCTCCACCCAAGCGGCCGG<br>AGAACCTGCGTGCAATCCATCTTGTTCAATCATGCGAAACGATCCTCATCCTGTCTCTTGATCA<br>GATCTTGATCCCCTGCGCCATCAGATCCTTGGCGGCAAGAAAGCCATCCAGTTTACTTTGCAG<br>GGCTTCCCAACCTTACCAGAGGGCGCCCCAGCTGGCAATTCCGGTTCGCTTGCTGTCCATAA<br>AACCGCCCAGTCTAGCTATCGCCATGTAAGCCCACTGCAAGCTACCTGCTTTCTCTTTGCGCT<br>TGCGTTTTCCCTTGTCCAGATAGCCCAGTAGCTGACATTCATCCGGGGTCAGCACCGTTTCT<br>GCGGACTGGCTTTCTACGTGTTCCGCTTCCTTTAGCAGCCCTTGCGCCCTGAGTGCTTGCG<br>GCAGCGTGAGCTTCAAAAGCGCTCTGAAGTTCCTATACTTTCTAGAGAATAGGAACTTCGTAC<br>CCAGCTTTCTTGTACAAAGTGGTCCCCAAGCTTCTGCAGAGCT |
| oAV_V1 | GGGTTCCGCGCACATTTCCCGCGCGCCCAGCTGTCTAGGG |
| oAV_V2 | GGCTCTTGTATCTATCAGTGCACAAATTTAAATCGTAATTATTGGGGACCCCTG |
| oAV_V3 | AATTACGATTTAAATTTGTGCACTGATAGATACAAGAGCCATAAGAACCT |
| oAV_V4 | CCCTAGACAGCTGGGCGCGCGGGAAATGTGCGCGGAACCC |
| oDE071 | TCGAGCTCGGTACCCGGGGAAGCAGGGATTCTGCAA |
| oDE072 | GGTTAGGCGCCATGCATCAGAAACGCAAAAAGGCCA |
| oDE094 | TTATTTGTACAATTCATCCATACCATG |
| oDE137 | CCGCCATAAACTGCCAGGAATTGGGGATCGGAGATATCGACGTCTTAAGACCC |
| oDE138 | TTATCCATTTTTGCCACCTGCGCCCGTCATTTGATGAATTCTTTTCTCTATCACTGATAG |
| oDE139 | ATCAAATGACGGGCGCAGGTGGCAAAAATGGATAAGAAATACTCAATAGGC |
| oDE140 | AGTTTAGGTTAGGCGCCATGCATCTCGAGGCATGCCTGCAATGCCTGGAGATCCTTACTC |
| oDE154 | ATTTGTCCTACTCAGGAGAGC |
| oDE195 | CGAACGTGAGGCATCCAGGCGCGGTATCAGCTCACTCAAA |
| oDE196 | GGTGAAGACGAAAGGGCC |
| oDE197 | AGAAAGAGGAGAAATACTAGATGAGTAAAGGAGAAGAACTTTTCA |

| | |
|---|---|
| oDE199 | TGGATGAATTGTACAAATAAACGCGTGCTAGAGGCATCAAATAAAACGAAAGGCTCAGTCGAAAG ACTGGGCCTTTCGTTTTATCTGTTGTTTGTCGGTGAACGCTCTCCTGAGTAGGACAAATCCGC CGCCCTAGACCTACAGGGGAGACCTGCGAACGTGAGGCATCCAGGC |
| oDE201 | ATTACTGATGGCAATGTGAT |
| oDE202 | ATCACATTGCCATCAGTAAT |
| oDE204 | GAGGCCCTTTCGTCTTCACCCAGGTCTCCAGTCTAATACGACTCACTATAGGGACAGTAACGTT AGCTAGCCTATCAGAACATGTTACATAGAATCACATTGCCATCAGTAATAACCCCTTGGGGCCTC TAAACGGGTCTTGAGGGGTTTTTTGGGAATGAGACCTCCGAACGTGAGGCATCCAGGC |
| oDE205 | GAGGCCCTTTCGTCTTCACCGAGGTCTCATGTATTGACAGCTAGCTCAGTCCTAGGGACTATG CTAGCTCTAGAGAAAGAGGAGAAATACTAG |
| oDE207 | CCCAACGAAAAGAGGGACCACATGGTCCTTC |
| oDE208 | GAAGGACCATGTGGTCCCTCTTTTCGTTGGG |
| oDE209 | CACTTGTCACTACTTTCGCGTTAGGTCTTCAATGCTTTGCG |
| oDE210 | CGCAAAGCATTGAAGACCTAACGCGAAAGTAGTGACAAGTG |
| oDE222 | CCTGTGAGACCCCTGTAGT |
| oDE223 | TTCCGGAGACCTTCCAAGTC |
| oDE225 | AAGACTTGGAAGGTCTCCGGAAAGGATGACGATGAGCGCATT |
| oDE226 | GACTACAGGGGTCTCACAGGTTAACGACCCTGCCCTGAAC |
| oDE252 | ATCAGAACATGTTACATAGAAT |
| oDE253 | ATAGTGCTGCTAACGTTCG |
| oDE254 | GGCCCTTTCGTCTTCACCGGTCTCCTACATAATACGACTCACTATAGGGACGAACGTTAGCAGC ACTATATCAGAACATGTTACATAGA |
| oDE276 | TGCTAGAGGCATCAAATAAAACG |
| oDE277 | AAAGGGCCTCGTGATACG |
| oDE278 | GGCGTATCACGAGGCCCTTTGAGGTCTCATGTATTGACAGCTAGCTCAGTCCTAGGTATAATAC TAGTATGTCTTCAGTCTAGAAGACGAGTTTTAGAGCTAGAAATAGCAAGTTAAAATAAGGCTAGT CCGTTATCAACTTGAAAAAGTGGCACCGAGTCGGTGCTGCTAGAGGCATCAAATAAAA |
| oDE282 | CTCGGTACCAAATTCCAGA |
| oDE283 | CTAGTATTTCCCCTCTTTCT |
| oDE284 | GAAAGAGGGGAAATACTAGATGAACACGATTAACATCGCT |
| oDE285 | CTGGAATTTGGTACCGAGTTACGCGAACGCGAAGTCC |
| oDE286 | GGAATTCGAGCTCGGTACCCAATTATTGAAGGCCTCCCT |
| oDE294 | GGTTAGGCGCCATGCATCCAGATAAAATATTTGCTCATGAGCCC |
| oDE311 | ACGAACGTTAGCAGCACTATNNNNNNNNNNNNNNNNNNNNACAGTAACGTTAGCTAGCCT |
| oDE336 | TTTCTAGCTCTAAAACTCGTCTTCGTGAGCGCAACGACATAAGC |
| oDE338 | GAAGACGAGTTTTAGAGCTAGA |
| oDE339 | GAAGACATACTAGTATTATACCTAG |
| oDE342 | GTCACACAGGAAAGTACTAAATGAT |
| oDE397 | GGGCCCAAGTTCACTTAAAAAGGAGATCAACAATGAAAGCAATTTTCGTACTGAAACATCTTAATC ATGCGATGGACGGTTTCTATGAACACGATTAACATCGCTAAG |
| oDE398 | TTAAACAAAATTATTTGTAGAGGCTGTTTC |
| oDE418 | GGCACCTTAAGAAGACGAAAAC |
| oDE433 | GGAGACGCGTCTCCTAGT |
| oDE516 | C*C*GAGTCTGGAAGCTGCTTTTGTTGATTACGGCGCTGAACGTCACAGTTTCCTCCCACTAAAA GAAATTGCCCGCGAATATTTCCCTGCT |
| oDE550 | CGTACGACGGAAGACATTCTC |
| oDE553 | TTGAGGTCTCATGTCTTCATTTCAGATAAAAAAAATCCTTAG |
| oDE554 | ATGAAGACATGAGACCTCAAAGGGCCTC |

| | |
|---|---|
| oDE583 | TTAAACAAAATTATTTGTAGAGGCTGTTTC |
| oDE659 | GAAGGTCTGATCAACAACCAG |
| oDE660 | CGGTACGAATAATCGCAGAAAC |
| oDE672 | CCTCTACAAATAATTTTGTTTAAAAGAATTCAAAAGATCTAGGAGG |
| oDE673 | CTTTTCTGGAATTTGGTACCGAGATGCCTGGAGATCCTTACTC |
| oDE690 | TAGTCCTGTTACAGTACCCACTTT |
| oDE691 | TAGTTCCAGCTCGATGTAAGCCAT |
| oDE699 | AAACAAAGTGGGTACTGTAACAGG |
| oDE700 | AAACATGGCTTACATCGAGCTGGA |
| oDE716 | TAGTGGTGAGTGCTACCTGAACTT |
| oDE717 | AAACAAGTTCAGGTAGCACTCACC |
| oDE718 | TAGTACGCCTTCTTTCAGAGCAGC |
| oDE719 | TAGTATGTTACGGTAGCGTGCGAT |
| oDE720 | TAGTAGACAATGCGGCCTTTGCTC |
| oDE721 | TAGTTCCGGCGCTTCGACTGTAAT |
| oDE722 | AAACGCTGCTCTGAAAGAAGGCGT |
| oDE723 | AAACATCGCACGCTACCGTAACAT |
| oDE724 | AAACGAGCAAAGGCCGCATTGTCT |
| oDE725 | AAACATTACAGTCGAAGCGCCGGA |
| oDE736 | GCCTCTACAAATAATTTTGTTTAAGTTATCCCTTCGGAGTTTTAGTC |
| oDE737 | CTTTTCTGGAATTTGGTACCGAGAGCACGGGACTTTGGTATTAAC |
| oDE744 | CTTAAATTGAACGCCTAAAAGATAAACGAGGAAACAAATGATTCCGGGGATCCGTCGACC |
| oDE745 | TTGGTATTAACCCTTAGGCTTATTTACCACGGGCTTCAATTGTAGGCTGGAGCTGCTTCG |
| oDE749 | TAGTGACGCTCGCCGCGAAGGTGC |
| oDE751 | TAGTTTACGCATCCACGTACGTTC |
| oDE752 | TAGTGTTATCCGGACAAAACGGTT |
| oDE753 | TAGTACCAAGACGCAGAGCGGGTC |
| oDE754 | TAGTAAGGACCTAACATCCAAGGG |
| oDE755 | AAACGCACCTTCGCGGCGAGCGTC |
| oDE757 | AAACGAACGTACGTGGATGCGTAA |
| oDE758 | AAACAACCGTTTTGTCCGGATAAC |
| oDE759 | AAACGACCCGCTCTGCGTCTTGGT |
| oDE760 | AAACCCCTTGGATGTTAGGTCCTT |
| oDE775 | TAGTAAATCTTCAGCAGCTTCAAC |
| oDE776 | TAGTTGTGCTTCTTTCAGAGCAGC |
| oDE777 | TAGTTGTGTGGGCTTCAGAGCAGC |
| oDE778 | TAGTTGATGGCAACGTTTCAAGAA |
| oDE779 | TAGTTGATTGCAACGTTTCAAGAA |
| oDE780 | TAGTAGTGATGAGCAGCACATAGT |
| oDE781 | TAGTCTCAAATGCGGCCTTTGCTC |
| oDE782 | TAGTCGTTTGGTGATATCAATTAC |
| oDE783 | TAGTCGTGCCATGATATCAATTAC |
| oDE784 | AAACGTTGAAGCTGCTGAAGATTT |
| oDE785 | AAACGCTGCTCTGAAAGAAGCACA |
| oDE786 | AAACGCTGCTCTGAAGCCCACACA |

| | |
|---|---|
| oDE787 | AAACTTCTTGAAACGTTGCCATCA |
| oDE788 | AAACTTCTTGAAACGTTGCAATCA |
| oDE789 | AAACACTATGTGCTGCTCATCACT |
| oDE790 | AAACGAGCAAAGGCCGCATTTGAG |
| oDE791 | AAACGTAATTGATATCACCAAACG |
| oDE792 | AAACGTAATTGATATCATGGCACG |

**Supplementary Table 6: Calculation of Per-experiment Cost**

| Reagent | Unit Cost | Unit Amount | Amount per Experiment | Cost per Experiment |
|---|---|---|---|---|
| PCA | *$129.00* | *25g* | *0.0231g* | $0.12 |
| Trolox | *$85.70* | *1g* | *0.015g* | $1.29 |
| 20X SSC | *$64.65* | *500mL* | *15.5mL* | $2.00 |
| Methanol | *$59.80* | *1L* | *12mL* | $0.72 |
| Acetone | *$64.60* | *1L* | *3mL* | $0.19 |
| rPCO enzyme | *$592.00* | *300U* | *18U* | $35.52 |
| Nuclease free Water | *$281.00* | *4L* | *112mL* | $7.87 |
| TCEP | *$146.00* | *10mL* | *2mL* | $29.20 |
| Triton X-100 | *$59.40* | *100mL* | *0.00615mL* | $0.00 |
| Ethylene carbonate | *$38.20* | *500g* | *6.605g* | $0.50 |
| Murine RNAse inhibitor | *$335.00* | *15000U* | *2000U* | $44.67 |
| Probes | *~$589.08 Avg Cost* | *~15nmol Avg Yield* | *1.5nmol* | $58.91 |
| *Upfront Cost (10^3 barcodes)* | $7,746.15 | | *Total Cost per Experiment* | $180.99 |
| *Upfront Cost (10^6 barcodes)* | $13,636.95 | | | |
| *Upfront Cost (10^9 barcodes)* | $19,527.75 | | | |

### Supplementary Discussion

In this supplementary discussion, we reflect and expand on several of the technical and biological insights made in the main text. First, we provide our view on the future applications of MARLIN, including alternative perturbation schemes, use in synthetic biology, and extension to other model organisms. We additionally discuss our (p)ppGpp regulatory model, relating it to previous regulatory hypotheses[32–34] and quantitatively connecting it to the proteome partitioning model of Scott *et al.* 2010.[32] Finally, we re-examine our library-scale nucleoid data to lend support to a previously proposed model of nucleoid morphology control.[35]

**MARLIN Limitations and Extensions**

In this work, we focused on using MARLIN to multiplex a CRISPRi library in *E. coli* to observe the resultant effects on division and growth. While this approach yielded rich information about gene function, CRISPRi has some interpretive limitations, most significantly polar effects. Extension of MARLIN to other pooled functional genomics techniques in *E. coli*, such as TNseq, CRISPR-associated TNs, and retron recombineering, would be straightforward and provide complementary information to the CRISPRi data gathered in this work.

Critically, MARLIN enables the measurement of long-term dynamics in single cells, which enabled characterization of growth dynamics. Another setting in which long-term dynamic observations are important is the study of protein expression regulation, especially with respect to stochastic fluctuations and stress response. Careful consideration of genetic architecture has led to the development of fluorescent reporters that respond to transcriptional, translational, and post-translational regulation in a handful of cases. Combining these reporter design principles with MARLIN would enable single-cell dynamic measurements of multiple levels of regulation, for potentially thousands of proteins.

In addition to functional genomics, we believe MARLIN has applicability in the context of synthetic biology, using *E. coli* as a chassis for synthetic circuit design. Synthetic circuits such as switches and oscillators must be characterized by long time-series observations, as their dynamics are typically on a generation or longer timescale. Moreover, single-cell observations are critical for characterizing the performance of circuits, especially their noise characteristics. As such, mother machine or similar microfluidics are typically used to assess circuits, but throughput limitations preclude systematic characterization of circuit design spaces. Employing MARLIN to multiplex such circuits would thus greatly aid in circuit optimization and understanding circuit design principles.

Aside from *E. coli*, we believe MARLIN can be readily applied to other bacteria to gain similar insights into their fundamental physiology. As with *E. coli*, achieving high multiplexing performance requires high levels of RNA barcode expression. This can be most easily achieved in organisms where high-strength, inducible expression systems are available, and will greatly depend on the copy number of the genetic locus hosting the barcode expression construct. Anecdotally, we have been able to achieve multiplexing using low copy plasmids in *E. coli*, after significant optimization of expression and imaging. While not suitable for all bacteria, many non-motile rod and coccoid bacteria can be cultivated in a standard mother machine architecture, suggesting that our high-throughput device design may be simply re-used in many cases.

**Mismatch-CRISPRi and Dynamic Phenotype Analysis**

Over the past few years, a handful of papers have reported systematic imaging of genetic perturbation libraries in bacteria.[36–38] These studies used either single or multiparametric analysis frameworks, most often focusing on endpoint or steady-state phenotypes. In this work, we were interested in the temporal evolution of phenotypes as essential genes were depleted. Thus, along with taking a multiparametric clustering approach, we measured phenotypic similarity using the full time-series representation. This enabled identification of common phenotypes and their gene depletions. While we focused on the mean behavior of different genetic variants, we note that this approach can be extended to analyze phenotypic heterogeneity as well. Anecdotally, several CRISPRi knockdowns caused the population to split into a phenotypically distinct fraction and a WT-like fraction, which may be evidence of ultrasensitivity and thus of feedback. However, due to limited sampling, we lack the statistical power to detect these cases in general. Future work to design more compact libraries could enable systematic identification of these cases.

**Plausible Mechanisms for a SpoT-dependent Translation Capacity Response**

Our results support a putative role for SpoT in degrading (p)ppGpp in proportion to the number of ribosomes in the charged tRNA-bound, translocating state (Fig. 6a). The most significant findings which support this role are (1) the increase in ribosome concentration as ribosomes are slowed through EF-G/FusA depletion and (2) the invariance of ribosome concentration under reduction of the initiation rate (Fig. 5d).

It has been suggested that the bifunctional (p)ppGpp synthase/hydrolase SpoT is the principal contributor to basal (p)ppGpp regulation and that RelA has a negligible contribution.[39] Further, it has been observed that reduced tRNA charging leads to the induction of (p)ppGpp in the absence of RelA.[39] Based on this and other observations[40,41], SpoT-dependent (p)ppGpp hydrolysis has been proposed to be inhibited by uncharged tRNAs, in a *ribosome-independent* manner. Such a mechanism would result in diminished (p)ppGpp/increased ribosome levels under reduction of translocation, as the concentration of uncharged tRNAs drops from reduced demand. Under this alternative model, reducing the rate of translation initiation should also promote increased tRNA charging, leading to a SpoT-dependent increase in RNA/Protein and reduction in (p)ppGpp. However, while a reduction in (p)ppGpp does occur under these conditions, it appears to be RelA-dependent (Fig. 6b) and is not corroborated by a major change in the RNA/Protein ratio, relative to the EV control. As such, we favor a model in which (p)ppGpp is both produced and degraded in a SpoT and *ribosome-dependent* manner. We attribute the small and RelA-dependent reduction in (p)ppGpp under reduced initiation to reduced RelA activity arising from high tRNA charging.

The mechanism by which the translocation rate is transduced into a change in (p)ppGpp via SpoT remains unclear. While direct binding of SpoT to translating ribosomes would be the simplest model for these observations, such an interaction has not been demonstrated, though SpoT has been found associated with both pre-50S particles[42] and the ribosome-associated GTPase CgtA.[43] Interestingly, pre-50S ribosomes are enriched during chloramphenicol treatment,[44] which reduces the elongation rate in a similar manner to EF-G/FusA knockdown (see "Nucleoid Morphology Under Elongation Reduction").

Another possibility is that reduction in the translocation rate provokes an accumulation of 70S ribosomes and polysomes which themselves precipitate a response that reduces (p)ppGpp levels. 70S ribosomes and polysomes have been found to bind to and inhibit the activity of RNase E; correspondingly, knockdown of RNase E results in filamentation and a reduced nucleoid to cell area ratio, which bears some similarity to morphological changes accompanying reduced translocation (compare Extended Data Fig. 6c, *RplA* and Extended Data Fig. 10a). Additional characterization of the cell state under reduced translocation, including identifying conditional interaction partners of SpoT, will be required to identify the molecular basis of this response.

Interestingly, RelA/SpoT homologous (RSH) enzymes also present in almost all bacterial species[45,46], suggesting that the ability to sense reduced translocation may be widely conserved. Consistent with this, evidence for upregulation of ribosome synthesis under reduced elongation has been uncovered in *B. subtilis*[47] and *H. influenzae*.[48] Replicating our study of ribosome regulation under different classes of translation perturbation, in the presence or absence of RSH enzymes, would provide the first step in determining if this behavior is indeed general.

**Global Proteome Control by Ribosome Kinetics**

The proposed mechanism for (p)ppGpp regulation by the ribosome-RelA-SpoT system provides an account for how the stringent response may be modulated by both amino acid supply (via RelA) and the translocation rate (via SpoT). Previously, this mechanism has been shown to provide an account for the well-known linear relationship between growth rate and ribosome concentration across differing nutrient conditions.[49] We expand on this work to show that it also explains the linear increase in ribosome content $R$ with respect to a decrease in the growth rate $\lambda$ under conditions of impeded elongation of the form:

$$R = R_{max} - k\tau_D\lambda \qquad (3)$$

where $\tau_D$ is the dwell time and k is a constant defined in the Supplementary Theory. In this view, the slope of the line reflects a linear scaling between the (p)ppGpp concentration and the elongation rate. Further, this slope is rendered steeper under conditions in which the ribosome dwell time $\boldsymbol{\tau_D}$ is higher, i.e., when there is a more limited supply of charged tRNAs (Supplementary Theory). This explains why $R$ curves constructed by chloramphenicol (cam) treatment are steeper in poor medium, previously leading to the terminology of "nutritional capacity".[32] In fact, this simple mechanism can reproduce all the features of the coarse-grained ribosome allocation model proposed in Scott *et al*. 2010, reparametrized in terms of the ribosome dwelling and translocation times (Supplementary Theory). Qualitatively, this accomplishes balance between the supply of charged tRNAs, which control dwell time, and the overall kinetics of ribosome translocation. Thus, this scheme provides a means of balancing amino acid and protein flux irrespective of other regulatory details.

**Nucleoid Morphology Under Low Translation Elongation Rates**

Our results indicate a strong degree of association between nucleoid compaction and perturbations to ribosome elongation (Extended Data Fig. 6c,g,h). The nucleoid morphology we observe bears similarity to reported nucleoid morphology under chloramphenicol (cam) treatment[35,50,51], which also acts to block elongation[52]. While several proposals have been made to explain this phenomenon[35,53–55], we will discuss two here: the Transertion Hypothesis and Nucleoid-Ribosome Mixing.

According to the Transertion Hypothesis, nucleoid morphology is determined by direct coupling of the chromosome to the IM via co-transcriptional translation and insertion of proteins into the IM.[54,55] Under this model, reduction in translation breaks these connections and produces nucleoid compaction, as we observe in the EF-G/FusA knockdowns. However, this is inconsistent with the observation that inhibition of translation initiation by InfA knockdown produces expanded nucleoids (Extended Data Fig. 6h), which also occurs when initiation is inhibited by kasugamycin (ksg).[35] Additionally, we observe no broad nucleoid compaction when directly inhibiting protein insertion via knocking down SRP or the Sec system (Extended Data Fig. 6f). Thus, Transertion is likely not the primary determinant of the compact nucleoid morphology observed under cam treatment.

Nucleoid-ribosome mixing has been proposed as an alternative and complementary mechanism to account for nucleoid compaction under cam treatment.[35] Under this model, conversion of 30S and 50S ribosomes to 70S polysomes promotes nucleoid compaction through a series of entropic and excluded-volume effects. This is consistent with our observation that reduction in elongation rate by EF-G/FusA knockdown, which promotes the formation of polysomes, results in compact nucleoids (Extended Data Fig. 6h). It also accounts for the expansion of nucleoids we observe under knockdown of the initiation factor InfA, which is expected to decrease the number of 70S polysomes (Extended Data Fig. 6h). Finally, nucleoid compaction appears to occur under reduction of tRNA charge and thus a reduced elongation rate in both *ΔrelA spoT*$^{+}$ and *ΔrelA ΔspoT* strains (Extended Data Fig. 6d,e), consistent with compaction occurring due to polysome formation and not a downstream effect of (p)ppGpp on global cell state. Overall, our data broadly support a model in which nucleoid morphology is primarily determined by nucleoid-ribosome mixing.

### Supplementary Note S1: Additional Cluster Details

In the main text, we focused our discussion of clustering on a handful of novel functional annotations and briefly outlined the phenotypic justification for each such assignment. Here, we expand on this to individually discuss each of our 13 curated clusters to both provide a comprehensive view of our dataset for interested researchers as well as a rationale for each gene function label we assign to a group. We show that each of these 13 groups presents a stereotyped phenotype within our landscape that can be rationalized with gene function. Further, we provide more complete phenotypic descriptions and background knowledge regarding the many biological hits identified in the main text. Finally, we examine the unexpected relationships between gene function and gross physiology leading to our clusters, and what each suggests about the coupling of fundamental cellular processes. In short, this note intends to explain the links between molecular functions and our phenotypes to better understand the features captured by clustering and how they might be leveraged in later screens to, e.g., annotate new genes or discover new physiological rules.

**Division Asymmetry Characterizes Nucleoid Segregation and Divisome Placement Genes**

We first discuss the three clusters associated with cell division, focusing on the distinction between division and division site placement clusters as well as the surprising association of anionic phospholipids with division. In the main text, we determined that, among our filamentous clusters, three exhibited no growth defect, which we subsequently annotated as defective in cell division (Fig. 4a and Extended Data Fig. 8a). To determine more specific functional annotations for each of these clusters, we inspected their gene membership. We noted that two of them contained all of the genes we associated with direct functions in promoting division, including members of the Divisome (*ftsB, ftsK, ftsN, ftsA,* and *zipA*) and most of the dcw cluster (O1-*murC* and O1-*lpxC*). In fact, excepting the *ftsEX* genes, which are non-essential under our growth conditions,[56] our division clusters captured all members of the Divisome as hits, suggesting that this identification was saturating for essential division genes. We correspondingly labeled these groups "Division I" and "Division II", reflecting their members direct role in promoting cell division.

While the third filamentous cluster exhibited increased length and normal growth, it also exhibited large errors in septum placement, leading to the label "Septum Error" (Extended Data Fig. 8a). In *E. coli*, errors in septum placement are principally associated with defects in Divisome localization. Proper Divisome placement involves both (1) nucleoid exclusion of FtsZ polymerization and (2) the Min system, which functions to inhibit FtsZ polymerization at the cell poles.[57–60] Consistent with this, we detected both a Min system gene (*minC*) as well as several genes involved in nucleoid occlusion via segregation (the O10-*mukB* operon, *parC,* and *parE*) within the "Septum Error" cluster. Notably, our ability to capture all major division localization systems in this group relied on time-lapse imaging to observe division asymmetry, which cannot be measured in snapshots, demonstrating a class of phenotype uniquely captured by MARLIN.

In addition to elevated septum error, the nucleoid morphologies within this group also diverged from the polynucleate morphologies of "Division I" and "Division II", which are discussed in the main text. However, upon inspection, the nucleoid morphologies in this group were consistent with the annotated functions of each gene. Depletion of MinC leads to several off-center divisions apparent in a representative kymograph (Extended Data Fig. 8f) without any change in the nucleoid, as expected of a defect purely impacting Divisome localization. On the other hand, MukB, ParC, and ParE, which promote segregation, exhibit a decrease in $N_n$. This reflects consolidation of replicated DNA into larger, unsegregating nucleoid masses and the formation of anucleate cells, as seen in a ParC knockdown (Fig. 3d). Incidentally, these anucleate cells, which form at a rate of 5% in some MukB mutants, were the object of an early microscopy screen which originally discovered the function of MukB.[61,62] This shows that MARLIN can recapitulate striking nucleoid phenotypes from other studies, without being limited to strictly focusing on those phenotypes. Further, while nucleoid information was not used for clustering in this study, as it was not available for each sgRNA, these observations provide proof of concept for making use of this information in future work. We anticipate that developments in imaging speed and resolution will facilitate both finer clustering of nucleoid phenotypes and a better understanding of the mapping of molecular function to nucleoid architecture.

**Depletion of Anionic Phospholipids Leads to Increased Cell Length**

While these division-deficient clusters clearly recapitulated known biology, we were also interested in finding genes with unexpected phenotypes, given their known molecular function, to identify underappreciated determinants of cell division. Previously, the major phospholipid (PL) - phosphatidylethanolamine (PE) - has been associated with cell division, through genetic data associated with its synthase PssA. Specifically, cells bearing temperature-sensitive *pssA* alleles become filamentous at the non-permissive temperature and contain an abnormal enrichment of cardiolipin (CL) in their inner and outer membranes.[63,64] Consistently, we find *pssA* in one of our division clusters (Extended Data Fig. 8a,c). On the other hand, *anionic* phospholipids such as phosphatidylglycerol (PG) and CL have not been associated with division defects. Moreover, cells bearing a null allele for *pgsA*, which is the upstream enzyme responsible for PG and CL synthesis, do not exhibit division defects.[65] Surprisingly, we also find *pgsA* among our division-defective clusters, with *pgsA* knockdown resulting in strongly increased cell length (Extended Data Fig. 8a,c).

Seeking to explain the division defect in each of these strains, we offer two possible explanations. The first is that in both cases, the physical properties of the inner and outer membranes are impacted such that the membrane remodeling during septation is less energetically favorable and inhibits the division process. An alternative explanation draws on a controversial hypothesis[66] that CL, which normally localizes to the cell poles, assists in mediating proper localization of the MinCDE system, possibly through an interaction with MinD. Under this model, it stands to reason that improper localization of CL (as in *pssA* defects) or the complete absence of CL (as in *pgsA* defects) would disrupt FtsZ localization and delay division as we see in a *minE* knockdown (Fig. 4a). Additional *pssA* and *pgsA* knockdown experiments with fluorescent reporters for Min system and FtsZ localization will be needed to test these hypotheses.

**The SpoVR Homolog *ycgB* Plays a Role in Division**

Excepting the association of anionic phospholipid disruption with division defects, the other novel annotations gleaned from our division clusters were for genes of unknown function. Within the "Division I" cluster, we detected four genes of unknown function with unambiguous division disruption phenotypes exhibiting polynucleate filaments: *yagH*, *ybcN*, *ycgB,* and *yecM* (Fig. 4a). Two of these genes (*yagH* and *ybcN*) are in prophages, leaving open the possibility that filamentation occurs indirectly through induction of the prophage instead of the direct activity of these gene products. Interestingly, YcgB is annotated as a homolog of the protein SpoVR, which plays an uncharacterized role in cortex formation during *B. subtilis* sporulation and is widely distributed in bacteria.[67] Spore cortex formation, like division, is primarily a process of peptidoglycan (PG) synthesis. Correspondingly, many other "SpoV" genes involved in cortex formation (*spoVB/murJ, spoVD/ftsI*, *spoVE/ftsW* are homologous to the enzymatic machinery of the Divisome. We conjecture that YcgB plays a role in septal PG synthesis during division, functionally equivalent to the role of SpoVR in cortex formation.

We also find, within the "Division II" cluster, an additional three genes of unknown function which, through filamentous, did not appear polynucleated in our HU-mCherry library. It is therefore likely that these genes do not impact the Divisome directly, but rather affect a related process, such as division precursor synthesis or chromosome duplication. Interestingly, *yeiH* is a predicted inner membrane (IM) protein, while a homolog of *yhcN* (BhsA) is a known outer membrane (OM) protein. Considering that division disruption accompanies defects in PL synthesis (*pssA* and *pgsA*), we speculate that depletion of these proteins may provoke functionally similar changes in membrane composition. Overall, we validated our division clusters by showing that they recapitulate nearly all essential factors involved in the Divisome, clarified that anionic phospholipid production is indeed required for proper division, and highlighted several genes of unknown function with division roles.

**Different Functions in Replication Distinguished by Cell Width**

Following our examination of the division-deficient clusters, we also compared the four clusters we annotated as defective in DNA replication in the main text, to determine what aspects of cell phenotype led to their distinction from each other. Interestingly, we found that these four groups were primarily distinguished by differences in cell width. To gain insight into the specific role in replication that was associated with different changes in width, we examined the genes in these clusters. The first group was characterized as having no change in width (Extended Data Fig. 8a) and was enriched for members of the Replisome (*dnaG, ssb, holA, holB, dnaN,* and *dnaX*) leading to the label "Replisome". Contrasting this, the next two clusters exhibited a notable decrease in cell width (Fig. 4c and Extended Data Fig. 8a) and contained much of the replication initiation machinery (*dnaA, dnaB,* and *dnaC*) as well as initiation-affecting genes (*gyrA* and *gyrB*).[68] Given their association with replication initiation, we designated these two clusters "Rep Init I" and "Rep Init II". Finally, we found that the last DNA replication cluster exhibited increased cell width (Extended Data Fig. 8a) and was composed of genes associated with synthesis of the dTTP DNA precursor (*thyA*, *folA*, *folC,* and *tmk*). Consistent with this, previous studies have associated an increase in cell width with defective ThyA activity[69,70]; as such, we designated this group with the label "ThyA-Like". In summary, the broad pattern among DNA replication defects was that decreasing dTTP precursor supply increased width, inhibiting elements of the replisome produced no change in width, and reducing replication initiation decreased width, which we leveraged to resolve these processes into distinct groups. Seeking an explanation for this pattern, we observe that reduced dTTP slows the replisome and promotes an increase in ploidy near *oriC*[70,71] while reducing the initiation rate decreases this ploidy, suggesting a relation between width and the structure of the chromosome that we discuss in the next section.

While genes directly involved in replication generally exhibited a reduced NC ratio and $N_n$, consistent with a cessation of replicative activity, some members of our replication clusters did not. However, many of these variants have previously been associated with disrupted replication, including *rpoC*[72,73] in the Replisome cluster and *rho/nusG*[74] in the Rep Init II cluster. We speculate that the expanded nucleoids we see in these variants result from reduced RNA synthesis, which is known to result in nucleoid expansion.[75] We therefore arrive at the surprising conclusion that defects in replication can in some cases be better captured by growth/morphology patterns than even direct labeling of the chromosome, illustrating the power of associative phenotyping.

While most of our replication-related genes have previously annotated functions, we found the gene of unknown function *yejF* in the Rep Init II cluster (Extended Data Fig. 8a). Homology-based annotation of YejF suggests it is an ATP-binding subunit of an oligopeptide transporter,[76] making its role in DNA replication unclear without further interrogation. We observed that YejF has a nucleoid phenotype similar to that of its cluster-mates, the transcription termination factor *rho* and the associated termination factor *nusG*, as well as *rpoC*. As previously discussed, we associate this phenotype with reduced RNA synthesis, suggesting YejF disruption leads to a perturbation in this process.

**An Unknown Mechanism Scales Width with Nucleoid Complexity**

While our clustering leveraged differences in cell width to classify different types of disruptions to DNA replication, the mechanistic origin of width variation in these groups remains unclear. Previously, it was observed that limitation of dTTP synthesis provokes an increase in cell width, increased chromosome duplication time, and a reduced DNA synthesis rate, consistent with the increased width we observe in the ThyA-Like cluster (Extended Data Fig. 8a).[69–71] More recently, a set of studies have proposed a physiological rule that, in *E. coli*, width scales positively with the nucleoid complexity ratio, defined as the amount of DNA associated with a single terminus.[70] This has been shown under various degrees of thymine limitation, where the nucleoid complexity ratio is increased by slowing the Replisome and promoting a higher multiplicity of replication forks, which precipitates a corresponding increase in cell width according to the rule.

However, it is not known whether the nucleoid complexity rule generalizes to conditions of altered ploidy other than reduced thymine, which has significant implications for how we interpret this rule mechanistically. In contrast to thymine limitation, inhibition of replication initiation is known to decrease the nucleoid complexity ratio, as ongoing replication events go to completion and leave complete circular chromosomes with no replication forks.[77] In this case, the nucleoid complexity rule would predict a *decrease* in cell width. Indeed, knockdowns in our library associated with the initiation machinery (*dnaA*, *dnaB*, and *dnaC*) consistently exhibit a decrease in cell width (Fig. 4c and Extended Data Fig. 8a). Hence, the nucleoid complexity rule is more general than previously appreciated, applying to both thymine limitation and reduced initiation and implying that the rule is a consequence of altered ploidy, rather than a thymine-specific effect. Though, it is still unknown whether this arises from differences in gene dosage, regulation by the replication machinery, or the biophysical properties of the nucleoid.

**Disruption of the Lol System Leads to Plasmolysis and Lysis**

Finally, we inspect the six clusters we implicated in synthesis of the cell envelope, discussing the distinction between clusters involved in weakening vs. strengthening the envelope, the phenotypic uniqueness of Lol system defects, and evidence for the increasingly appreciated load-bearing role of the outer membrane (OM). As discussed in the main text, we found 4 clusters with increased width which contained several genes involved in envelope synthesis (Extended Data Fig. 8a). Generally, genes in these groups are responsible for synthesizing load-bearing elements of the envelope such as the OM or the cell wall, and their depletion results in an expected increase in width due to the loss of envelope rigidity.[78–80]

Examining these groups more closely, they were further distinguished from each other by either the magnitude of the cell width defect or the measured septum placement error. Two of these clusters exhibited a moderate increase in cell width with no septum placement defect and contained genes involved in diverse envelope synthesis processes, including PG synthesis (*murA*, *murB*, and *murI*), LPS synthesis/trafficking (*lpxA*, *lpxB*, *lpxD*, *lptA,* and *lptG*), and production of the PG/LPS precursor UDP-GlcNAc (*glmM*, *glmS, dxs, dxr, ispH*) (Extended Data Fig. 8a). On this basis, we named these two clusters "Envelope I" and "Envelope II". In contrast to these groups, the third wide-cell cluster was characterized by more extreme increases in width, with cells in this group essentially reaching the width limit imposed by confinement within our microfluidic device. This group was solely composed of core members of the Elongasome (*mrdAB* operon O13 and *mreBCD* operon O14), a multi-protein complex responsible for incorporating new strands of PG into the cell wall, leading us to label this group as the "Elongasome" cluster.

The last wide-cell group differed from the preceding three by a large increase in the measured error in septum placement. Examining the genes in this group, we found that they corresponded to elements of the Lol system (*lnt, lspA*, *lolA*, and the *lolCDE* operon), which is responsible for trafficking lipoproteins into the OM, leading us to label this cluster "Lol System". Since septum placement is not a known function of the Lol system, we inspected Lol system knockdowns to confirm our measurement (Extended Data Fig. 8f). However, in disagreement with our quantification, direct inspection of cell kymographs in this group did not reveal obvious septum placement defects. Instead, we observed many instances in which cell cytoplasm signal disappeared in consecutive timepoints, indicative of lysis, which often accompanies Lol disruptions (Extended Data Fig. 8f).[81,82] The apparent increase in septum error characteristic to this group may thus be explained by high rates of lysis, which can drive false detection of asymmetric divisions.

In addition to increased lysis, our inspection of the "Lol System" cluster revealed indentations in the cytoplasm periphery (Extended Data Fig. 8f, *bottom*), which are indicative of plasmolysis, a process by which local bulges in the periplasm provoke increased separation of the cytoplasm from the OM. This is consistent with observations from *V. cholerae* where disruption of the Lol system resulted in the appearance of plasmolysis sites, which then developed into extrusions of OM material known as outer membrane vesicles (OMVs).[83] In *E. coli*, loss of the major Lol target protein Lpp, which anchors the cell wall to the OM, results in an increase in OMV production.[84] Based on our data, we hypothesize that OMV production in *E.* coli follows the same developmental pattern as in *V. cholerae*, nucleating nascent OMVs at the observed plasmolysis sites.

**Cell Width Decreases Associated with Disruptions that Increase OM Rigidity**

Finally, in addition to groups with increased cell width, we were surprised to find 2 clusters exhibiting a *decrease* in cell width, in the absence of changes in length and growth rate (Extended Data Fig. 8a). Correspondingly, we labeled these final groups "Narrow I" and "Narrow II". Interestingly, we found that these groups contained many genes involved in envelope synthesis, including several directly involved in OM synthesis, including *rfaE*, *lpcA*, the *waaFCL* operon O19, *ftsH*, and *wcaJ* (Extended Data Fig. 8a). Broadly, these targets are involved in exopolysaccharide synthesis, with most playing a role in LPS synthesis, while *wcaJ* participates in colanic acid synthesis.

To rationalize these observations, we note that it is increasingly appreciated that LPS plays a major role in determining OM stiffness, contributing to its roles in both load-bearing and cell-shape determination.[79,80] Hypomorphic *mreC* alleles, which possess cell-shape defects due to low cell wall synthesis, can be suppressed by a reduction in function of the essential protease FtsH.[80] This drives stabilization of the FtsH target LpxC and a resultant overproduction of LPS, increasing OM stiffness. The decrease in cell width we observe after FtsH depletion may therefore be due to an increase in OM stiffness from LPS overproduction. Consistent with this model, other knockdowns associated with increased OM stiffness, such as *waaC*[85], produce the same effect (Extended Data Fig. 8a). Cell width also decreases under knockdown for RfaE or LpcA, which synthesize the ADP-L-glycero-β-D-manno-heptose precursor used for all inner core heptose additions. These results demonstrate that modification of OM stiffness can not only rescue cell shape defects, but also modify cell width in both the positive direction by eliminating LPS synthesis and the negative direction by increasing LPS synthesis or modifying LPS structure (Extended Data Fig. 8a). How this relates to and interacts with other LPS functionality, such as determining OM permeability to antibiotics, remains to be explored.

To conclude this note, the evidence presented here suggests that our clustering approach captures extremely subtle distinctions in gene function, by leveraging several underappreciated links between function and cell phenotype. While we focus on well-characterized essential genes, we anticipate that the essential function landscape outlined here can be used in later MARLIN iterations to (1) further understand the physiological impacts of genes with poorly annotated functions and (2) pinpoint the functional impact of more complex perturbations, such as antibiotics. Moreover, by developing this approach in other bacteria, and comparing their phenotypic landscapes, we hope MARLIN will also help identify universal physiological rules such as e.g., the nutrient and ribosome growth laws and help uncover their underlying mechanisms.

### Supplementary Note S2: Theory Summary

**(p)ppGpp Control of Ribosomes**

It has previously been shown that ribosome concentration shows positive-linear scaling with growth rate, under varying nutrient conditions, and negative-linear scaling under varying concentrations of the elongation-targeting antibiotic chloramphenicol.[32] In addition, our data suggests that ribosome levels do not vary under growth defects incurred by inhibiting translation initiation. Here, we show how all three of these behaviors may be derived from the (p)ppGpp regulatory model discussed in the main text.

Under this model, (p)ppGpp synthesis is hypothesized to occur in proportion to the concentration of uncharged ribosomes $R_U$ competent to bind an uncharged tRNA and stimulate RelA, while (p)ppGpp degradation occurs in proportion to the concentration of charged ribosomes $R_C$ with a charged A-site tRNA awaiting translocation. The steady state concentration of (p)ppGpp $g$ is thus the ratio of these concentrations, scaled by a constant $k$ equal to the ratio of rate constants associated with synthesis and degradation:

$$g = k \cdot \frac{R_U}{R_C} = k \cdot \frac{\tau_D}{\tau_T}$$

During elongation, ribosomes switch from the uncharged to charged state with an average dwell time $\tau_D$, followed by switching back into the uncharged state once again after an average translocation time $\tau_T$. At equilibrium, elongating ribosomes obey detailed balance and satisfy the condition $R_U\tau_D^{-1} = R_C\tau_T^{-1}$, meaning that we may freely substitute ribosome concentrations with the kinetic rates of dwelling and translocation; this form of $g$ will thus allow us to connect (p)ppGpp levels to translation kinetics. In comparison, a simple first-order model that has SpoT-dependent (p)ppGpp synthesis repressed by charged ribosomes leads to the following steady-state $g_{Syn}$:

$$g_{Syn} = -k \cdot R_C + C = -k \cdot R_{active}\left(1 + \frac{\tau_D}{\tau_T}\right)^{-1} + C$$

where we have used detail balance and the identity $R_{active} = R_U + R_C$. Notably, this model results in a dependence of $g_{Syn}$ on the concentration of active ribosomes $R_{active}$, which contradicts the observed invariance of ribosome levels under reduced initiation. As such, we favor the first model, i.e., stimulation of SpoT hydrolysis by charged ribosomes.

Previously, the behavior of this model was analyzed under variable dwelling time, which was associated with conditions of varying nutrient quality and tRNA charging.[49] The form of $g$ under variable dwell time was given as a function of the overall translation elongation rate $\varepsilon = (\tau_D + \tau_T)^{-1}$, with the maximum elongation rate $\varepsilon_{max} = \tau_T^{-1}$ associated with the limit $\tau_D \rightarrow 0$:

$$g_{Dwell}(R, \varepsilon) = k \cdot \left(\frac{\varepsilon_{max}}{\varepsilon} - 1\right)$$

We extend this model to varying initiation and translocation rates by considering model behavior when tRNA charging is already high – i.e., under a reference condition of rapid growth. Under these conditions, dwell time will be primarily limited by the diffusion of ternary complexes[86] and only weakly dependent on tRNA charge. Therefore, despite the expected increase in tRNA charge under initiation defects, the resulting change in dwell time will be small while the translocation time will be unaffected. As such, (p)ppGpp levels would be expected to remain constant under initiation knockdown:

$$g_{Init}(R, \varepsilon) = g_{ref}$$

Finally, modeling translocation defects by varying $\tau_T$ under a fixed dwelling time $\tau_D^0$, we arrive at the following relationship between (p)ppGpp levels and the elongation rate:

$$g_{Elong}(R, \varepsilon) = c \cdot \varepsilon \tau_D^0$$

Previously, it was shown that the relationship between ribosome levels $R$ and growth rate $\lambda$, under variable dwell time (i.e., obeying $g_{Dwell}$) could be obtained by satisfying a system of equations including the (p)ppGpp repressive function on ribosomes $R(g)$ and the dependence of growth rate on the ribosome concentration and elongation rates $\lambda(R, \varepsilon)$. We adopt a similar approach here to model $R(\lambda)$ under variable initiation and translocation. Motivated by the empirical scaling of (p)ppGpp with ribosome levels[87] we assume it acts as a simple monovalent repressor of ribosome synthesis:

$$R(g) = \frac{k}{k + g} \cdot (R_{max} - R_0) + R_0$$

As alluded to in the main text, this form follows from a monovalent model of ribosomal promoter repression and the fact that ribosomes are diluted through first-order kinetics by cell growth, making ribosome levels proportional to ribosomal promoter activity.

The growth rate is proportional to the product of the number of active ribosomes $(R - R_{inact})$ engaged in translation and the elongation rate:

$$\lambda(g) = a \cdot \varepsilon(g) \cdot (R(g) - R_{inact})$$

In the simplifying case where $R_{inact} = R_0$ – which appears to be satisfied over most of the range of growth rates examined – solving this system of equations yields $R(\lambda)$ under both varying initiation and translocation. In the main text, we treat the special case of this approximation where $R_{inact} = R_0 = 0$, but emphasize that our results apply so long as $R_{inact} = R_0$ is satisfied.

Combining this with the previously derived form under varying dwelling time results in the following three functions:

$$R_{Dwell}(\lambda) = R_0 + \varepsilon_{max}^{-1}\lambda \qquad (Varying\ Dwelling)$$

$$R_{Init}(\lambda) = R_{ref} \qquad (Varying\ Initiation)$$

$$R_{Elong}(\lambda) = R_{max} - \frac{c\tau_D}{ak}\lambda \qquad (Varying\ Elongation)$$

**Scaling of Unregulated Proteins**

We hypothesize that the set of proteins whose activities are limiting for division are only passively regulated by (p)ppGpp, influenced primarily through competition with (p)ppGpp-regulated transcripts for the translation machinery. To formalize how (p)ppGpp regulation will impact this set of proteins and modulate size, we use a conventional proteome allocation framework.[32,88] In this framework, the cell's total proteome is partitioned into functionally distinct "sectors," each occupying a fraction $\phi_i$ of the total proteome. Each such sector is further divided into a basal $\phi_i^0$ and an additional $\Delta\phi_i$ allocation capturing any changes from this reference. We will specify the following two coarse-grained proteome sectors: a (p)ppGpp-repressed sector $\phi_C$ and an unregulated sector $\phi_{NC}$ – which we hypothesize contains the set of limiting division proteins assigned to a sub-sector $\phi_D$. Due to the finite size of the proteome, additional $\phi_C$ will come at the expense of $\phi_{NC}$ with each sharing a fixed 'flexible' portion of the proteome $\Delta\phi^{max}$ according to $\Delta\phi_C + \Delta\phi_{NC} = \Delta\phi^{max}$.

To link the fraction of unregulated proteins $\phi_{NC}$ directly to the growth rate, we first introduce the following relationship between the ribosome fraction $\phi_R$ – which depends on growth as described previously – and the overall fraction of genes repressed by (p)ppGpp $\phi_C$:

$$\Delta\phi_C(\lambda) = f_R^{-1}\Delta\phi_R(\lambda)$$

Or equivalently:

$$\phi_C(\lambda) = f_R^{-1}\Delta\phi_R(\lambda) + \phi_C^0 = a_1\phi_R(\lambda) + b_1$$

In other words, we assume that the change in the $\phi_C$ sector is linearly proportional to the change in the $\phi_R$ sector, according to a constant $f_R^{-1}$. We may obtain this condition if we assume that (p)ppGpp-regulated genes are controlled according to functions of the same form as the ribosome repression function $R(g)$ introduced earlier – with the (p)ppGpp binding constant $k$ held constant, i.e., assuming (p)ppGpp affinity for RNAP is not strongly impacted by promoter sequence. Combining this result with the constraint of a finite proteome produces the following dependence of $\phi_{NC}$ on growth rate:

$$\Delta\phi_{NC}(\lambda) = \Delta\phi^{max} - f_R^{-1}\Delta\phi_R(\lambda)$$

We may specify $\Delta\phi_R(\lambda)$ according to the linear relationship of ribosome concentration and growth rate $R(\lambda)$ under tRNA charge depletion and elongation inhibition, described in the previous section:

$$\phi_{NC}^{Dwell}(\lambda) = \phi_{NC}^0 + \Delta\phi^{max} - f_R^{-1}\beta_{tRNA}\cdot\lambda \qquad (Varying\ Dwelling)$$

$$\phi_{NC}^{Init}(\lambda) = \phi_{NC}^0 + \Delta\phi^{max} - f_R^{-1}\Delta\phi_R^{Ref} \qquad (Varying\ Initiation)$$

$$\phi_{NC}^{Elong}(\lambda) = \phi_{NC}^0 - f_R^{-1}\beta_{Elong}\cdot\lambda \qquad (Varying\ Elongation)$$

where each $\beta$ refers to the slope of the corresponding ribosome fraction-growth relationship $\phi_R(\lambda)$. In general, all three equations correspond to a linear form:

$$\phi_{NC}^{Class}(\lambda) = a_2^{Class}\cdot\lambda + b_2^{Class}$$

with perturbation-specific slopes and intercepts.

**Negative Hyperbolic Size-Growth Scaling**

Finally, to link the abundance of division proteins to cell size, we will need a rough model for cell division. To this end, we will require that the division volume $V_D$ scales inversely with the concentration of division proteins $\phi_D$ – which being a subset of the unregulated sector $\phi_{NC}$ is equivalent to:

$$V_D \propto 1/\phi_{NC} = 1/(1-\phi_C)$$

Note that this criterion is generally obtained for the division accumulator family of size control models – while replication-driven models exhibit a more complex dependence of $V_D$ on the abundance of the replication-initiating protein and the growth rate. This condition will hold for the average cell volume $V$, which is proportional to $V_D$ for cells growing at steady state – with the exact proportionality set by the conditions of observation, e.g., a sample from liquid culture vs. trapped lineages (Supplementary Theory).

We integrate our proteome model with this size-concentration relation – substituting $\Delta\phi_C$ with $f_R^{-1}\Delta\phi_R$ – to arrive at the following negative hyperbolic function describing the explicit dependence of cell size on ribosome concentration:

$$\frac{V}{V_0} = \left(1 - \theta \cdot \frac{\Delta\phi_R}{\phi_R^{max}}\right)^{-1} \qquad \theta = \frac{f_R^{-1}\phi_R^{max}}{\phi_{NC}^{max}} \qquad \phi_i^{max} = \phi_i^0 + \Delta\phi^{max}$$

which is equivalent to the relation $V \propto (1 - c\phi_R)^{-1}$ from the main text. Qualitatively, the parameter θ serves to convert a given percentage change in the ribosome sector, normalized to its maximum size, into an equivalent percentage change in the unregulated sector. This is then converted into a change in volume, with the normalization condition that the minimum volume is achieved when the unregulated sector is maximized ($\phi_{NC}^{max}$). Applying the condition-dependent forms of $\Delta\phi_R(\lambda)$ from the previous section to this equation, we arrive at a distinct linear ribosome-growth relationship for each translation perturbation class:

$$\frac{V_{Dwell}}{V_0}(\lambda) = \left(1 - \theta \cdot \frac{\beta_{Dwell}}{\phi_R^{max}} \cdot \lambda\right)^{-1} = (1 - \kappa_{Dwell} \cdot \lambda)^{-1} \qquad (\mathit{Varying\ Dwelling})$$

$$\frac{V_{Init}}{V_0}(\lambda) = \left(1 - \theta \cdot \frac{\Delta\phi_R^{Ref}}{\phi_R^{max}}\right)^{-1} = \frac{V_{Dwell}}{V_0}(\lambda^{Ref}) \qquad (\mathit{Varying\ Initiation})$$

$$\frac{V_{Elong}}{V_0}(\lambda) = \left(\frac{\phi_{NC}^0}{\phi_{NC}^{max}} - \theta \cdot \frac{\beta_{Elong}}{\phi_R^{max}} \cdot \lambda\right)^{-1} = (\alpha_{Elong} - \kappa_{Elong} \cdot \lambda)^{-1} \qquad (\mathit{Varying\ Elongation})$$

Examining the specific case of tRNA synthetase knockdowns – which exhibit positive growth-size scalings of the SMK type – we can see directly how this classic law emerges naturally from our integrated model:

$$V_{Dwell} = V_0 \exp(\kappa_{Dwell} \cdot \lambda) \exp\left(\frac{1}{2}(\kappa_{Dwell} \cdot \lambda)^2 + O(\lambda^3)\right)$$

In other words, our model form can be approximated as exponential in growth rate when the change in $\Delta\phi_R$ over conditions is small (e.g., $\kappa_{Dwell} \cdot \lambda$ is small). From this we also see that the deviation from an equivalent exponential fit will be second-order in the growth rate, reflecting an increased relative curvature in the negative hyperbolic model.
