## Supplementary Theory for "Essentialome-Wide Multigenerational Imaging Reveals Mechanistic Origins of Cell Growth Laws"

Here we expand on the (p)ppGpp control model, introduced in [60] and develop several derivative results for model behavior under various conditions of translation inhibition related to our experimental data in the main text. We begin by reviewing the growth theories of [53] and [52], discussing their empirical motivations, assumptions and ultimate limitations in accounting for our data. Motivated by these limitations, we reconstruct the model of ribosome-intrinsic (p)ppGpp regulation, first introduced in [60], which we show resolves many of these experimental inconsistencies. Using simple assumptions, we derive the expected dependence of ribosome concentration on growth rate, under variable translocation time and translation initiation rate. We then show that these results, along with the equivalent dependence under variable dwell time from [60], are sufficient to reconstruct the empirical behaviors originally motivating the growth theory of [53]. Finally, we show that these predictions are largely borne out by our RNA/Protein and (p)ppGpp data under translation limitation, touching on the special cases of *relA* deletion and translation initiation limitation. Overall, this model of (p)ppGpp regulation and the associated results are in agreement with our conclusion in the main text that SpoT mediates degradation of (p)ppGpp under conditions of diminished translation elongation.

**7 Conclusion** **26**

**References** 28

### 1. STRENGTHS AND LIMITATIONS OF PRIOR GROWTH THEORY

In the main text, we observe three distinct responses of the ribosome content, measured by RNA/Protein, to various perturbations of the translation machinery: 1) increased RNA/Protein under inhibition of translocation, 2) decreased RNA/Protein under reduced tRNA charging and 3) mostly invariant RNA/Protein under impaired translation initiation. We further show that these behaviors are generally accompanied by concomitant changes in (p)ppGpp, a repressor of ribosome synthesis. The first two behaviors were anticipated by the coarse-grained gene regulatory model proposed in [53], which we reproduce following the original presentation in Scott et al. 2010. We further discuss why the observed invariance of RNA/Protein under reduced translation initiation cannot be rationalized under this model, but is in fact well addressed by the model in [60].

**Fig. S1. RNA/Protein ratio vs growth rate measurements from Scott et al. 2010.** A) Measurements for WT cell, where differently colored points/lines represent different growth media under various degrees of translation inhibition by chloramphenicol (0,2,4,8 or 12$\mu M$ Cm). Circles with black outlines correspond to untreated cultures and the black line to a fit of these points. All fits used weighted least squares regression with weights scaled to RNA/Protein measurement error. B) Similar measurements for mutant strains bearing slowly elongating ribosomes grown in different growth media. Differently colored points/lines represent different strains with elongation rates of 15 (blue), 7.8 (orange) and 5 a.a./s (green) [46].

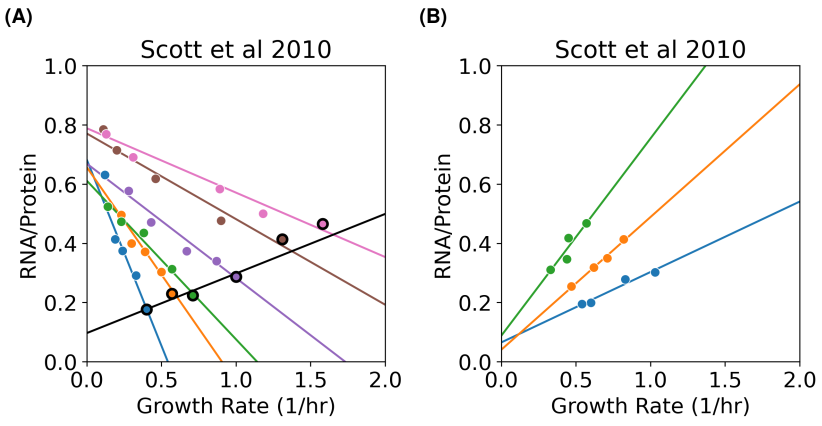

#### A. Ribosome Content Under Varying Nutrients, Translational Capacity

The growth theory introduced in [53] was primarily motivated as a means of modeling the relationship between ribosome content and the growth rate while varying both (1) nutrient quality and (2) the translation capacity. More specifically, it has long been observed [38] that ribosome content exhibits a positive, linear relationship with the growth rate, when varying the nutrient composition of the growth medium (Figure S1A). In contrast, it has also been shown that the opposite trend, that is, increasing ribosome content with decreasing growth rate, is provoked when limiting translation by either the introduction of sub-lethal quantities of the translation inhibitor chloramphenicol (Figure S1A) or through the presence of certain translation-deficient ribosome mutants (Figure S1B). Formally, we say that for exponentially growing cells in various media supporting different growth rates, the RNA/Protein ratio, $R$ is related to the growth rate $\lambda$ as:

$$R = R_0 + \kappa_t^{-1}\lambda \tag{S1.1}$$

where $R_0$ is the intercept at zero growth and $\kappa_t$ is the inverse slope. On the other hand, for cells with a variable translation capacity due to either the presence of the translation inhibitor chloramphenicol or mutant ribosomes, the RNA/Protein ratio, $R$ is related to the growth rate $\lambda$ as:

$$R = R_{max} - \kappa_n^{-1}\lambda \tag{S1.2}$$

where $R_{max}$ is the intercept at zero growth and $\kappa_n$ is the inverse slope. Interestingly, when such lines were constructed using chloramphenicol inhibition, under differing nutrient conditions (Figure S1A), it was found that $\kappa_n$ scaled positively with the growth rate in untreated media. As such, $\kappa_n$ was designated as representing the "nutritional capacity" supported by the medium. Similarly, when nutrient conditions were varied to provoke the scaling S1.1 in cells bearing mutant ribosomes with different elongation rates, $\kappa_t$ scaled positively with the elongation rate, leading to its designation as reflecting "translational capacity". In the following sections, we review the molecular rationale provided in [53] to explain each of these behaviors.

#### B. Growth-Ribosome Scaling Under Varying Nutrient Quality

We may reproduce the empirical relation of equation S1.1 from first principles assuming that ribosomes are limiting for growth and exhibit a constant translation rate, following the argument in [53]. Under these assumptions we may express the mass synthesis rate of a given proteome fraction as:

$$\frac{d}{dt}M_X = f_X \cdot \varepsilon \cdot N_R \tag{S1.3}$$

where $M_X$ is the mass of a proteome fraction $X$, $f_X$ is the fraction of ribosomes allocated to that proteome fraction, $\varepsilon$ is the elongation rate and $N_R$ is the number of ribosomes. In [53], the proteome is divided into three components R, P and Q, which will be specified later to describe the origin of the $R_{max}$ term in equation S1.2. It follows that the total protein mass $M = M_P + M_Q + M_R$ is produced according to:

$$\frac{d}{dt}M = \varepsilon \cdot N_R \tag{S1.4}$$

Specifying that the R fraction of the proteome corresponds to ribosomal or "R"-proteins as well as other translation-related proteins and that all ribosomes are engaged in translation, it follows that $M_R = m_R \cdot N_R$, where $m_R$ is the amino acid mass of the ribosome and its associated components, taken as 7336 a.a./ribosome in [53]. If we assume the system is in the exponential steady-state we have $M_X(t) = M_X(0) \cdot e^{\lambda t}$ for each proteome fraction, with the following relations for the fractions R, P, Q and the total protein mass:

$$\lambda = f_R \cdot \varepsilon / m_R \tag{S1.5}$$

$$\lambda M_P = f_R \cdot \lambda M \tag{S1.6}$$

$$\lambda M_Q = f_Q \cdot \lambda M \tag{S1.7}$$

$$\lambda M = \varepsilon \cdot M_R / m_R \tag{S1.8}$$

Focusing on equation S1.5, we see that this can be expressed in terms of the ribosomal mass fraction $\phi_R = M_R/M$ and rearranged as:

$$\phi_R = \frac{\lambda}{\varepsilon / m_R} \tag{S1.9}$$

with the conversion to RNA/Protein $R$, given by $\phi_R = \rho \cdot R$, where $\rho \approx 0.76$ $\mu g$ protein / $\mu g$ RNA [53]. We see that this is equivalent to equation S1.1, without an offset $R_0$, which results from the fact that not all ribosomes are actively engaged in translation. Relating this to $\kappa_t$ from equation S1.1, we have:

$$\kappa_t = \rho \cdot \varepsilon / m_R \tag{S1.10}$$

Based on this relation, it follows that the slope $\kappa_t^{-1}$ in equation S1.1 is inversely dependent on the elongation rate $\varepsilon$. In [53], the authors demonstrate this to be the case by gathering ribosome content vs growth curves under variable nutrient conditions, for strains possessing slowly-elongating ribosome mutants (Figure S1B). The primary assumption leading to equation S1.9 is that the elongation rate $\varepsilon$ is constant across varying nutrient conditions, irrespective of the particular regulation that achieves that condition. Equivalently, this model takes the overall ribosome concentration as being essentially rate-limiting for growth, meaning that any increase in the growth rate necessitates a linearly proportional increase in ribosome levels, as opposed to an increased elongation rate.

Empirically, $\varepsilon$ appears to be constant, under differing nutrient conditions, at growth rates from roughly 0.5 to 2 doublings per hour [10]. Since the elongation rate $\varepsilon$ has a Michaelis-Menten dependence on tRNA charge, it is somewhat surprising that $\varepsilon$ is maintained at a maximal rate over a 4-fold range in growth, despite large changes in the ribosome concentration. It has been suggested that ribosome abundance and activity is, in fact, regulated to maintain this maximum elongation rate to optimize the overall rate of growth, a notion we will revisit when discussing (p)ppGpp-dependent regulation of ribosome synthesis.

#### C. Rationale for the Maximum Ribosome Fraction $\phi_R^{max}$

Continuing with the presentation from [53], the second notable behavior the authors sought to model was the existence of a maximum ribosome fraction at $\phi_R^{max} \approx 55\%$. They suggested a three component model of the proteome, with R being the ribosome-affiliated fraction, previously discussed, and another two fractions P and Q. They took Q to be a growth-rate independent sector occupying a mass fraction:

$$\phi_Q = 1 - \phi_R^{max} \tag{S1.11}$$

Additionally, they took P to be a growth-rate variable component which is diminished as the ribosome fraction $\phi_R$ increases:

$$\phi_P = \phi_R^{max} - \phi_R \tag{S1.12}$$

Thus, in this view, $\phi_R^{max}$ is established as a consequence of a large fraction of proteins Q, that are maintained at the same level, regardless of growth rate. It appears self-evident that such a fraction must exist to carry out the activities of biosynthesis that are not directly related to protein synthesis. While other implications of this fraction are detailed in [53], the most important consequence of this result is that the R-protein concentration will be limited to the value $\phi_R^{max}$ corresponding to the value $R_{max}$ in equation S1.2.

#### D. $\kappa_n$ Originates from Allocation to Precursor Synthesis

Finally, to capture the scaling behavior of equation S1.2, it was argued in [53] that such a relation may be expected from the effects of a metabolic bottleneck on amino acid synthesis. They assumed a growth bottleneck consisting of a single rate-limiting enzyme, E. Considering the mass of the bottleneck enzyme to be $M_E$ it follows that the growth rate will be determined by the flux $J$ of the growth limiting precursor metabolized by E:

$$\lambda M = c \cdot J = c \cdot k_E \cdot M_E \tag{S1.13}$$

where $c$ scales the quantity of the product of E to the overall growth rate and $k_E$ is the kinetic rate constant of E. We can recover equation S1.2 if we additionally assume that the abundance of E scales inversely with the ribosome concentration, i.e., that E belongs to the P-sector of the proteome. It follows that the mass fraction of E is:

$$\phi_E = M_E / M = \alpha_E \cdot \phi_P \tag{S1.14}$$

where $\alpha_E$ is the fraction of P composed of E. Substituting equation S1.14 into equation S1.13 results in:

$$\lambda = c \cdot \alpha_E \cdot k_E \cdot \phi_P \tag{S1.15}$$

which, along with the existence of a maximum R-sector allocation $\phi_R^{max}$ described by equation S1.12 is equivalent to equation S1.2 with the nutritional capacity $\kappa_n$ identified as:

$$\kappa_n = c \cdot \alpha_E \cdot k_E \cdot \rho \tag{S1.16}$$

We see from this relation that the increase in $\kappa_n$ (and $\lambda$) associated with media of higher quality arises naturally from increasing values of the $c \cdot \alpha_E \cdot k_E$ term. Thus, according to this model, under conditions of reduced translation capacity, the R-fraction will expand at the expense of the P-fraction. This decrease in the P-fraction will then result in a linear decrease in the overall rate of nutrient influx, leading to the linearity in equation S1.2.

#### E. Complete Proteome Model from Scott et al. 2010

Combining the relations specified by equations S1.1, S1.2, and S1.12 we arrive at the three component growth theory posed by [53]:

$$\phi_R = \rho \cdot \lambda / \kappa_t + \phi_0 \tag{S1.17}$$

$$\phi_P = \rho \cdot \lambda / \kappa_n \tag{S1.18}$$

$$\phi_P + \phi_R = \phi_R^{max} \tag{S1.19}$$

Our molecular interpretation of this growth theory rests on the previous stated assumptions underlying the specification of the parameters $\kappa_t$, $\kappa_n$ and $\phi_R^{max}$. In particular, these relations arise naturally when decreases in either the R or P-fractions of the proteome relate *linearly* to the growth rate. Thus, we treat ribosome concentration as linear with growth to link equation S1.17

to conditions of variable nutrient quality and treat nutrient influx enzyme concentration as linear with growth to map equation S1.18 onto conditions with variable elongation rates. For instance, we cannot have that a decrease in the P-fraction under reduced translation capacity results in a non-linear reduction in growth e.g., due to the relief of product inhibition.

As discussed in [52], these conditions are satisfied under an allocation scenario in which the amino acid pool is 1) small enough to avoid end product inhibition of nutrient influx and 2) large enough to maintain a maximum ribosome elongation rate $\varepsilon$. Such an allocation would be desirable from a fitness perspective, as it results in maximal flux into protein synthesis. In this context, the invariance of $\varepsilon$ over a wide range of growth rates is taken as evidence that the cell follows such an allocation strategy, maintaining amino acid pools high enough to support this maximum elongation rate.

In [52] it was suggested that such an optimal allocation could be achieved by an increasing regulatory function $\chi_R(a)$ that scales the amount of ribosome synthesis $\chi_R$ to the steady-state amino acid level $a$. The authors suggested that repression of ribosome synthesis by (p)ppGpp could provide a reasonable implementation of such an increasing regulatory function. That is, under conditions where amino acid levels $a$ increase, the level of tRNA charging will increase and lower the overall rate of (p)ppGpp synthesis through RelA. In turn, this will de-repress ribosome synthesis, leading to a positive relationship between ribosome levels and the amino acid pool size. Thus, in this view, (p)ppGpp would mediate adjustment of the R and P fractions such that flux is balanced, the ribosome elongation rate is maintained at a near-maximum level, and growth is maximized.

In the absence of such regulation, cells would have a fixed rate of ribosome synthesis and nearly always suboptimally allocate to the R and P-fractions, resulting in translation being limited by either protein synthesis or amino acid supply. Consistent with this, it has been shown that strains which cannot produce (p)ppGpp exhibit a fixed ribosome concentration at many growth rates from 0.7 to 2 doublings per hour [42], implying that the ribosome elongation rate $\varepsilon$ must be scaling with growth rate. Thus, the validity of equation S1.9 (and therefore equation S1.1), likely depends on (p)ppGpp regulation to maintain constant $\varepsilon$ across differing nutrient conditions.

#### F. Limitations of the Growth Theory

Fundamentally, the growth theory is jointly motivated by the *a priori* goal of fitness maximization [52] and the empirical response of the proteome to perturbations to nutrient availability, translation and over-expression of inert proteins [53].

However, there remain questions about some assumptions underlying this model. In particular, the condition of linearity between growth and both the R and P-fractions is critical, as the molecular mechanisms invoked to explain equations S1.17 and S1.18 rely on this. In other words, when nutrient quality is varied, the elongation rate cannot change and when the elongation rate is varied, the rate constant of nutrient influx cannot change. It remains unclear whether this is indeed the case under the experimental conditions interrogated.

In fact, under chloramphenicol treatment, the fraction of tRNAs in the charged state increases, up to 100% [35], strongly indicative that translation has become solely rate-limiting for growth under this condition. In other words, it is likely that this bottleneck provokes an increase in the amino acid pool size such that the non-linear effect of product inhibition cannot be ignored. This complicates the previous molecular interpretation of $\kappa_n$ in equation S1.18, which relies on a fixed kinetic rate of nutrient influx enzymes. As we shall later see, our interpretation of equation S1.18 is also problematized by the observation that decreasing translation through a decreased initiation rate does not strongly impact the R-sector, despite this being the expectation under the prior growth theory. Thus, while several of the empirical claims of the growth theory of [53] and [52] remain true, the mechanism implementing $\kappa_n$ and the reason for inconsistency between different translation defects remains unclear. We will show in the next section that the explicit ribosome regulatory model of [60], further developed here, resolves these inconsistencies, while recapitulating all the empirical claims of the previous growth model.

### 2. MODEL FOR RIBOSOME-INTRINSIC (P)PPGPP REGULATION

The (p)ppGpp regulatory model introduced in [60] addresses many of the limitations of the prior growth model and, as we will show, provides excellent predictive power over a large range of perturbations to translation. The key assumption of this model is that translocating ribosomes provoke a previously unappreciated (p)ppGpp degradation activity, which based on our results we assign to SpoT. In the following sections, we describe this regulatory model and produce the resultant predictions for ribosome and (p)ppGpp scaling with growth rate under different conditions of impaired translation.

#### A. (p)ppGpp Regulation in WT *E. coli*

We begin by reproducing the model, reviewing its assumptions, and re-deriving the resulting relationship between elongation rate and (p)ppGpp levels. In [60] the authors simplify the kinetics of ribosome elongation into two steps, a dwelling step in which ribosomes await successful capture of a charged tRNA in the A-site and a translocating step in which an amino acid is transferred to the elongating peptide chain, the tRNA in the A-site is transferred to the P-site and the ribosome moves to the succeeding codon.

Based on biochemical and structural evidence, it is widely accepted that the (p)ppGpp synthase RelA is stimulated by interacting with uncharged tRNAs in the A-site, a state that should be proportional to the number of dwelling ribosomes. Motivated by the empirical relationship between elongation rate and (p)ppGpp levels, they postulate a secondary activity of the ribosome in degrading (p)ppGpp, in proportion to the number of ribosomes in the translocating state. They further speculated that this type of activity could be mediated by the (p)ppGpp hydrolase/synthase SpoT.

To model this effect, the authors specify that dwelling ribosomes at a concentration $R_D$ produce (p)ppGpp at some rate $\alpha$ and that charged tRNA bound or "translocating" ribosomes $R_T$ degrade (p)ppGpp at some rate $\beta$. This results in the following steady-state value of (p)ppGpp, $g(R_D, R_T)$:

$$g(R_D, R_T) = c\frac{R_D}{R_T} \tag{S2.1}$$

where $c = \alpha/\beta$. Equivalently we can state this in terms of the fraction of ribosomes in the dwelling state $f_D$ and the fraction of ribosomes in the translocating state $f_T$:

$$g(f_D, f_T) = c\frac{f_D}{f_T}$$

illustrating that g is sensitive to the kinetics of translation, rather than the total concentration of ribosomes. If we take $\tau_T$ and $\tau_D$ as the dwell times of the translocating and dwelling states, respectively, from detail balance we have:

$$R_D\tau_D^{-1} = R_T\tau_T^{-1} \tag{S2.2}$$

Substituting this into equation S2.1, we have:

$$g(\tau_D, \tau_T) = c\frac{\tau_D}{\tau_T} \tag{S2.3}$$

The kinetic steps of the ribosome cycle can then be related to the elongation rate $\varepsilon$ as:

$$\varepsilon^{-1} = \tau_T + \tau_D \tag{S2.4}$$

In [60] it was assumed that under conditions where tRNA charge is limiting, $\tau_D$ varied while $\tau_T$ remained constant. Also, it was assumed that $\tau_D \longrightarrow \tau_D^0$ as $\varepsilon \longrightarrow \varepsilon_{max}$, where $\tau_D^0 \ll \tau_T$. This led to $\varepsilon_{max} = \tau_T^{-1}$, allowing equation S2.3 to be re-written as:

$$g_D(\varepsilon) = c\frac{\tau_D}{\tau_T} = c(\varepsilon^{-1} - \varepsilon_{max}^{-1})\varepsilon_{max} = c(\frac{\varepsilon_{max}}{\varepsilon} - 1) \tag{S2.5}$$

Experimental evidence from [60] further demonstrated that, empirically, (p)ppGpp levels indeed exhibit an inverse dependence on the elongation rate under different growth rates and during diauxic shifts from glucose to glycerol. Critically, under the ribosome-intrinsic model of (p)ppGpp regulation specified by equation S2.1, this relationship is a consequence of variable ribosome dwelling time. As such, we may expect a very different relationship to arise when, instead, we vary the translocation rate as in our EF-G and ribosomal protein knockdowns.

To reformulate this (p)ppGpp regulatory model to include the effects of reduced translocation time, we once again begin with equation S2.3. We add the constraint that the amino acid supply is not limiting, and the overall elongation rate is totally limited by the translocation time, that is $\varepsilon^{-1} = \tau_T$. These are reasonable assumptions for the medium to fast growth rate regime, and we will assume $\lambda > 0.75h^{-1}$ in all cases where we limit growth by inhibiting translocation. These assumptions lead to the following (p)ppGpp vs elongation relationship:

$$g_T(\varepsilon) = c\frac{\tau_D}{\tau_T} = c\tau_D\varepsilon \tag{S2.6}$$

We later show that both elongation rate $\varepsilon$ to (p)ppGpp concentration functions, $g_D(\varepsilon)$ and $g_T(\varepsilon)$, combined with plausible ribosome regulatory functions $R(g)$, lead to distinct ribosome concentration to growth rate relationships.

#### B. Special Case: Chloramphenicol Treatment

Interestingly, an additional source of evidence for the empirical relation specified by equation S2.5 from [60] was the effect of chloramphenicol (Cm) treatment on elongation rate and (p)ppGpp levels. Specifically, it was found that reduced (p)ppGpp concentration under Cm treatment was accompanied by an increase in the measured elongation rate, following equation S2.5.

However, based on biochemical evidence [58], Cm is well-known to inhibit peptidyl transfer but not the initial binding of charged tRNAs to the A-site. As such, it is unclear that such behavior would be well-modeled by equation S2.5, which assumes an increase in the ribosome dwell time, not an increase in the translocation time. We favor reconciling these observations by invoking the model of Cm inhibition specified in [10], which was used to rationalize similar elongation rate measurements under Cm treatment.

In both [60] and [10], elongation rates are measured using the LacZ induction method. Briefly, this assay measures elongation rate by quantifying the time it takes the ribosome to translate a *lacZ* transcript. This may be measured by inducing *lacZ* transcription at time 0 and subsequently measuring LacZ expression by beta-galactosidase assay over time. The timepoint at which LacZ expression begins to rise provides a measure of elongation time across the transcript, which is then normalized by transcript length to produce the elongation rate.

During Cm treatment, when a translating ribosome is bound by a single Cm molecule, it will remain bound for some time with a complex lifetime of ~12 mins [19]. [10] further argues that, in 99% of cases, this ribosome will stall and initiate RNase degradation of the transcript. Any ribosomes downstream of this stalled complex will finish translation without any impedance, while upstream ribosomes will also stall and eventually recycled after the transcript is degraded. Because the LacZ induction assay only measures complete LacZ molecules, the apparent elongation rate will only reflect the activity of unimpeded ribosomes and appear high. But, in actuality, the average elongation rate including stalled ribosomes will be even lower than that in untreated cells.

Therefore, if we consider Cm treatment to decrease the average elongation rate, it no longer follows that it obeys the relation S2.5. Given that the mechanism of Cm inhibition is to increase $\tau_T$ through inhibited peptidyltransfer, we should instead expect this treatment to obey equation S2.6. That is, as the Cm concentration is increased, $\varepsilon$ should decrease resulting in diminished (p)ppGpp. Thus, this formulation both recapitulates the known drop in (p)ppGpp under Cm treatment, reconciles it with the molecular model specified by equation S2.1 and remains consistent with our molecular view of Cm inhibition.

### 3. RIBOSOME-GROWTH SCALING

Here, we determine the consequences for the proposed ribosome regulatory model in terms of the overall scaling between ribosome concentration and growth. We show that these predictions are consistent with observations under various perturbations to translation from the main text including reduced tRNA charging, reduced elongation/translocation and reduced translation initiation.

#### A. Scaling Under Nutrient Limitation

In [60], the authors incorporated feedback between (p)ppGpp and ribosome synthesis/activity into their model, yielding a linear relationship between ribosome content and growth rate under nutrient limitation. We reproduce this derivation in abbreviated form here. [60] first specified the following relationship including the effects of (p)ppGpp, ribosome synthesis, and ribosome sequestration by hibernation factors on the active concentration of ribosomes able to engage in translation at an elongation rate $\varepsilon(g)$, potentially also dependent on (p)ppGpp concentration:

$$\lambda(g) = \varepsilon(g)\Big(R(g) - H(g)\Big) = \varepsilon(g)\left(\frac{a}{g} - bg\right) \tag{S3.1}$$

where $R(g)$ and $H(g)$ were functions capturing the (p)ppGpp-dependence of the total and inactive ribosome concentrations, respectively. Substitution of the $R(g)$ term with equation S2.5 resulted in the following equation, in the limit of slow elongation:

$$a = c \cdot R_0 \cdot (h - 1) \tag{S3.2}$$

with $h = \varepsilon_{max}/\varepsilon_0$. It was further argued that, as $\lambda$ approaches 0, $\varepsilon_0 > 0$ and so it must be that $H(g) = R(g)$ in this limit (i.e., that all ribosomes are hibernating in this limit). Combining this assumption with substitution of the $H(g)$ term with equation S2.5 yields:

$$b = \frac{R_0}{c \cdot (h - 1)} \tag{S3.3}$$

Substituting $R(g) = a/g$, equation S3.2, and equation S3.3 into equation S3.1, we obtain:

$$\lambda = \varepsilon_{max} \frac{R^2 - R_0^2}{R + R_0 \cdot (h - 1)} \tag{S3.4}$$

Finally, if we assume that $\varepsilon_0 : \varepsilon max = 1 : 2$, which is supported by the measurements in [60] then $h = 2$ resulting in:

$$R = R_0 + \varepsilon_{max}^{-1} \lambda \tag{S3.5}$$

where the slope of the $R$ vs $\lambda$ is positive and corresponds to $\varepsilon_{max}^{-1}$. This is functionally equivalent to the classical linear dependence between the ribosome concentration and the growth rate observed across different nutrient conditions. Comparing this relation to equation S1.1, we see that it takes the same form with $\varepsilon_{max}$ taking the place of $\kappa_t$. That is, under this model, the translation capacity merely reflects the maximum elongation rate supported by the cell's ribosomes. Indeed, values for $\kappa_t$ were collected under conditions of variable elongation rate (Figure S1B), supporting the identification of $\kappa_t$ with $\varepsilon_{max}$.

### B. Scaling Under Slow Translocation

We were interested in modeling how our predicted (p)ppGpp vs elongation rate relationship under slow translocation, $g_T(\varepsilon, \tau_D)$, would constrain the scaling between growth and ribosome content. Like [60] we required a regulatory function relating (p)ppGpp to ribosome content. It is convenient in this case to model (p)ppGpp as a simple monovalent repressor of ribosome synthesis:

$$R(g) = \frac{k}{k + g}(R_{max} - R_0) + R_0 \tag{S3.6}$$

where $R_{max}$ is the maximum ribosome concentration and $R_0$ is the basal ribosome concentration at high (p)ppGpp levels. Examining the (p)ppGpp vs RNA/Protein data from [60] we see that this function provides a fit of similar quality to that of the $a/g$ regulatory function suggested previously. Further, for similar data originally from [1, 2, 3, 4, 5, 6, 10, 14, 16, 20, 23, 27, 29, 30, 34, 47, 48, 51, 53, 54, 55, 60, 61, 62] and collected in [8], our repressor model provides an even better fit than the simple $a/g$ form.

Finally, we assume the growth model from equation S3.1, with the additional assumption that $H(g) = R_0$. In other words, we assume that (1) the concentration of inactive ribosomes $H(g)$ is constant and (2) that it is roughly equivalent to the basal expression of ribosomes at high (p)ppGpp levels $R_0$. (1) is true as $H(g) = 0.03$, having no growth rate dependence at moderate to fast growth rates, where the doubling time $\lambda > 0.5h^{-1}$ [10]. Given this, to satisfy (2), we must have that $R_0 = 0.03$. A fit to the $R$ vs $\lambda$ curve in [10] (Figure S3) shows that, for the linear trend over the $\lambda > 0.75h^{-1}$ portion of the domain, the corresponding intercept $R_0 = 0.03$. Since we are specifically modeling the medium to fast growth regime, we may take this as the value of $R_0$, satisfying (2) and justifying our assumption $H(g) = R_0$.

As an additional validation of this claim, we examined the $R_0$ determined from our previously fitted repressor functions (Figure S2), which took values of 0.04 and 0.07. Thus, in either case, the value was reasonably close to 0.03, further justifying our substitution of $H(g) = R_0$. As a result we have:

$$\lambda(g) = \varepsilon(g)\Big(R(g) - R_0\Big) \tag{S3.7}$$

Substituting equation S2.6 with this relationship yields:

$$g_T = \frac{c\tau_D \lambda}{R - R_0} \tag{S3.8}$$

**Fig. S2.** **RNA/Protein vs relative (p)ppGpp measurements.** A) Blue line indicates the model fit (with fixed $R_{max}$ = 0.793) for our repressor model, orange line indicates the fit for the $a/g$ model from Wu et al. 2022. The inset box indicates the fitted model parameters for our repressor model. B) RNA/Protein vs relative (p)ppGpp measurements, aggregated by growth rate windows of 0.1, derived from data collected in Chure and Cremer 2023. Models the same as in Figure S2A, fitted on these data.

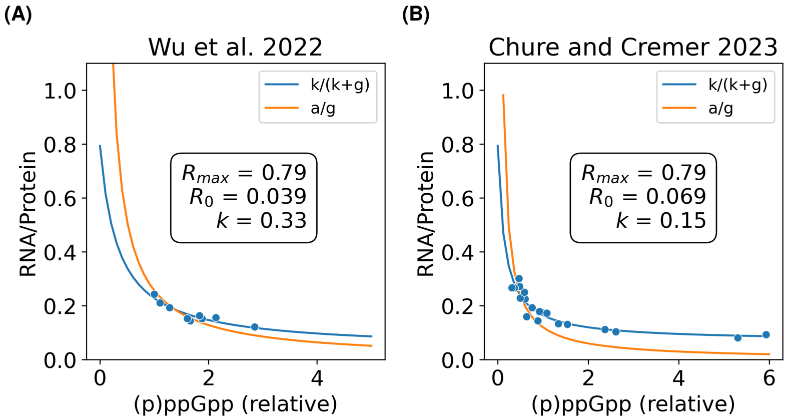

**Fig. S3.** **Inactive Ribosome Measurements from Dai et al. 2016.** A) RNA/Protein ratio vs growth rate measurements from Dai et al. 2016. Black line is a linear fit for points with a growth rate over $0.75h^{-1}$, $R_0$ is the intercept for this fit. B) RNA/Protein contributed by inactive ribosomes vs growth rate measurements from Dai et al. 2016. The dotted line indicates the mean RNA/Protein (inactive) value $H_0$ for points with a growth rate over $0.75h^{-1}$.

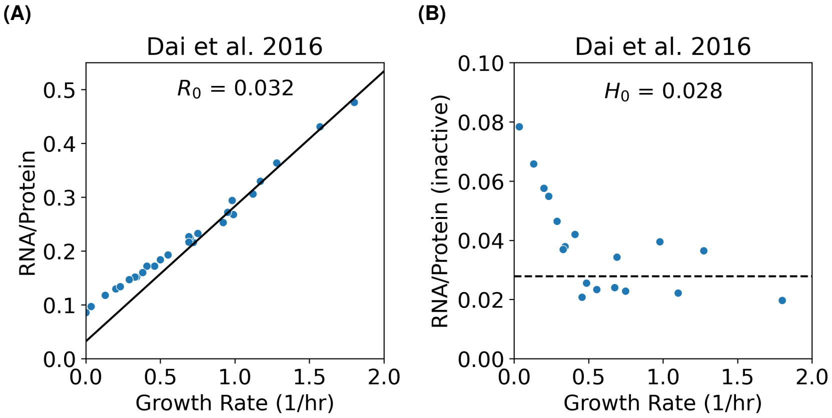

Further, we can rearrange equation S3.6 into the following form and then substitute $g_T$ to produce a ribosome vs growth relationship:

$$R = R_{max} + \frac{(R_0 - R)}{k} g_T = R_{max} + \left(\frac{c\lambda\tau_D}{k}\right)\left(\frac{R_0 - R}{R - R_0}\right)$$

Simplifying this yields a final, relatively simple form for ribosome-growth scaling under reduced translocation:

$$R(\lambda) = R_{max} - \frac{c\tau_D}{k}\lambda \tag{S3.9}$$

where $\frac{c\tau_D}{k}$ is positive. So this predicts that the relationship between ribosome content and growth, under translocation limitation, will be linear and negative. In fact, this equation takes the same form as equation S1.2, which served to model ribosome-growth scaling under sub-lethal chloramphenicol treatment or elongation rate-slowing mutations. Recall that the inverse slope in this relation, $\kappa_n$ scales with the unperturbed growth rate or "nutritional capacity" of the medium. Based on the argument in "Special Case: Chloramphenicol Treatment", it is likely that sub-lethal chloramphenicol treatment specifically slows ribosome elongation via increased $\tau_T$. Similarly, slow ribosome mutants also exhibit higher accuracy [46], indicating an extended proofreading period to sample tRNA-carrying ternary complexes (i.e., increased $\tau_T$). Therefore, all of these data could be fully explained by the proposed (p)ppGpp degradation activity of the ribosome, resulting in the following relation between $\kappa_n$ and $\tau_T$:

$$\kappa_n = \frac{k}{c\tau_D} \tag{S3.10}$$

Thus, revisiting [53] in the context of our model, we may produce an equivalence between a phenomenological parameter $\kappa_n$ and a kinetic rate parameter $\tau_D^{-1}$. Re-examining the translation inhibition curves from [53], reproduced in Figure S1A, we can now view the increase in steepness $\kappa_n^{-1}$ at low $\lambda$ as a consequence of the larger basal (p)ppGpp pool being produced under conditions of low $\lambda$ and high $\tau_D$. As a result, the same marginal increase in $\tau_T$ leads to a much larger absolute change in (p)ppGpp concentration and, as a result, a larger increase in ribosome concentration $R$. We explore the consequences of this result for the empirical determination of $R_{max}$ in section F.

#### C. Complete Proteome Model

As reviewed in section 1, the phenomenological relations specified by equations S1.1 and S1.2 may be used to constrain a complete model of the proteome (equations S1.17, S1.18, and S1.19) in terms of ribosome-affiliated "R-class" proteins, "P-class" proteins which share a proteome fraction with the R-class, and Q-class proteins which remain constant under the conditions investigated. Similarly, we may combine equations S3.5, S3.9, and the proteome constraint specified by equation S1.19 to produce a similar proteome model. We first reorganize equation S3.9 into the following form, by again specifying $\rho = \phi_R / R$:

$$\phi_R = \phi_R^{max} - \rho \frac{c\tau_D}{k}\lambda \tag{S3.11}$$

combining this with equation S1.12 gives:

$$\phi_P = \rho \frac{c\tau_D}{k}\lambda \tag{S3.12}$$

Similarly, equation S3.5 can be rearranged as:

$$\phi_R = \rho \varepsilon_{max}^{-1}\lambda + \phi_0 \tag{S3.13}$$

where $\phi_0 = R_0/\rho$. Equations S3.12, S3.13 and S1.19 form a set of constraints of similar form to equations S1.17, S1.18 and S1.19, with $\frac{k}{c\tau_D}$ taking the place of the nutrient capacity $\kappa_n$ and $\varepsilon_{max}$ taking the place of the translation capacity $\kappa_t$. The solution to the above system of equations is also of the same form:

$$\lambda\left(\varepsilon_{max}, \tau_D, \phi_R^{max}\right) = \frac{\phi_R^{max} - \phi_0}{\rho} \cdot \frac{\kappa_t \kappa_n}{\kappa_t + \kappa_n} = \frac{\phi_R^{max} - \phi_0}{\rho} \cdot \frac{\varepsilon_{max} k}{\varepsilon_{max} c\tau_D + k} \tag{S3.14}$$

$$\phi_R\left(\varepsilon_{max}, \tau_D, \phi_R^{max}\right) = \left(\phi_R^{max} - \phi_0\right) \cdot \frac{\kappa_n}{\kappa_t + \kappa_n} + \phi_0 = \left(\phi_R^{max} - \phi_0\right) \cdot \frac{k}{\varepsilon_{max} c\tau_D + k} \tag{S3.15}$$

$$\phi_P\left(\varepsilon_{max}, \tau_D, \phi_R^{max}\right) = \left(\phi_R^{max} - \phi_0\right) \cdot \frac{\kappa_t}{\kappa_t + \kappa_n} = \left(\phi_R^{max} - \phi_0\right) \cdot \frac{\varepsilon_{max} c \tau_D}{\varepsilon_{max} c \tau_D + k} \tag{S3.16}$$

substitution of equation S3.14 into equation S3.15, holding $\tau D$ constant results in equation S3.13 and substitution of equation S3.14 into equation S3.15, holding $\varepsilon_{max}$ and $\phi_R^{max}$ constant, results in equation S3.12. Other results in terms of the parameters $\kappa_t$ and $\kappa_n$ found in [53] can also be reparameterized in terms of $\varepsilon_{max}$ and $\tau_D$. For instance, we may also specify equation S3.13 such that the growth rate $\lambda$ takes units of doubling time:

$$\phi_R = \varepsilon_{max}^{-1} m_R \lambda + \phi_0 \tag{S3.17}$$

where $m_R$ is the amino acid mass of the ribosome and its associated components, taken as 7336 amino acids/ribosome in [53]. Following a similar approach, we can also express equation S3.11 with $\lambda$ in doubling units:

$$\phi_R = \phi_R^{max} - \frac{c\tau_D}{k} m_R \lambda \tag{S3.18}$$

Critically, the features of this model reproduce several of the RNA/Protein trends we observe in the main text. Under reduced translocation, due to FusA/EF-G knockdown, we find that the RNA/Protein ratio increases (Figure 5D), as it does in the model under decreased $\varepsilon_{max}$ with fixed $\tau_D$ (equation S3.9). Further, under reduced tRNA charging in a PheT knockdown, we observed a lower RNA/Protein ratio, again consistent with model behavior (equation S3.13). In the main text, we also examine the relationship of ribosome content with growth rate under reduced translation initiation, a distinct case for which we develop model predictions in the following section.

### D. Effect of Reduced Initiation

In our experiments, we observed that both length (Figure 5C) and the RNA/Protein ratio (Figure 5D) were largely independent of growth rate, when we varied the initiation rate. We can see simply from equation S2.3 that, since reduction of initiation does not change ribosome elongation kinetics, this perturbation should not change the (p)ppGpp level or the total ribosome content $R$. If we take $C_{inf}$ to be the concentration of some limiting initiation factor we can modify equation S3.7 as:

$$\lambda = \varepsilon(R - R_0(C_{inf})) \tag{S3.19}$$

where we assume that $R_0$ is now a monotonic, decreasing function of $C_{inf}$. So, consistent with our data, $\lambda$ will decrease with decreasing $C_{inf}$, without any change in $R$, by increasing the fraction of inactive ribosomes $R_0$. Thus, $R$ will simply always take its reference value in a given set of growth conditions $R_{ref}$:

$$R = R(\lambda = \lambda_{ref}) = R_{ref} \tag{S3.20}$$

To compare equations S3.9, S3.13 and S3.20, we baselined all of these functions to a common reference state $R_{ref}$ based on equation S3.17 at $\lambda = 2$:

$$R_{ref} = R(\lambda = 2) = \rho^{-1} \varepsilon_{max}^{-1} m_R \lambda + R_0 = 0.3$$

with $R_0 = 0.03$, $\varepsilon_{max}$ = 72000 a.a./hr, $m_R$ = 7336 a.a./ribosome and $\rho$ = 0.76, with choices for $\rho$ and $m_R$ based on [53]. Additionally, taking $R_{max}$ = 0.793 to specify equation S3.18, also based on [53], we produced the following trendlines for each of the models:

We note that, while our RNA/Protein measurements (Figure 5D) were fully consistent with invariant $R$ under reduced translation initiation, we did however note a roughly 50% reduction in (p)ppGpp under the same conditions (Figure 6B), where $g$ should be invariant. Reconciling these observations, we also observed that this does not occur in a Δ*relA* strain, indicating that this reduction is RelA-dependent. As mentioned in the Extended Discussion, we speculate that this effect occurs due to an un-modeled decrease in $\tau_D$, resulting from an increase in ternary complex availability under reduced ribosome competition. This is plausible as these complexes are considered to be limiting for translation [10, 26] and any reduction in availability will alter elongation kinetics.

#### *D.1. Comparison to Previous Initiation Knockdown Data*

This line of thinking also illuminates the moderate disagreement of these observations with a handful of previous works that observed increases in ribosome levels under inhibition of translation initiation by depletion IF-2 or IF-3, encoded by *infB* and *infC* respectively [9, 39, 53]. These previous results would suggest that the reduction in (p)ppGpp under reduced translation

**Fig. S4. Model Predictions Under Various Translation Limitations.** A) Predicted relationship between the RNA/Protein ratio and growth rate, under variable translocation time (blue), dwell time (orange) and initiation factor concentration. B) Predicted relationship between the relative (p)ppGpp concentration and growth rate, under variable translocation time (blue), dwell time (orange) and initiation factor concentration. All values based on a reference state of $R(\lambda = 2) = 0.3$

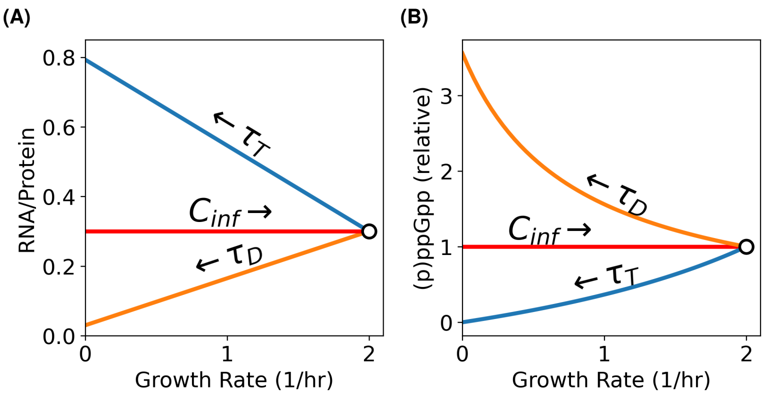

initiation is greater than that which we have observed. Critically, our experiments were all conducted in rich media at rapid growth rates, where all amino acids are supplied by the extracellular environment. As such, tRNA charge is expected to be relatively high and RelA activity correspondingly low. Consistently, comparing basal (p)ppGpp levels in WT and Δ*relA* strains (Figure 6B), we observe that RelA supplies around half of the basal (p)ppGpp in the cell.

In comparison, previous experiments in which initiation factor knockdown has been considered have been conducted in relatively poor media, with basal RNA/protein ratios varying between 0.37-0.38 in comparison to our basal level of 0.58. In these poorer conditions, the impact of modulating RelA activity will be far greater on the absolute change in (p)ppGpp. Therefore, we would expect that the effect of reduced translation initiation on (p)ppGpp and ribosome levels would be magnified - consistent with previous observations of increased RNA/protein up to 0.62 after initiation factor depletion under these conditions.

Importantly, the evidence that ribosome levels are mostly unaffected by significant reductions in the translation initiation rate is problematic for the generality of the amino-acid pool centric model of ribosome regulation provided in [52, 53]. Under such a model, any reduction in the translation rate per ribosome will engender an equivalent regulatory response, which is clearly not the case under these conditions. Moreover, any model which only considers amino acid pool size [52, 53] or tRNA charge ([8]) as the primary variable relating translation status to (p)ppGpp/ribosome levels will fail to account for this behavior without additional considerations. Thus, in light of these data, such models will most likely need to be revised to include other variables e.g., the translocation rate.

### E. Effect of Ribosome Stalk Knockdown

As noted in the main figures, the GTPase-stimulating L7/L12 ribosome stalk RplL exhibits exceptionally invariant size and ribosome levels compared to most ribosomal proteins (Figures 5C,D). Based on these data and our other results, it appears that RplL knockdowns exhibit invariant (p)ppGpp levels. Through its GTPase-stimulating activity, RplL promotes translocation (though EF-G/FusA), charged tRNA capture (through EF-Tu) and initiation (through IF-2/InfB) [18, 21, 32]; all three of the major activities we identify as impacting (p)ppGpp homeostasis. All these rates have been shown to be highly defective in ribosomes lacking RplL *in vitro*, from either decreasing the effective availability of these GTPases or by stimulating GTPase activity [21, 32]. It somewhat surprising that the net effect of ribosome stalk loss, which will impact all these processes, exhibits little to no impact on (p)ppGpp-dependent regulation.

Previously, when we considered the effects of knocking down proteins involved in initiation, elongation and tRNA charging, we could model the consequences as smoothly varying defects in the kinetic rates of their associated steps in the translation cycle. Unlike these factors which are catalytic for singular steps in translation, the ribosome stalk is associated with ribosomes for long timescales [11], and will persistently impact the kinetic behavior of a specific fraction of the ribosome population. Explicitly modeling this behavior using our current approach is not possible because the heterogeneity in the ribosome population makes it impossible to proceed without an explicit account of queuing.

Nonetheless, our previous theoretical results suggest that a partial account of these defects is straightforward. As we have seen previously, defects in the initiation of new rounds of translation are not predicted to greatly affect (p)ppGpp levels. Ribosomes without ribosome stalks are known to initiate at a 40-fold diminished rate in vitro [21] and so the first-order effect of stalk depletion is to render that fraction of the ribosome population essentially incompetent for initiation. Since ribosomes must engage in translation to exert effects on (p)ppGpp levels under our model, initiation defects will effectively be epistatic to this class of defect. As such, we may expect ribosome stalk depletions to primarily mirror initiation defects with little change in (p)ppGpp-dependent regulation, as observed.

We note, however, that this does not account for the increase in cell length observed under ribosome stalk knockdown in a Δ*relA* strain (Extended Data Figure 6H), which according to our previous argument suggests an impact on (p)ppGpp from reduced translocation. While this is plausible given RplL function in translocation, a complete account of this multifaceted knockdown requires more information about its impact on each kinetic rate in translation and a full model for ribosome queueing.

#### F. Predicted Linearity of R and $\kappa_n$ and Consequences for $R_{max}$

A major consequence of the proteome model from [53] was the specification of a unique $\phi_R^{max}$ in equation S1.11 corresponding to an $R_{max}$ of around 0.72. Recall that this was previously argued to arise from a fixed entitlement of the proteome to a so-called Q sector, which was composed of proteins that are expressed at a fixed concentration under all growth conditions. Examining the data from [53], we see that while $R_{max}$ values are somewhat narrowly distributed across different growth media, they do in fact vary, ranging from 0.6-0.78 (Figure S1A). The origin of this variability has not been explained thus far.

In our model, which does not take proteome constraints into account, $R_{max}$ corresponds to the maximum rate of ribosome synthesis. This is reached when the repressor (p)ppGpp is zero and all ribosomal operons are completely de-repressed. While we previously took $R_{max}$ to be fixed, in reality the relative copy number of ribosomal operons will change at different growth rates [15], which will impact $R_{max}$.

Introducing this dependence on growth $R_{max}(\lambda)$ poses a difficulty since the exact relationship between growth, the effective ploidy of ribosomal operons and the resultant dependence of $R_{max}$ is complex and likely depends on (p)ppGpp regulation [15]. Nonetheless, the relationship between ribosome content and growth under translation inhibition is remarkably linear over the range of growth rates previously examined (Figure S1A). Given this, we may reasonably approximate $R(\lambda)$ under elongation inhibition by linearizing equation S1.2, at the growth rate under conditions of no elongation inhibition $\lambda^o$, where according to equation S3.5 $R$ is:

$$R^o = R_0 + \varepsilon_{max}^{-1}\lambda^o \tag{S3.21}$$

If we assume that $R_{max}$ changes very little near this point (i.e., $\frac{dR_{max}}{d\lambda} = 0$), linearizing S1.2 at $\lambda^o$ results in:

$$R(\lambda) = R_{max}(\lambda^o) - \kappa_n^{-1}\lambda \tag{S3.22}$$

So, in this view the $R_{max}$ is some function of the growth rate under conditions without translocation inhibitors. Note that we are neglecting the impact of these inhibitors on $R_{max}$, despite those inhibitors effect on (p)ppGpp homeostasis and therefore replication [15], since we are only modeling the local linear behavior of translation inhibition near $\lambda^o$. If we take the value of this function at $R^o$ and $\lambda^o$, and rearrange, we can solve for this $R_{max}$:

$$R_{max}(\lambda^o) = R^o + \kappa_n^{-1}\lambda^o \tag{S3.23}$$

In other words, if we know the translation inhibition slope in each medium, we can determine $R_{max}(\lambda^o)$ using this information, along with $R^o$, and $\lambda^o$, which are given by equation S1.1. Recall that the negative inverse of this slope $\kappa_n$ is referred to as the nutritional capacity of the medium, as it is known to increase in media which supports increasing growth rates. According to the growth model presented in [53], a form for $\kappa_n$ can be obtained by rearranging S3.14, leading to:

$$\lambda^o = \lambda_c \frac{\kappa_n}{\kappa_t + \kappa_n} \quad \text{(S3.24)}$$

where $\lambda_c = \kappa_t(R_{max} - R_0)$ represents the maximum possible growth rate under the proteome partitioning model. The resulting relationship between the growth rate in media with different nutrient composition and $\kappa_n$ is shown in Figure S5A. We can then use equation S3.23 to determine $R_{max}(\lambda^o)$ and plot the resulting curves on the data from Figure S1A. Examining these curves in Figure S5B, we see that even estimated using this approach - which allows for $R_{max}$ to vary, the estimated $R_{max}(\lambda^o)$ values do not populate the empirical range from 0.6-0.78.

**Fig. S5. Relationship Between Nutritional Capacity and Growth.**

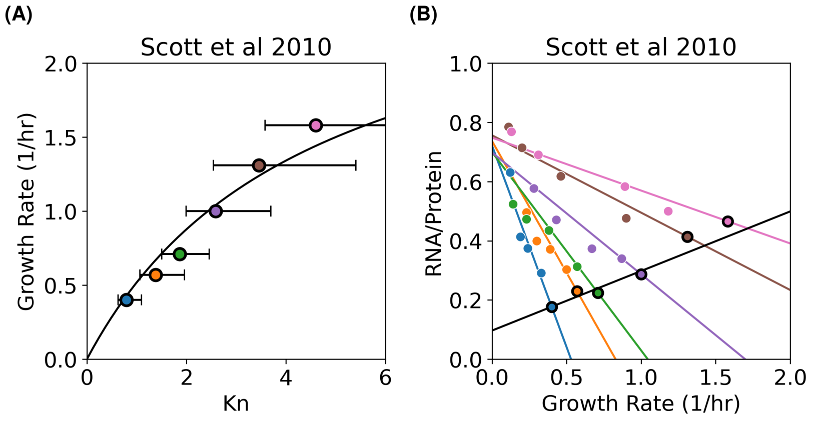

We noted that, while the proteome model predicted a Michaelis-Menten relationship between $\kappa_n$ and $\lambda^o$, the scaling in reality appears linear, through the exact dependence is difficult to determine due to uncertainty in fitting the linear $\kappa_n$ term. We asked whether our model, which does not take proteome constraints into account, could reproduce this linear behavior.

Recall that, in our model, $\kappa_n$ is linearly dependent on the dwell time $\tau_D$ according to S3.10. We can specify $\tau_D$ according to the effective concentration of ternary complexes (aa-tRNA-EF-Tu-GTP complexes), following the argument in [10]:

$$\tau_D = \frac{1}{k_{on}[TC_{eff}]} \quad \text{(S3.25)}$$

where $[TC_{eff}]$ is the effective concentration of all ternary complexes in the cell and $k_{on}$ is the forward rate constant of TC-ribosome binding. In [10] it is further verified that there is a direct proportionality between $[TC_{eff}]$ and the RNA/Protein ratio, $R$, of the form $[TC_{eff}] = C \times R$. Combining this with the previous equation results in:

$$\tau_D = \frac{1}{k_{on}C * R} \quad \text{(S3.26)}$$

Based on equation S3.10, we can substitute and rearrange this in the following form:

$$\kappa_n = \left(\frac{k * k_{on}C}{c}\right) R \quad \text{(S3.27)}$$

Thus, our model for (p)ppGpp regulation, together with the TC vs R relationship from [10], predicts that ribosome levels in media without translocation inhibitors $R^o$ should be linear with $\kappa_n$. It follows from S3.5 that $\kappa_n$ will also be linear with the growth rate in the same conditions $\lambda^o$ - consistent with Figure S5A.

Examining $\kappa_n$ vs $R^o$ values from [53], we see that the relationship is indeed linear (Figure S6A). That is, there is a direct proportionality between the basal R-sector allocation and the empirical "nutrient quality" of a given medium. Similarly to Figure S5B, we can fit this relationship to predict $\kappa_n$, determine $R_{max}(\lambda^o)$ and plot the resulting curves according to equation S3.22 (Figure S6B). We find that this approach reproduces more reasonably the spread of $R_{max}$ values observed empirically, and does especially well predicting the slope of points with lower degrees of translation inhibition.

**Fig. S6. Linear Relationship Between Nutritional Capacity and Ribosome Content.** A) Nutrient capacity ($\kappa_n$) vs RNA/Protein ratio for uninhibited points in Figure S1A. Line derived from a linear fit of these points. Error bars correspond to a 95th percentile confidence interval of the fitted $\kappa_n$, same as in Figure S5A. B) Same data as in S1A, with lines corresponding to S3.22 setting $\kappa_n(R^o)$ according to the linear fit in panel A and $R_{max}$ according to S3.23.

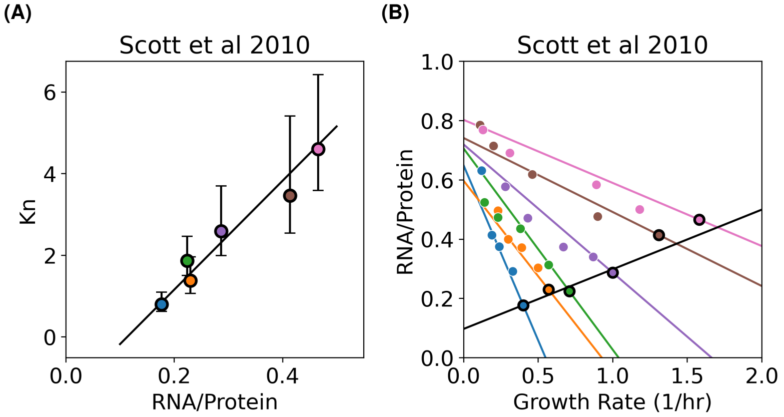

As previously mentioned we neglect the effects of translation inhibition on $R_{max}$. However, it is worth noting that low (p)ppGpp levels are associated with increases in the ori/ter ratio [15]. Since ribosomal operons tend to cluster near the origin, the expected outcome of inhibiting translocation, and thus lowering (p)ppGpp, would be an increase in $R_{max}$ - which is qualitatively consistent with the upward curvature in ribosome vs growth data at high levels of translation inhibition (Figure S5B). Additional efforts to model this non-linear impact on the (p)ppGpp to ribosome regulatory function would likely produce even greater consistency with the empirical data.

### 4. SIZE-GROWTH SCALING

While numerous studies have sought a direct regulatory interaction by which (p)ppGpp modulates cell size, none have conclusively demonstrated such a mechanism. Given the extensive proteome alterations resulting from ppGpp-mediated repression of ribosome synthesis, it is equally likely that (p)ppGpp impacts division indirectly. Proteome allocation models provide a compelling means to evaluate whether indirect regulatory effects—such as reallocating the protein synthesis machinery toward regulated components at the expense of constitutively expressed ones—can adequately explain observed size variations. In this section, we develop a simple proteome model that captures these indirect effects on division protein expression and show that this model can quantitatively capture all of our size and ribosome abundance data.

#### A. Proteome Model for (p)ppGpp Regulation

To examine the hypothesis that (p)ppGpp action to repress ribosomes is sufficient to explain our size scaling behaviors, we partition the proteome into two distinct fractions: a (p)ppGpp-repressed proteome fraction $\phi_P$ and an unregulated fraction $\phi_{NP}$. We will further assume that the proteome fraction made up of ribosomal proteins $\phi_R$ is a subset of $\phi_P$ while the proteome fraction comprised of the set of limiting division proteins $\phi_D$ is a subset of $\phi_{NP}$.

We will separate each such sector into two components:

$$\phi_i = \phi_i^0 + \Delta\phi_i \tag{S4.1}$$

where $\phi_i^0$ represents the basal allocation to each sector and $\Delta\phi_i$ captures changes between different conditions. As a result of the constraint that proteome fractions sum to 1, we obtain the following relation:

$$\Delta\phi_P + \Delta\phi_{NP} = \Delta\phi^{max} \tag{S4.2}$$

where the constant $\Delta\phi^{max} = 1 - \phi_{NP}^0 - \phi_{NP}^0$.

### B. Linear Growth Scaling in the (p)ppGpp Response

Previous studies have been able to operationally define proteome sectors based on correlated protein expression, varied through diverse genetic and environmental perturbations [22, 33, 53]. Interestingly, for several growth-limiting defects in nutrient assimilation and translation elongation, proteome sectors exhibit largely linear responses to changes in growth rate, each characterized by a unique slope and intercept:

$$\phi_i = \phi_i^0 + \nu_i \lambda \tag{S4.3}$$

Analogous to the translation flux argument from Scott et al. [53], such linearity arises naturally when sectors are broadly interpreted as reflecting course grained metabolic functions stoichiometric with mass growth, and the catalytic efficiency of "unperturbed" sectors is held constant [22]. As discussed previously, it is uncertain whether this condition is valid under our translation perturbations. However, our previous analysis of (p)ppGpp regulation gives us some insight into the linearity observed specifically under conditions of inhibited translation elongation.

Based on equation S3.18, we expect that $\phi_R$ should be a linear function of the growth rate. Therefore, we can similarly obtain this condition for $\phi_{NP}$, and by extension $\phi_D$, if we assume that changes in $\phi_P$ are proportional to changes in $\phi_R$:

$$\Delta\phi_R = f_R \Delta\phi_P \tag{S4.4}$$

This constraint may be easily obtained from the functional form of the (p)ppGpp repression function S3.6, previously used to model ribosome regulation, where we have substituted the ribosome concentration $R$ for the expression of an arbitrary (p)ppGpp-regulated protein $P_i$:

$$P_i(g) = \frac{k_i}{k_i + g}(P_i^{max} - P_i^0) + P_i^0 \tag{S4.5}$$

Under the condition that (p)ppGpp binding to RNAP is not strongly impacted by the specific sequence content of the promoter, i.e., that $k_i = k$ for all such (p)ppGpp-repressed promoters, it follows that expression of any two promoters $P_i$ and $P_j$ will be related as:

$$P_i = c \cdot P_j + (P_i^0 - c \cdot P_j^0)$$

where the constant $c = (P_i^{max} - P_i^0)/(P_j^{max} - P_j^0)$. In other words, the fractional abundance of all (p)ppGpp-repressed proteins should satisfy $\Delta P_i = c \cdot \Delta P_j$. Therefore changes in the fractional sums over these proteins represented by $\phi_R$ and $\phi_P$ should also be related linearly, satisfying the constraint S4.4.

We believe that the assumption that the (p)ppGpp binding constant $k$ is independent of promoter sequence is reasonable given that (p)ppGpp interacts with conserved sites on RNAP and DksA, distal to the DNA binding site, and that this interaction still occurs in the absence of promoter binding [37, 44, 45]. Critically however, we note that this independence will not be required for the validity of our overall size scaling model, as long the constraint S4.4 is obeyed empirically - which we demonstrate is the case in Extended Data Figure 11. It follows from this constraint and S4.2 that:

$$\Delta\phi_{NP}(\lambda) = \Delta\phi^{max} - f_R^{-1} \Delta\phi_R(\lambda) \tag{S4.6}$$

In this formulation, the linearity in the sector to growth function does not arise from growth as an independent variable *per se*, but rather on the regulatory action of (p)ppGpp, which provokes distortions in target genes proportional to those in r-protein expression. This interpretation aligns with previous observations: under varying (p)ppGpp conditions, growth and cell size exhibit several scaling behaviors, both when (p)ppGpp levels are synthetically manipulated [6] and in our translation perturbations affecting (p)ppGpp (Figure 5C).

### C. Simple Threshold Accumulator

To connect our proteome-partitioning model to quantitative predictions of cell size—and thus link size directly to growth—we require a plausible function linking the abundance of division proteins to cell size. Currently, the most well-supported quantitative model for the action of division proteins on size is the threshold accumulator model, which posits that cell division occurs once some limiting divisome component $D$ reaches a critical abundance $D*$. While the precise identity of the limiting division protein or proteins remains ambiguous, a threshold accumulator-type mechanism is supported by the observations that 1) divisome proteins are not known to be post-translationally modified - making their abundance equivalent to their activity, 2) such a mechanism is consistent with the cell size adder i.e., the observed statistical independence of the cell added volume from birth volume, during exponential growth, and 3) mild repression/overexpression of division protein concentration produces complementary changes in size [7, 31, 57]. Adopting a threshold accumulator model, it follows that we may express $D*$ in terms of our coarse-grained division sector:

$$D* = f_D \phi_{NP} \cdot V_D$$

where $V_D$ is the volume of the cell at division and $f_D$ is the fraction of the $\phi_{NP}$ made up of $\phi_D$. The resulting relationship between division size and the division sector is thus:

$$V_D = V_{D,0} \cdot \frac{\phi_{NP}^{max}}{\phi_{NP}}$$

with $V_{D,0}$ being the minimum cell volume when the $\phi_{NP}$ sector is maximized (i.e. at $\phi_{NP}^{max} = \phi_{NP}^0 + \Delta\phi^{max}$). $V_D$ is related to the average cell volume during exponential growth, with $\bar{V} = \frac{3}{4} V_D$, based on the theoretical distribution of cell volumes during exponential growth [43]. We expect a similar proportionality, with a different scaling factor, to hold in the mother machine. So we may re-write the previous relation as:

$$\bar{V} = \bar{V}_0 \frac{\phi_{NP}^{max}}{\phi_{NP}} = \bar{V}_0 \frac{\phi_{NP}^{max}}{\phi_{NP}^0 + \Delta\phi_{NP}}$$

Substituting $\Delta\phi_{NP}(\Delta\phi_R)$ S4.6 into our equation for $\bar{V}$ and rearranging yields:

$$\bar{V} = \bar{V}_0 \left(1 - \theta \cdot \frac{\Delta\phi_R}{\phi_R^{max}}\right)^{-1} \tag{S4.7}$$

where $\theta = (f_R^{-1} \phi_R^{max}) / \Delta\phi^{max}$ and $\phi_R^{max} = \phi_R^0 + f_R \Delta\phi^{max}$. That is, as expected, cell volume exhibits an inverse proportionality to the max-normalized change in the $\phi_R$ sector, scaled according to the degree that this change distorts the $\phi_{NP}$ sector by $\theta$.

#### D. Condition-specific Size Scaling

Recall that, according to our (p)ppGpp regulatory model, we expect perturbations to translation to yield three distinct ribosome-growth scaling behaviors. Specifically, inhibiting tRNA charging, translation elongation or translation initiation provokes scaling behaviors equivalent to the following specifications of $\Delta\phi_R$:

$$\begin{aligned}
\Delta\phi_R^{Dwell} &= \beta_{Dwell}\lambda \quad \beta_{Dwell} > 0 \\
\Delta\phi_R^{Init} &= \Delta\phi_R^{ref} \\
\Delta\phi_R^{Elong} &= (\phi_R^{max} - \phi_R^0) + \beta_{Elong}\lambda \quad \beta_{Elong} < 0
\end{aligned}$$

Therefore, we may substitute these terms into $\bar{V}(\Delta\phi_R)$ S4.7 to yield a function $\bar{V}(\lambda)$ describing the resulting size-growth scaling behavior in each condition. This results in the following perturbation-specific size scaling behaviors:

$$\bar{V}_{Dwell} = \bar{V}_0 \cdot \left(1 - \theta \frac{\beta_{Dwell}}{\phi_R^{max}} \cdot \lambda\right)^{-1} \tag{S4.8}$$

$$\bar{V}_{Init} = \bar{V}_0 \cdot \left(1 - \theta \frac{\Delta\phi_R^{ref}}{\phi_R^{max}}\right)^{-1} = \bar{V}_{Ref} \tag{S4.9}$$

$$\bar{V}_{Elong} = \bar{V}_0 \cdot \left(\frac{\phi_{NP}^0}{\phi_{NP}^{max}} - \theta \frac{\beta_{Elong}}{\phi_R^{max}} \cdot \lambda\right)^{-1} \tag{S4.10}$$

### E. Fitting Model to Size Data

In practice, we fit our size scaling data on our model scalings for tRNA charge depletion and translation elongation inhibition, according to the following equivalent equations:

$$\bar{V}_{Dwell} = \bar{V}_0 \cdot \Big(1 - \kappa_{Dwell}\lambda\Big)^{-1} \tag{S4.11}$$

$$\bar{V}_{Elong} = \bar{V}_0 \cdot \Big(\alpha_{Elong} - \kappa_{Elong}\lambda\Big)^{-1} \tag{S4.12}$$

where $\alpha_{Elong} = (\phi^0_{NP}/\phi^{max}_{NP})$ and $\kappa_i = \theta(\beta_i/\phi^{max}_R)$ for both perturbation classes.

In the context of jointly fitting a model for both scaling behaviors, we can reduce the model fit to only three parameters: $V_0$, $\kappa_{Dwell}$, and $\kappa_{Elong}$ by requiring that the two curves intersect at the observed reference (i.e. no inhibition) growth rate $\lambda^{ref}$. As a result, $\alpha_{Elong}$ becomes the following function of the $\kappa_i$:

$$\alpha_{Elong} = 1 + (\kappa_{Elong} - \kappa_{Dwell})\lambda^{ref}$$

leading us to only have to fit three free parameters to specify our size scaling data, while fixing $\lambda^{ref}$ based on our control non-targeting sgRNAs.

We may use a similar intersection condition to obtain the relationship between our fitted size parameters and the form of ribosome growth scaling. From the condition that $\Delta\phi^{tRNA}_R(\lambda = \lambda^{ref}) = \Delta\phi^{Elong}_R(\lambda = \lambda^{ref})$, it follows that:

$$\beta_{Dwell} = \frac{\phi^{max}_R - \phi^0_R}{\lambda^{ref}} + \beta_{Elong}$$

rearranging and taking advantage of the identity $\beta_{Dwell}/\beta_{Elong} = \kappa_{Dwell}/\kappa_{Elong}$, we get the following fitted value for $\beta_{Dwell}$:

$$\beta_{Dwell} = \frac{\phi^{max}_R - \phi^0_R}{\lambda^{ref}}\Big(1 - \frac{\kappa_{Elong}}{\kappa_{Dwell}}\Big)^{-1} \tag{S4.13}$$

We can similarly obtain $\beta_{Elong}$ from the same equivalence to the $\kappa_{Dwell}/\kappa_{Elong}$ ratio:

$$\beta_{Elong} = \frac{\phi^{max}_R - \phi^0_R}{\lambda^{ref}}\Big(\frac{\kappa_{Dwell}}{\kappa_{Elong}} - 1\Big)^{-1} \tag{S4.14}$$

Therefore, once we have obtained our $\kappa$ values from our fitted size scaling model, we may convert them into $\beta$ slopes specifying the ribosome level to growth relationship. Note that for this we must specify values for $\phi^{max}_R$ and $\phi^0_R$ which we obtain empirically from our ribosome scaling data.

### F. Comparison to Nutrient Growth Law

We interpret $\bar{V}_{Dwell}$ as expressing the typical size growth dependence under conditions of decreasing nutrient quality - which has been shown to similarly result in lower elongation rates and heightened (p)ppGpp levels [10, 41]. Compare this to the classic form of size scaling law of Schaechter, Maaløe and Kjeldgaard [49]:

$$\bar{V}_{SMK} = \bar{V}_0 2^{(\lambda/\lambda_0)}$$

where $\lambda_0 = 1$ doubling/hr. While the $\bar{V}_{Dwell}$ S4.12 function is not obviously similar to the exponential form specified by the classic growth law, it in fact behaves roughly exponentially under typical experimental conditions - to see this we can examine the Maclaurin series expansion of $\log \bar{V}_{Dwell}$:

$$\begin{aligned}\log \bar{V}_{Dwell} &= \log \bar{V}_0 - \log(1 - \kappa_{Dwell}\lambda) \\ &= \log \bar{V}_0 + \kappa_{Dwell}\lambda + \frac{(\kappa_{Dwell}\lambda)^2}{2} + O(\lambda^3) \\ &= \log \bar{V}_0 \sum_{n=1}^{\infty} \frac{(\kappa_{Dwell}\lambda)^n}{n}\end{aligned}$$

Exponentiating $\log \bar{V}_{Dwell}$ results in a functional form that is equivalent to the classic growth law, up to a first-order approximation:

$$\bar{V}_{Dwell} = \bar{V}_0 exp(\kappa_{Dwell}\lambda) \cdot exp(\frac{(\kappa_{Dwell}\lambda)^2}{2} + O(\lambda^3)) \quad \text{(S4.15)}$$

As such, for relatively small deviations in growth rate, second-order and higher terms will become insignificant - yielding the observed exponential scaling.

### 5. PPGPP-GROWTH SCALING

#### A. Scaling Under Nutrient Limitation

To complement our examination of $R$ to $\lambda$ scaling, under various types of translation inhibition, we conclude with a short discussion of the predicted scaling of (p)ppGpp ($g$) with $\lambda$ under each of these conditions. Beginning with the condition of nutrient/tRNA charge limitation, [60] produced predictions for the resulting (p)ppGpp to growth rate relationship, which we reproduce here. First, simple substitution of equation S3.2 into the regulatory function $R(g) = a/g$ results in:

$$R = c \cdot R_0 \cdot \frac{h-1}{g}$$

It follows that in the limit $\lambda \to 0$ we have $g_0 = c(h-1)$, where $g_0$ is the (p)ppGpp concentration at a growth rate of zero. Substituting this back into the previous relationship we obtain:

$$R = R_0 \frac{g_0}{g_D} \quad \text{(S5.1)}$$

If we substitute equation S5.1 into equation S3.4 we obtain the relationship between growth rate and (p)ppGpp concentration predicted by the model:

$$g_D(\lambda) = \frac{g_0}{1 + \lambda/(\varepsilon_{max} R_0)} \quad \text{(S5.2)}$$

#### B. Scaling Under Slow Translocation

Similarly, our model produces a prediction for the relationship between the growth rate and (p)ppGpp concentration. To obtain this we first rearrange equation S3.8:

$$R = \frac{c\tau_D \lambda}{g} + R_0$$

If we then substitute this into equation S3.9 and rearrange, we may produce the following relation between growth rate and (p)ppGpp:

$$\frac{kg_T}{k + g_T} = \frac{c\tau_D \lambda}{R_{max} - R_0}$$

Since k and g are positive, it follows that the left-hand size of the equation is positive and monotonic with increasing g, implying that the relationship between growth rate and g is indeed positive, as expected. Further rearrangement yields our final relation between growth rate and (p)ppGpp:

$$g_T(\lambda) = \frac{\theta}{(1/\lambda) - (\theta/k)} \quad , \quad \theta = \frac{c\tau_D}{R_{max} - R_0} \quad \text{(S5.3)}$$

We may compare the behaviors of (p)ppGpp concentration with respect to growth rate under either restricted tRNA charge or slow elongation using equations S5.1 and S5.3. To do so, we first need to specify model parameters and then set reference values for our initial growth condition, as was done for the $R$ vs $\lambda$ model comparison. Like this previous comparison, we specify $R_0 = 0.03$, $R_{max} = 0.793$ and $\varepsilon_{max} = 72000 a.a./hr$. We will also set $k$ based on our model fit of equation S3.6 to data from [60]. We may then use the reference state $\lambda = 2$ to recover the values of $g_0$ as well as $c\tau_D$ according to the following procedure. From our reference state we have that $R(\lambda = 2) = 0.3$, which we substitute into equation S3.9 resulting in an estimate of $c\tau_D$:

$$c\tau_D = 0.082$$

further substituting this into equation S5.3 produces our curve for (p)ppGpp concentration under slow translocation:

$$\theta = \frac{0.082}{R_{max} - R_0} \approx 0.11 \quad , \quad g_T(\lambda) = \frac{0.11}{(1/\lambda) - (0.11/k)} \tag{S5.4}$$

From this we can compute a consistent reference value for $g_0$ in the restricted tRNA charge model by computing the value of $g_T(\lambda = 2)$ according to equation S5.4 and then use it to set $g_D(\lambda = 2)$ equal to this value:

$$g_T(\lambda = 2) = g_{ref} = \frac{0.11}{(1/2) - (0.11/k)} \approx 0.61$$

Rearranging equation S5.1 and plugging in values for $\varepsilon_{max}$, $R_0$, and $g_{ref}$ we may compute a reference value for $g_0$:

$$g_0 = (1 + \frac{\lambda}{\varepsilon_{max} R_0}) g_{ref}$$

Now, we may examine the curves for equations S5.1 and S5.3, consistently estimated and baselined, which are overlaid with a flat line corresponding to the case of reduced initiation discussed previously (Figure S4). Interestingly, the model exhibits a roughly 3-fold dynamic range in (p)ppGpp concentration under nutrient limitation, though empirically we observe variability on the order of 15-fold (Figure 6B). As such, while the model is broadly predictive of cellular ribosome content, it appears to be unable to quantitatively capture changes in (p)ppGpp concentration. Further development of this model, likely also accounting for the apparent timescale separation of RelA and SpoT activities discussed in [60], will be necessary before such a quantitative prediction is possible.

### 6. THE CONSEQUENCES OF RELA DELETION

The growth model above, centered around the regulation of (p)ppGpp levels according to equation S2.3,provides an internally consistent account for the diversity of scaling behaviors that we documented in WT cells. In Δ*relA* cells, however, it is not clear that this model can account for the observed inversion of both size and ribosome scaling, under conditions of tRNA synthetase depletion. In this final section, we identify this phenotype with the previously characterized "relaxed" response to amino acid depletion, and propose modifications of our main (p)ppGpp regulatory scheme that better capture the observed behaviors. Specifically, we consider the additional effects of ribosome queueing as a possible explanation for the relaxed response and propose that RelA responds non-linearly to codon-specific dwell times to counteract this effect in WT cells.

#### A. (p)ppGpp Regulatory Function in Δ*relA*

The simple model of (p)ppGpp regulation specified by equation S2.1 does not fully account for observations in a Δ*relA* strain, where despite the absence of RelA-dependent (p)ppGpp synthesis, (p)ppGpp levels do not drop to 0. Rather, the residual synthesis activity of SpoT appears to account for the remaining (p)ppGpp, which for simplicity we model as a fixed production rate, $c* = bc$ where $c$ is the same as in equation S2.1 and is scaled by an arbitrary constant $b$. Including this term, equation S2.1 can be rewritten:

$$g(R_D, R_T) = c\frac{R_D + b}{R_T} \tag{S6.1}$$

In a Δ*relA* strain, production of (p)ppGpp by dwelling ribosomes is eliminated resulting in:

$$g_{\Delta relA}(R_T) = c\frac{b}{R_T} = \frac{c*}{R_T} \tag{S6.2}$$

making (p)ppGpp levels strictly dependent on the concentration of translocating ribosomes $R_T$. For simplicity, we neglect the contribution of the extra $b$ term in WT cells. We note that, however, SpoT-dependent (p)ppGpp synthesis likely contributes to $\sim 50\%$ of total (p)ppGpp in WT cells, as the concentration of (p)ppGpp in $\Delta relA$ is roughly half that of WT (Figure 6B).

While naively it appears that $g_{\Delta relA}$ is no longer dependent on $R_D$, considering that ribosome elongation still obeys detail balance between the dwelling and translocating states, (p)ppGpp levels should still be sensitive to the ratio of $\tau_D$ and $\tau_T$. We can see this if we decompose the active ribosome pool into $R_T$ and $R_D$ components:

$$R_{active} = R_T + R_D$$

substituting equation S2.2 into this, we see that $R_T$ can be expressed in terms of the active ribosome fraction $R_{active}$, $\tau_D$, and $\tau_T$:

$$R_T = R_{active} \frac{1}{\tau_D/\tau_T + 1}$$

Substituting this into equation S6.2 we arrive at the following relation:

$$g_{\Delta relA}(\tau_D, \tau_T) = c * R_{active}^{-1}(1 + \frac{\tau_D}{\tau_T}) \tag{S6.3}$$

demonstrating that $g_{\Delta relA}$ retains the positive-linear dependence on $\frac{\tau_D}{\tau_T}$ as in $g_{WT}$.

Comparing this to our data, we find that the reduction in (p)ppGpp accompanying reduced elongation is not affected by the deletion of *relA* (Figure 6B), indicating that equation S6.3 is at least qualitatively consistent with the expected impact of low elongation (increased $\tau_T$). As we discuss in the main text, the lack of any other known (p)ppGpp hydrolase or synthase in *E. coli* implies that $R_T$-associated (p)ppGpp regulatory activity must be a result of *spoT*.

### B. Ribosome Queuing as a Mechanism for the Relaxed Phenotype

It is less clear why we observe that in Δ*relA* cells, single tRNA synthetase knockdowns result in both elevated size and RNA/protein ratios, suggesting diminished (p)ppGpp (Figure 6B,C). This puzzling decrease in (p)ppGpp has been previously described as the so-called "relaxed" response, which can also be provoked by depleting cells for single amino acids, which results in the similar consequences for tRNA charge [36]. The forms of equations $g_{WT}$ S2.3 and $g_{\Delta relA}$ S6.3 would suggest that that both WT and Δ*relA* cells should exhibit a positive dependence of (p)ppGpp with increasing $\tau_D$, and presumably increase under conditions of diminished tRNA charge. This is the simple interpretation that we focus on in the main text.

However, this means that the "relaxed" response exhibited in Δ*relA* is not explicitly accounted for by our form for $g_{\Delta relA}$ without additional considerations. One hint lies in the fact that, while single amino acid deprivation decreases (p)ppGpp in Δ*relA* cells, instead withdrawing multiple amino acids results in a more typical induction of (p)ppGp synthesis [36]. Indeed, we also observe that blocking amino-acid synthesis more broadly, e.g. by knockdowns targeting glycolysis, results in decreased size even in Δ*relA* cells (Figure 6C).

A major difference between these conditions lies in the expected impact on the *codon-specific* $\tau_D$ - withdrawing a single amino acid results in the appearance of so-called "hungry" codons, with long $\tau_D$. To gain an intuition for how ribosome stalling on hungry codons will impact the overall rate of protein synthesis, we break down $\tau_D$ into the average dwelling time of codons specifying each of the 20 amino acids. For simplicity, we will assume even codon usage among all amino acids:

$$\tau_D = \frac{\sum_{n=20} \tau_{D,i}}{20} \tag{S6.4}$$

where $\tau_{D,i}$ is the average dwell time for codons of a specific amino acids. Assuming that all codons start with the same dwell time, we will have:

$$\tau_D^0 = \frac{\sum_{n=20} \tau_D^0}{20}$$

Now, consider the consequences of withdrawing one of the 20 amino acids to the point where the elongation rate drops by 50%. Breaking down equation S6.4 into 19 amino acids we will hold at a constant dwell time $\tau_D^0$ and 1 which we will knockdown to some lower dwell time $\tau_D^{KD}$, we have:

$$\tau_D^f = \frac{19\tau_D^0}{20} + \frac{\tau_{D,KD}}{20} \approx \tau_D^0 + \frac{\tau_{D,KD}}{20}$$

where we rounded the first term from 0.95 to 1 for simplicity. Now, considering that we inhibit elongation by 50% it follows that $\tau_D^f = 2\tau_D^0$ and, from the previous relation:

$$20\tau_D^0 = \tau_{D,KD}$$

Therefore, in this simple scenario, though elongation and growth only drop by 50%, and the average dwell time only doubles, there is a 20-fold increase in the dwell time of codons corresponding to the amino-acid being depleted.

Importantly, the long-lived stalling events induced by a hungry codon not only affect the kinetics of the dwelling ribosome, but also promote queueing of upstream ribosomes, which has been observed in several ribosome profiling studies [28, 56]. Critically most of the ribosomes in these queues will be locked in the translocating state, resulting in an increase in $\tau_T$. Therefore, according to our equation for $g_{\Delta relA}$ S6.3, we may expect a decrease in (p)ppGpp under single tRNA synthetase knockdown due to this elevated $\tau_T$.

Here, in lieu of the stochastic simulations typically used to study queueing, we will introduce a simple toy model to understand the effects of queuing on $\tau_T$ and $\tau_D$. Once again assuming that the codons corresponding to all 20 amino acids are balanced, we will model the behavior of a single hungry codon with an upstream ribosome queue of length $Q$. Instead of modeling the transcript and context-dependent ribosome flux into the queue, we will simply set the queueing rate to be the initiation rate of translation $I$, which is the main context-independent determinant of the queueing rate. Treating this situation as a simple Markovian M/M/1 queue, the probability of there being $q$ ribosomes in the queue will be:

$$P(Q = q) = (1 - \rho)\rho^i$$

where $\rho = I \cdot \tau_{D,KD}$. It is known that ribosome rescue pathways are strongly induced by the presence of ribosome queues, and as such most queues are of length one [56]. For simplicity, we will treat the effects of ribosome rescue as effectively truncating the queue distribution to a length of one, where the probability of queue lengths of two or greater are reassigned to a length of one. As a result the revised probability of our queue will be:

$$P(Q = q) = \begin{cases} (1 - \rho) & q = 0 \\ \rho & q = 1 \end{cases}$$

Finally, we will assume that, for every dwelling ribosome there are another 19 ribosomes that are either in the queue or freely translating, reflecting the proportion of unaffected codons in the system. Under these assumptions, we will have the following formulas for the average $\tau_D$ and $\tau T$:

$$\tau_{T,Q} = \begin{cases} ((19 + (1 - \rho))\tau_T^0 + \rho\tau_{D,KD})/20 & \rho < 1 \\ (19\tau_T^0 + \tau_{D,KD})/20 & \rho \geq 1 \end{cases} \tag{S6.5}$$

$$\tau_{D,Q} = (19\tau_D^0 + \tau_{D,KD})/20 \tag{S6.6}$$

In order to compare this to an equivalent model without a ribosome queue, we will use the same set of assumptions with a queue size of zero, resulting in average $\tau_D$ and $\tau_T$ of:

$$\tau_{T,NQ} = \tau_T^0 \tag{S6.7}$$

$$\tau_{D,NQ} = (19\tau_D^0 + \tau_{D,KD})/20 \tag{S6.8}$$

Unlike our previous model, we will not attempt to model the regulatory effect of (p)ppGpp on the ribosome concentration and will simply set growth based on the some fixed initial ribosome fraction, according to equation S1.9:

$$\lambda = \phi_R^{init} \cdot (\varepsilon(\tau_{D,KD})/m_R)$$

Comparing the scaling of $\tau_{T,Q}$ (Figure S7A ,orange) and $\tau_{T,NQ}$ (Figure S7A ,light orange) with the growth rate, while varying $\tau_{D,KD}$, we see that as growth rate decreases the $\tau_{T,Q}$ is increasingly dominated by the effects of ribosome queueing. Moreover this effect can be quite significant under experimentally relevant decreases in growth on the order of 50%. We note that the effects of queueing exhibit a sharp change in behavior at $\rho = I \cdot \tau_{D,KD} \geq 1$, which is due to the previously described clipping of our queueing model to ribosome queue sizes of at most one.

The consequences of this increase in $\tau_{T,Q}$ for (p)ppGpp regulation are quite significant. Comparing the resulting behavior of our (p)ppGpp regulatory function S2.3, we see that the additional consideration of queueing leads to an inversion in the scaling behavior of (p)ppGpp with growth

**Fig. S7. Effects of Ribosome Queueing on (p)ppGpp Regulatory Model.** A) Average translocation time vs growth rate, when varying dwelling time on a hungry codon. Lines corresponding to cases with (orange) and without queueing (light orange) considered are shown. Grey shaded area corresponds to values of $\rho$ greater than or equal to 1. Model parameters are $\tau_D^0 = 1/25\,s^{-1}$, $\tau_T^0 = 1/100\,s^{-1}$, $phi_R^{init} = 0.2$ and $I = 1/2\,s^{-1}$. B) Relative (p)ppGpp levels vs growth rate, for different (p)ppGpp regulatory models. Lines correspond to models with linear (p)ppGpp synthesis, with respect to dwell time, either with (orange) or without (light orange) queueing. An additional line corresponds to a model with squared (p)ppGpp synthesis, with respect to dwell time, and queueing (green). Same model parameters as in panel A.

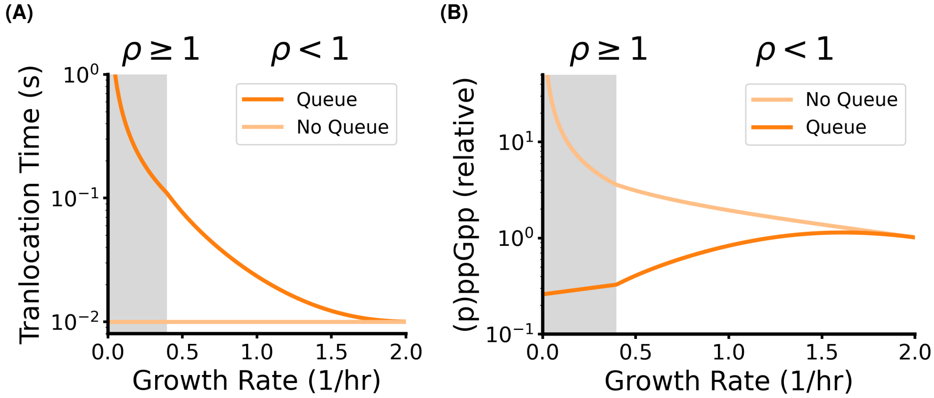

under varying $\tau_{D,KD}$ (Figure S7B ,orange). In other words, the increase in translocation time owing to ribosome queueing, under conditions of single amino acid depletion, results in a drop in (p)ppGpp levels. The drop occurs as long as $\tau_T^0 < \tau_D^0$, which is generally the case [50]. We can see when we plug this model into equation S2.3 for $\rho \geq 1$:

$$g_Q(\tau_{D,KD}) = g_0\left(\frac{\tau_T^0}{\tau_D^0}\right)\left(\frac{19\tau_D^0 + \tau_{D,KD}}{19\tau_T^0 + \tau_{D,KD}}\right) \tag{S6.9}$$

As $\tau_{D,KD} \to \infty$, $g_Q \to g_0(\tau_T^0/\tau_D^0)$, which will be less than $g_0$ for $\tau_T^0 < \tau_D^0$. Notably, while this is true for the single ribosome queue considered here, admitting longer queues up to length $q$ will result in the average translocation time tending towards $(20-q)\tau_T^0 + q\tau_{D,KD}$ resulting in $g_Q \to g_0(\tau_T^0/q\tau_D^0)$, meaning that a decrease would be expected in this more general case for $\tau_T^0 < q\tau_D^0$.

Therefore, with the added consideration of ribosome queueing, both our WT $g_{WT}$ S2.3 and $\Delta relA$ $g_{\Delta relA}$ (p)ppGpp regulatory models predict a relaxed response i.e., a decrease in (p)ppGpp at low growth rates under conditions of single amino acid depletion. Moreover, it is expected that knocking down amino acid levels more evenly, which does not strongly impact $\tau_T$ due to a lack of queueing, will provoke increased (p)ppGpp consistent with both our (Figure 6B) and previously published observations [36]. This is also consistent with ribosome growth scaling being preserved in Δ*relA* strains [55], as varying growth media will generally modulate the flux of multiple amino acids.

What remains unresolved, however, is why in WT strains do we not observe the same relaxed response under tRNA synthetase knockdown, as suggested by the dependence of $g_{WT}$ S2.3 on the $\tau_D/\tau_T$ ratio. One answer may be that RelA activity does not linearly scale with the average dwelling time $\tau_D$. To see why a non-linear RelA response would in fact be the expectation, consider the RelA activation mechanism. It is known that RelA is activated through a bimolecular interaction with dwelling, uncharged ribosomes and uncharged tRNAs. Critically, recent studies have suggested that RelA activation only occurs if the dwelling ribosome is paused on a codon *that is complementary* to the anti-codon of the uncharged tRNA [40, 59]. As such, under the scenario where charging of a single tRNA becomes limiting for growth, the fraction of RelA allocated to that uncharged tRNA becomes:

$$[RelA]_{KD} = \frac{t_{KD,U}}{\sum_{i\in 1,\ldots,19} t_{i,U} + t_{KD,U}}[RelA]_{tot} \tag{S6.10}$$

where uncharged tRNAs of other types with concentrations $t_{i,U}$ effectively compete with the concentration of uncharged target tRNA $t_{KD,U}$ for the total RelA population $[RelA]_{tot}$.

At this point, it is not entirely clear that there is a simple way to relate this form to the translation kinetic parameters of our (p)ppGpp control model. However, if we assume that single amino acid depletions result in uncharged tRNA levels well below the the Km of the cognate synthetase, we may approximate charging kinetics as a zero-order reaction - an approach introduced previously in Elf and Ehrenberg 2005 [13].

$$v_{KD} = k_{cat}e_{KD}$$

where $k_{cat}$ is the catalytic rate of the cognate tRNA synthetase and $e_{KD}$ is the concentration of that synthetase. At the same time, all other tRNAs are should be charged according to typical Michaelis–Menten kinetics:

$$v_i = \frac{k_{cat}e_0 t_{U,i}}{Km + t_{U,i}}$$

Where we take $k_{cat}$ as the same across all synthetases - though $k_{cat}$ generally ranges from 1 to 100 $s^{-1}$ across synthetases [17]. At steady state, $v_{KD} = v_i = \varepsilon/20$, assuming that translation consumes amino acids roughly equally, leading to the following form for the elongation rate:

$$\varepsilon = 20k_{cat}e_0 \frac{t_{U,i}}{Km + t_{U,i}}$$

Identifying $\varepsilon \to \varepsilon_0$ as the maximum elongation rate when $t_{U,i} \gg Km$, this relationship can be rearranged as:

$$t_{U,i} = \frac{Km\varepsilon}{\varepsilon_0 - \varepsilon}$$

This finally allows us to write our equation S6.10 for the active RelA concentration allocated to the knocked down tRNA, in terms of the elongation rate:

$$[RelA]_{KD} = \frac{t_{KD}^{tot}(\varepsilon_0 - \varepsilon)}{19Km\varepsilon + t_{KD}^{tot}(\varepsilon_0 - \varepsilon)}[RelA]_{tot} \tag{S6.11}$$

Where we set $t_{KD,U} = t_{KD,tot}$ i.e., the total concentration of the target tRNA - reasoning that nearly all tRNAs charged by the target synthetase will be uncharged in this zero-order kinetic scenario. We can see that this function exhibits significant nonlinearity across plausible parameter regimes (Figure S7A).

Conveniently, this expression simplifies significantly if $t_{KD,tot} = 19\mu M$, which is reasonably centered in the empirical range of 2-30$\mu M$ [12]:

$$[RelA]_{KD} = \frac{\varepsilon_0 - \varepsilon}{((Km - 1)\varepsilon) - \varepsilon_0}[RelA]_{tot}$$

We can simplify this even further, if we similarly take a middling value of $Km = 1\mu M$ for the tRNA synthetase where $Km$ ranges from 0.1 to 2 [25]:

$$[RelA]_{KD} = \left(1 - \frac{\varepsilon}{\varepsilon_0}\right)[RelA]_{tot} \tag{S6.12}$$

Consider in turn how this impacts the functional form of (p)ppGpp regulation. Again, RelA goes by bimolecular interactions between codons and cognate tRNAs, so the synthesis of (p)ppGpp will be:

$$\left(\frac{dg}{dt}\right)_{KD} = k_1 \cdot [RelA]_{KD}R_D^{KD} - k_2 R_T g$$

where $g$ once again denotes the (p)ppGpp concentration. At steady state this results in a similar form as our original (p)ppGpp regulatory model:

$$g_{KD} = c \cdot \frac{[RelA]_{KD}R_D^{KD}}{R_T}$$

The fraction of dwelling ribosomes on the limiting codon will be in proportion to the dwell time on that codon according to the following relation:

$$R_D^{KD} = R_D \cdot \frac{\tau_{D,KD}}{19\tau_D^0 + \tau_{D,KD}}$$

Substituting this into our previous equation, we may obtain an expression for ppGpp in terms of kinetic parameters:

$$g_{KD} = c \cdot \frac{R_D}{R_T}(1 - \frac{\varepsilon}{\varepsilon_0})(\frac{\tau_{D,KD}}{19\tau_D^0 + \tau_{D,KD}}) = c \cdot \frac{\tau_D}{\tau_T}(1 - \frac{\varepsilon}{\varepsilon_0})(\frac{\tau_{D,KD}}{19\tau_D^0 + \tau_{D,KD}})$$

If we once again allow for ribosome dwelling, and therefore take the average dwelling time to follow equation S6.6, we can obtain this equation in terms of the dwell time of the limiting codon:

$$g_{KD} = c * \cdot \frac{\tau_{D,KD}}{\tau_T}(1 - \frac{\varepsilon}{\varepsilon_0}) \quad \text{(S6.13)}$$

where we have absorbed additional constant terms into the parameter $c$, resulting in $c*$.

**Fig. S8. More Realistic RelA Activity Restores WT-like Behavior to Queueing Model.** A) Effective RelA concentration allocated to a growth-limiting tRNA vs the resultant elongation rate. Lines describe this trend for different total concentrations in this limiting tRNA, noting that in our model we do not distinguish between tRNA isoacceptors for simplicity. In practice, these values range from 4.88 $\mu m$ for to 44.38$\mu m$ for Leucine tRNAs at fast growth [12]. B) Semi-log plot of the predicted (p)ppGpp vs growth rate dependence for our queueing model with (orange line) and without (blue line) the proposed RelA activity. Same parameters as in Figure S7B. As before, grey shaded area corresponds to values of $\rho$ greater than or equal to 1.

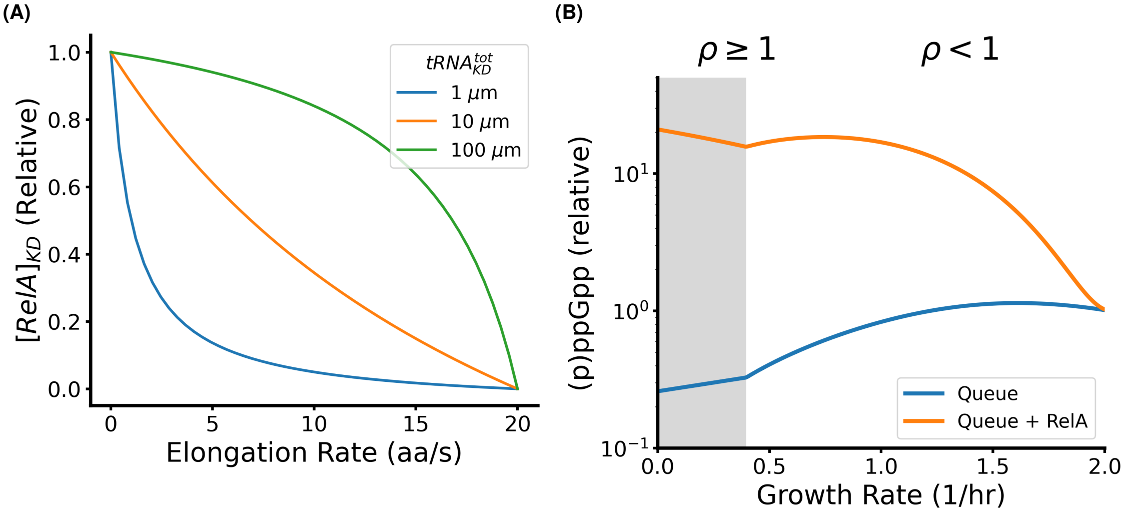

Compare the growth vs (p)ppGpp behavior of our more realistic model for RelA activation (Figure S8B, orange), represented by equation S6.13, to our previous model (Figure S8B, blue) - which only incorporated a simple, first-order relation between RelA and the dwelling time. It is apparent that this updated model recovers the negative scaling of (p)ppGpp with growth, under limiting tRNA charge for a single amino acid - while the first-order model does not. That is, this more realistic RelA activation model counters the effects of increased translocation time from queueing and qualitatively reproduces the empirical scaling of (p)ppGpp in WT cells. Elimination of this non-linear dependence in $\Delta relA$ is consistent with the "relaxed" phenotype in these strains, which we have seen can be explained by queueing effects.

### 7. CONCLUSION

In summary, we reviewed the ribosome regulatory model introduced in Scott et al. 2010 [53], which accurately accounts for ribosome-growth scaling under conditions of varying nutrient quality and elongation rate. However, we found that it could not account for our observation of mostly invariant ribosome content under conditions of reduced translation initiation. We determined that the more recent model of Wu and Balakrishnan et al. 2022 [60] accurately predicts this behavior. Further, we extended this model to show that it can also account for the ribosome-growth scalings previously observed in [53]. We showed that this model predicts a linear relationship between a medium's nutrient quality $\kappa_n$ and the ribosome content $R$ it supports without elongation inhibition, which is largely borne out by the data in [53]. Overall, our data are consistent with a model in which translocating ribosomes promote (p)ppGpp degradation, as proposed in [60].

We prospectively attribute this activity to SpoT, primarily due to strong elongation-dependent (p)ppGpp regulation in a Δ*relA* strain, where SpoT is the only remaining (p)ppGpp synthase/hydrolase. The mechanism mediating this regulation remains unclear, since SpoT does not bind directly to elongating ribosomes, though there is some evidence for its association with a pre-50S particle on sucrose gradients [24]. As such, a mechanism which measures elongation by proxy, for instance through the rate of ribosome stalling and rescue, appears to be more likely at this time. Future biochemical work will thus be needed to evaluate these possibilities.

We noted that this model does not provide a full accounting for the phenotypic differences between WT and Δ*relA* strains. We speculated that these differences can be resolved through the addition of two additional elements: ribosome queueing on hungry codons stimulating SpoT and RelA activation via only cognate uncharged-tRNA-codon binding. Direct experimental validation of these hypotheses is therefore a promising direction for future work.
